# Using conditional Generative Adversarial Networks (GAN) to generate *de novo* synthetic cell nuclei for training machine learning-based image segmentation

**DOI:** 10.1101/2022.11.12.516283

**Authors:** Mehmet Ilyas Cosacak, Caghan Kizil

**Author notes:** Twitter: @micosacak (Mehmet Ilyas Cosacak).

## Abstract

Generating masks on training data for augmenting machine learning is one of the challenges as it is time-consuming when performed manually. While variable random images can be generated by Generative Adversarial Networks, an image-to-image translation is needed to generate both images and ground truth data. To generate cells and their corresponding masks, we used a new approach to prepare the training data by adding masks on 4 different channels preventing any overlapping between masks on the same channel at an exactly 2-pixel distance. We used GAN to generate nuclei from only two images (415 and 435 nuclei) and tested different GANs with alternating activation functions and kernel sizes. Here, we provide the proof-of-principle application of GAN for image-to-image translation for cell nuclei and tested variable parameters such as kernel and filter sizes and alternating activation functions, which played important roles in GAN learning with small datasets. This approach will decrease the time required to generate versatile training datasets for various cell types and shapes with their corresponding masks for augmenting machine learning-based image segmentation.

## INTRODUCTION

Image segmentation through automated quantification of objects is one of the main challenges in machine learning. Generalizing a method that can be commonly used is still a need, even the existing ones requires generation of new training data depending on the tissue type, cell types or complexity of the image (Stringer et al., 2020; Stringer & Pachitariu, 2022). As a result, generating relevant training data is the key step in training neural networks (NNs). In the last decade, Convolutional Neural Networks (CNNs) have been successfully applied to solve image segmentation by using pixel-wise information classification. Currently, U-Net is a widely used method among the state-of-the-art models for image segmentation (Ronneberger et al., 2015). The U-Net architecture is composed of encoders (convolution layers) and decoders (deconvolution layers) with skip connections between encoder and decoders. Many models have been developed based on U-Net architecture: UNet++ (Zhou et al., 2018), UNet3+ (Huang et al., 2020), DC-UNet (Lou et al., 2021), MultiResUNet (Ibtehaz & Rahman, 2019), and Half-UNet (Lu et al., 2022). In almost all of these models, the number of parameters was decreased without compromising the accuracy of segmentations. Additionally, some of the U-Net-derived models have already been used in image segmentation, such as StarDist (Schmidt et al., 2018) and Cellpose (Stringer et al., 2020; Stringer & Pachitariu, 2022). However, despite the existing U-Net architectures, there is still room to generate more models derived from U-Net. For instance, in addition of the U-Net architectures aforementioned, chaining of U-Nets (Alom et al., 2018; Laibacher et al., 2018; Zhuang, 2018) is another option to generate different U-Net architectures. These models can be adapted to various datasets and every one of them may perform significantly better than other variants depending on the training datasets used. While most of these models have been tested for image segmentation, these models may be further used as generative models for GANs. Therefore, in our current study, in addition to classical U-Net architecture, we improved the U-Net architecture by generating 3x-U-Net and evaluated its performance by comparing it with classical 4, and 6 skip-connected U-Nets.

Training data, besides the architecture of the Neural Networks, is a key step for image segmentation. Training data generation requires manual segmentation to construct the ground truth data, which is labor-intensive. Evaluation of most models is performed on unseen images, for which the test images are not used for training the models. As a result, the generation of the unseen image is also needed to evaluate the predictive performance of models on diverse image datasets. Generative Adversarial Networks (GANs), especially conditional GANs (Isola et al., 2016, 2017a), can be used to generate data for image augmentation, either to use as training data or as test data for evaluation. While GANs can be used to generate random realistic images (Goldsborough et al., 2017; Han et al., 2018; Mannam et al., 2021; Osokin et al., 2017), conditional GANs can be used to generate segmented cells (masks) and their images (Baniukiewicz et al., 2019). In the latter study, single cells and their intracellular structure were generated. However, to our knowledge, there is no study that generates microscopic cell images from their masks, especially if the masks are crowded and overlapping. Here, we propose a new way of generating masks for cell nuclei and distributing these masks on 4 different channels to prevent any overlapping between any adjacent masks on the same channel. Our training data is composed of 850 cells from 2 images, augmented by generating small images from the bigger images. To generate a model that can perform better image-to-image translation, we used U-Net and concatenated U-Nets to generate 2x and 3x U-Nets. Thus, in this study, we are proposing a new and quick strategy for the segmented mask to target image generation, so that U-Nets can perform better than regular U-Net architecture. Our approach is a proof-of-principle for surrogate realistic microscopic image data generation that can be used as synthetic test images or training image datasets. We believe that this approach can be further developed to generate irregular shapes, which are common for various cell types such as astroglia or microglia.

## MATERIAL AND METHODS

### Generating Masks and Training Images

We cropped two regions (629×1014 and 527×1383 pixels) (**Fig. S1**) from an image of a zebrafish (20X objective) telencephalon and generated masks for each DAPI staining (hereafter cell nucleus). Then, we arranged all masks on 4 channels to prevent any overlapping masks on the same channel (**Fig. 1**). Thus, we randomly inserted a mask on the first channel and located the remaining masks one by one as follows; if the mask overlapped with the masks on the first channel, we tried to insert it on the second channel, otherwise, we tried to locate it on the third or fourth channel. We repeated this until all masks were stacked on all layers. If any masks could not be fitted on a channel, we randomized the masks and started to arrange them again as above. Based on the image regions and cell density in the two images, 4 channels were enough for the current analyses. To create small images for training, we generated images 128×128 pixels sub-images from the two images, by slicing horizontally and/or vertically 4 pixels, which generated 4110 128×128 pixel images. The masks on these images were again randomized 10 times and images were further inverted and rotated 90, 180, and 270 degrees. In total, we used 205500 images with 128-pixel sizes for training. During randomization, some images were redundant because of non-overlapping masks, thus all masks were on channels 1 and/or 2. These images were not excluded from the training data.

**Figure 1:**
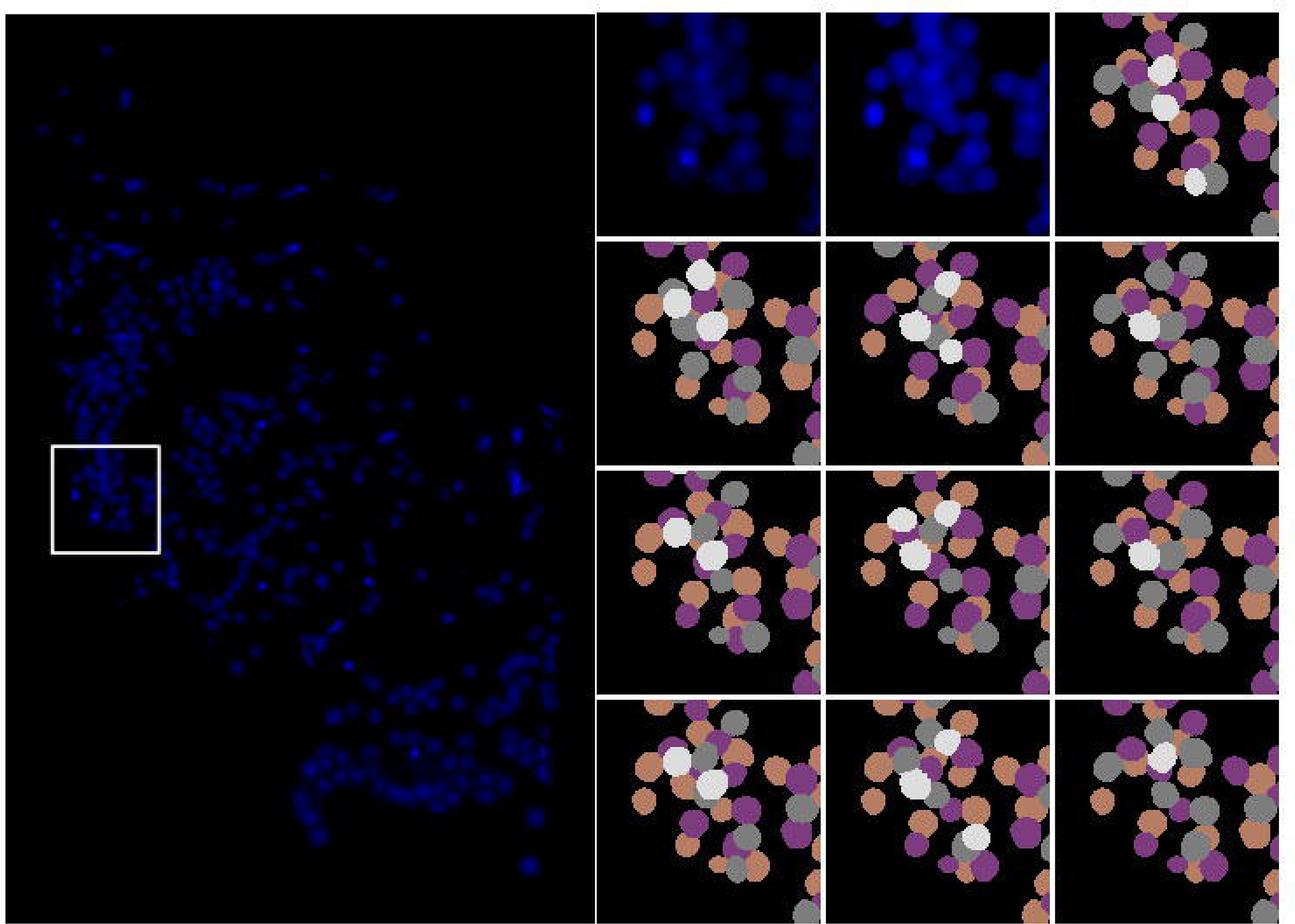
Image 1 of the 2 images used to generate masks. (A) the image with original exposure, (B) an example area from the original image with original exposure, (C) an example area from the original image with high exposure for visibility, (D) original masks generated by a human, (E-M) the masks have been randomized and 9 different masks distribution has been shown. The white masks belong to the 3^rd^ layer, and different masks have been shown each time. Each color indicates a different layer.

For mouse brain nuclei, we used two images (**Supp_Figs. 2a, 3a**) taken at 40X with relatively visible intranuclear structures. We cropped a less crowded 3451×949 image (**Supp_Fig. 2b**) area from one image (**Supp_Fig. 2a**) to be used as training data and another 1457×1049 image area from **Supp_Fig. 3a** to be used as test data (**Supp_Fig. 3b**). An additional 1024×1024 (**Supp_Fig. 3c**) pixel area from the CA3 of the hippocampus, which has a relatively high cell density area was cropped to generate another test data. We manually generated masks on each image as described above. **Supp_Fig. 2b** was used to generate 256×256 pixel sub-images by moving 8-pixel vertically and/or horizontally and 10 different images by randomizing masks were generated as described above. **Supp_Fig. 3b** was used to generate 256×256 pixels to be used as testing outputs of the models at each epoch. **Supp_Fig. 3c** was used directly to challenge models at epoch 25.

### Generative Adversarial Neural Networks

We used the implemented version of image-to-image GAN (or pix2pix) from the TensorFlow website and previous publications (Isola et al., 2017a, 2017b; Mirza & Osindero, 2014). In brief, we used 4 or 6 layers of convolution and deconvolutions (or fractionally strided convolutions) with batch normalization and dropout (**Fig. 2**). Striding the U-Net encoders resulted in 1×1 pixel at the lowest layer with 128×128 pixel images for 6 layers U-Net. We used the same U-Net and concatenated its output layer with the input layer and passed the resulting layer in a second U-Net (2x U-Net). Similarly, we concatenated the output of the 2x U-Net with the input layer and pass it to another U-Net (3x U-Net). The output layer of the first U-Net of 2x and the second U-Net of 3x U-Net were alternating between “*tanh*” or “*gelu*” activation functions. Additionally, we used 16, 32, and 64 filter sizes for 1x U-Net, while 16 filter sizes for 2x and 3x U-Net and 3, 5, and 7 kernel sizes for all models. The 64 filter-sized 1x U-Net with 7 kernel sizes has many parameters, which took hours for 1 epoch training, so we skipped these conditions for testing. We ran all analyses on V100-SXM2-32GB GPUs. Each model had a different running time per epoch, ranging from 1 min to 25 mins depending on the number of total variables, concatenations and the training data.

**Figure 2:**
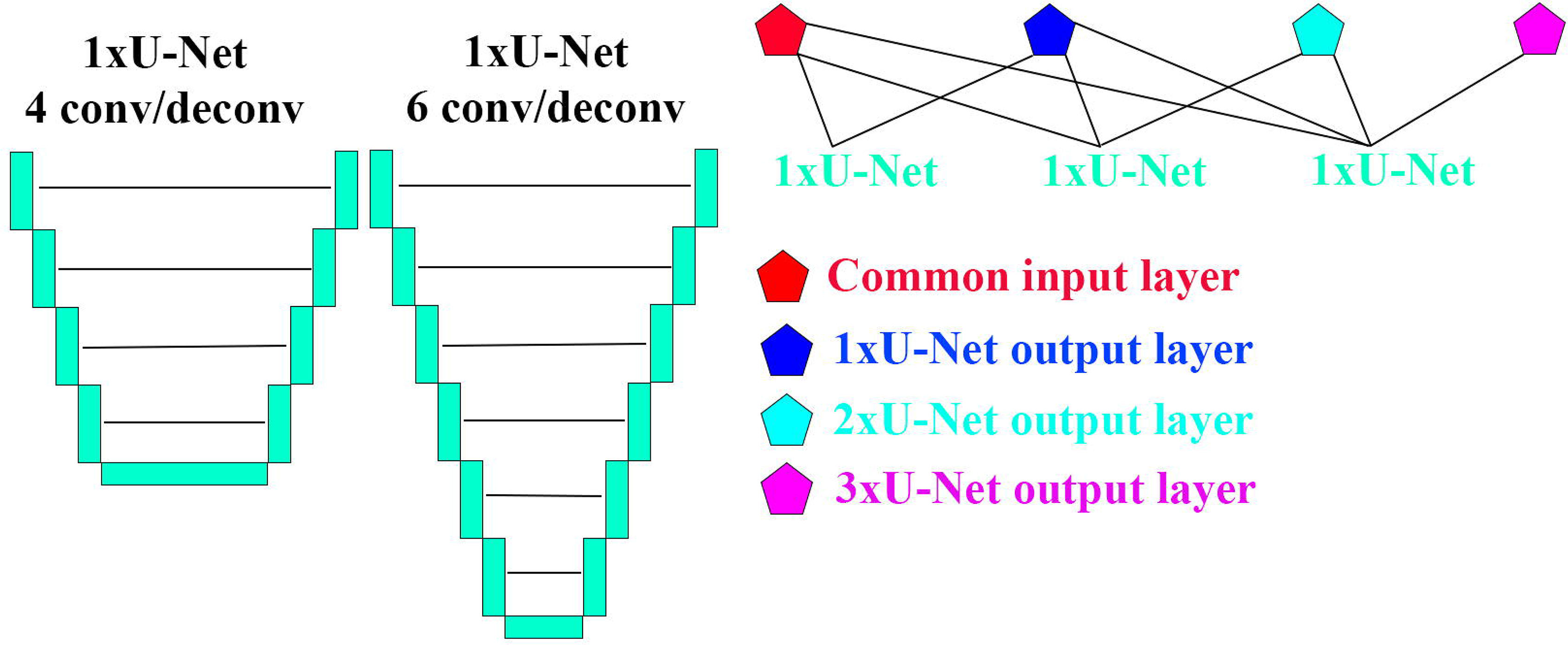
Architecture of U-Nets. (A) 1x-U-Net with 4 or 6 convolutions/deconvolutions (fractionally-stridden convolutions) without input and output layers. Each layer has batch normalization and some of them have a dropout (see the source code for the details). The lines indicate the skip-connections, 4 vs 6 skip-connections. (B) 1x-U-Net, 2x-U-Net, and 3x-U-Net architecture. Lines from polygons to 1x-U-Net are inputs and lines from 1x-U-Net to polygons are the outputs. Multiple inputs have been concatenated before using as input for the 1x-U-Net.

## RESULTS

In general, for image segmentation, original images and their ground truth (hereafter masks) data are needed to train convolutional neural networks (CNNs), which generate rules by adjusting parameters to find a general rule. However, generating masks on images requires expertise and is time-consuming. This is one of the rate-limiting and at the same time challenging steps in training NNs, which depends on good and variable training data. To solve this problem, GANs are commonly used to generate synthetic images for enhancing machine learning. While GANs have been successfully applied to generate images for biomedical and fluorescent images (Baniukiewicz et al., 2019; Han et al., 2018; Isola et al., 2017b; Mannam et al., 2021; Mirza & Osindero, 2014; Osokin et al., 2017), to our knowledge, it has not been applied to generate training data for complex high cell-density data. As a proof-of-principle, we have designed a strategy to generate cell nuclei in the zebrafish brain. We achieved this by generating a new way of training data and systematically evaluating several parameters on the U-Net architecture. As a proof-of-principle, we used zebrafish brain image (20X objective) cells stained DAPI, which shows a smooth distribution without visible intranuclear structures. Additionally, we used a mouse CA3 region from a hippocampal image taken at 40X with visible intranuclear structures (nuclear domains with high chromatin concentration). We started analyses by using Zebrafish brain nuclei to evaluate the U-Net models and then we further evaluated the models by using relatively complex cell nuclei from mice.

### Randomizing the masks increases the number of variable training data

For machine learning, cell nuclei and their masks are required for training a model. In general, masks are generated on images, and a model is used to learn how to generate masks from the images. To generate more images, we started from an inverse path, thus we used masks to train a model to generate images (here cell nuclei). To generate training data, we used 2 images (**Fig. S1**). We put the masks on 4 channels such that the minimum distance between 2 adjacent masks on the same channel was 2 pixels. We sought that masks can be distributed differently on channels, so we randomized the masks and arranged them on channels which increased the number of variable training data (**Fig. 1**). We also generated another training data that has one additional channel, which is the merge/cumulative sum of 4 channels. In such a case, we aimed to check if adding additional information to training data would improve generator performance. We generated 128×128 pixel images, the minimum size required to train the 6-layer U-Net models. We generated images by moving 4 pixels horizontally and/or vertically. The resulting 4,110 images were 10 times randomized, inverted, and rotated 3 times 90 degrees, which in total resulted in 205,500 images. Here, we started analyses with only 2 images which in total have 850 cell nuclei, and by randomizing masks we augmented data that generated variable cell nuclei shapes and distributions. This is a new way of augmenting images to generate training data. We did not augment images by image augmentation in TensorFlow, which might further add variation in training data that might improve GAN performance (Ching et al., 2018). We trained all models with 4,110 images by inverting and rotating as above, to check if augmenting images by just randomizing the masks improves the performance of GAN models.

### Generating random test images by GANs to check the performance of GANs

To generate random test images that have different mask distributions, and densities, we used all 850 masks generated from zebrafish images to train a GAN model that can generate random masks from random noise. Thus, we used all 850 masks generated above and randomly put them in 28×28 pixel images to build test images. In this way, the masks were either in the 28×28 pixel area or partly outside of the area that generated variable masks. We trained a neural network to generate random masks. These random masks were laid on 512×512 pixel images to generate low-density to high-density masks. The high-density masks must have had at least 200 masks on 512×512 pixel images. As the masks were randomly generated and densely distributed, we believe that this already constituted unseen/test data to evaluate the performance of GANs. This trained model to generate random masks can be used to generate variable dense masks with random distribution, which was one of our aims to generate variable nuclei images from these masks.

### Chaining U-Net architectures with alternating activation functions generate better images

U-Net is one of the most used Convolutional Neural Networks (CNNs) for image segmentation and image generation (Ronneberger et al., 2015). We started using U-Net as a generator and checked if it could learn and generate cell nuclei from the masks. Based on some preliminary analyses (not shown), we realized that we needed to change parameters (kernel size, filter size, and activation functions) and the architecture of U-Net. The U-Net that we used had either 4 or 6 convolutions and deconvolutions (fractionally-strided convolutions), hereafter 1x U-Net (**Fig. 2**). Thus, we tried to test as many different conditions as possible in order to find conditions that can be further evaluated on different datasets. To check if chaining 2 or more U-Nets can perform better in learning and generating nuclei from the masks, we concatenated the output of the 1x U-Net with the input layer and used it as input for a second U-Net (2x U-Net), and we concatenated the outputs of 1x and 2x U-Nets with the input layer and used as input for a third U-Net (3x U-Net). In total, we used 3 models as generators and a common discriminator for all these models. For 1x U-Net, we used 16, 32, and 64 filter sizes, while for 2x and 3x U-Net 16 filter sizes were used. Additionally, all models were tested for 3, 5, and 7 kernel sizes, except 1x U-Net with 64 filter sizes and kernel size 7. Furthermore, the first U-Net of 2x and the second of 3x U-Nets have either “*tanh*” or “*gelu*” activation functions in the output layer. All U-Nets and their conditions are summarized in **Table 1**. All conditions in **Table 1** were further trained on 4 channels and/or merged of 4 channels. As a result, we checked for 40 different conditions and evaluated them on 128×128 and 512×512 test images. We started first with visual examinations of the generated cell nuclei in 128×128 pixel images by all models and their parameters. If the models did not have a remarkable difference by visual evaluation, then we tested on 512×512 test images.

**Table 1:** The number of total parameters for each model with different options. FS; filter size, KS; kernel size, AF; activation function, 4,6-layers are for each 1x-U-Net layer of all models, 4/5-channels are the number of input channels. Note; the 1^st^ chanel of 5-channels is the cumulative sum of the last 4-channels.

|  |  |  |  | 4-layers |  | 6-layers |  |
| --- | --- | --- | --- | --- | --- | --- | --- |
|  | FS | KS | AF | 4-channels | 5-channels | 4-channels | 5-channels |
| 1x U-Net | 16 | 3 | tanh | 883.873 | 884.017 | 14.164.129 | 14.164.273 |
|  | 32 | 3 | tanh | 3.528.001 | 3.528.289 | 56.630.593 | 56.630.881 |
|  | 64 | 3 | tanh | 14.097.025 | 14.097.601 | 226.470.529 | 226.471.105 |
|  | 16 | 5 | tanh | 2.450.081 | 2.450.481 | 39.323.297 | 39.323.697 |
|  | 32 | 5 | tanh | 9.789.761 | 9.790.561 | 157.264.193 | 157.264.993 |
|  | 64 | 5 | tanh | 39.137.921 | 39.139.521 | 628.998.785 | 629.000.385 |
|  | 16 | 7 | tanh | 4.799.393 | 4.800.177 | 77.062.049 | 77.062.833 |
|  | 32 | 7 | tanh | 19.182.401 | 19.183.969 | 308.214.593 | 308.216.161 |
|  | 64 | 7 | tanh | 76.699.265 | 76.702.401 | 1.232.791.169 | 1.232.794.305 |
| 2x U-Net | 16 | 3 | tanh, tanh | 1.767.890 | 1.768.178 | 28.328.402 | 28.328.690 |
|  | 16 | 3 | gelu, tanh | 1.767.890 | 1.768.178 | 28.328.402 | 28.328.690 |
|  | 16 | 5 | tanh, tanh | 4.900.562 | 4.901.362 | 78.646.994 | 78.647.794 |
|  | 16 | 5 | gelu, tanh | 4.900.562 | 4.901.362 | 78.646.994 | 78.647.794 |
|  | 16 | 7 | tanh, tanh | 9.599.570 | 9.601.138 | 154.124.882 | 154.126.450 |
|  | 16 | 7 | gelu, tanh | 9.599.570 | 9.601.138 | 154.124.882 | 154.126.450 |
| 3x U-Net | 16 | 3 | tanh, tanh, tanh | 2.652.051 | 2.652.483 | 42.492.819 | 42.493.251 |
|  | 16 | 3 | tanh, gelu, tanh | 2.652.051 | 2.652.483 | 42.492.819 | 42.493.251 |
|  | 16 | 5 | tanh, tanh, tanh | 7.351.443 | 7.352.643 | 117.971.091 | 117.972.291 |
|  | 16 | 5 | tanh, gelu, tanh | 7.351.443 | 7.352.643 | 117.971.091 | 117.972.291 |
|  | 16 | 7 | tanh, tanh, tanh | 14.400.531 | 14.402.883 | 231.188.499 | 231.190.851 |
|  | 16 | 7 | tanh, gelu, tanh | 14.400.531 | 14.402.883 | 231.188.499 | 231.190.851 |

### Additional information, kernel sizes, and chaining more U-Nets generate better images

1x U-Net is commonly used for GANs, therefore we first checked using 16, 32, and 64 filter sizes and 3, 5, and 7 kernel sizes. Increasing kernel size and/or filter size increases the time required for each epoch. As a result, we decreased the filter size from 64 to 32 and 16 for the first convolution layer and doubled it at each layer. Interestingly, kernel size 7 was the best for filter sizes 16 and 32, while kernel size 7 with filter size 64 was not tested because of the long-running time. The 1xU-Net with 64 filter sizes may need more epochs to perform as well as 16 and 32 (not shown here). However, we do not believe this is because of the number of total parameters that 1xU-Net with 64 filter size has, as 3xU-Net with almost 8-fold number of parameters learns better (with the same data and number of epochs) than 1xU-Net which has ~14×10e6 total parameters performance (**Figs. 3, 4, 5, and 6**).

**Figure 3:**
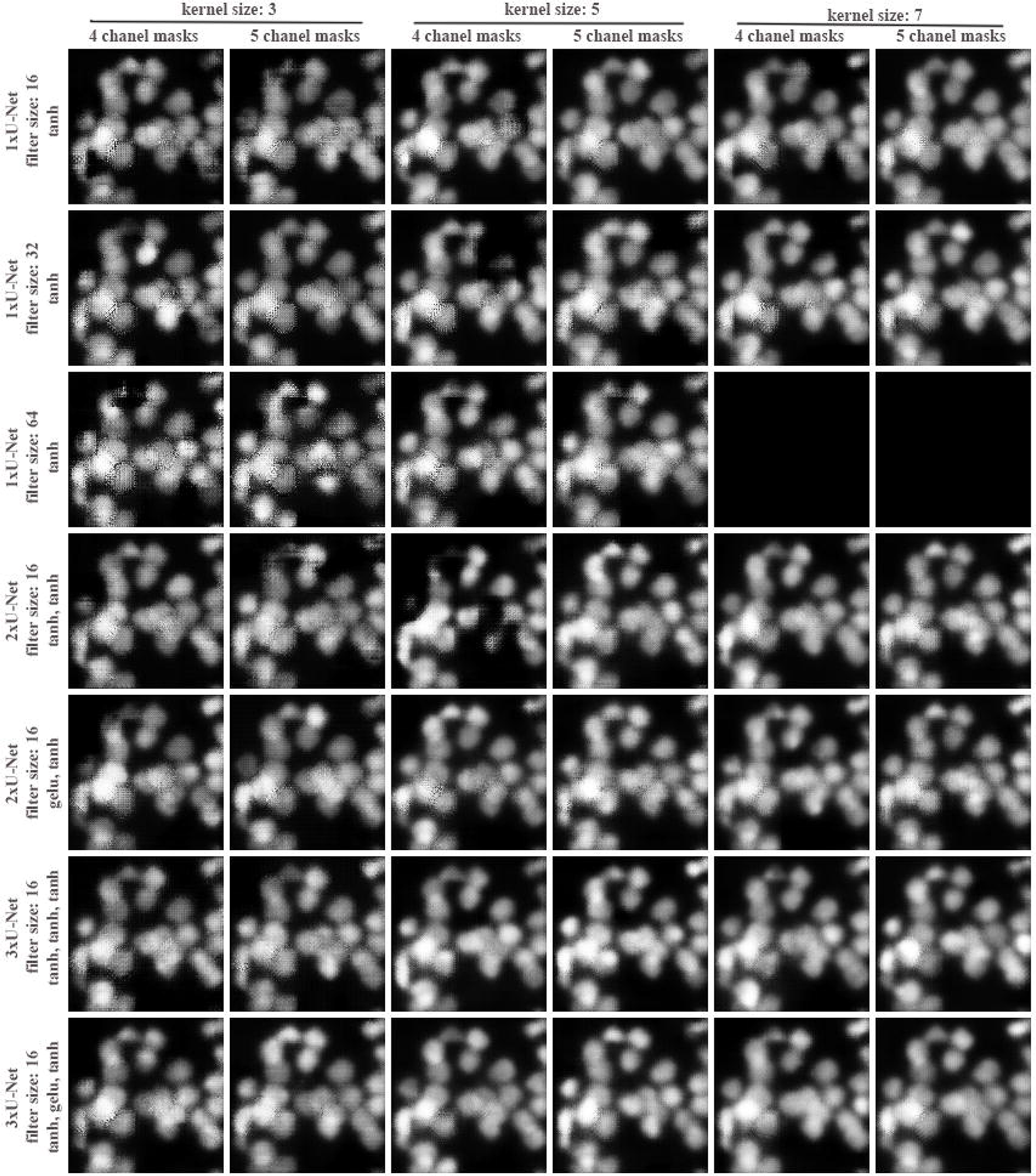
Generation of de-novo 128×128 pixel cell nuclei from the same test inputs by all models and conditions. U-Nets with 4 convolutions and deconvolutions have been used for all U-Net models. The original masks have been used for training directly without randomizing the masks 10 times. 4 channel masks; 4 mask layers have been used, 5 channel masks, and a merged channel of 4 channel masks was additionally added to the training data. The filter size and activation information are given on the left side of the figure. Note that 1xU-Net with 64 filter sizes and kernel size 7 were not tested.

**Figure 4:**
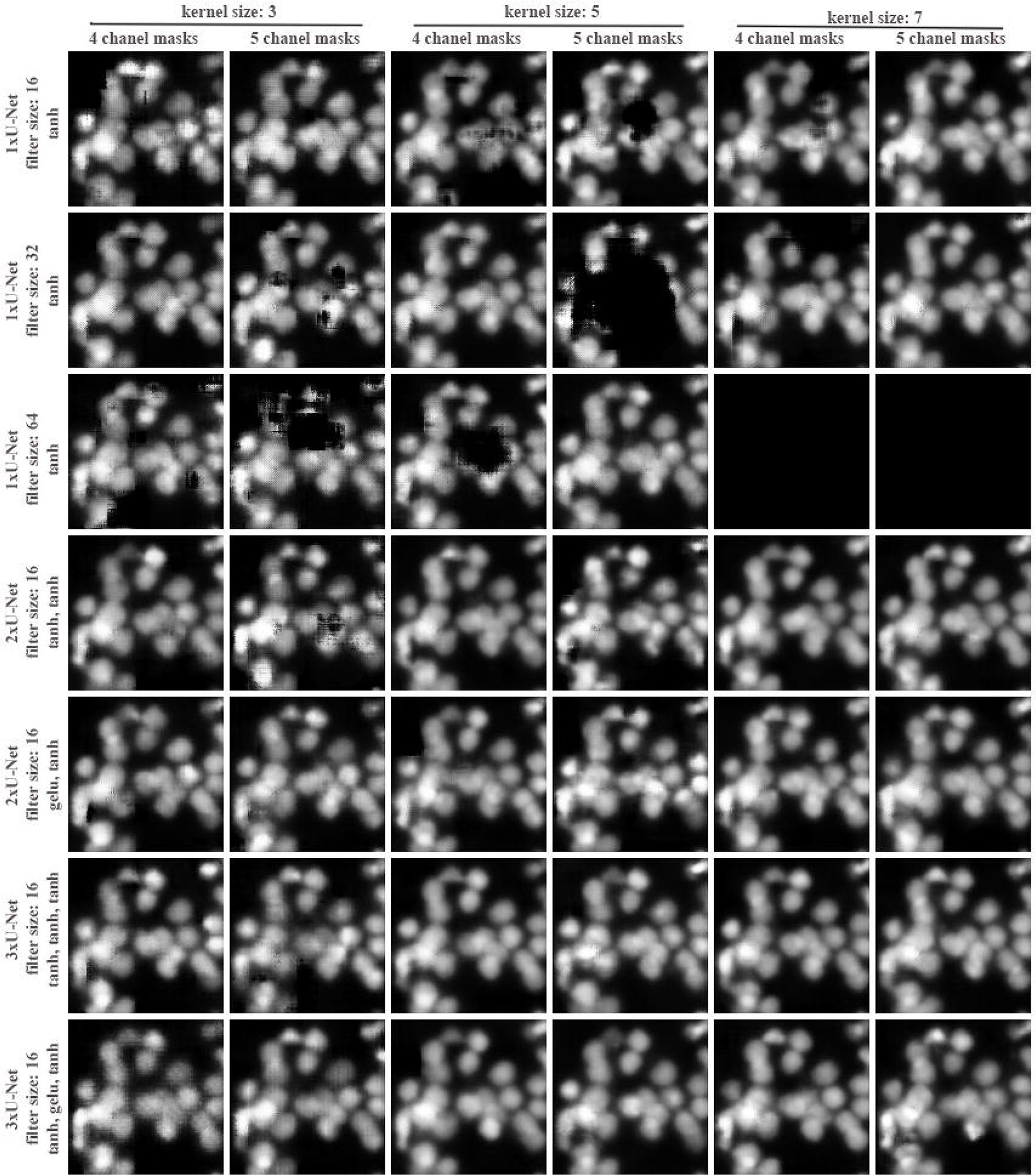
Generation of de-novo 128×128 pixel cell nuclei from the same test inputs by all models and conditions. U-Nets with 4 convolutions and deconvolutions have been used for all U-Net models. The original masks have been randomized 10 times. 4 channel masks; 4 mask layers have been used, 5 channel masks, and a merged channel for 4 channel masks was additionally added to the training data. The filter size and activation information are given on the left side of the figure. Note that 1xU-Net with 64 filter sizes and kernel size 7 were not tested.

**Figure 5:**
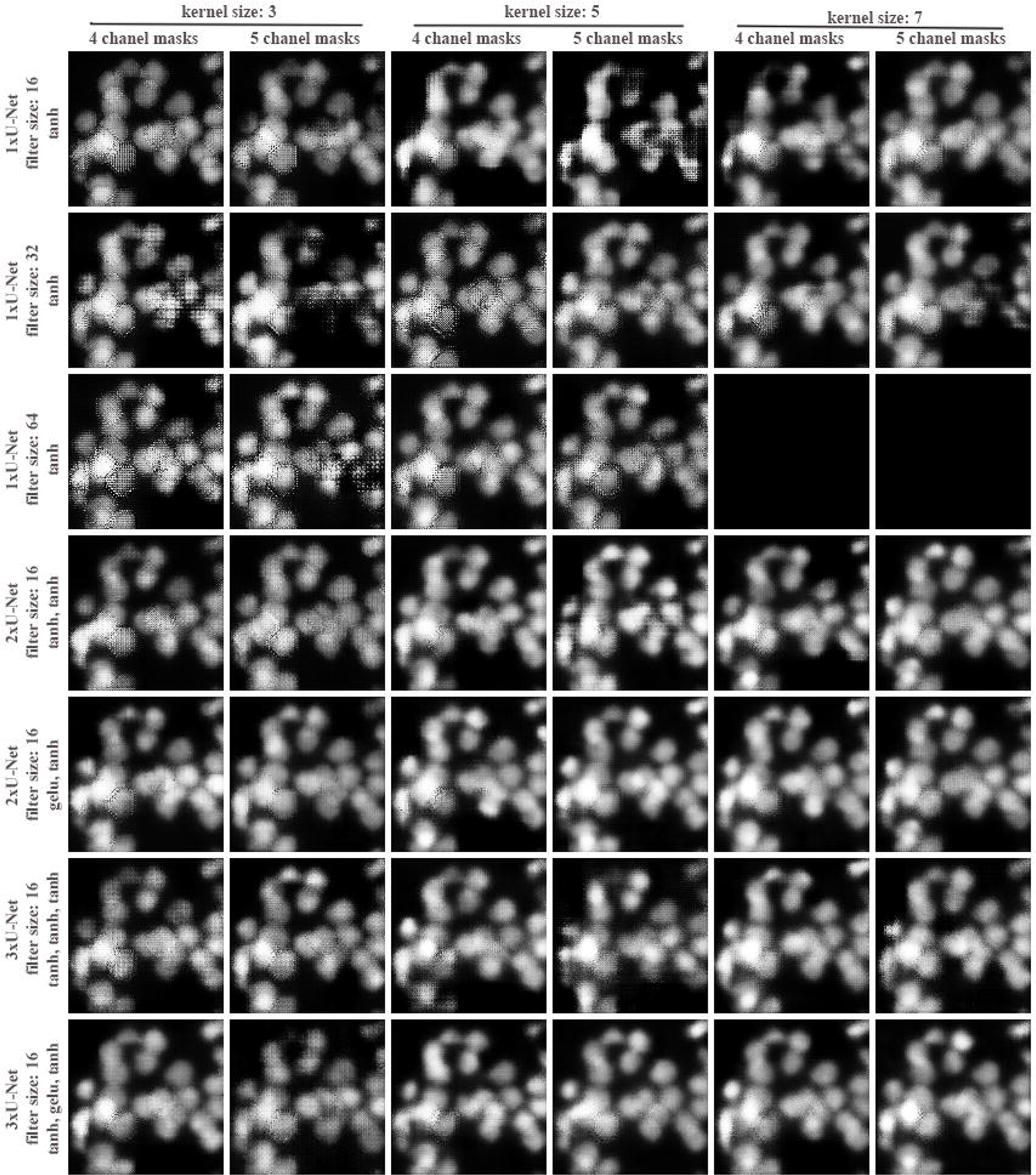
Generation of de-novo 128×128 pixel cell nuclei from the same test inputs by all models and conditions. U-Nets with 6 convolutions and deconvolutions have been used for all U-Net models. The original masks have been used for training directly without randomizing the masks 10 times. 4 channel masks; 4 mask layers have been used, 5 channel masks, and a merged chanel for 4 channel masks was additionally added to the training data. The filter size and activation information are given on the left side of the figure. Note that 1xU-Net with 64 filter sizes and kernel size 7 were not tested.

**Figure 6:**
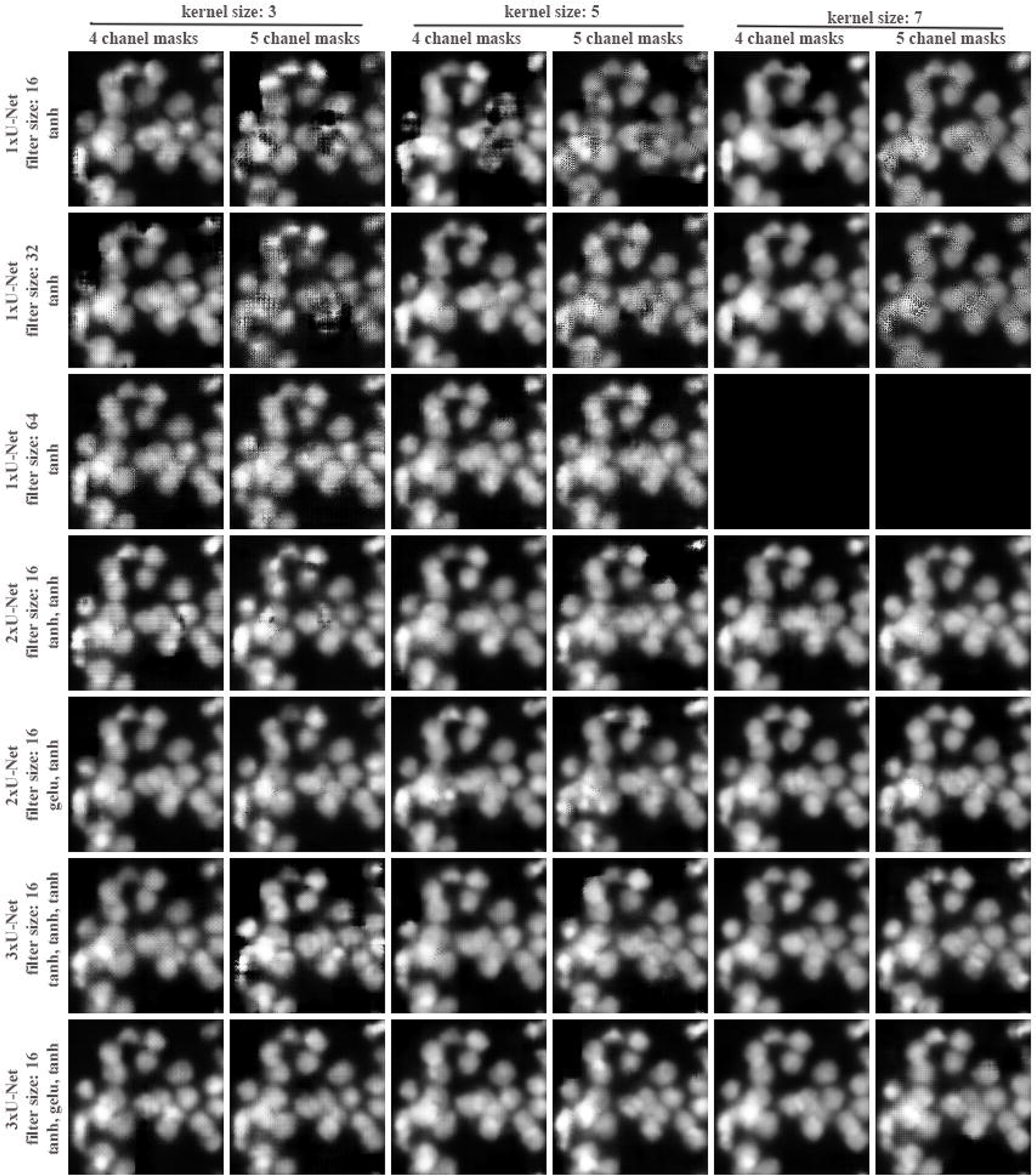
Generation of de-novo 128×128 pixel cell nuclei from the same test inputs by all models and conditions. U-Nets with 6 convolutions and deconvolutions have been used for all U-Net models. The original masks have been randomized 10 times. 4 channel masks; 4 mask layers have been used, 5 channel masks, and a merged chanel for 4 channel masks was additionally added to the training data. The filter size and activation information are given on the left side of the figure. Note that 1xU-Net with 64 filter sizes and kernel size 7 were tested.

As 1xU-Net with filter size 16 and kernel size 7 performed better, we generated 2x and 3x U-Nets and compared their performance. 1xU-Net with 16 filter sizes with kernel size 3 performs better than 2xU-Net with kernel size 3. However, increasing kernel size to 5 and 7, improved the performance of 2x U-Net. Additionally, adding a merged layer of masks to the training data further improved 2xU-Net performance (**Figs. 3, 4, 5, and 6**).

We checked if the increasing chain of U-Nets will further improve learning by testing 3x U-Net with the same conditions above. We found that 3xU-Net performs even better than 2xU-Net. In all of these cases, “*tanh*” was used for the output layers of all U-Nets. We asked whether changing the “*tanh*” to “*gelu*” in the first U-Net of 2x and the second U-Net of 3x U-Net would improve the performance, and found that adding an alternating activation function further improved the performance of the 2x and 3x U-Nets (**Figs. 3, 4, 5, and 6**) while using “*gelu*” as the last U-Net of 1x and 3x U-Net completely abolished the performance of 2x and 3x U-Nets (not shown).

The conclusions above were based on 128×128 pixel test images to evaluate the performance of all models and conditions indicated. To enhance the qualitative differences between the predictions of different models, we challenged the models by using 512×512 pixel images for evaluations, as 128×128 pixel images for evaluations might not have provided the required challenge for the models. As expected, challenging the models with larger images further enhanced the models’ performance. Interestingly, 3xU-Net with “*tanh*”, “*gelu*”, and “*tanh*” activation functions, performed remarkably better than other models (**Fig. S4**). This further supports that 3xU-Net can be trained with small-sized images, while it can be used to generate larger images.

We also tested whether randomizing masks to generate more training data had any impact on GAN performance. We used each original image one time and inverted, rotated 3 times 90 degrees. Then, we compared the GAN performance tested on 1 time versus 10 times randomized masks on each image. Adding more variation by randomizing the masks further improved the performance of GAN (see **Figs 3 and 4**, **Figs 5 and 6**).

In summary, based on our results from relatively simple cell nuclei, we concluded the followings; (i) increasing kernel sizes improves the performance of GANs, (ii) adding additional layers (6-layer U-Net) further improves the performance of GANs, (iii) 2x and 3x U-Nets perform better than regular U-Net architecture, (iv) adding an additional layer can improve the performance of 6-layer U-Net, but not 4-layer U-Net. However, further evaluation of these conditions on different datasets is required, as a result, we tested the same conditions on mouse brain nuclei which have more visible intranuclear structures.

### 3x U-Net can generate nuclei with intranuclear structures as well

In order to test the same conditions with different cell nuclei, we used nuclei images from a hippocampal area of the mouse brain. We selected 2 image areas (**Sup Fig. 2a** and **3a**), less crowded and relatively highly crowded areas.

We trained U-Nets with training images generated from **Sup Figs. 2b** and **3b** and used sub-images from **Sup Fig. 3c** to compare the generated images. As the number of sub-images generated from **Sup Fig 2b** was too many, we did not rotate or invert images to perform faster training. Contrary to Zebrafish brain nuclei, 4-layer U-Nets performed better than 6-layer U-Nets. Especially, kernel sizes 5 and 7 did not generate clear intranuclear structures compared to 4-layer U-Nets (**Figs. 7, 8**). These contrary results show that CNNs may perform differently on different datasets (cell shapes, etc.) and further evaluation of the 4 and 6-layer U-Net is required for different datasets. A further challenge of the trained models on **Sup Fig 3b**, further supported these conclusions. As **Sup Fig 3b** had a relatively high cell density data with 1024×1024 pixel size (4 fold compared to the training data), the difference between 4- and 6-layer U-Nets is not because of the number of cells but the performance differences (**Sup Figs. 5, 6, 7, 8**). We generated two further challenges for the models, by generating random masks and arranging them on 4 channels in an image of 1024×1024 pixels (**Sup Figs. 7, 8**) and an image similar to the dentate gyrus of the hippocampus (**Sup Figs. 9, 10, 11**). These challenges further supported that 2x and, especially, 3x U-Nets perform better than regular U-Nets, and combinations of activation functions further improve the performance of CNNs.

**Figure 7:**
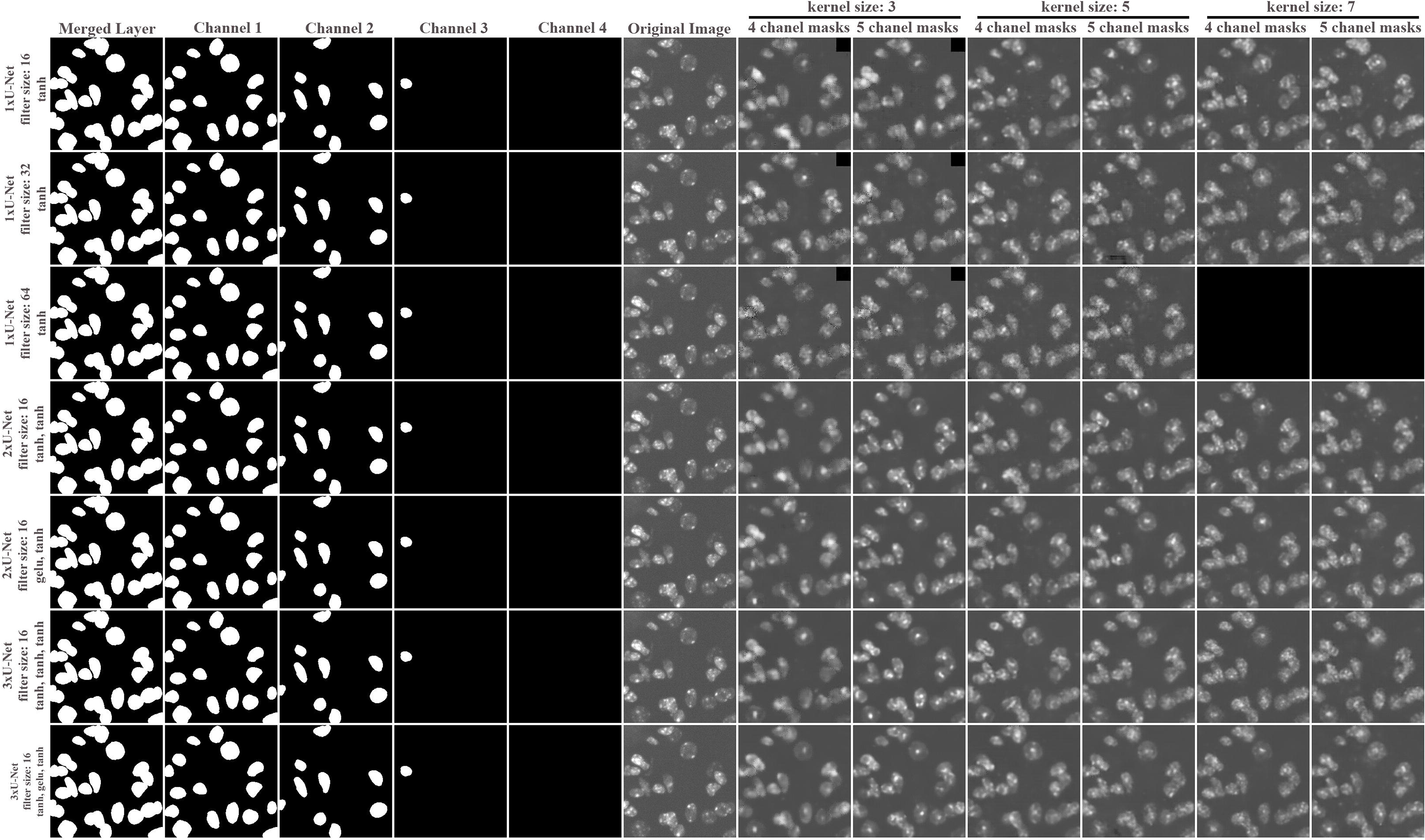
Generation of de-novo 256×256 pixel mouse cell nuclei from the same test inputs by all models and conditions. U-Nets with 4 convolutions and deconvolutions have been used for all U-Net models. The original masks have been randomized 10 times. 4 channel masks; 4 mask layers have been used, 5 channel masks, and a merged chanel for 4 channel masks was additionally added to the training data. The filter size and activation information are given on the left side of the figure. Note that 1xU-Net with 64 filter sizes and kernel size 7 were not tested. The original image and masks used for testing have been provided on the left side of the image (Merged Layer, Chanel 1-4, Original Image).

**Figure 8:**
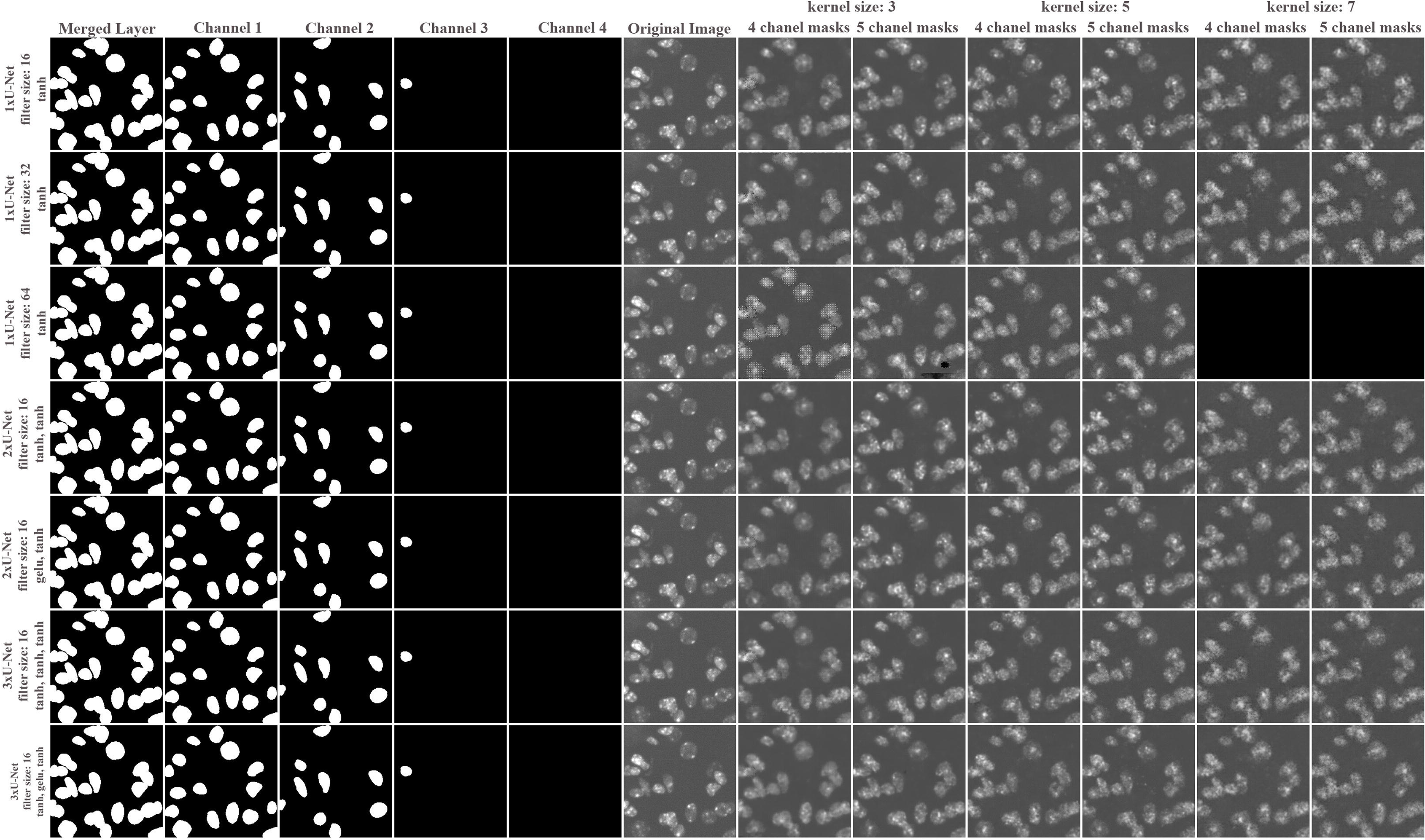
Generation of de-novo 256×256 pixel mouse cell nuclei from the same test inputs by all models and conditions. U-Nets with 6 convolutions and deconvolutions have been used for all U-Net models. The original masks have been randomized 10 times. 4 channel masks; 4 mask layers have been used, 5 channel masks, and a merged chanel for 4 channel masks was additionally added to the training data. The filter size and activation information are given on the left side of the figure. Note that 1xU-Net with 64 filter sizes and kernel size 7 were not tested. The original image and masks used for testing have been provided on the left side of the image (Merged Layer, Chanel 1-4, Original Image).

## DISCUSSION

Augmenting images for machine learning by using GAN has been done on real-life images, biomedical images, and fluorescent images (Baniukiewicz et al., 2019; Goldsborough et al., 2017; Han et al., 2018; Isola et al., 2017b; Johnson et al., 2017; Mannam et al., 2021; Osokin et al., 2017). Here, we not only generate new images by GAN but provide their masks (ground truth or segmented images) as well. In that way, we decreased the time required to generate training data. The strategy may be further adapted for irregular cell shapes.

To optimize GANs, we further evaluated many parameters and U-Net architecture, to find the best model for such kinds of analyses. One of the parameters we tested is kernel size, increasing kernel size increases running time. To compensate for it, we decreased the filter size (features) which decreased the total number of parameters and running time. Kernel size has also been an important parameter in CNN learning (Ding et al., 2022; Hu et al., 2020). However, increasing the kernel size increases the running time required for each epoch. For the current study, kernel size 5 was sufficient to generate simple cell nuclei, while using 7 gave slightly better results. However, different kernel sizes can be further tested for different cell sizes and shapes.

Our reference of comparison was to compare with 1xU-Net, especially with filter size 64, which is commonly used. Concatenating 1xU-Nets and adding alternating functions created a model that performed better. This might be because of concatenating output layers with the inputs and the addition of the “*gelu*” function, both of which may add variation in feature selection that improves the learning ability of the model.

U-Net architecture has been used to generate different U-Net models. These models have not been tested here; however, 3x U-Net with alternating activation functions was sufficient to generate variable cell nuclei. Further detailed comparisons of all available models as generators on our dataset will further help to improve GANs to generate variable data. Thus, understanding how and why each CNNs based on U-Net architectures learns from the same data will help us to develop a general model that can be used for variable tasks.

Our approach works for zebrafish and mouse brains’ cell nuclei. However, cells may have different shapes or intracellular structures in different organisms. Therefore, further evaluation and optimization may be needed to apply this strategy to different cell shapes with intracellular structures.

One limitation of the current study is the generation of only 2D images instead of 3D images. We will adapt this strategy to generate 3D images as well. Additionally, while evaluating the models and their parameters, our main aim was to use a minimum number of epochs, training data, and time per epoch. However, we do not exclude the possibility that all models could perform similarly by running with more epochs, adjusting hyperparameters and especially with more variable training data (Lucic et al., n.d.-a). Finally, one of the challenges in assessing the quality of the GANs is the qualitative approach that can be applied to GANs. Although there are some quantitative approaches to measure the performance of GANs, (Alaa et al., 2021; Borji, 2019; Heusel et al., n.d.; Lucic et al., n.d.-b; Salimans et al., n.d.; Xu et al., 2018), we challenged the models by using with large-sized images which clearly shows the difference between the model evaluated here.

## Code Availability

The Source code is available on https://github.com/micosacak/de-novo-cell-nuclei-by-GAN.

## Author Contributions

Conceptualization, formal analysis, investigation, methodology, resources, writing-original draft M.I.C.; funding C.K., review and editing, M.I.C., C.K. Both authors read and agreed to the published version of the manuscript.

## Acknowledgment

We would like to thank Dr. Gerd Kempermann for the GPU computer used for the preliminary analyses. Main analyses were performed on HPC clusters at German Center for Neurodegenerative Diseases (DZNE).

## Supplementary Figure Legends

**Fig. S1**: The original dapi staining of 2 cropped images (left images), with their masks (right images). Blue, Green, Red, and Light Gray in channels 1, 2, 3, and 4, respectively.

**Fig. S2:** Generation of training images from mouse hippocampal area. **(a)** the original image and the area (right-bottom) selected to generate training data **(b)** the dapi and masks on channels (msk0, msk1, msk2, msk3).

**Fig. S3:** Generation of training and test images from mouse hippocampal area. **(a)** the original image and the areas selected to generate training data, **(b)** a 1024×1024 pixel area used to challenge the models after 25 epochs of the training. **(c)** an area used to generate sub-images for testing models’ performance at each epoch; the dapi, and masks on channels (msk0, msk1, msk2, msk3).

**Fig. S4**: 512×512 pixel images test images. All models with kernel size 5 were tested on **(a)** 1-time randomized masks, **(b)** 10-times randomized masks, **(c)** 1-time randomized and with an additional merged layer, **(d)** 10-times randomized and with an additional merged layer.

**Fig. S5**: Generation of training data from mouse hippocampus. **(a)** the mouse hippocampus, **(b)** the area of the mask used to add as many masks as possible, **(c)** the masks, **(d)** the generated image by 3xU-Net, with kernel size 3, the addition of merged masks, 4-layer U-Nets. Note: The black area in the generated image is caused by zero-padding to the training data which generated the black area around nuclei. The original image has a background that has not been removed.

**Fig. S6:** A custom image generated by generating random masks and arranged on 1024×1024 pixel image, cell nuclei were generated by the models trained on **Sup Fig. 2b, (a) 4-layer U-Nets**, **(b) 6-layer U-Nets**.

**Fig. S7:** A custom image generated by generating random masks and arranging them on 1024×1024 pixel image, cell nuclei were generated by models were trained on **Sup Fig. 3b, (a) 4-layer U-Nets**, **(b) 6-layer U-Nets**.

**Fig. S8:** The models trained on Sup Fig 2b have been challenged on a relatively crowded area in Sup Fig 3b, **(a)** 4-layer U-Nets, **(b)** 6-layer U-Nets.

**Fig. S9:** Generation of dentate gyrus (DG) area of mouse hippocampus. **(a)** a mask area from crowded cell area in **Sup Fig. 3a**, **(b)** randomly generated masks arranged and enriched in the DG, **(c)** cell nuclei generated by 3x-U-Net with “gelu” activations and kernel size 5 **(d)** cell nuclei generated by 3x-U-Net with “gelu” activation and kernel size 7.

**Fig. S10**: Generation of dentate gyrus (DG) area of mouse hippocampus using the model trained on **Sup Fig. 2b.**

**Fig. S11**: Generation of dentate gyrus (DG) area of mouse hippocampus using the model trained on **Sup Fig. 3b.**

