## Supplementary figures and images for "Using conditional Generative Adversarial Networks (GAN) to generate *de novo* synthetic cell nuclei for training machine learning-based image segmentation"

### hc_more_test_mask2dapi_001_px_FS_16_KS_3_ML_True_4.jpg

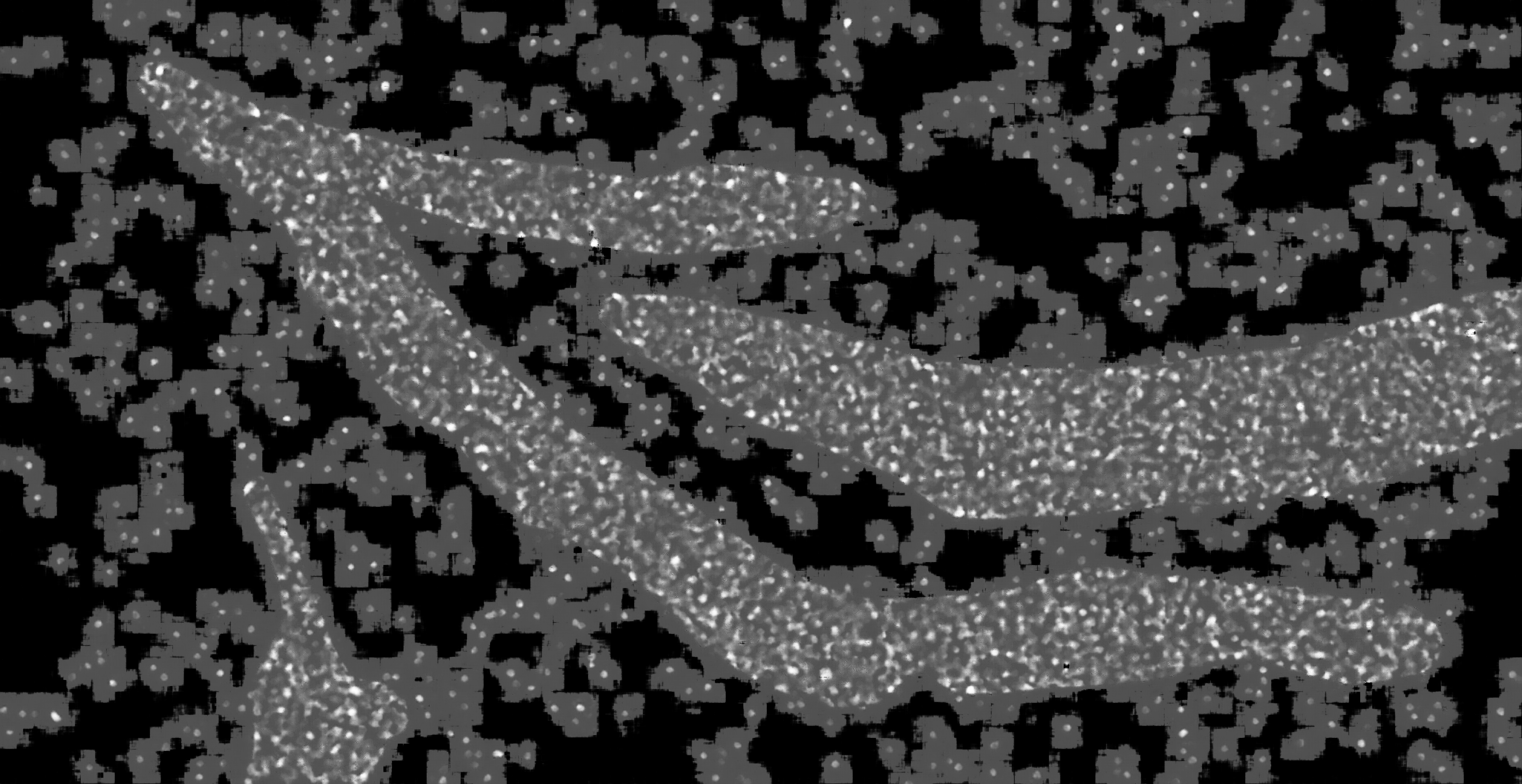

### hc_more_test_mask2dapi_001_px_FS_16_KS_5_ML_False_4.jpg

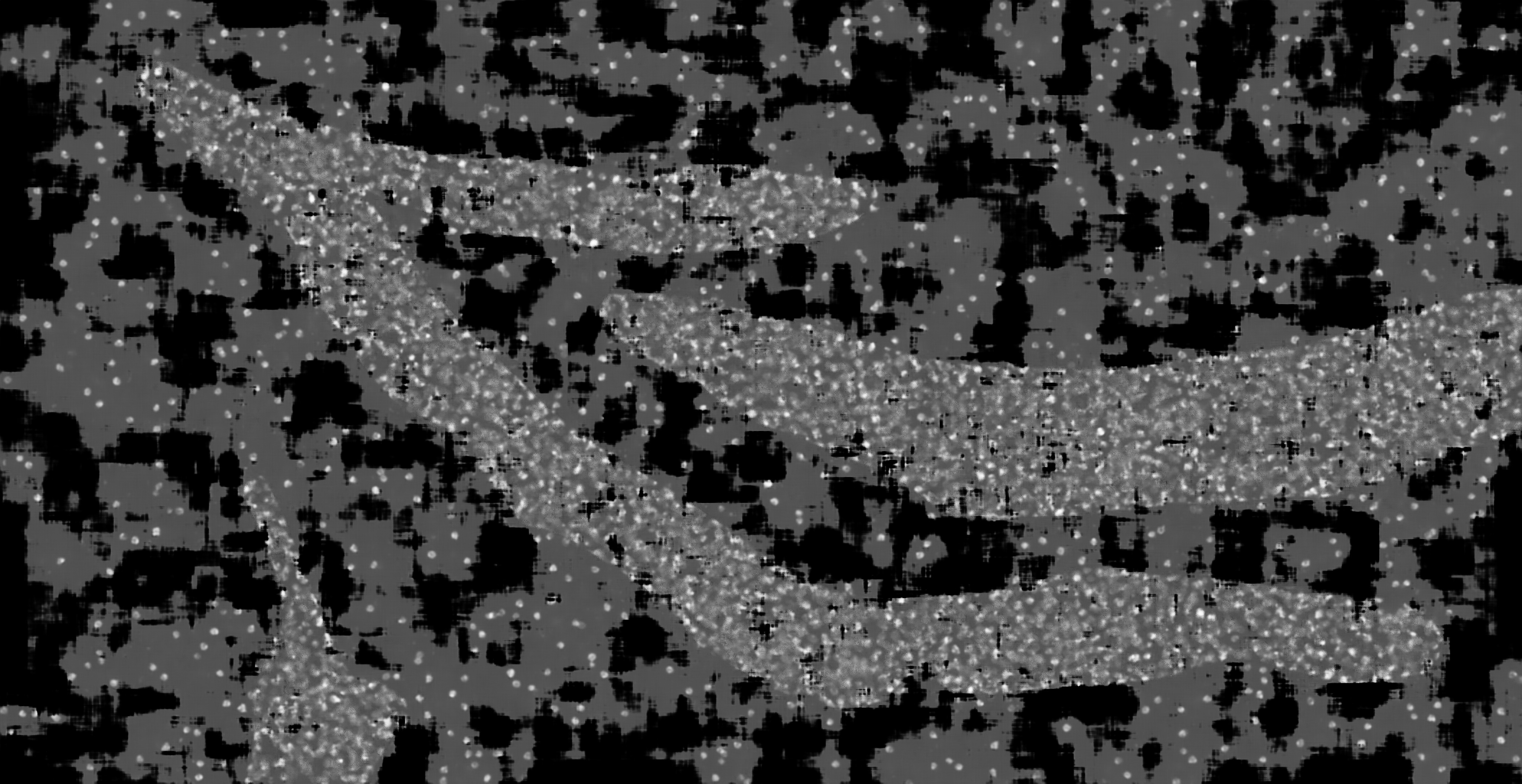

### hc_more_test_mask2dapi_001_px_FS_16_KS_7_ML_True_4.jpg

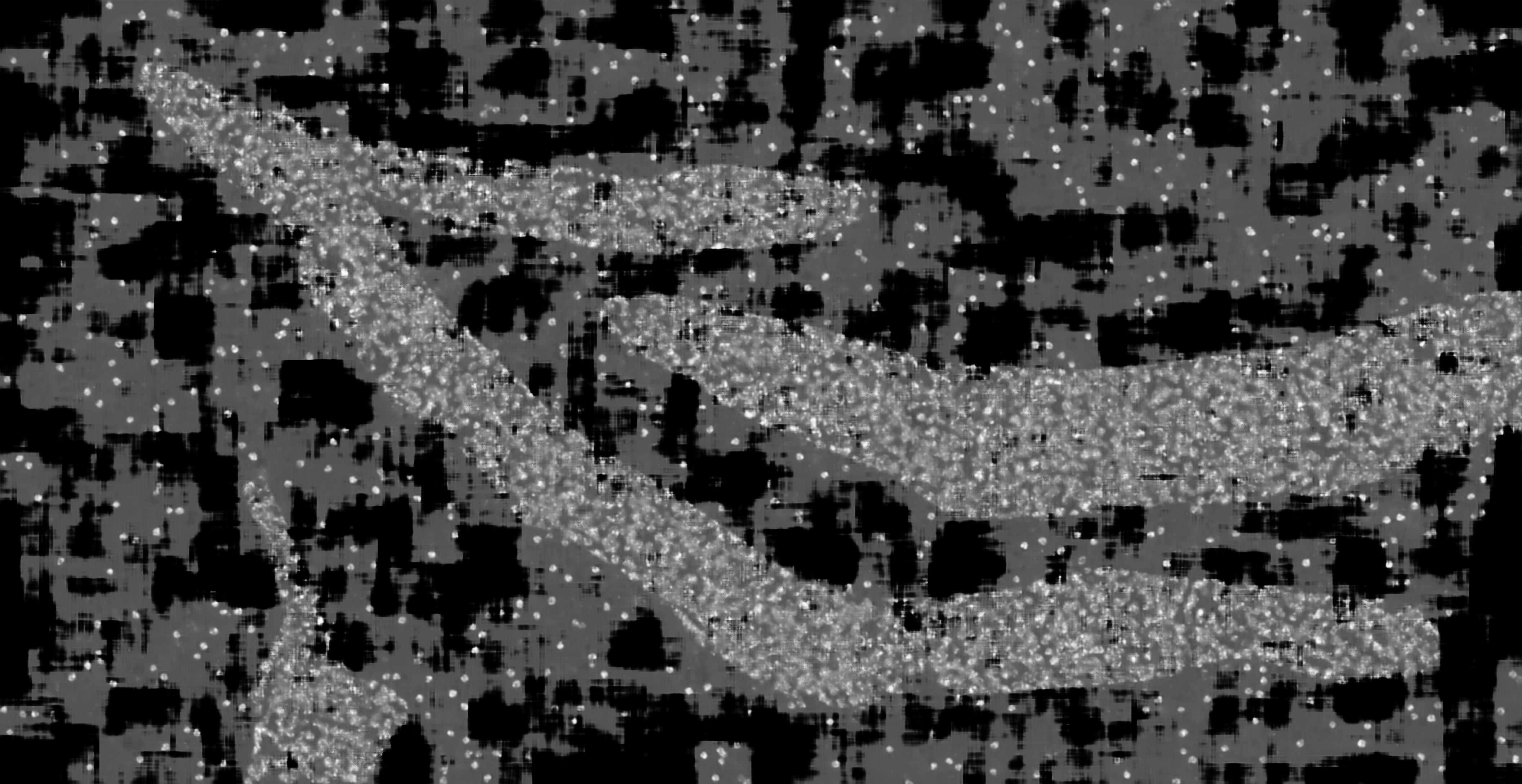

### hc_more_test_mask2dapi_001_px_FS_32_KS_3_ML_False_4.jpg

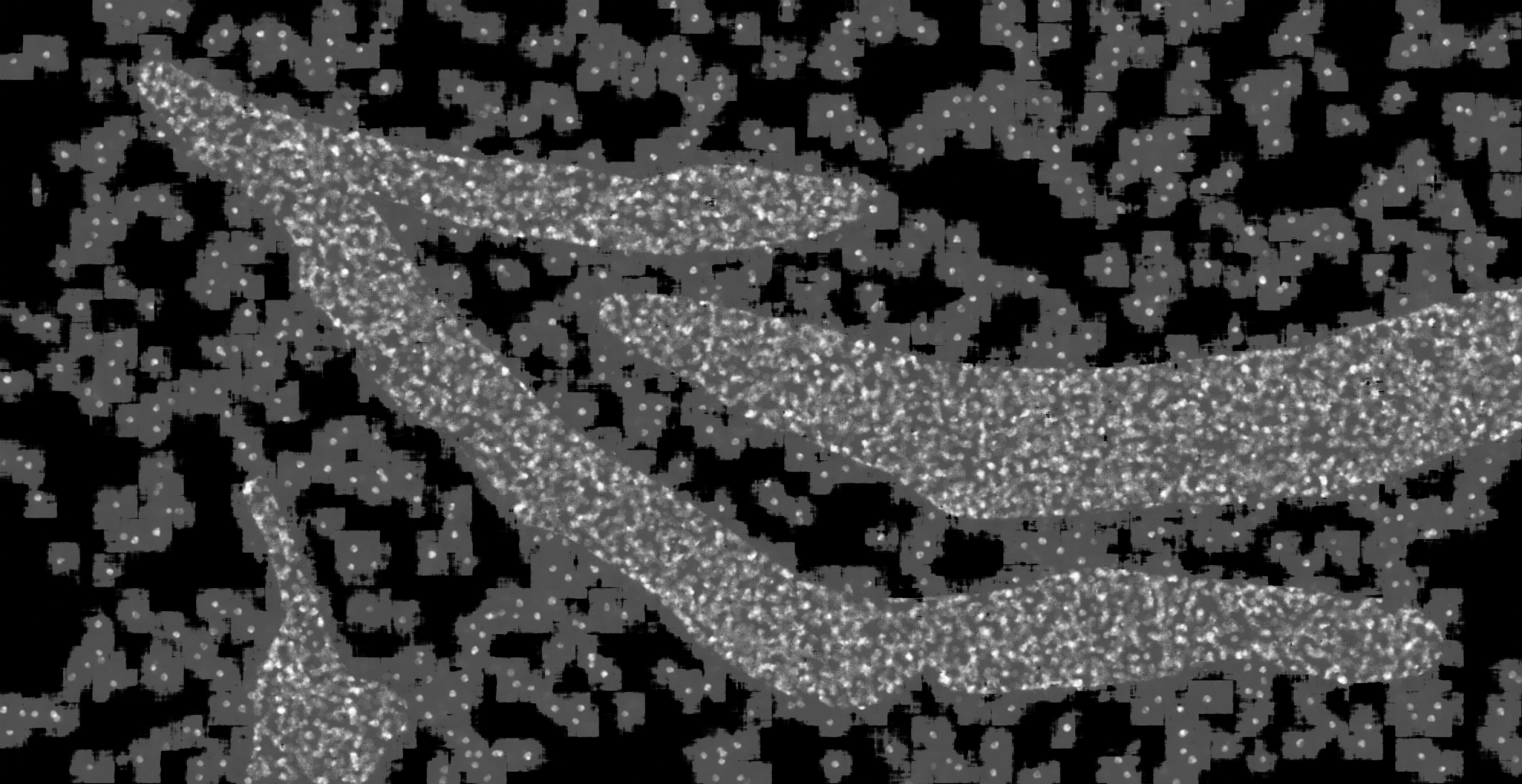

### hc_more_test_mask2dapi_001_px_FS_32_KS_5_ML_False_4.jpg

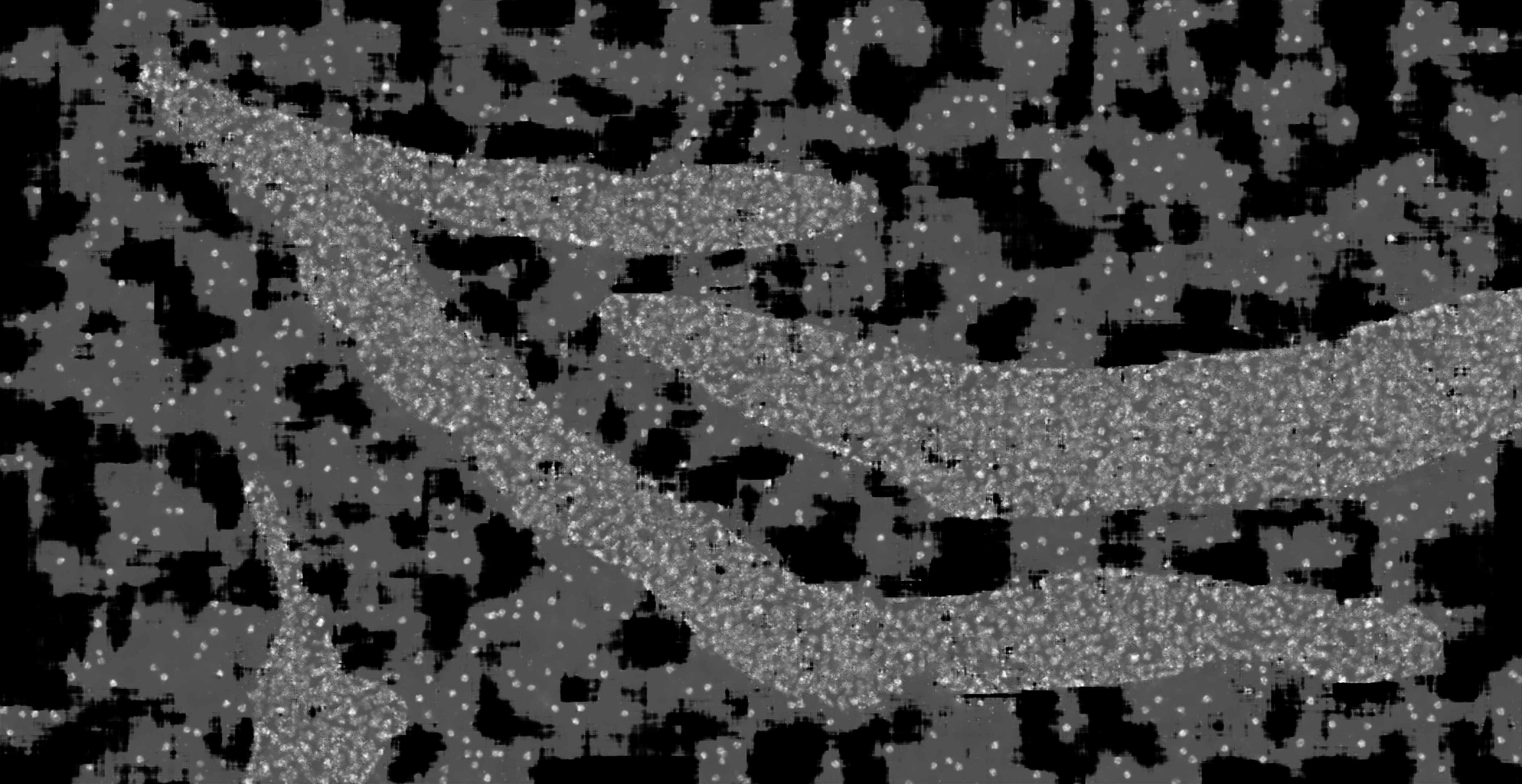

### hc_more_test_mask2dapi_001_px_FS_32_KS_5_ML_True_4.jpg

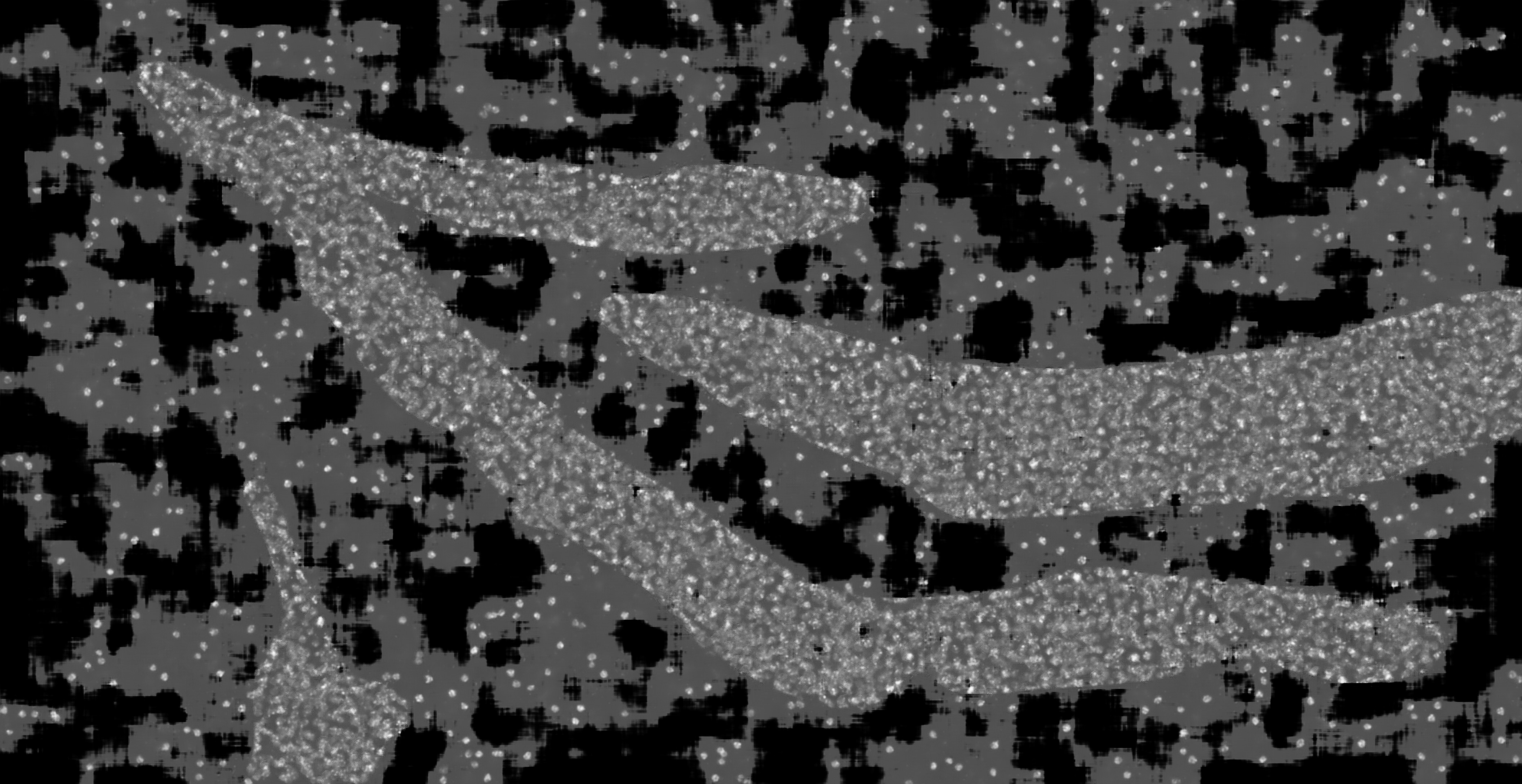

### hc_more_test_mask2dapi_001_px_FS_32_KS_7_ML_False_4.jpg

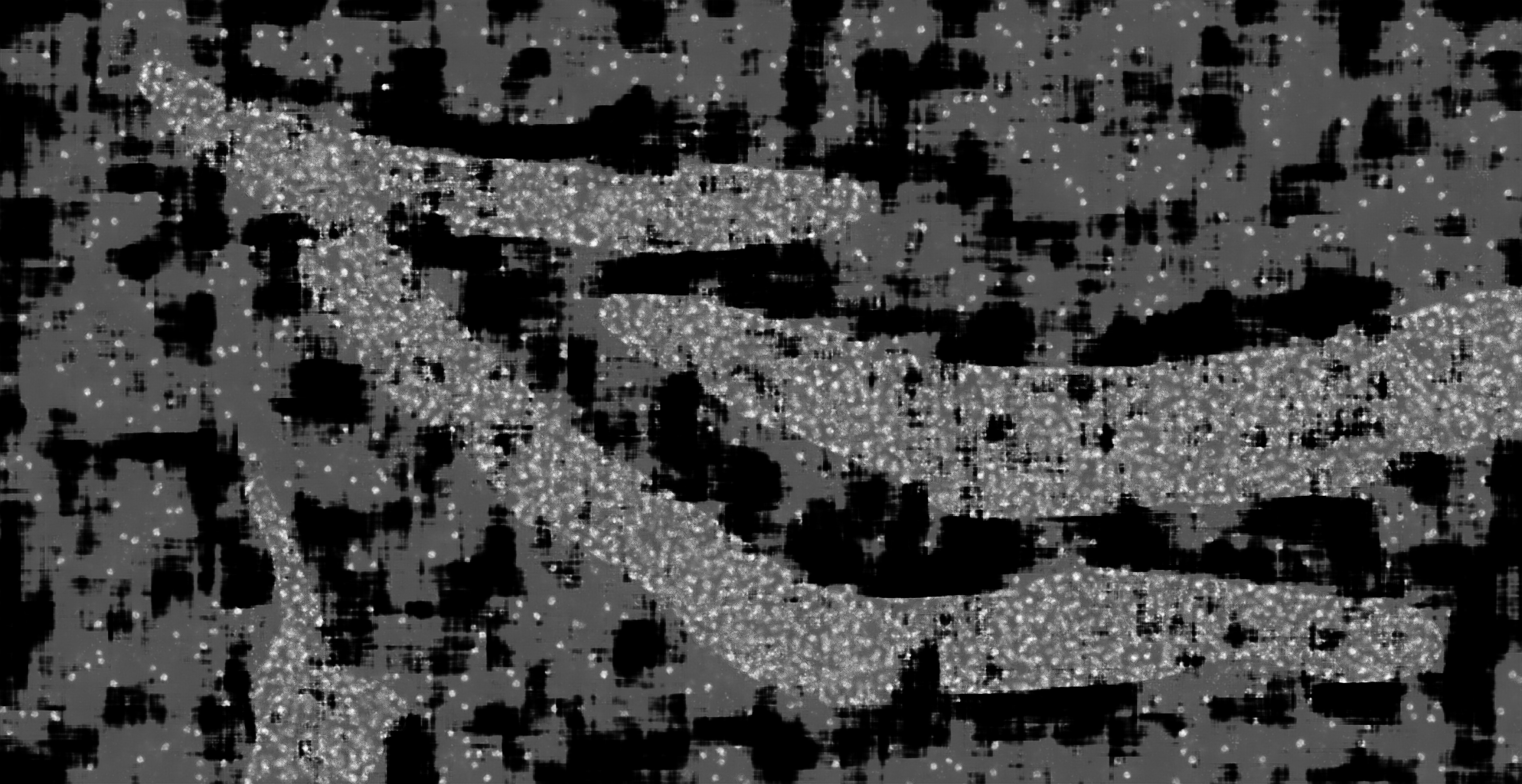

### hc_more_test_mask2dapi_001_px_FS_64_KS_3_ML_False_4.jpg

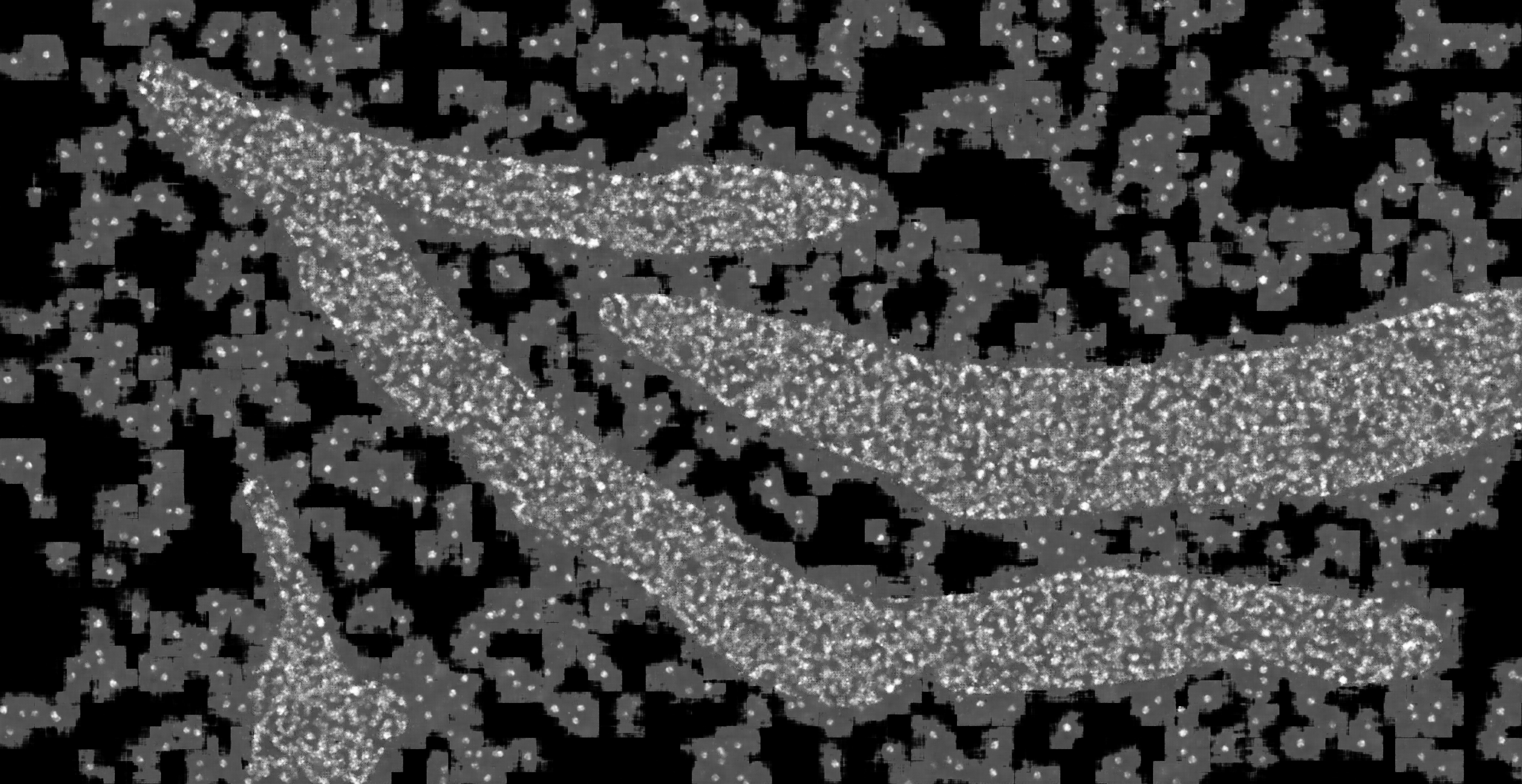

### hc_more_test_mask2dapi_001_px_FS_64_KS_3_ML_True_4.jpg

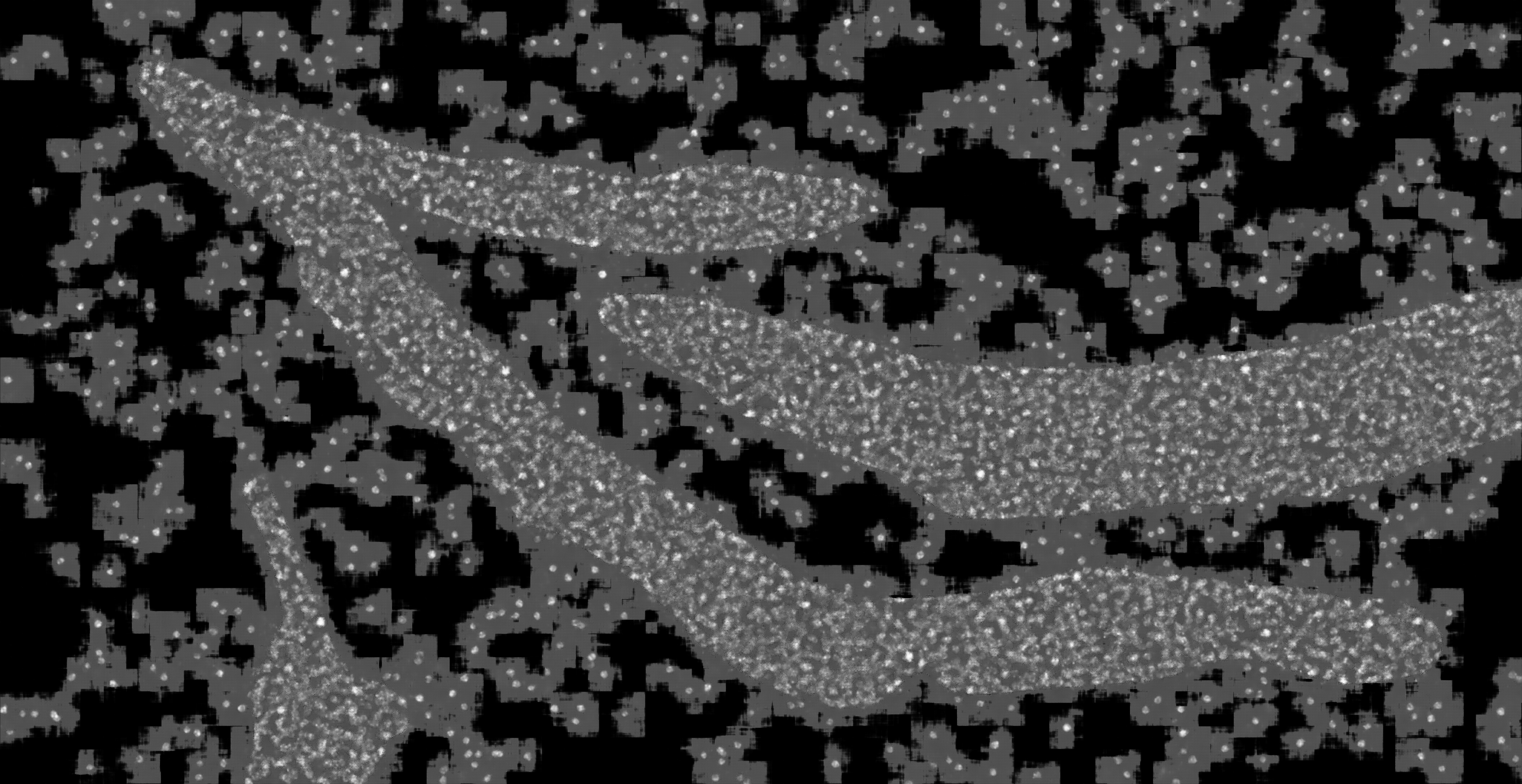

### hc_more_test_mask2dapi_001_px_FS_64_KS_5_ML_False_4.jpg

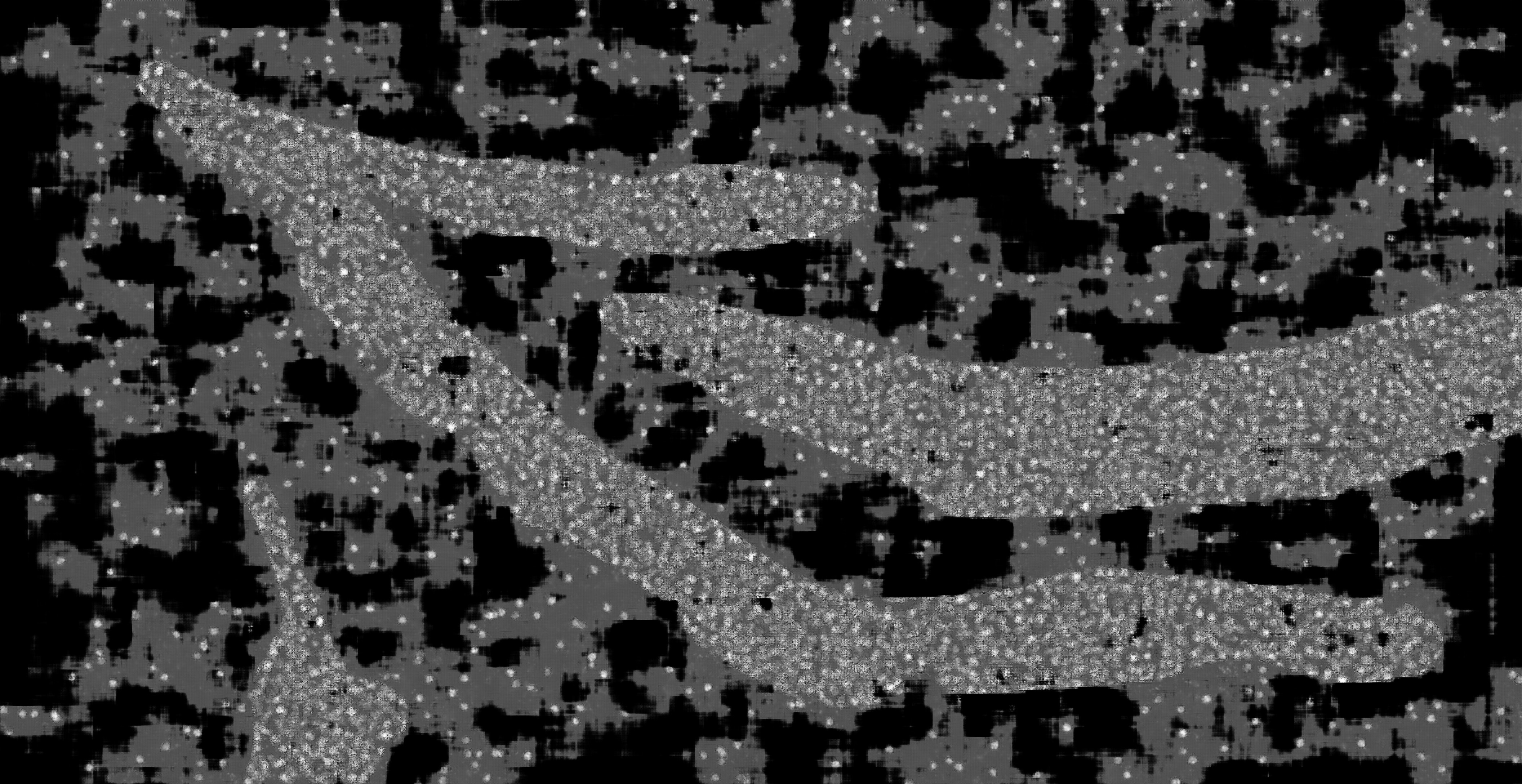

### hc_more_test_mask2dapi_001_px_FS_64_KS_5_ML_True_4.jpg

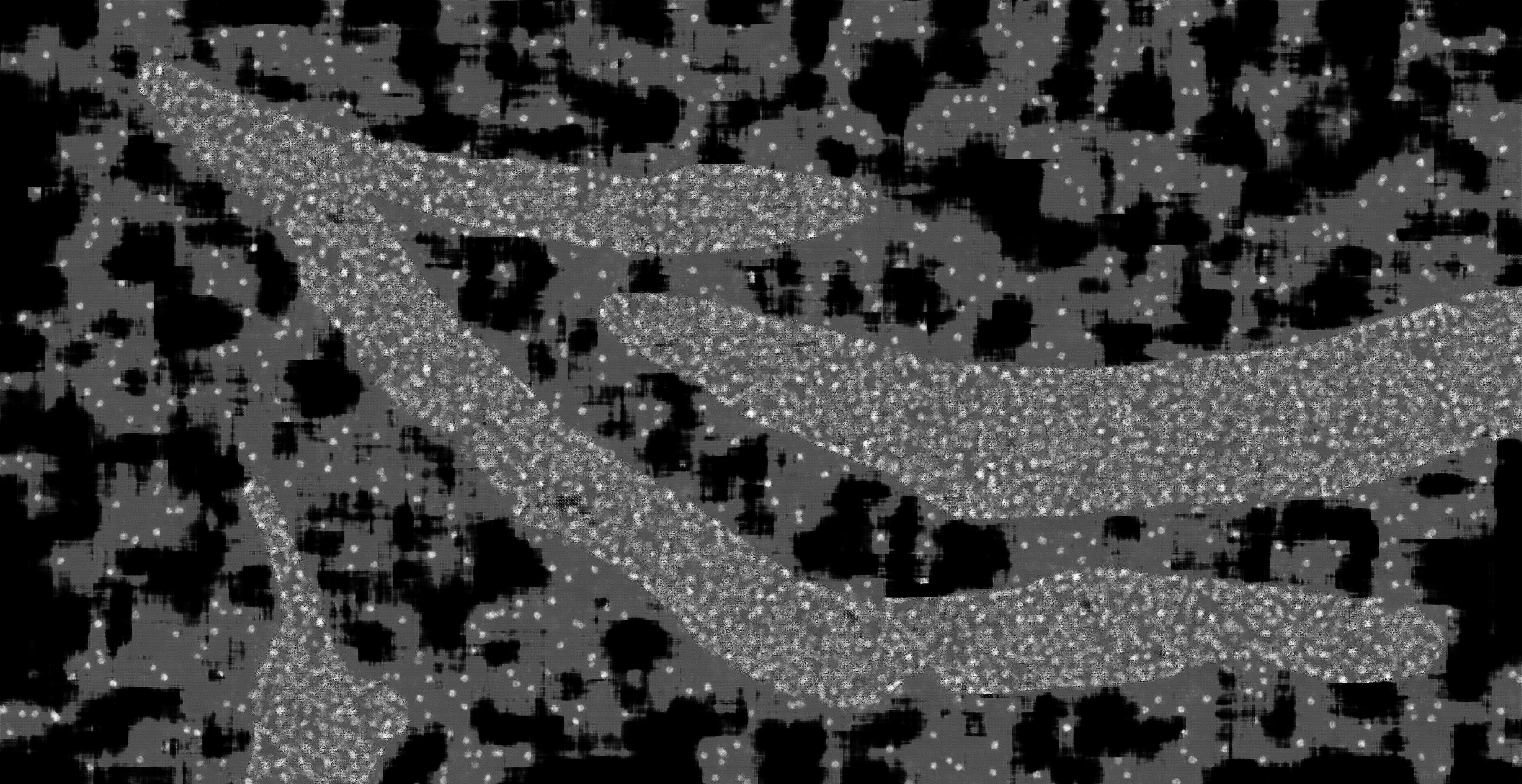

### hc_more_test_mask2dapi_002a_px_FS_16_KS_3_ML_False_4.jpg

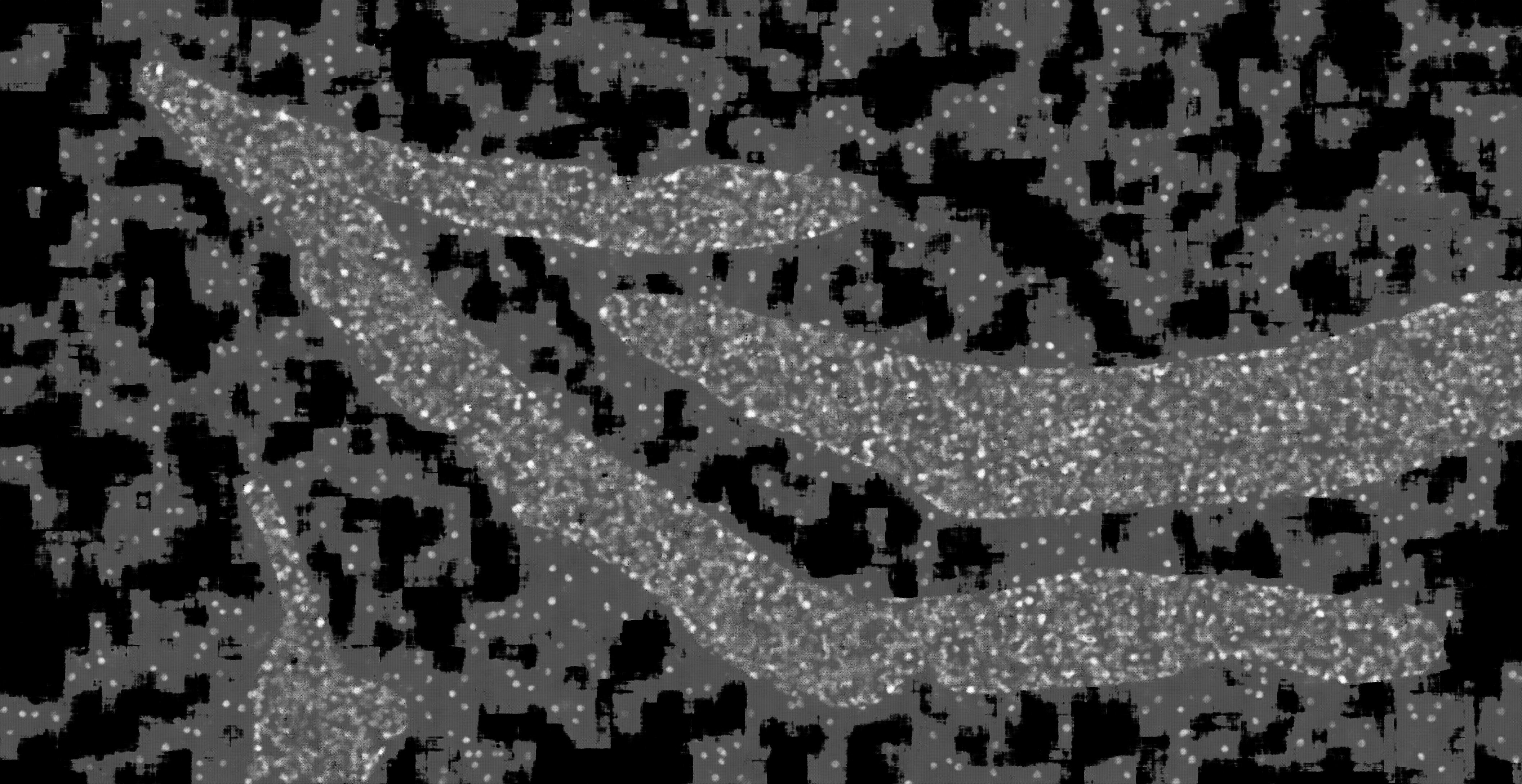

### hc_more_test_mask2dapi_002a_px_FS_16_KS_5_ML_True_4.jpg

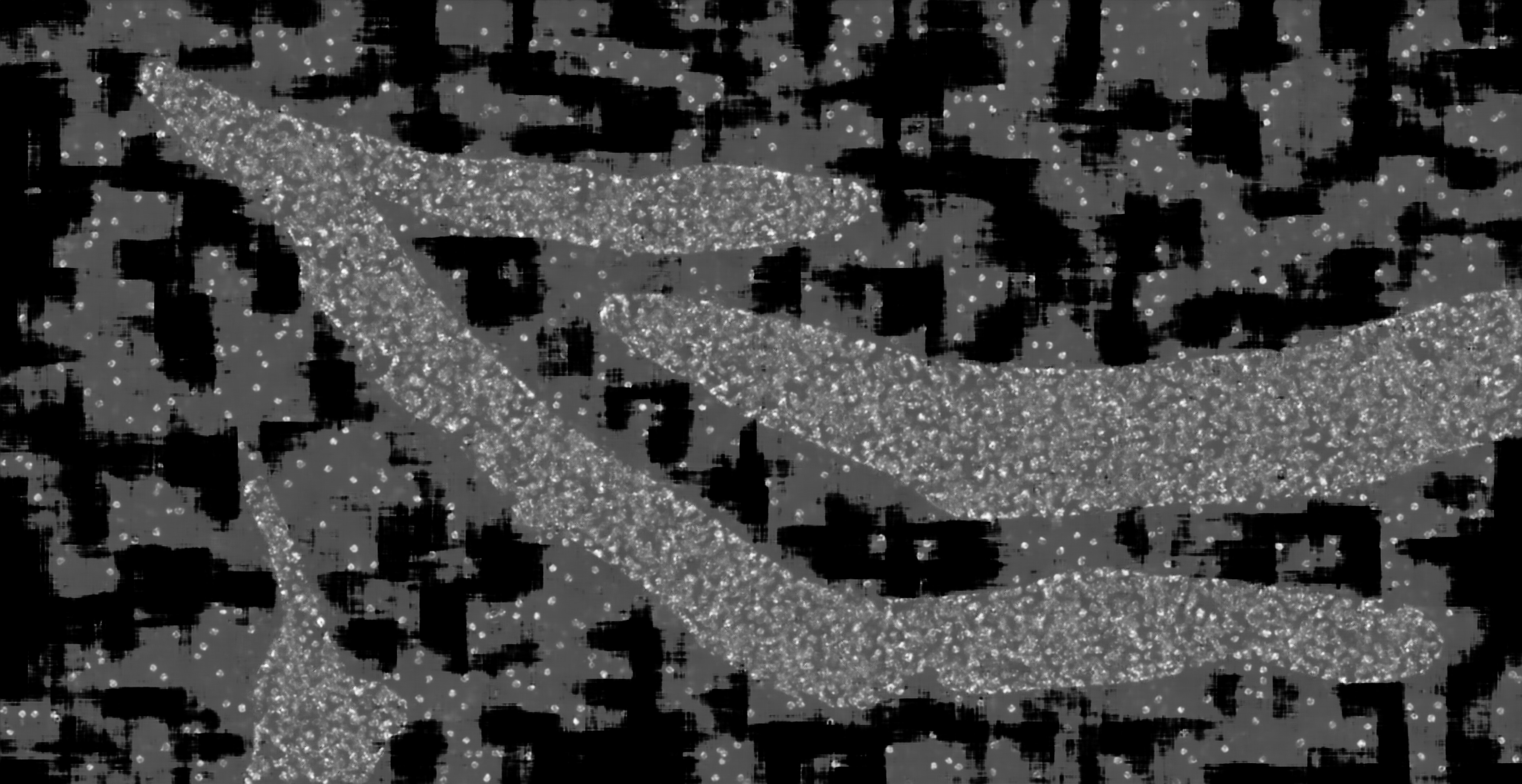

### hc_more_test_mask2dapi_002b_px_FS_16_KS_3_ML_True_4.jpg

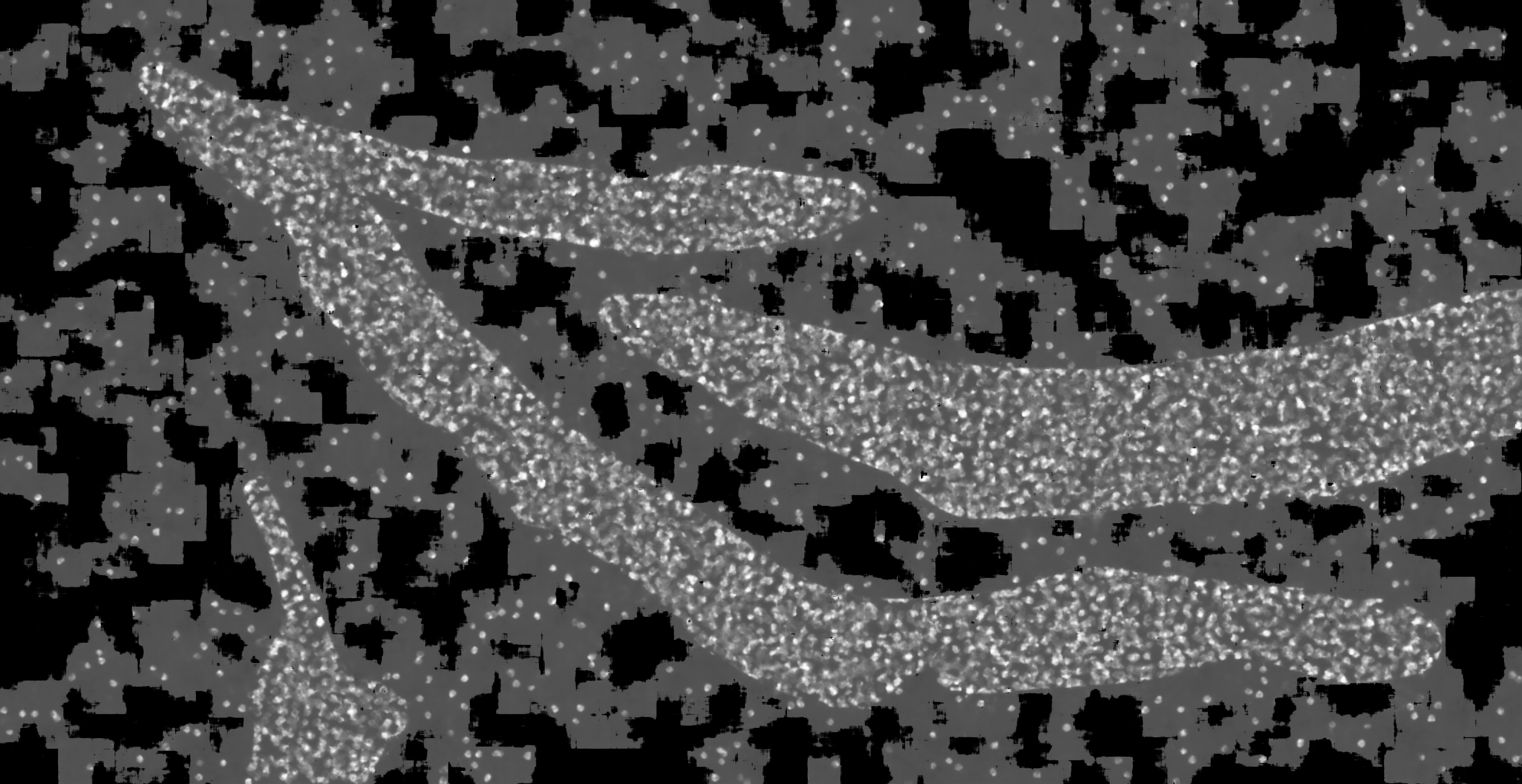

### hc_more_test_mask2dapi_002b_px_FS_16_KS_5_ML_False_4.jpg

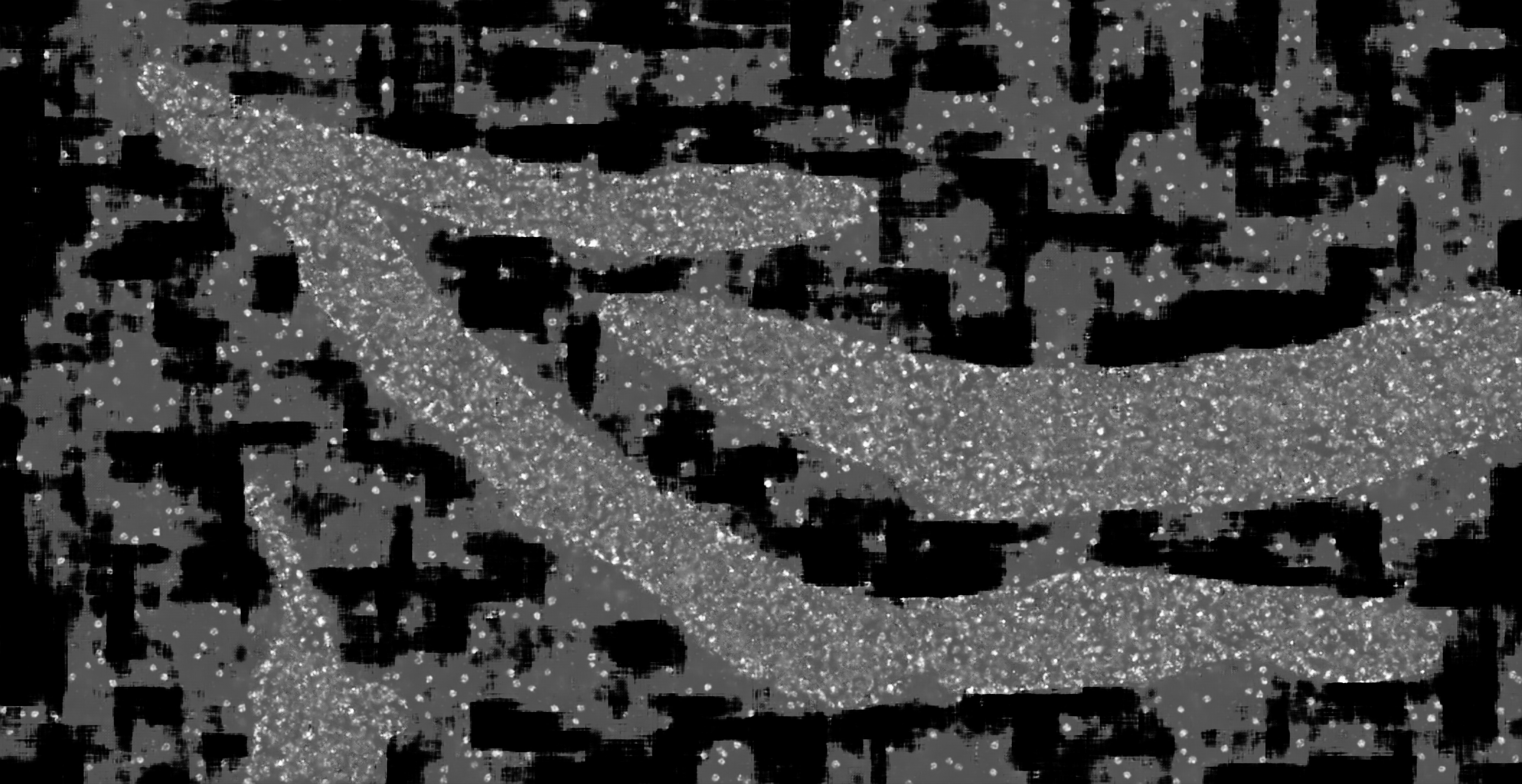

### hc_more_test_mask2dapi_002b_px_FS_16_KS_7_ML_True_4.jpg

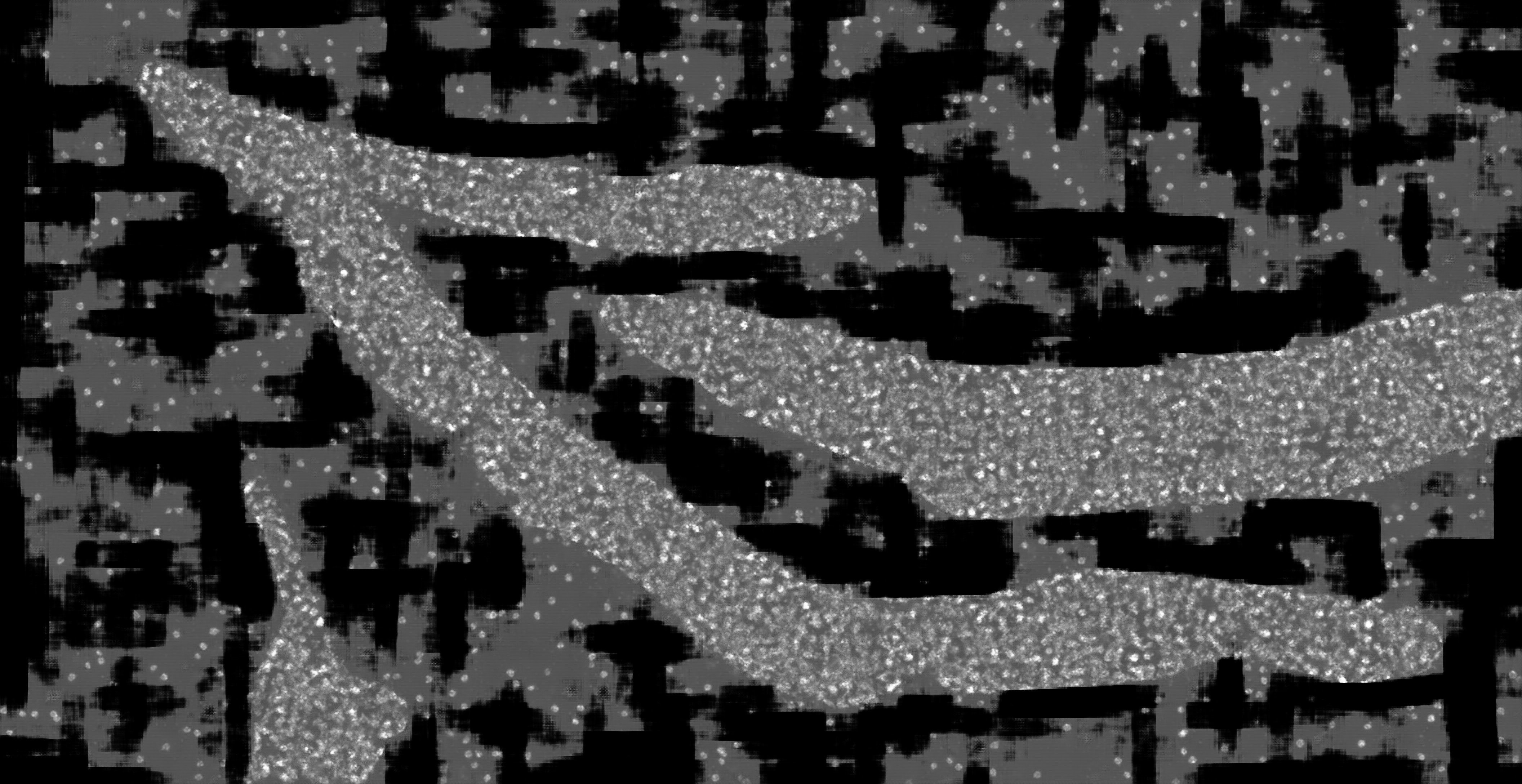

### hc_more_test_mask2dapi_003a_px_FS_16_KS_3_ML_True_4.jpg

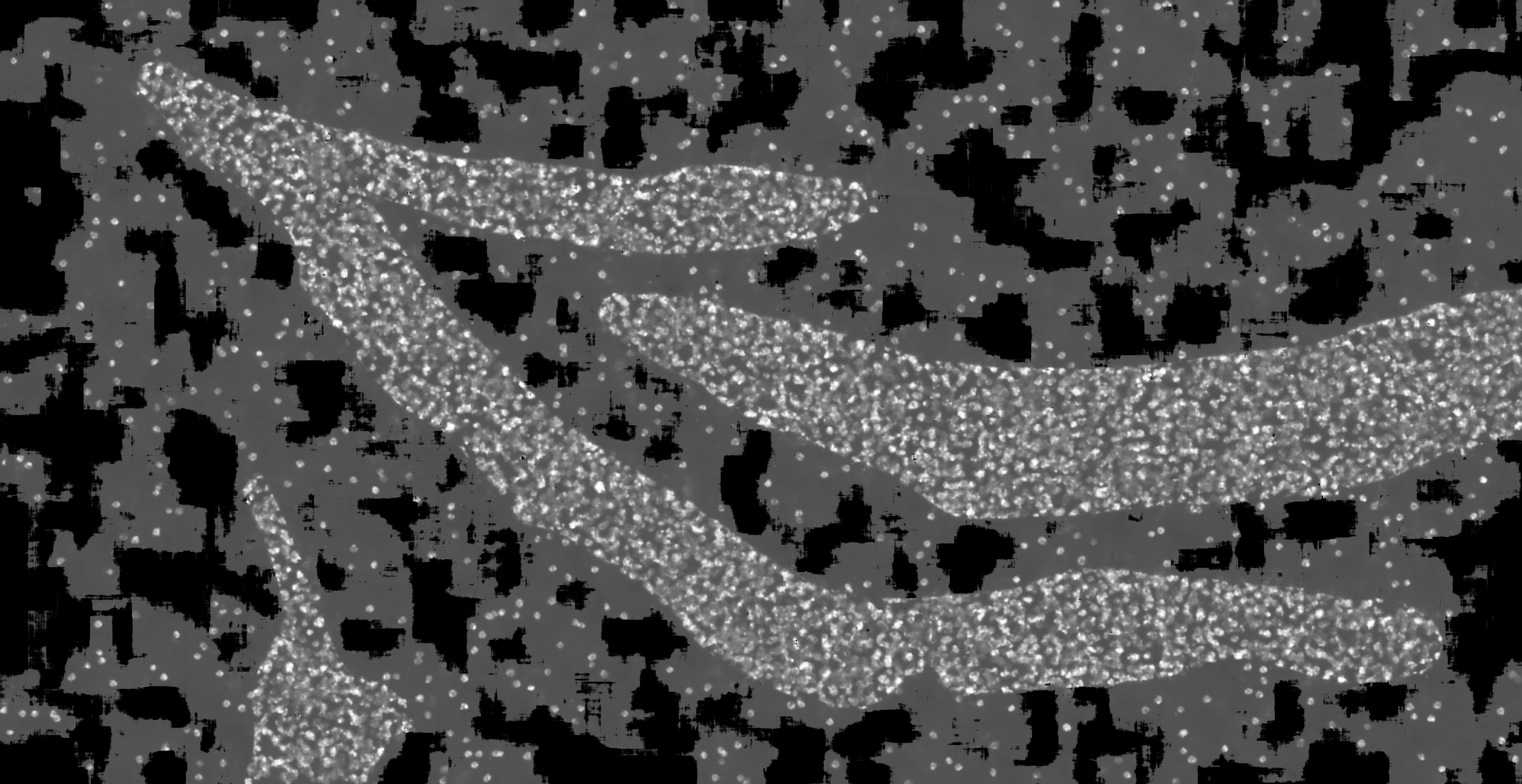

### hc_more_test_mask2dapi_003a_px_FS_16_KS_5_ML_True_4.jpg

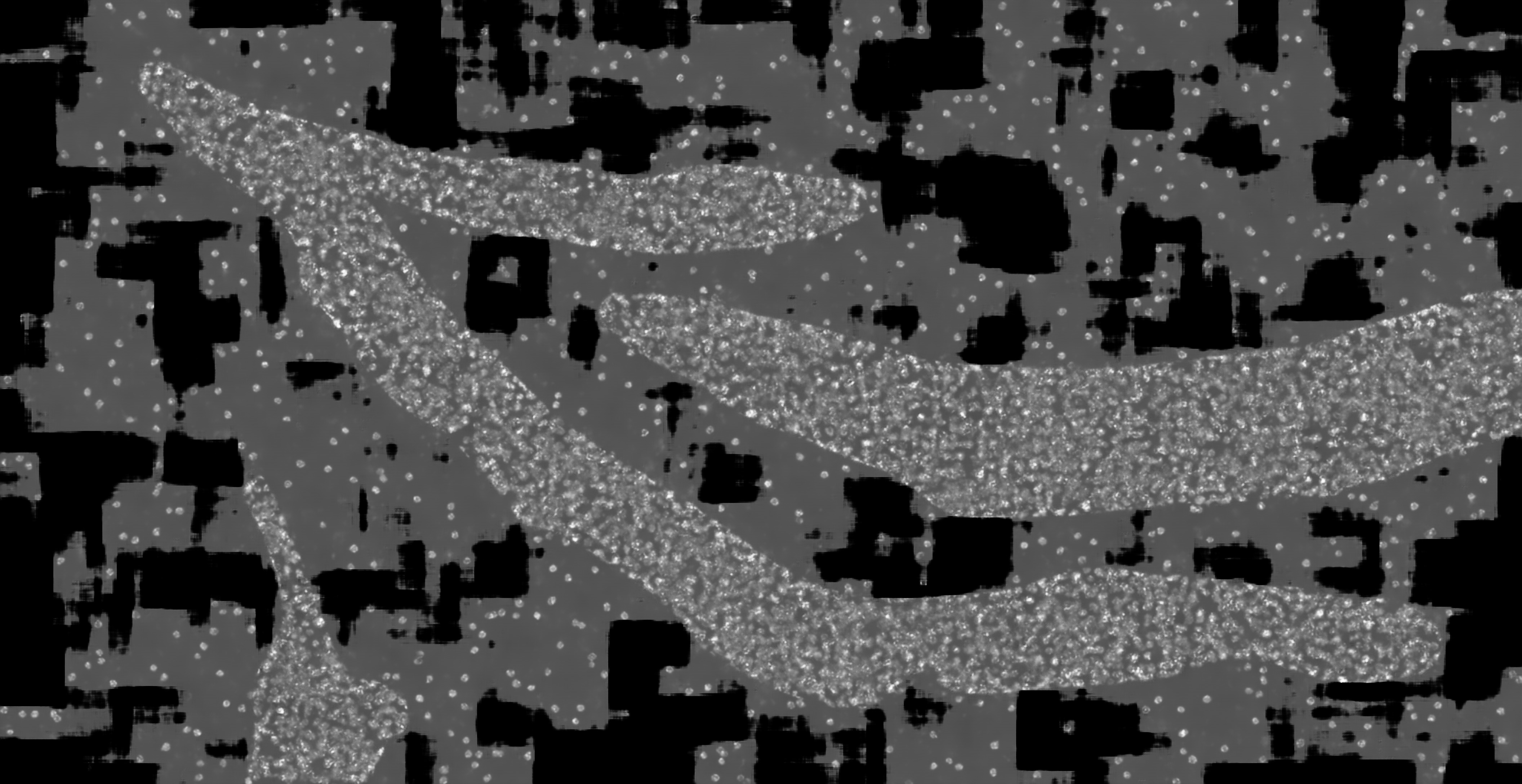

### hc_more_test_mask2dapi_003a_px_FS_16_KS_7_ML_False_4.jpg

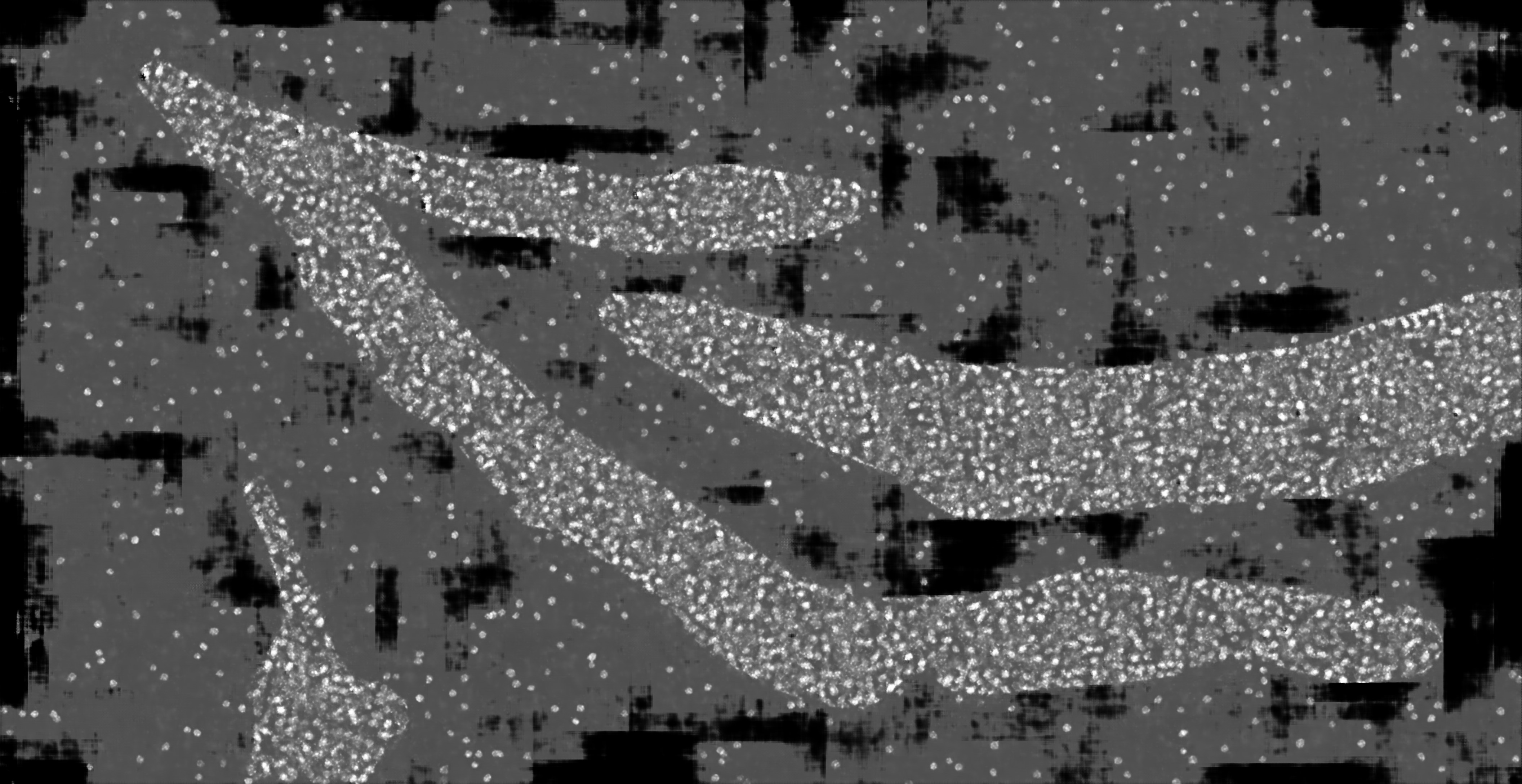

### hc_more_test_mask2dapi_003a_px_FS_16_KS_7_ML_True_4.jpg

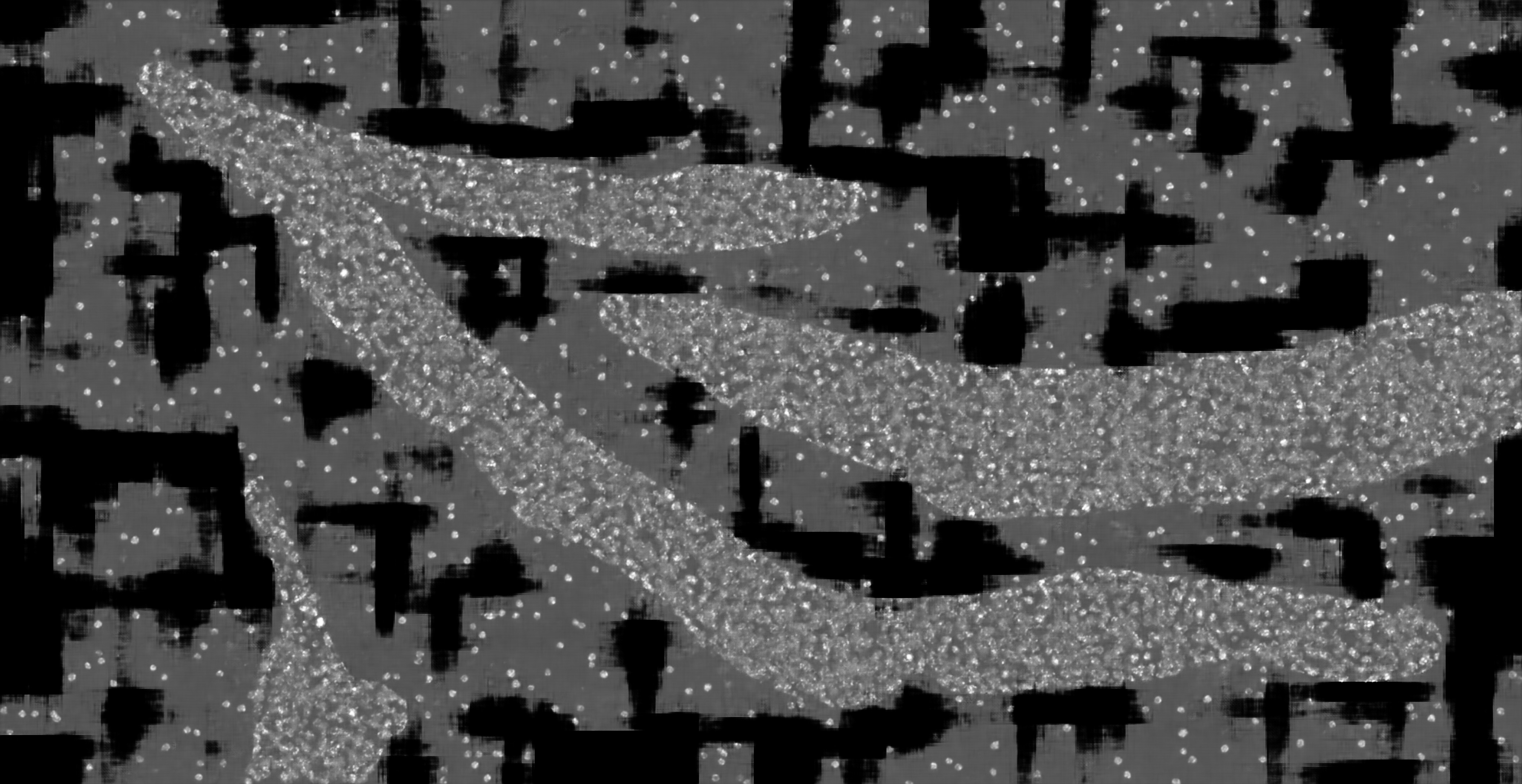

### hc_more_test_mask2dapi_003b_px_FS_16_KS_5_ML_True_4.jpg

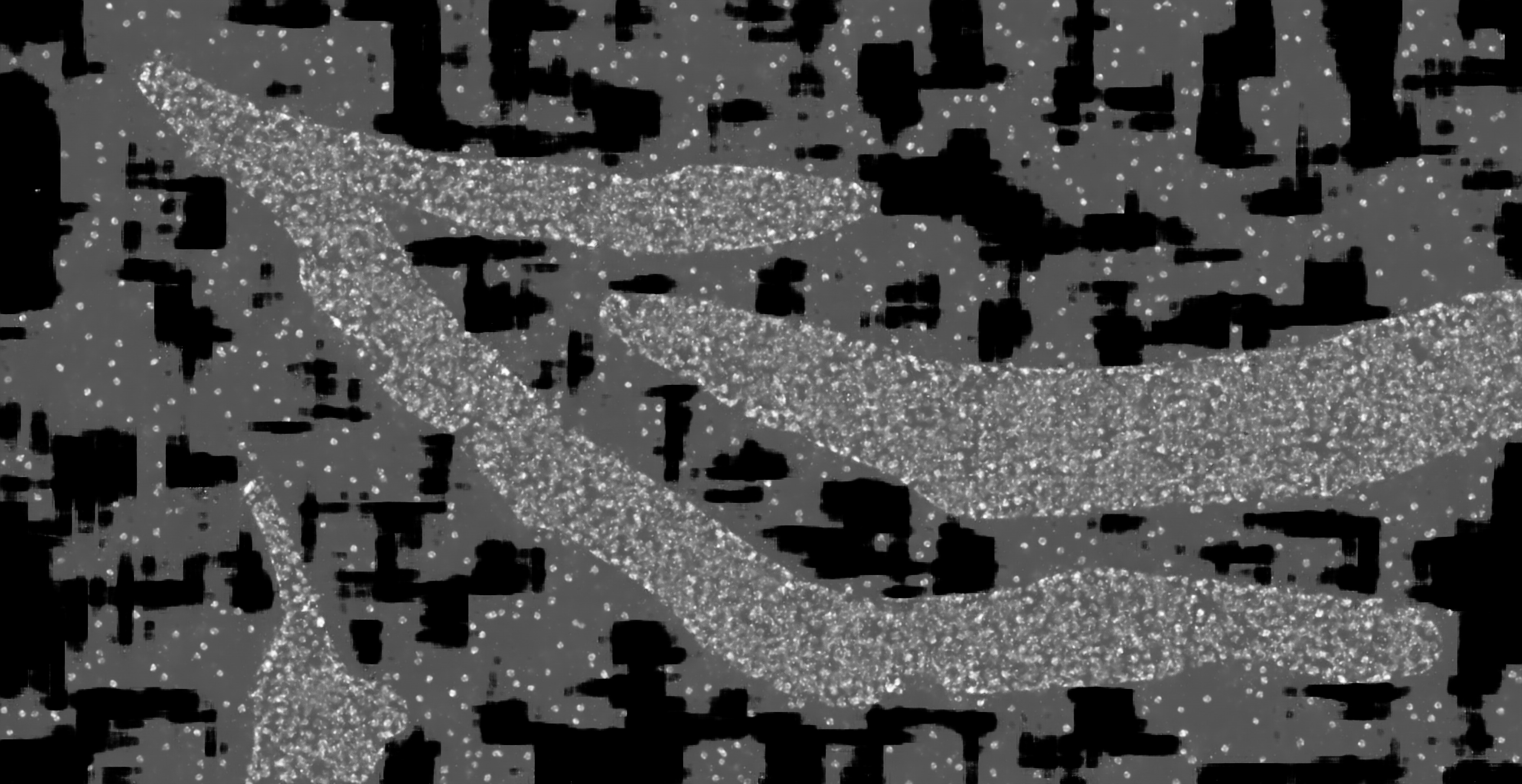

### hc_test_mask2dapi_001_px_FS_16_KS_3_ML_False_4.jpg

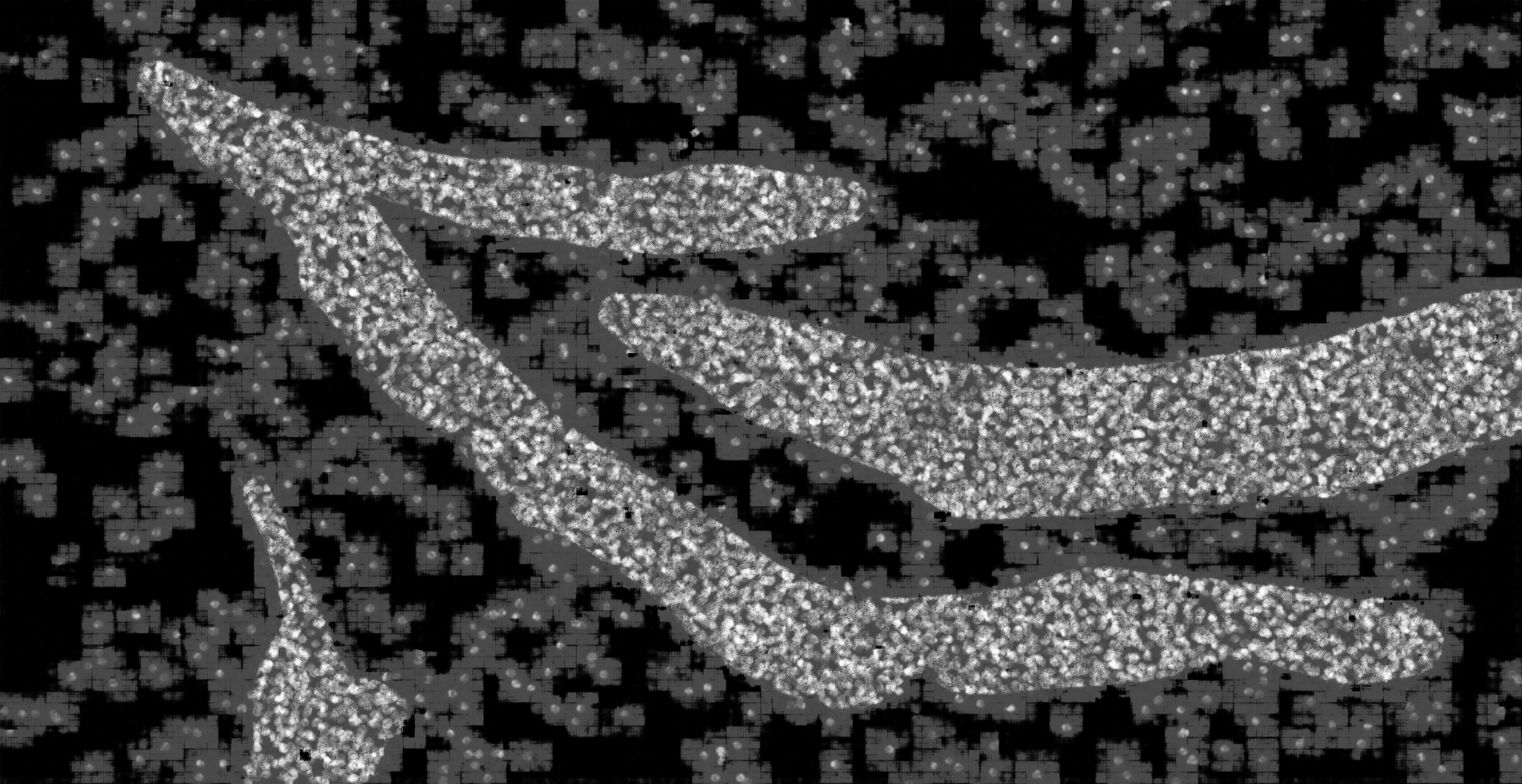

### hc_test_mask2dapi_001_px_FS_16_KS_3_ML_True_4.jpg

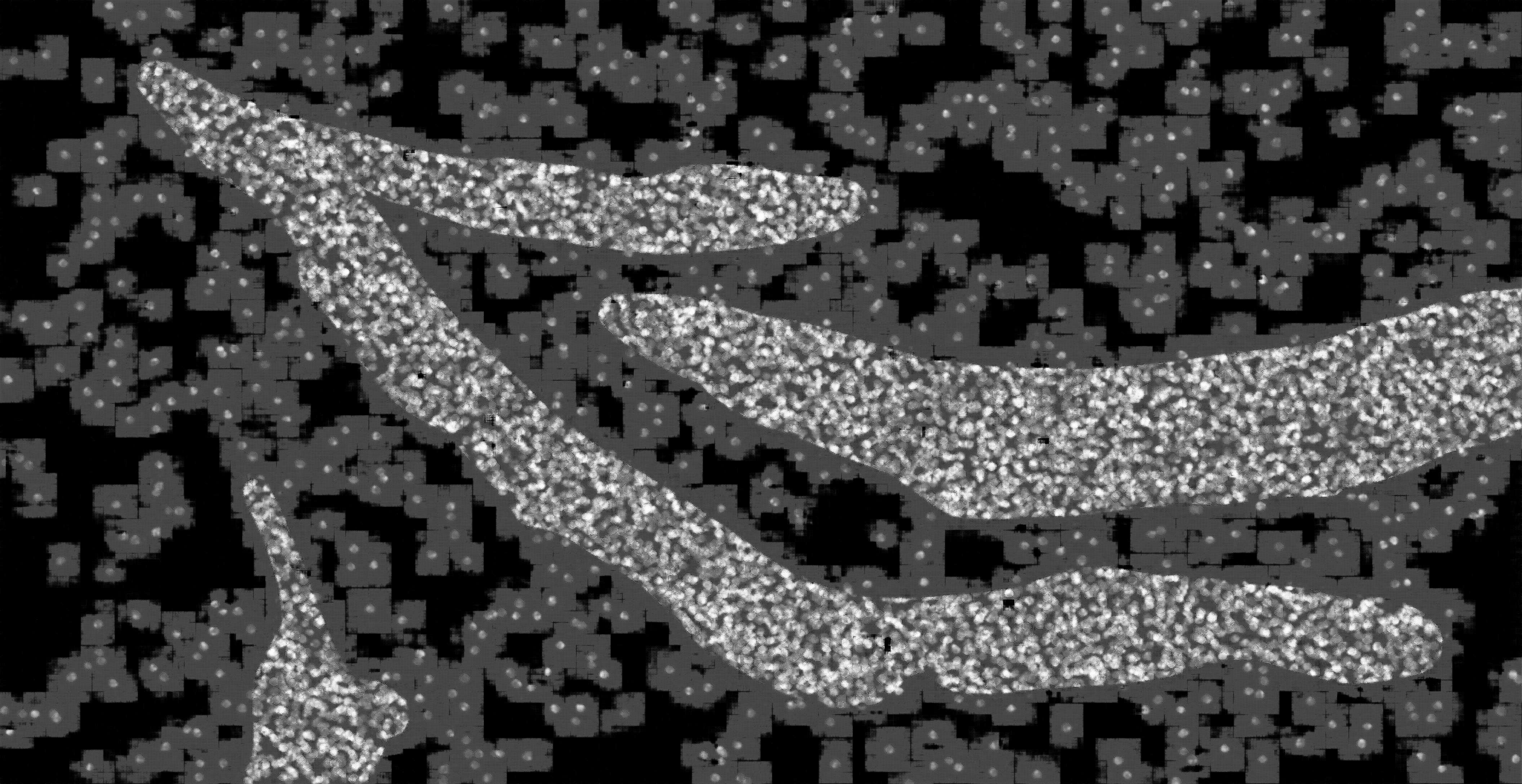

### hc_test_mask2dapi_001_px_FS_16_KS_5_ML_False_4.jpg

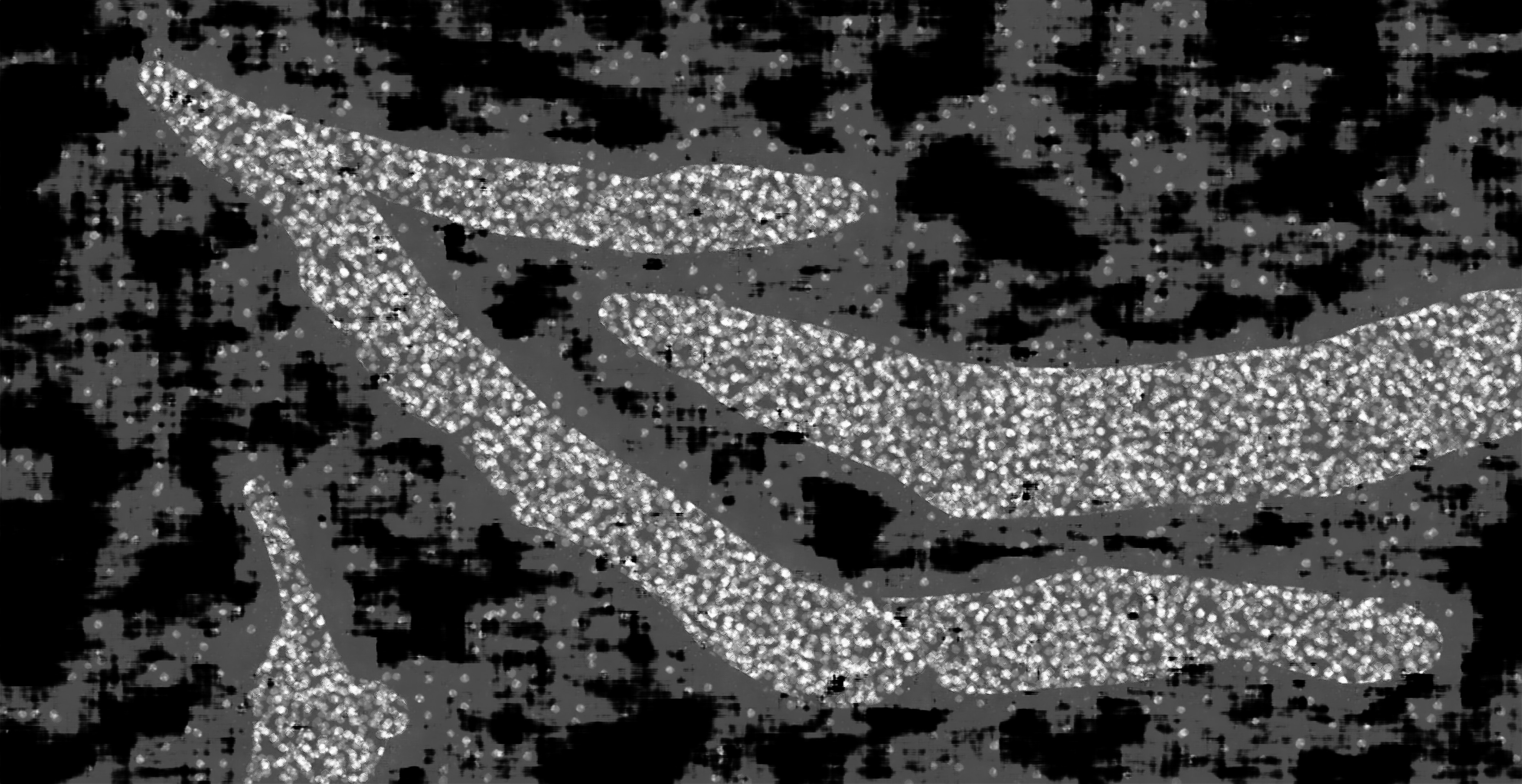

### hc_test_mask2dapi_001_px_FS_16_KS_5_ML_True_4.jpg

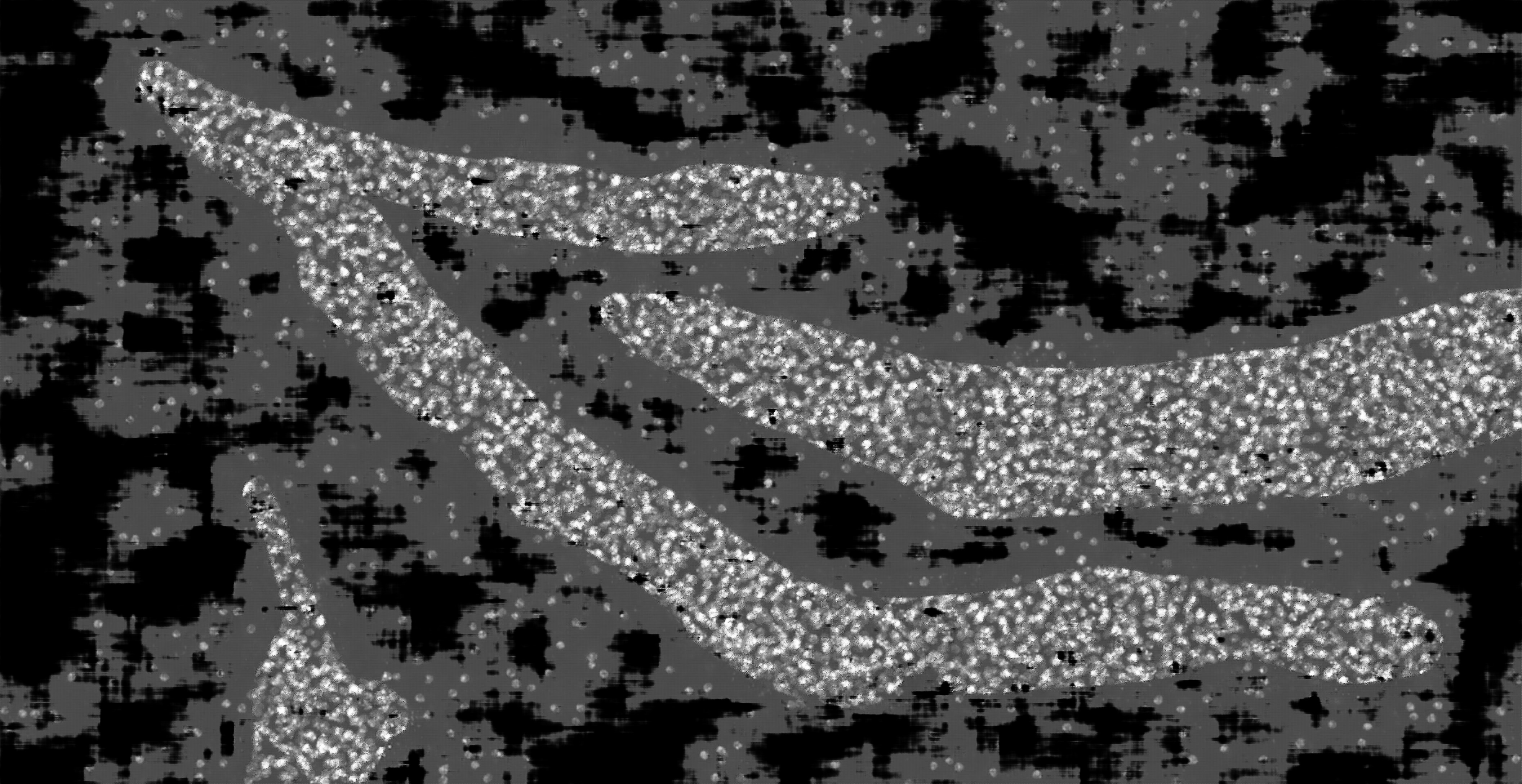

### hc_test_mask2dapi_001_px_FS_16_KS_7_ML_False_4.jpg

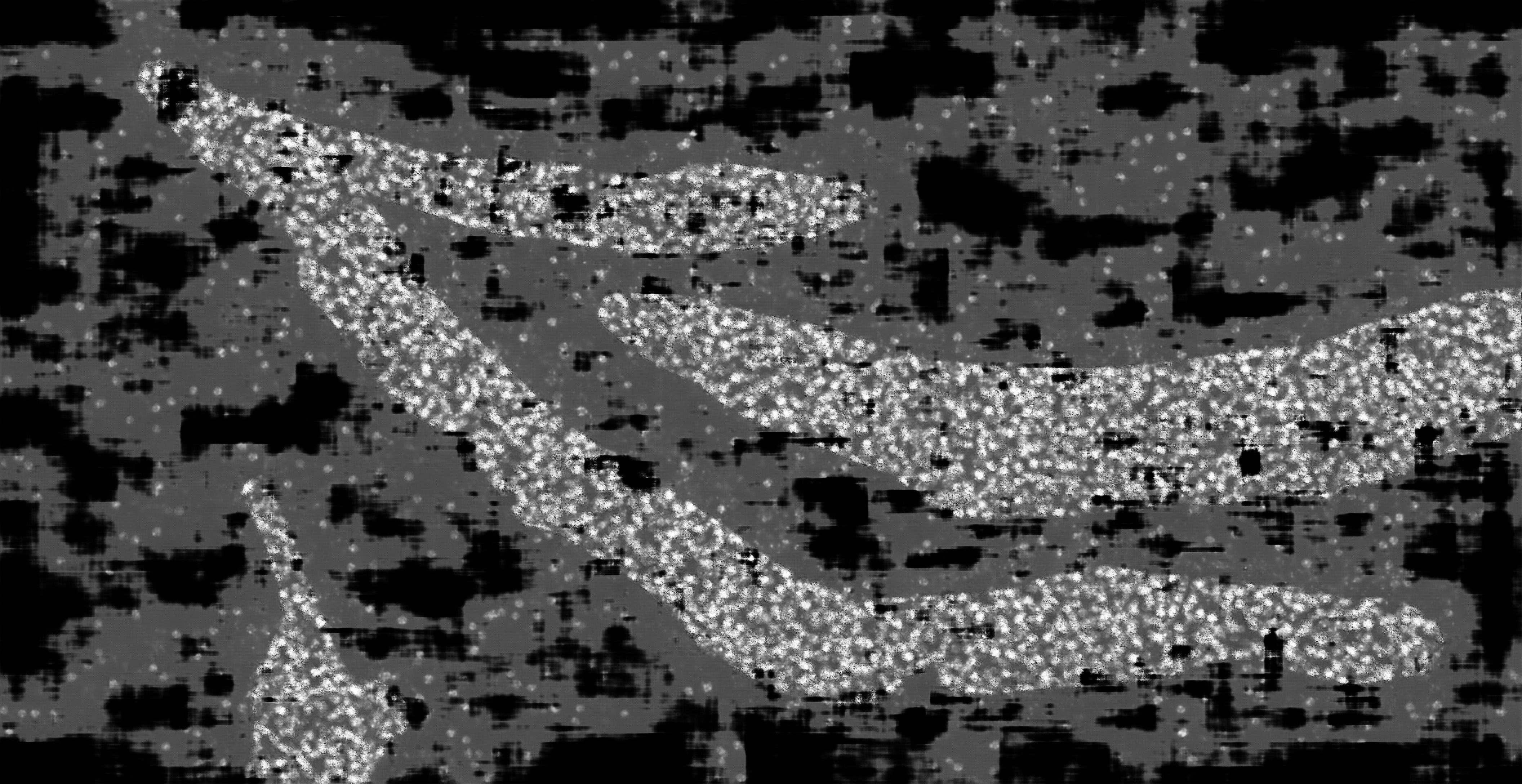

### hc_test_mask2dapi_001_px_FS_16_KS_7_ML_True_4.jpg

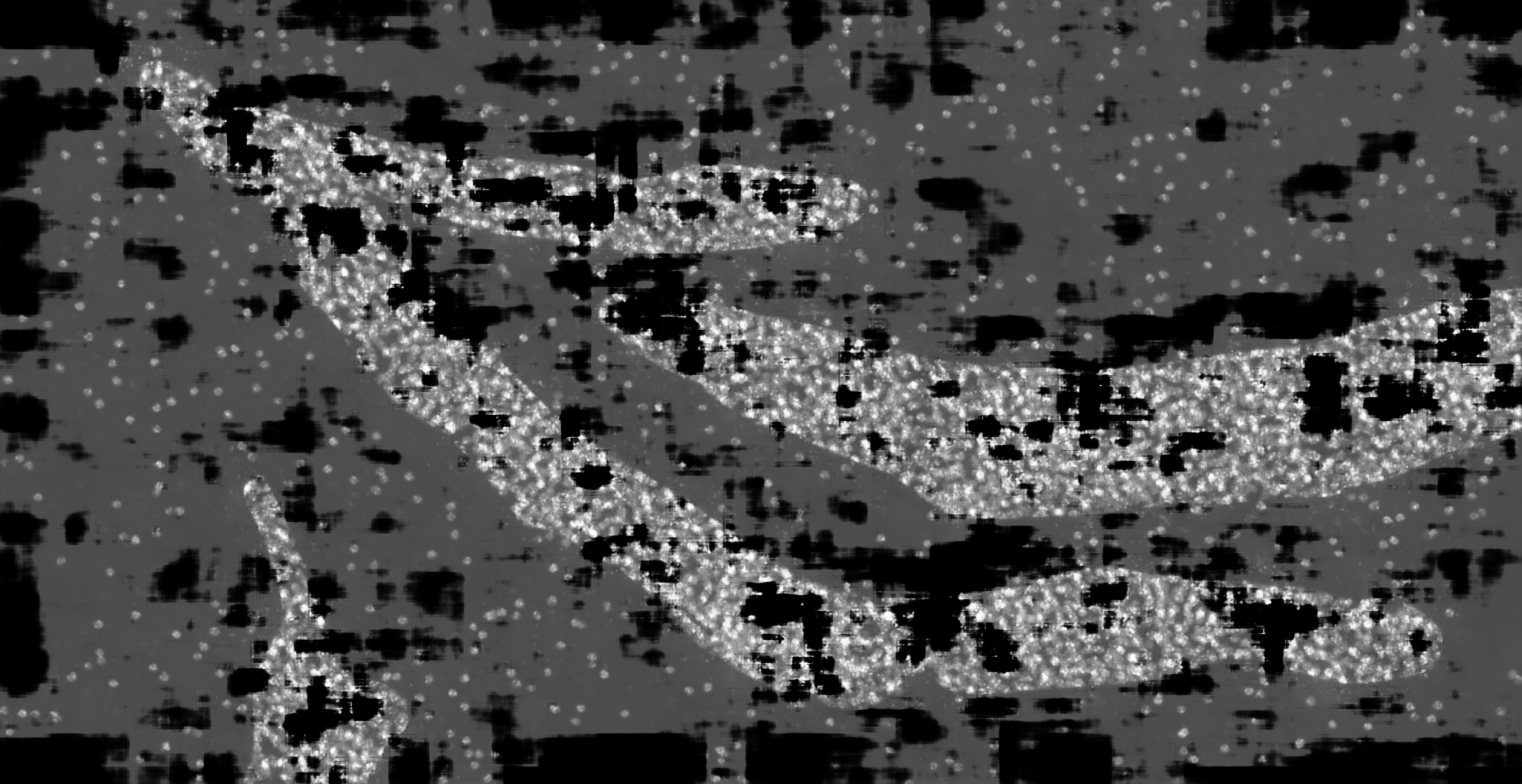

### hc_test_mask2dapi_001_px_FS_32_KS_3_ML_False_4.jpg

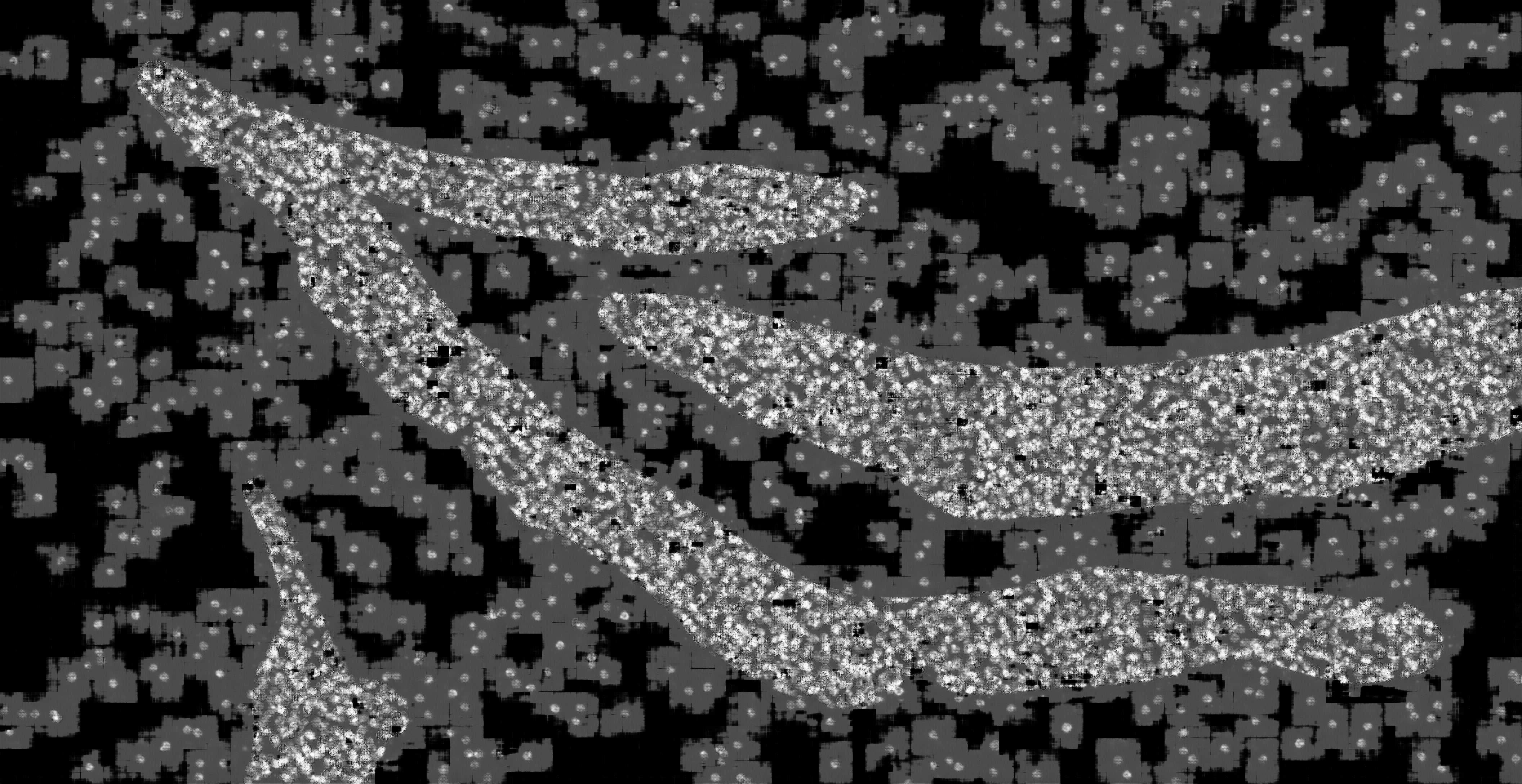

### hc_test_mask2dapi_001_px_FS_32_KS_3_ML_True_4.jpg

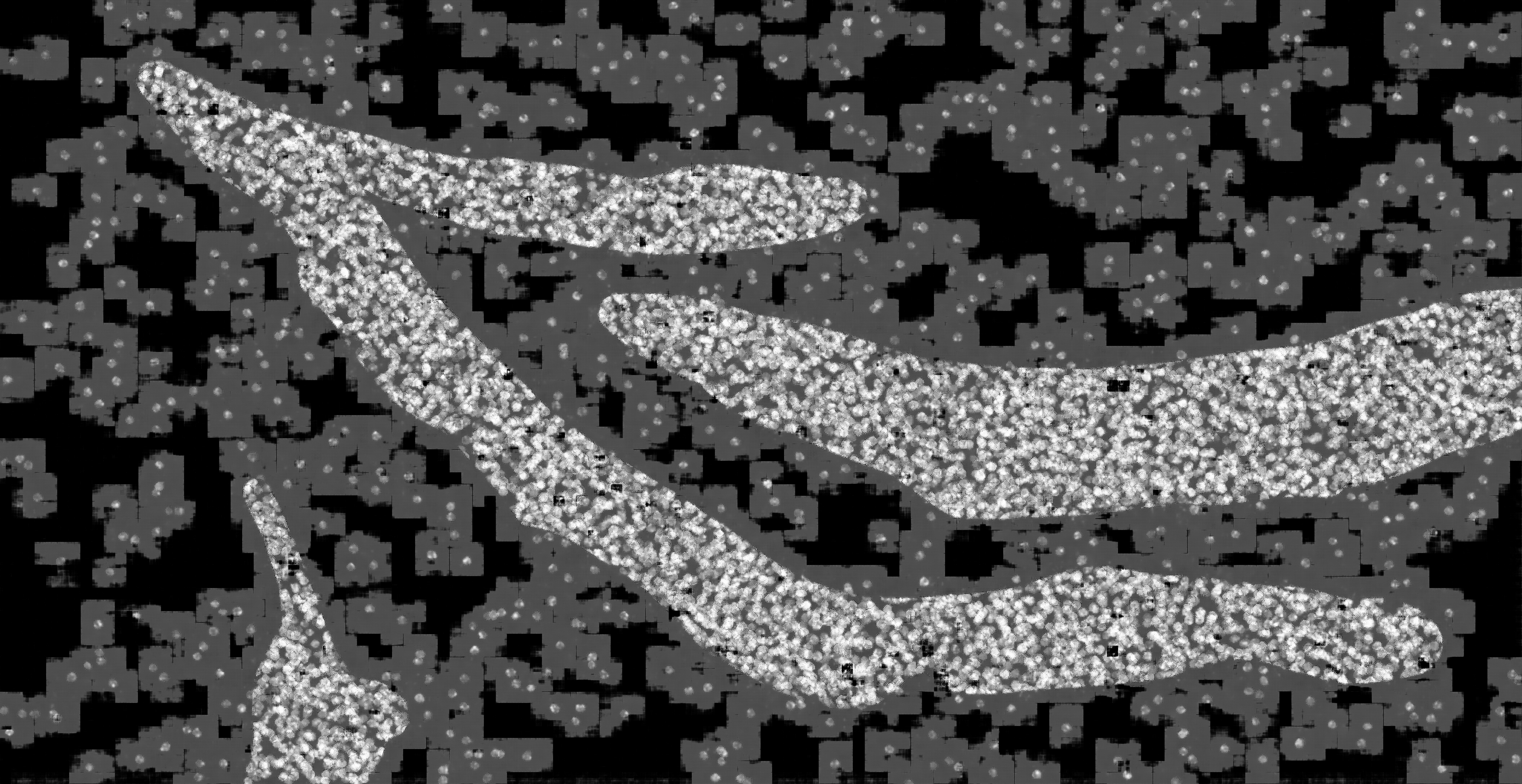

### hc_test_mask2dapi_001_px_FS_32_KS_5_ML_False_4.jpg

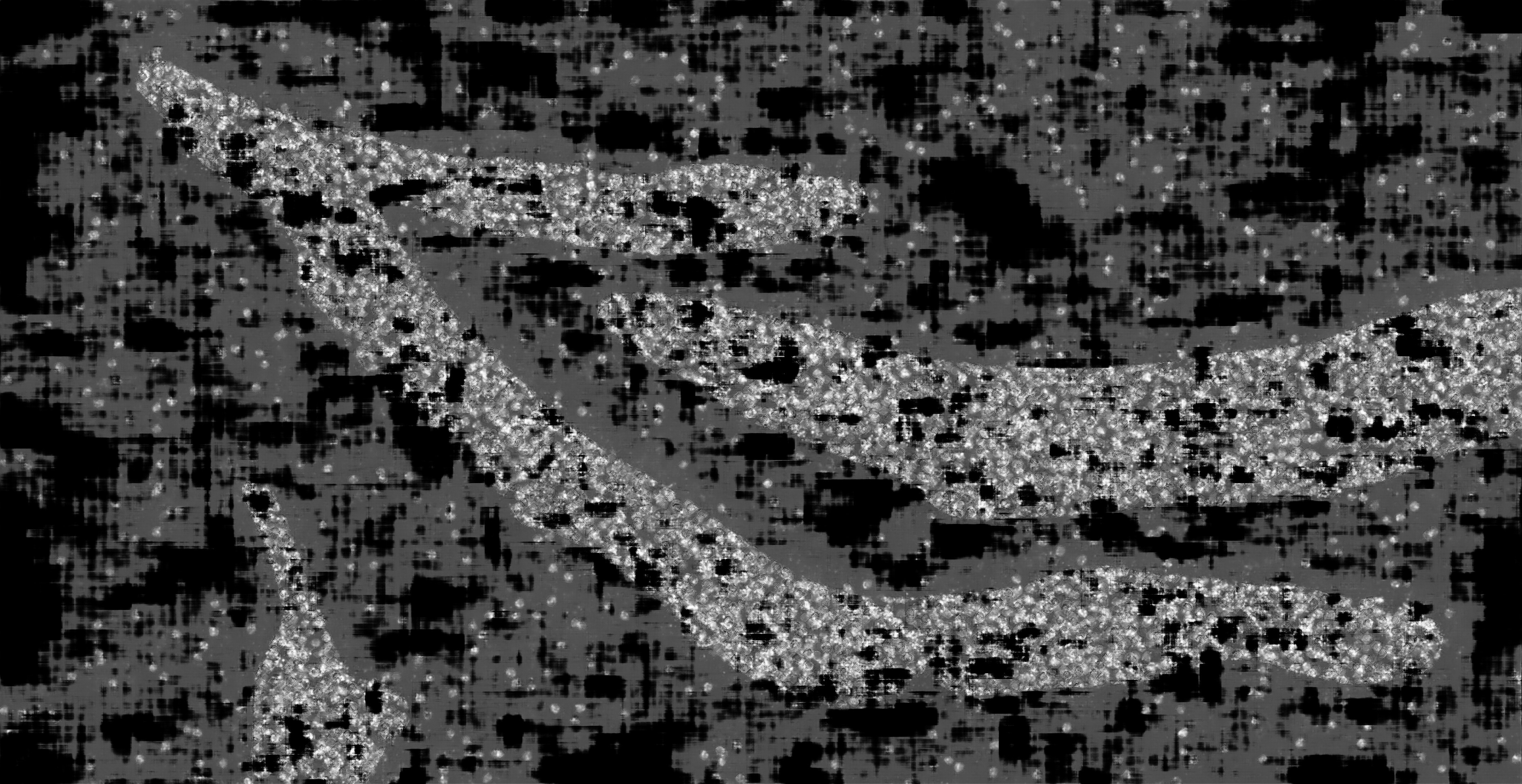

### hc_test_mask2dapi_001_px_FS_32_KS_5_ML_True_4.jpg

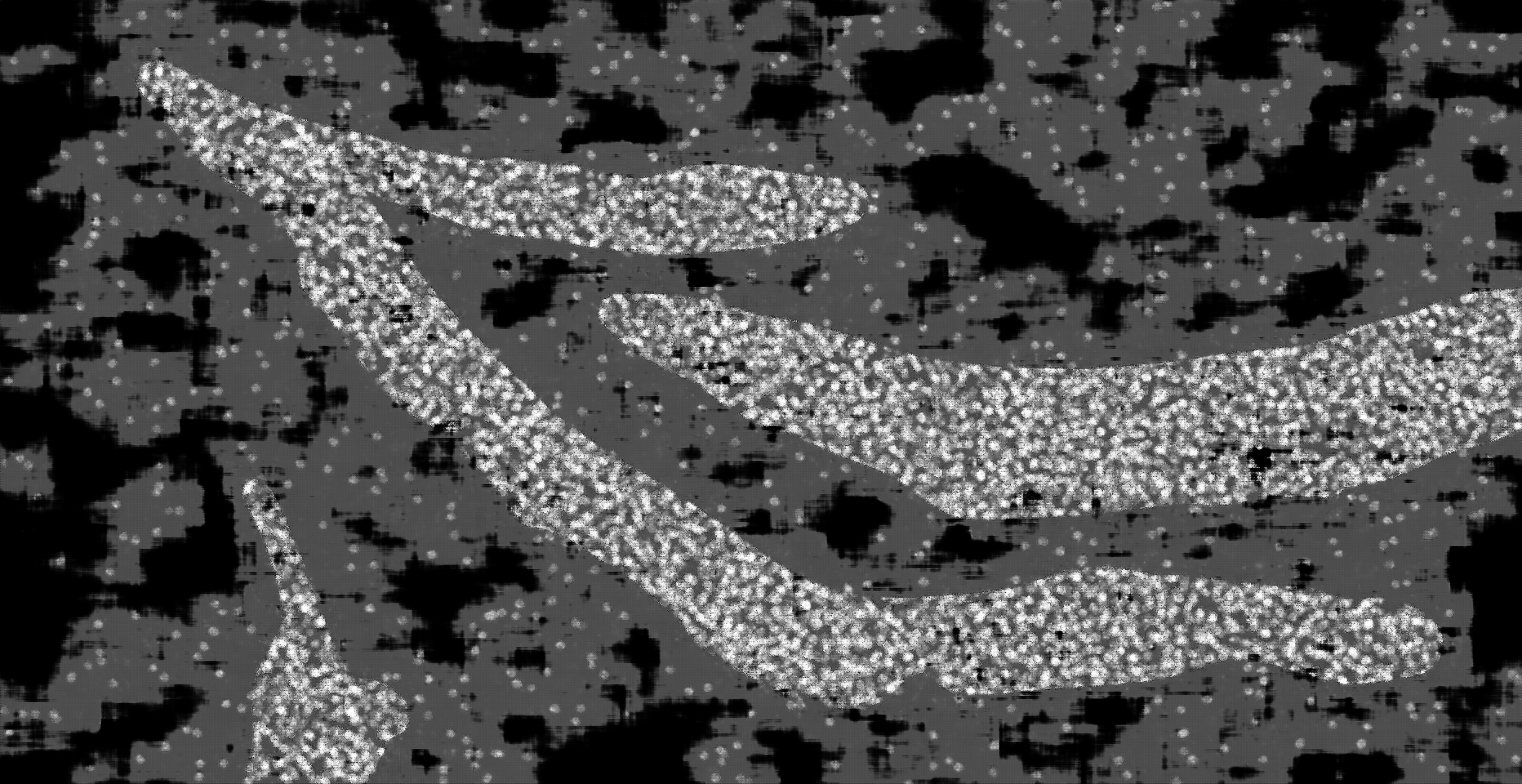

### hc_test_mask2dapi_001_px_FS_32_KS_7_ML_False_4.jpg

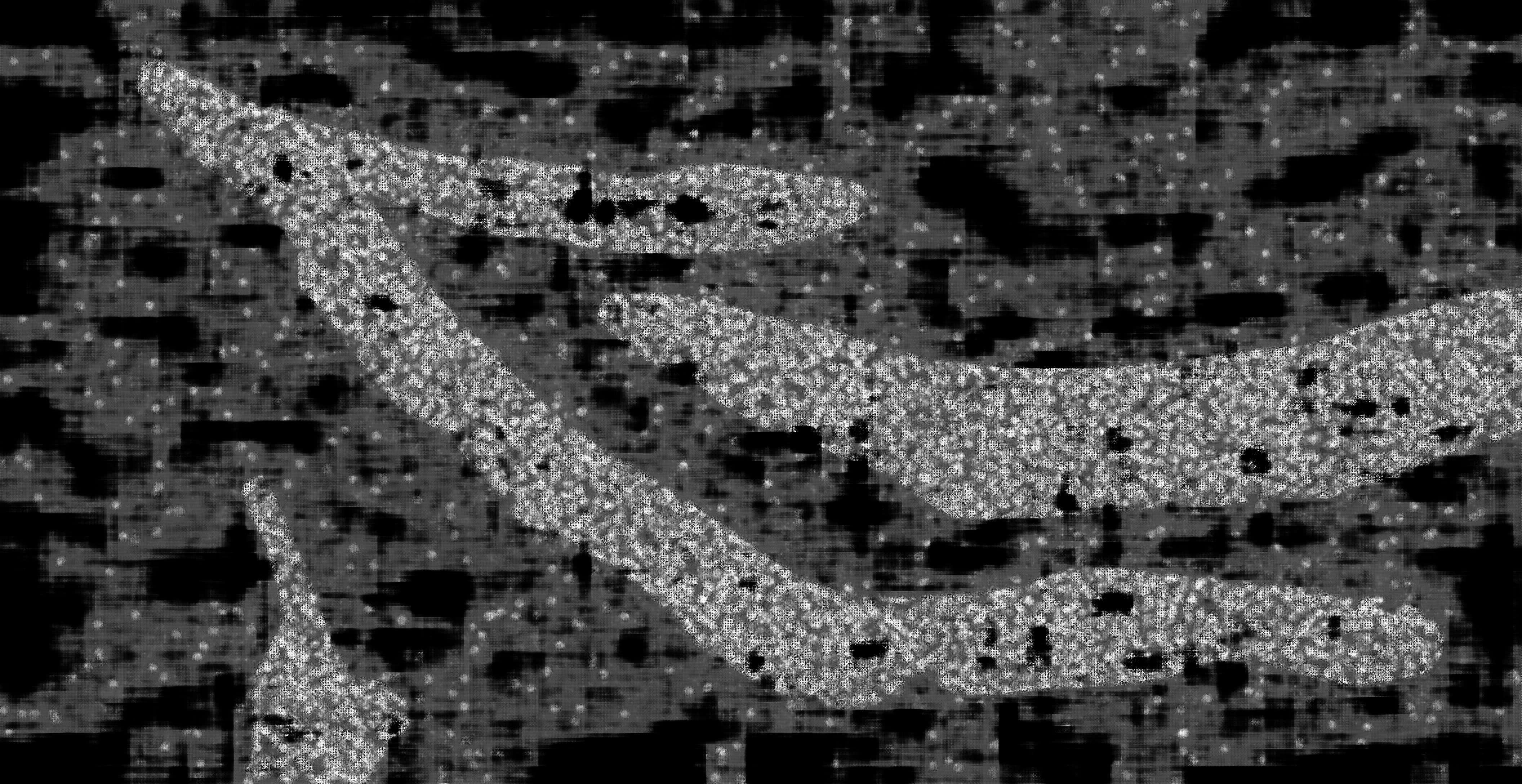

### hc_test_mask2dapi_001_px_FS_32_KS_7_ML_True_4.jpg

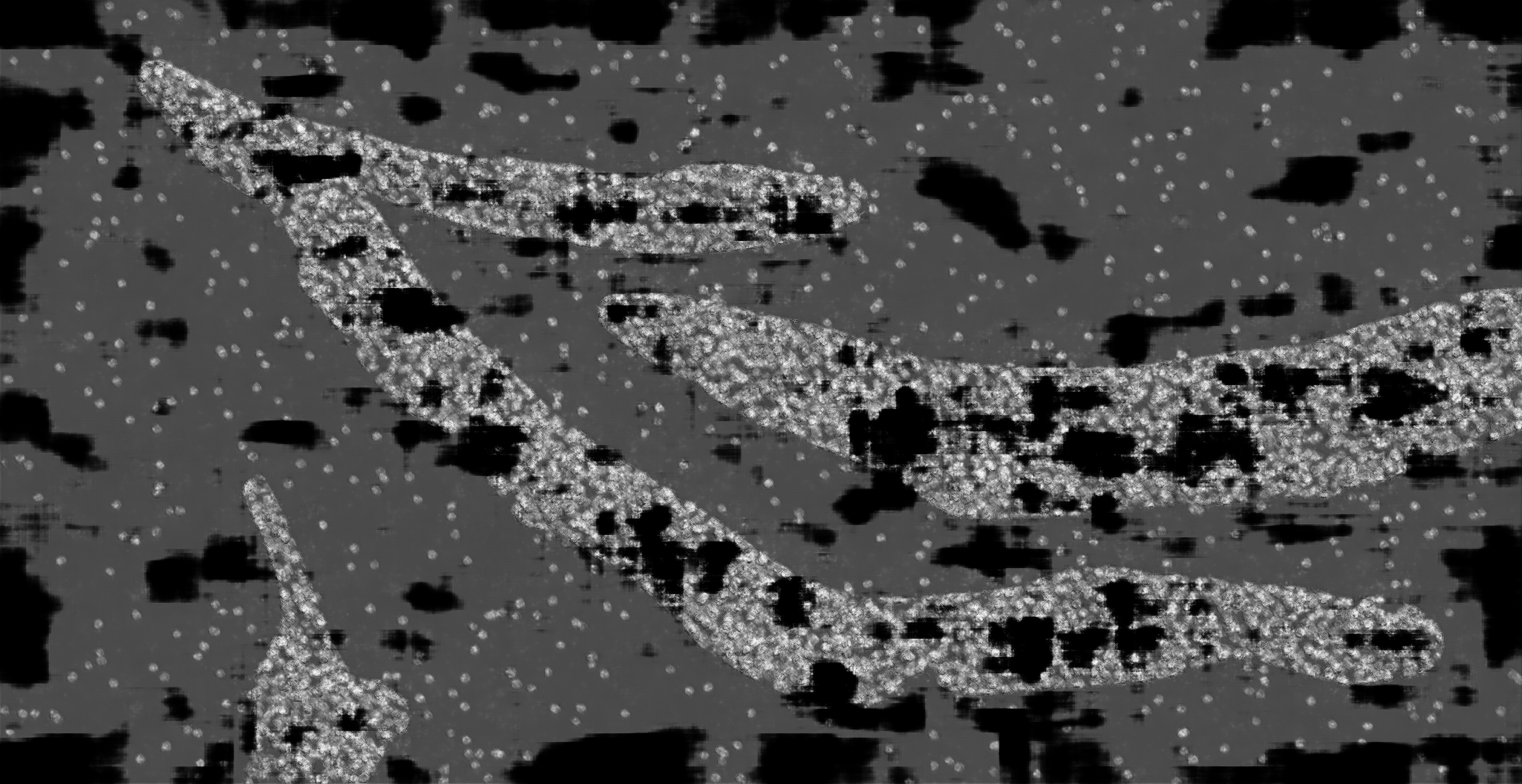

### hc_test_mask2dapi_001_px_FS_64_KS_3_ML_False_4.jpg

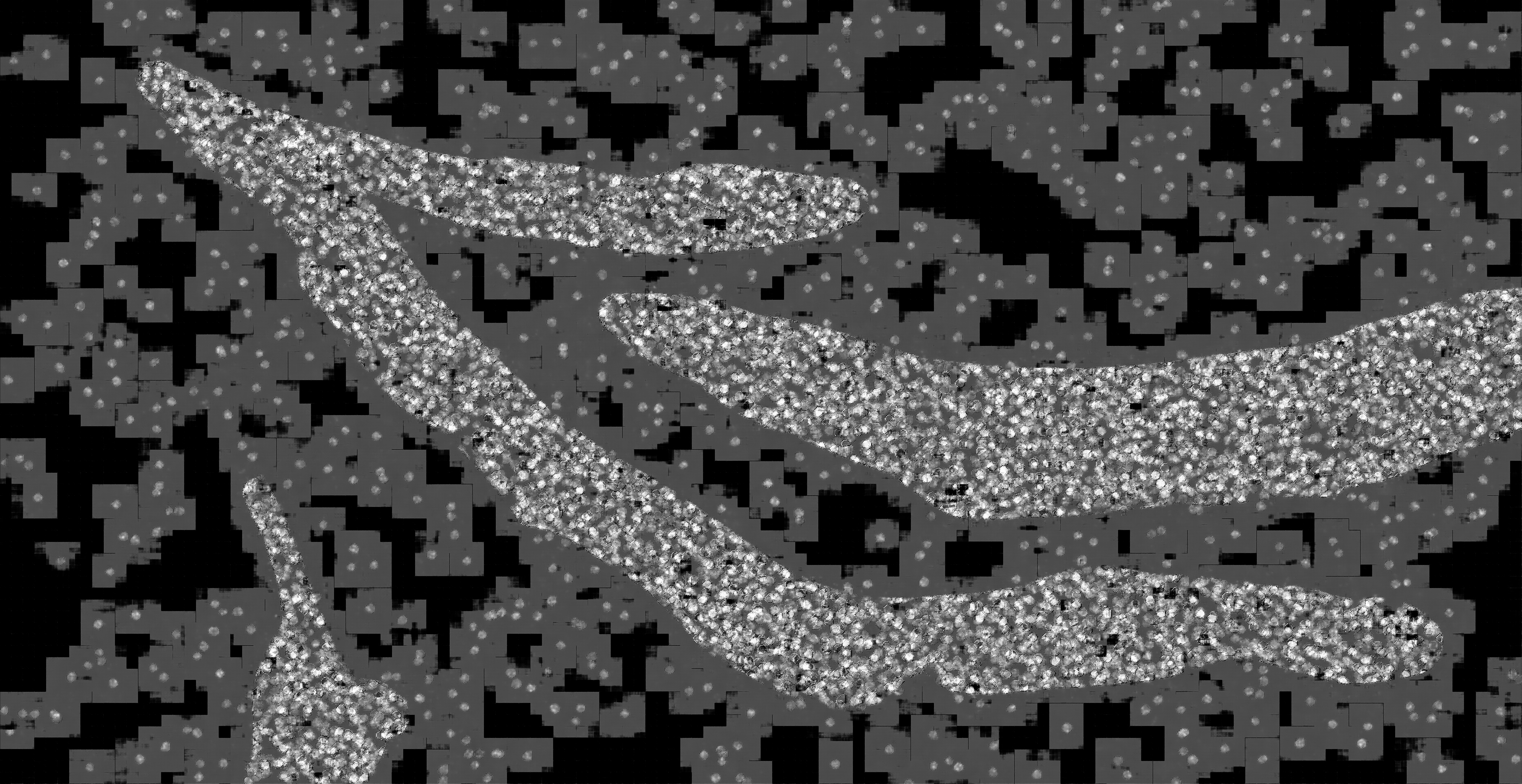

### hc_test_mask2dapi_001_px_FS_64_KS_3_ML_True_4.jpg

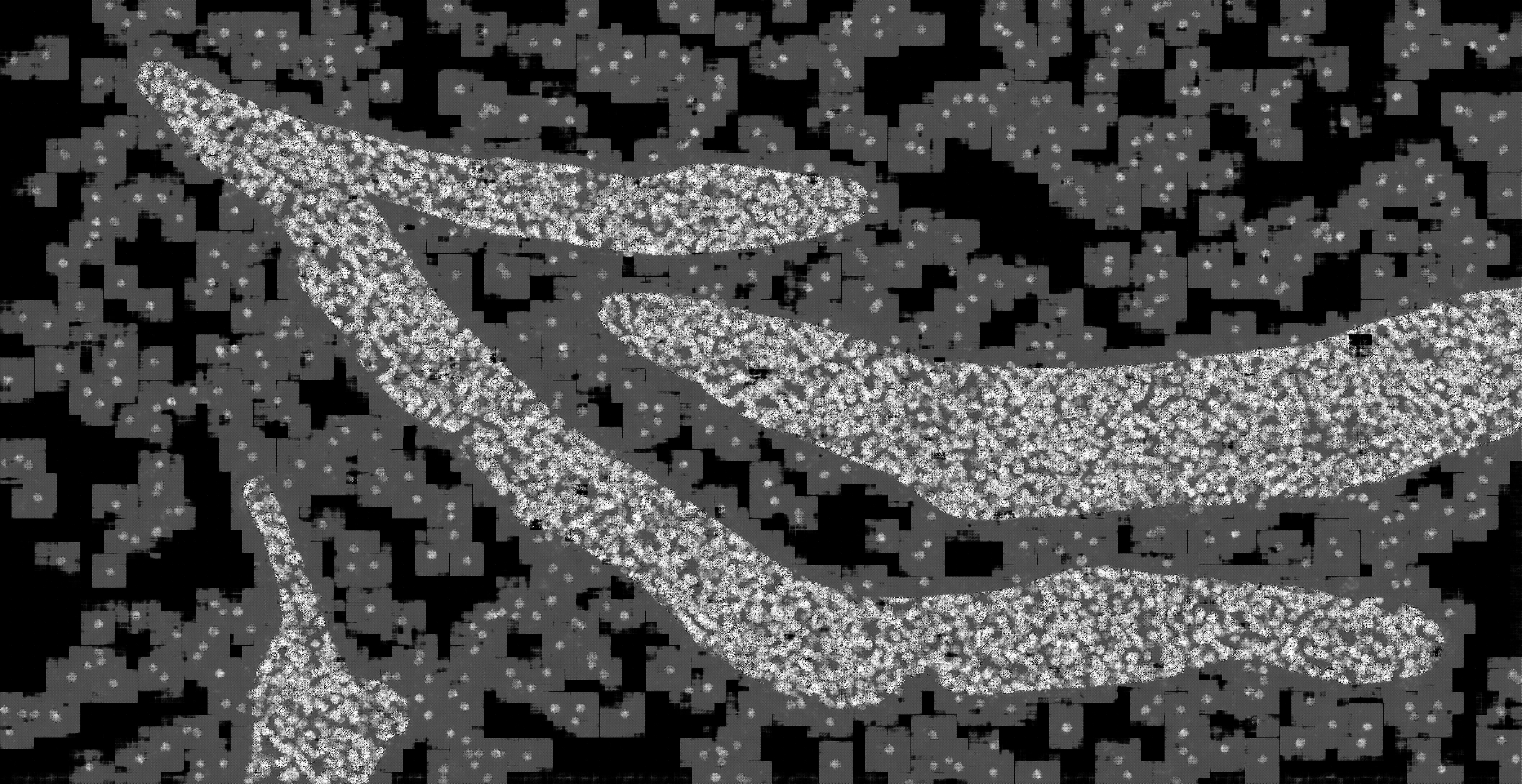

### hc_test_mask2dapi_001_px_FS_64_KS_5_ML_False_4.jpg

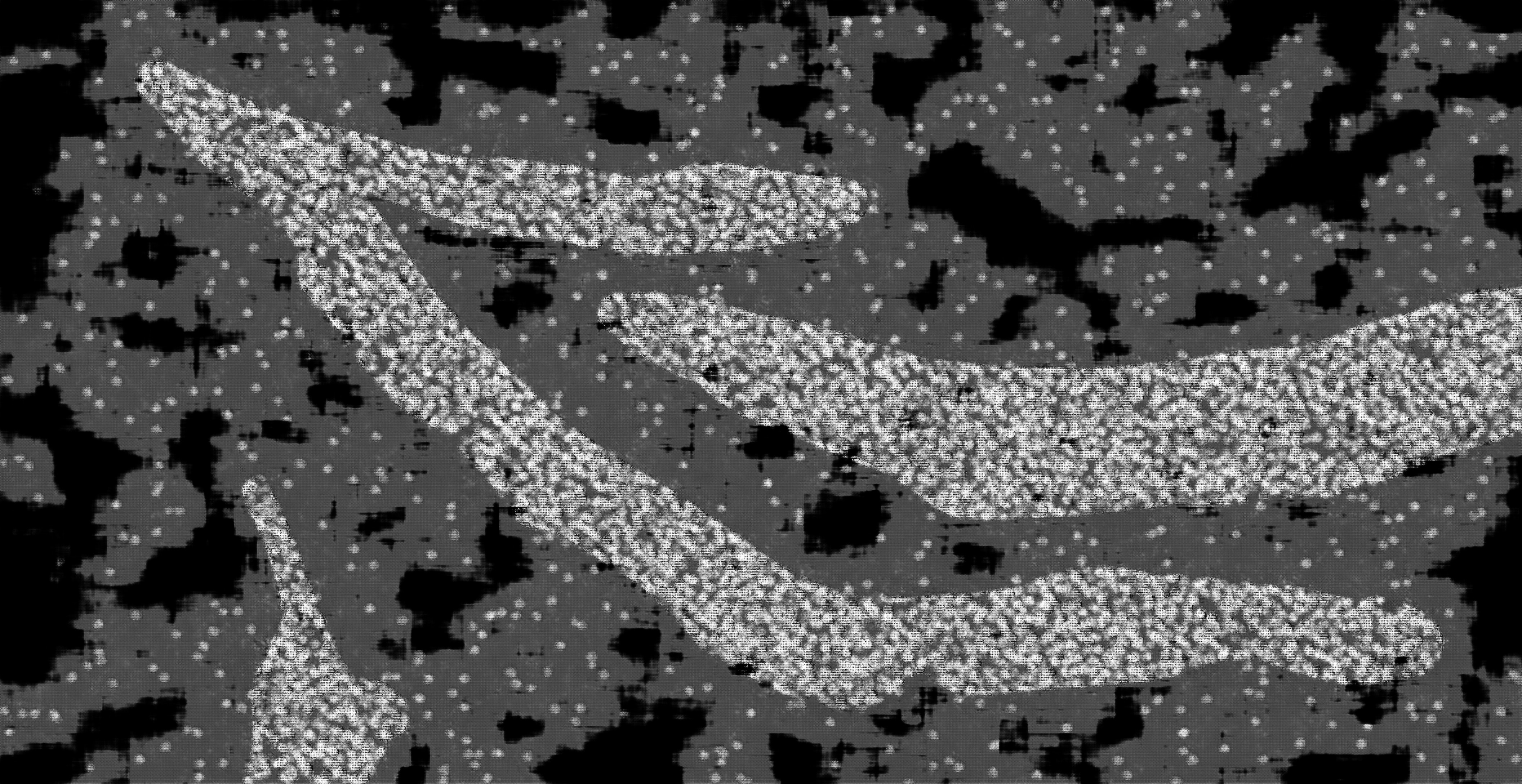

### hc_test_mask2dapi_001_px_FS_64_KS_5_ML_True_4.jpg

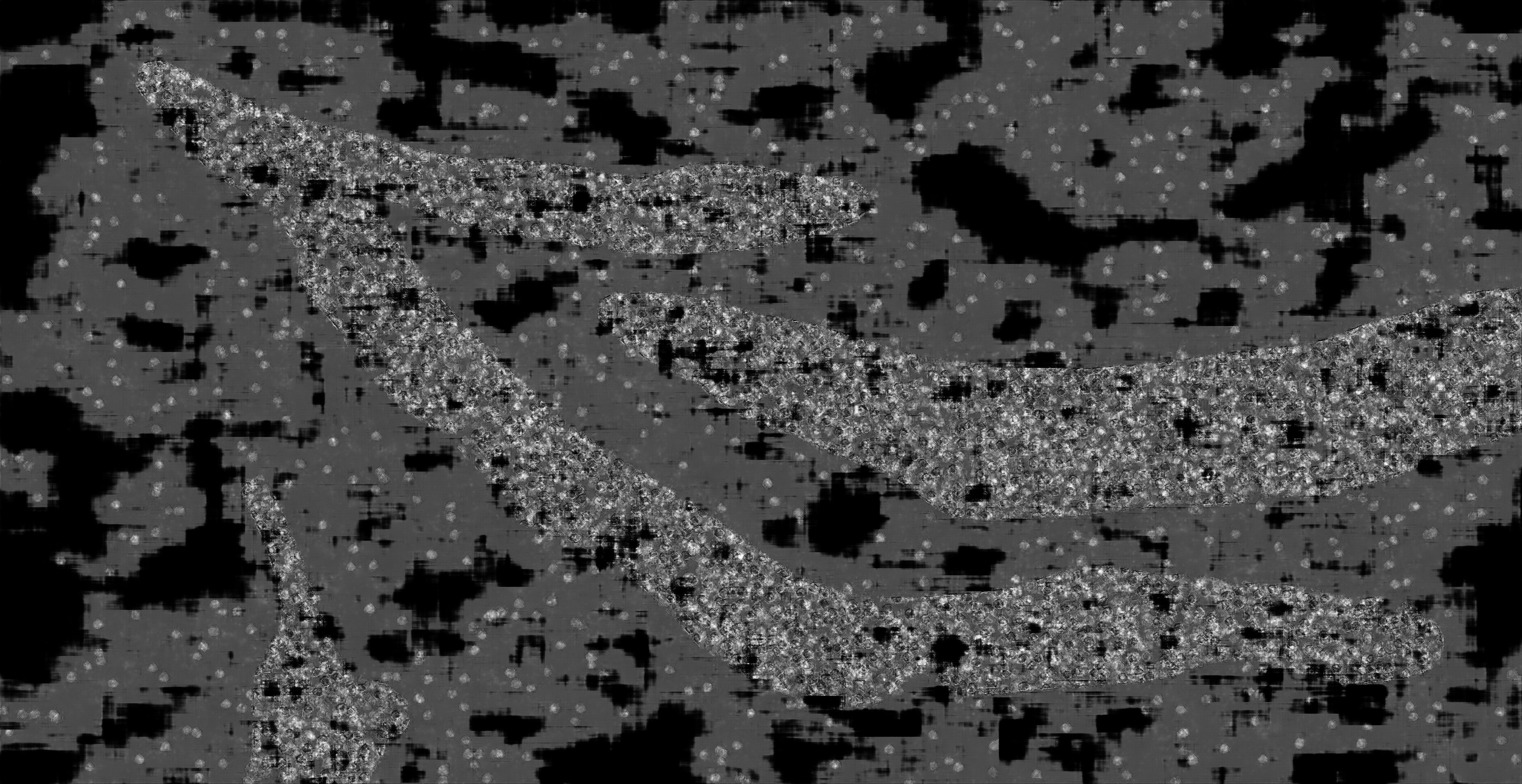

### hc_test_mask2dapi_002a_px_FS_16_KS_3_ML_False_4.jpg

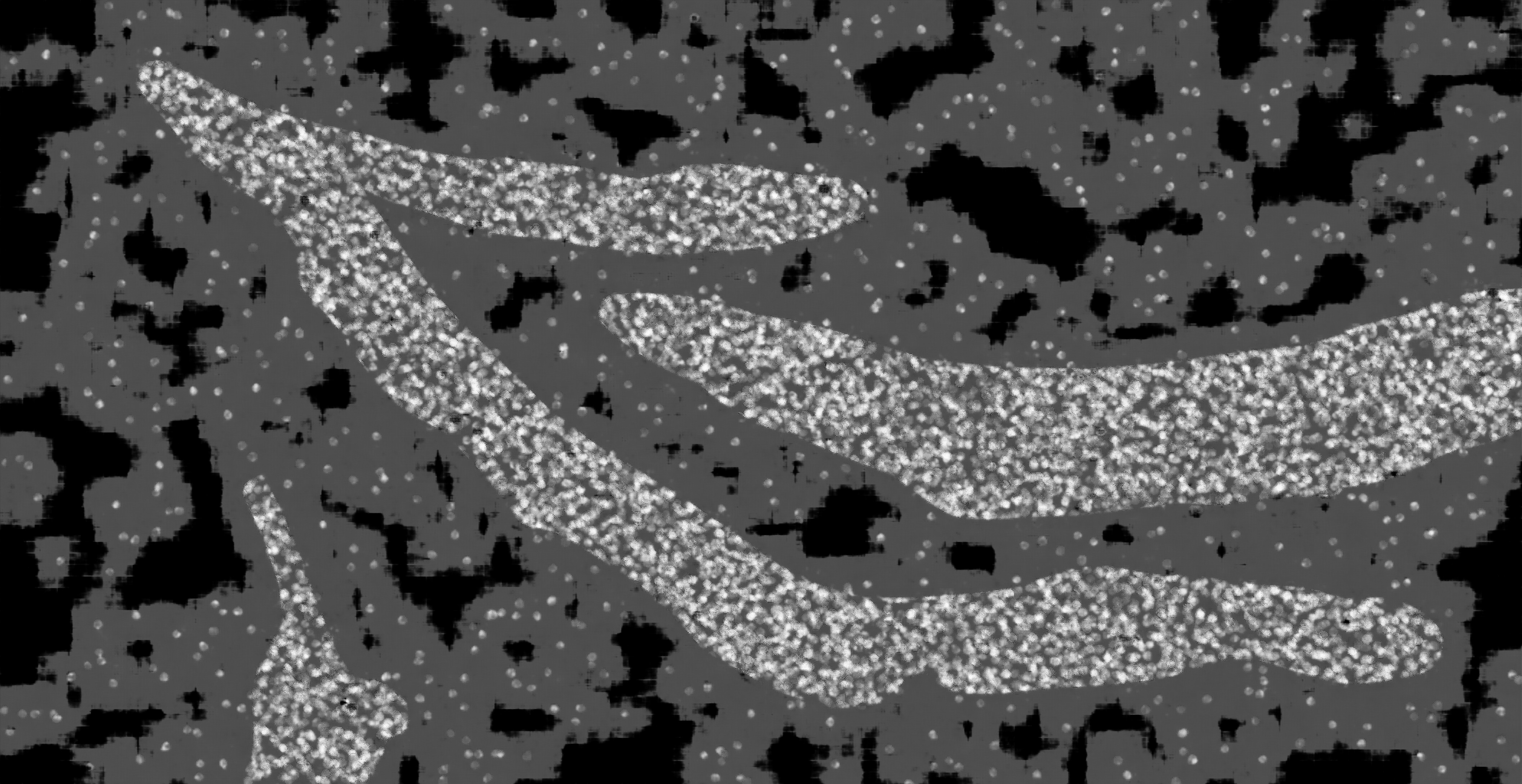

### hc_test_mask2dapi_002a_px_FS_16_KS_3_ML_True_4.jpg

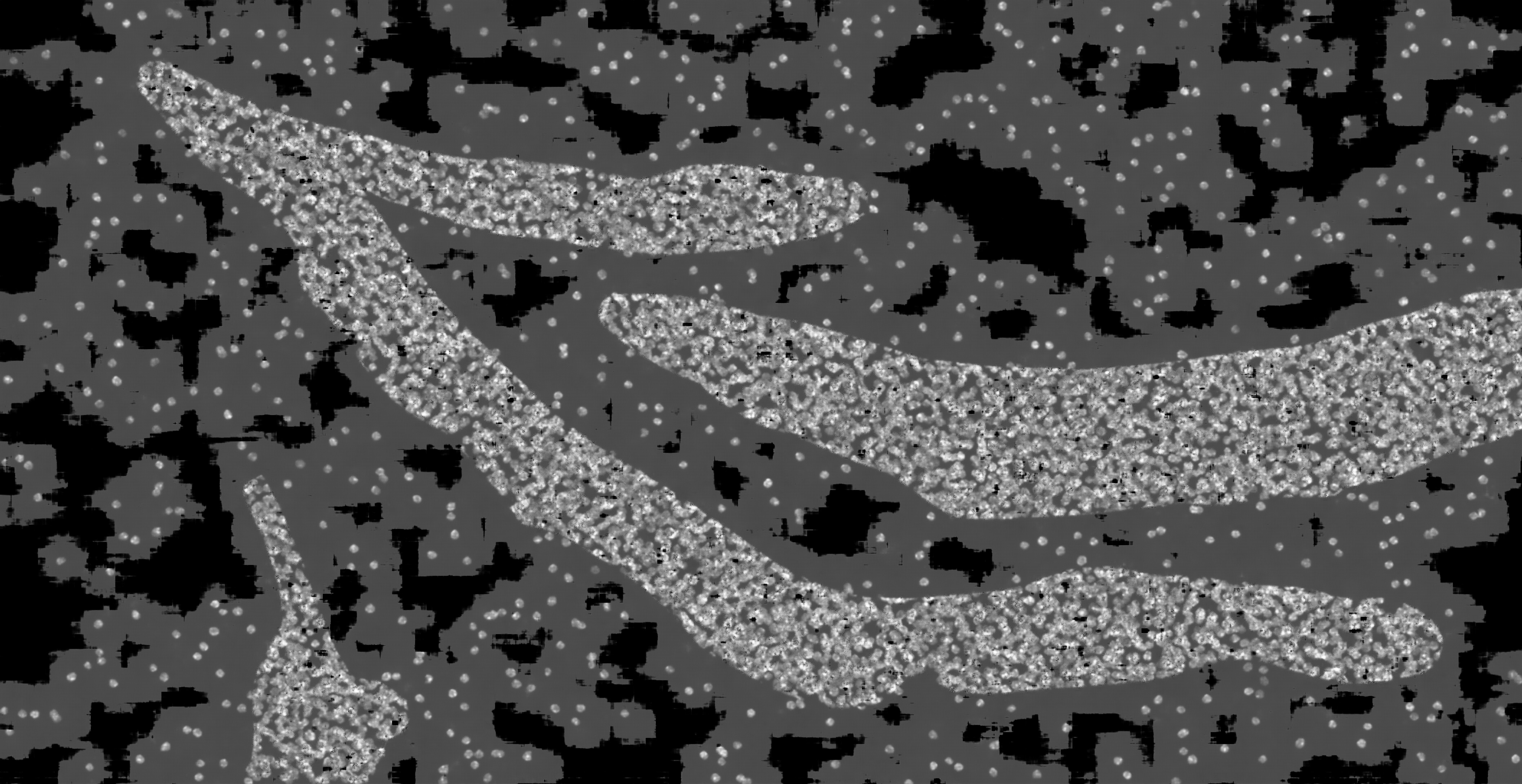

### hc_test_mask2dapi_002a_px_FS_16_KS_5_ML_False_4.jpg

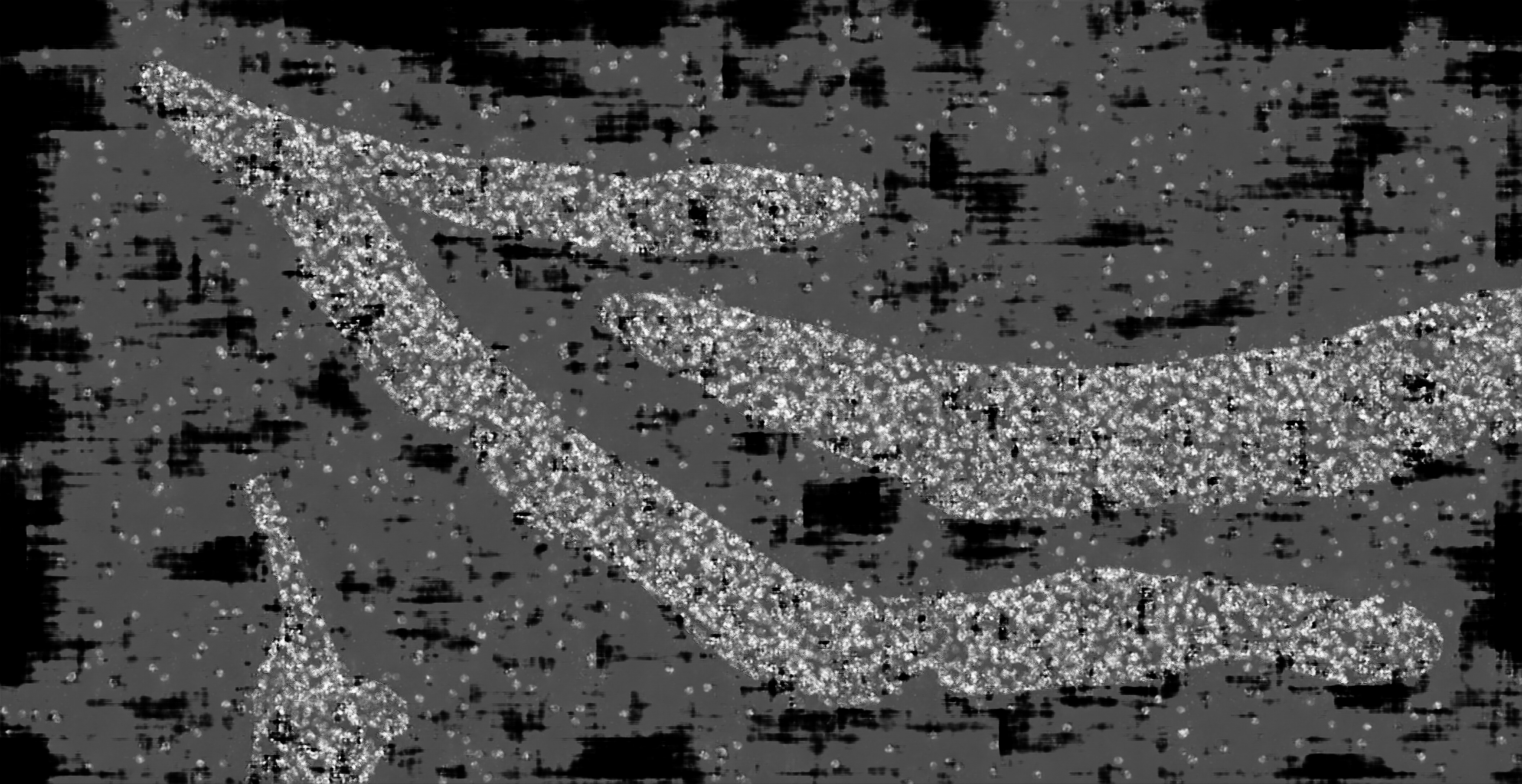

### hc_test_mask2dapi_002a_px_FS_16_KS_5_ML_True_4.jpg

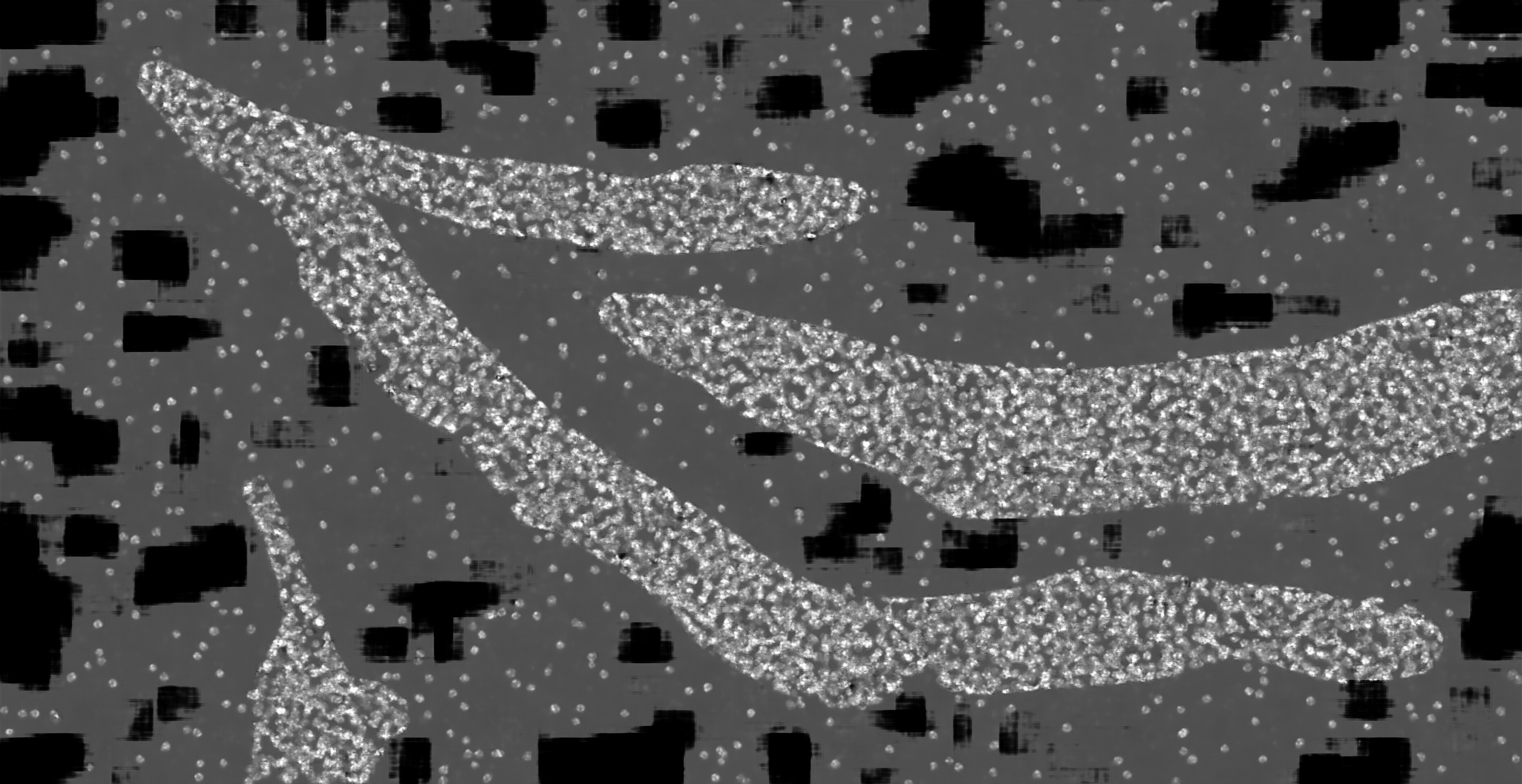

### hc_test_mask2dapi_002a_px_FS_16_KS_7_ML_False_4.jpg

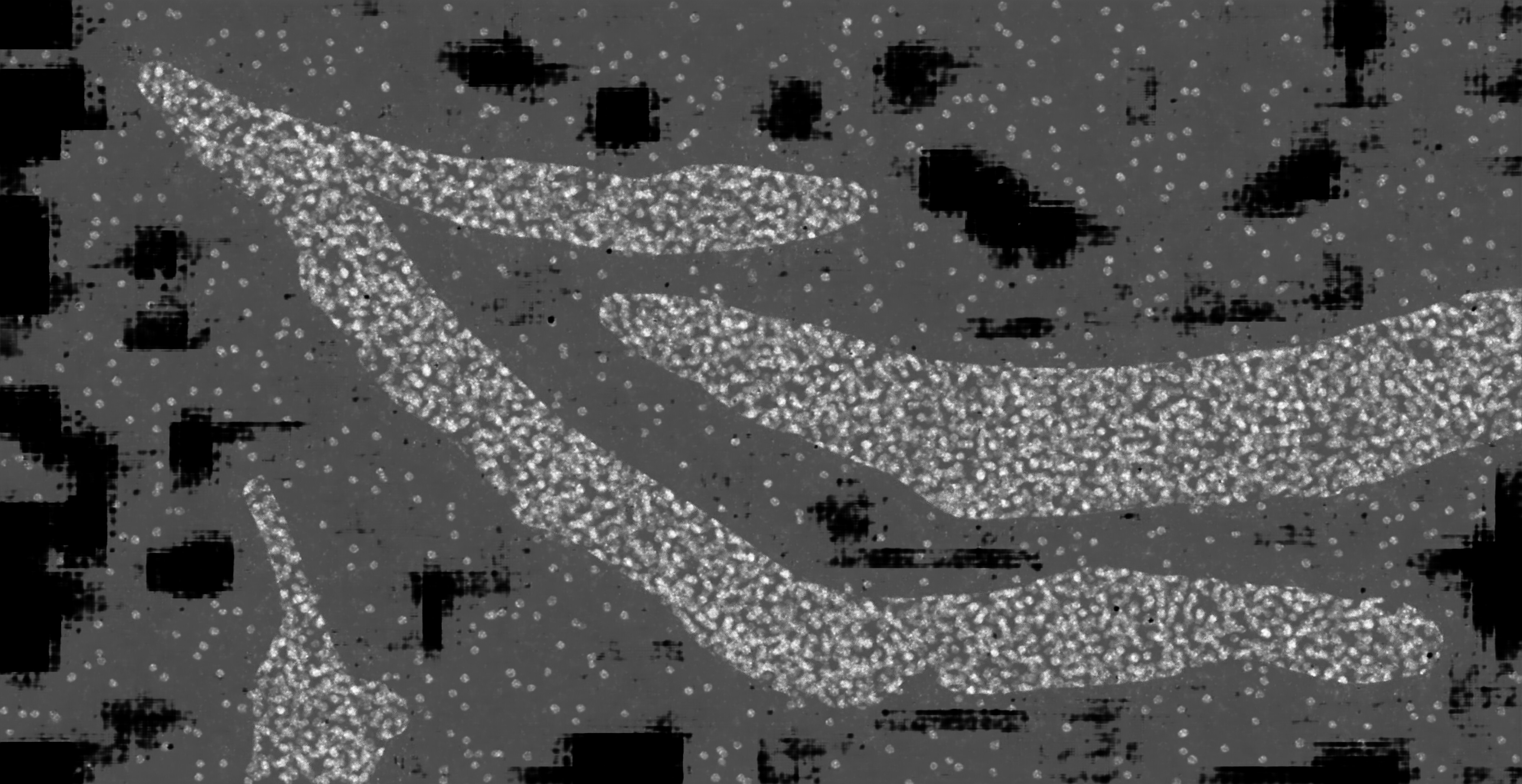

### hc_test_mask2dapi_002a_px_FS_16_KS_7_ML_True_4.jpg

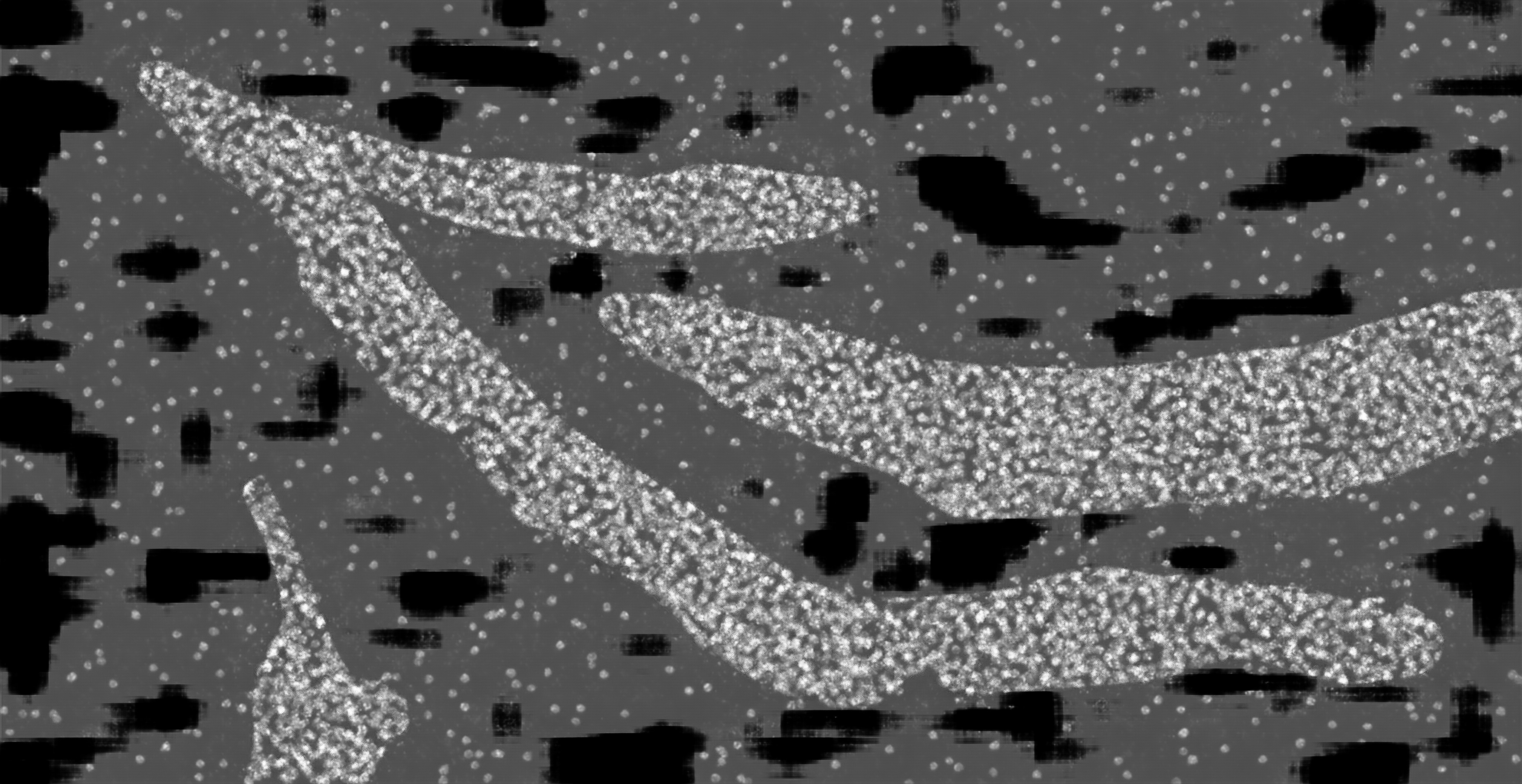

### hc_test_mask2dapi_002b_px_FS_16_KS_3_ML_False_4.jpg

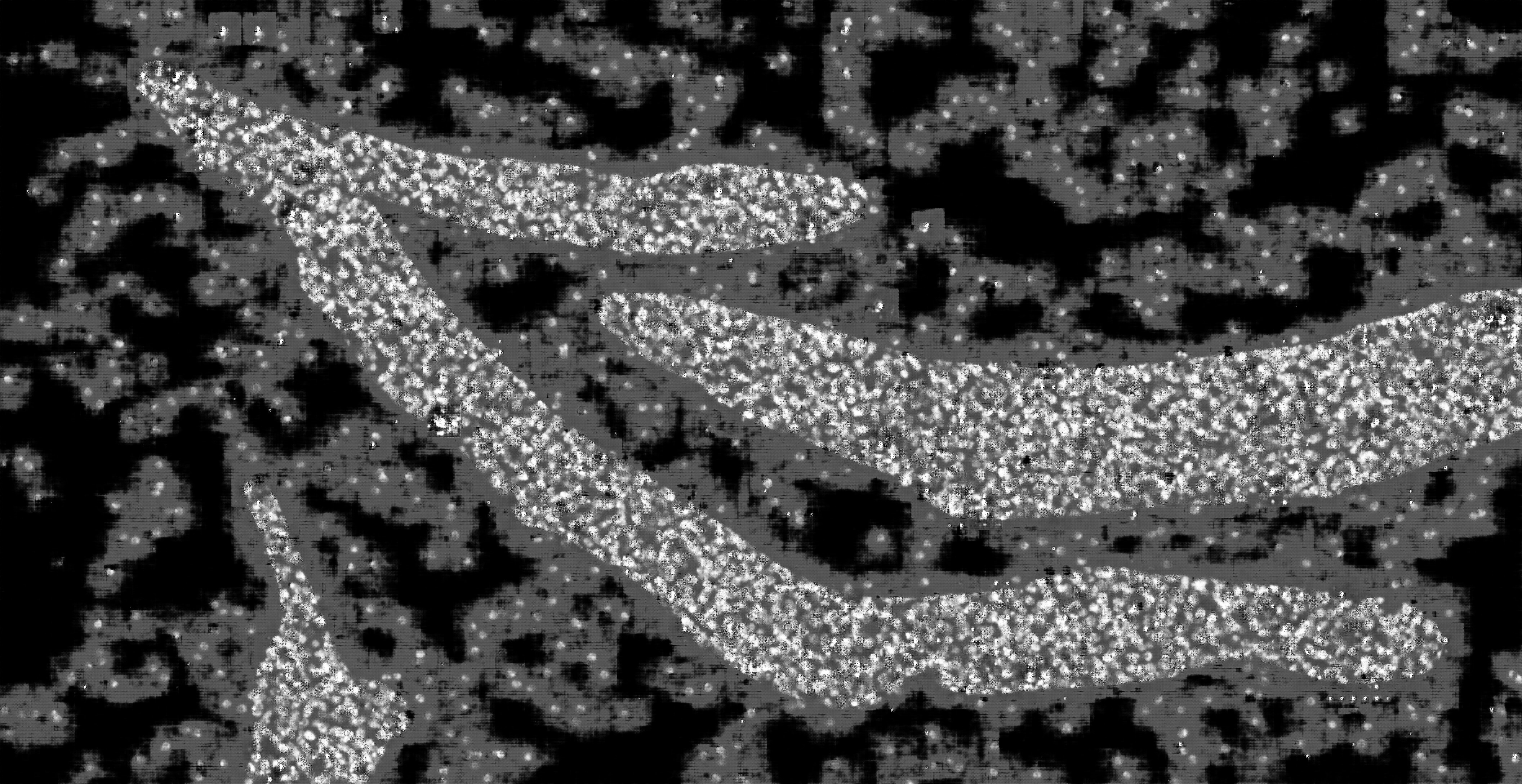

### hc_test_mask2dapi_002b_px_FS_16_KS_3_ML_True_4.jpg

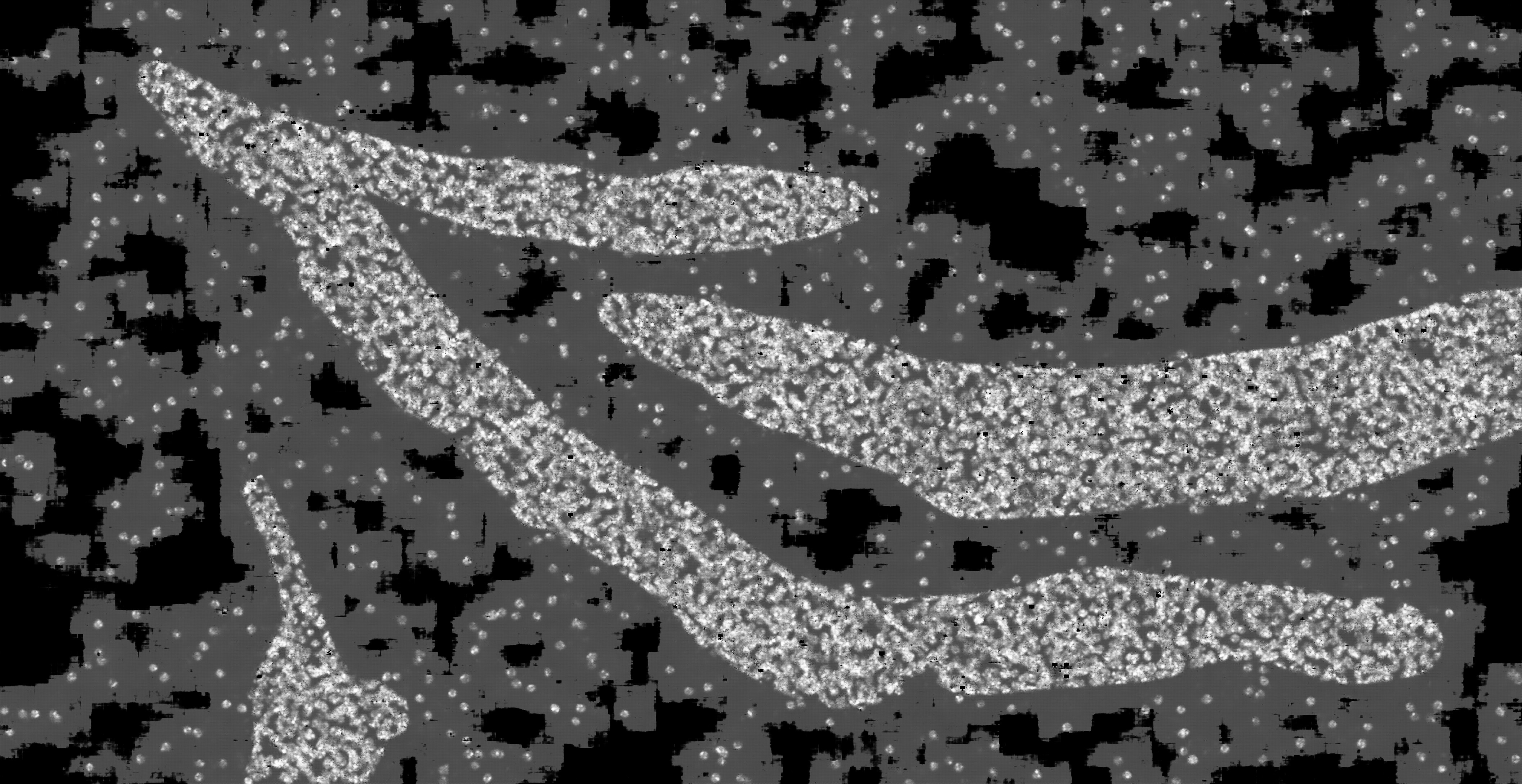

### hc_test_mask2dapi_002b_px_FS_16_KS_5_ML_False_4.jpg

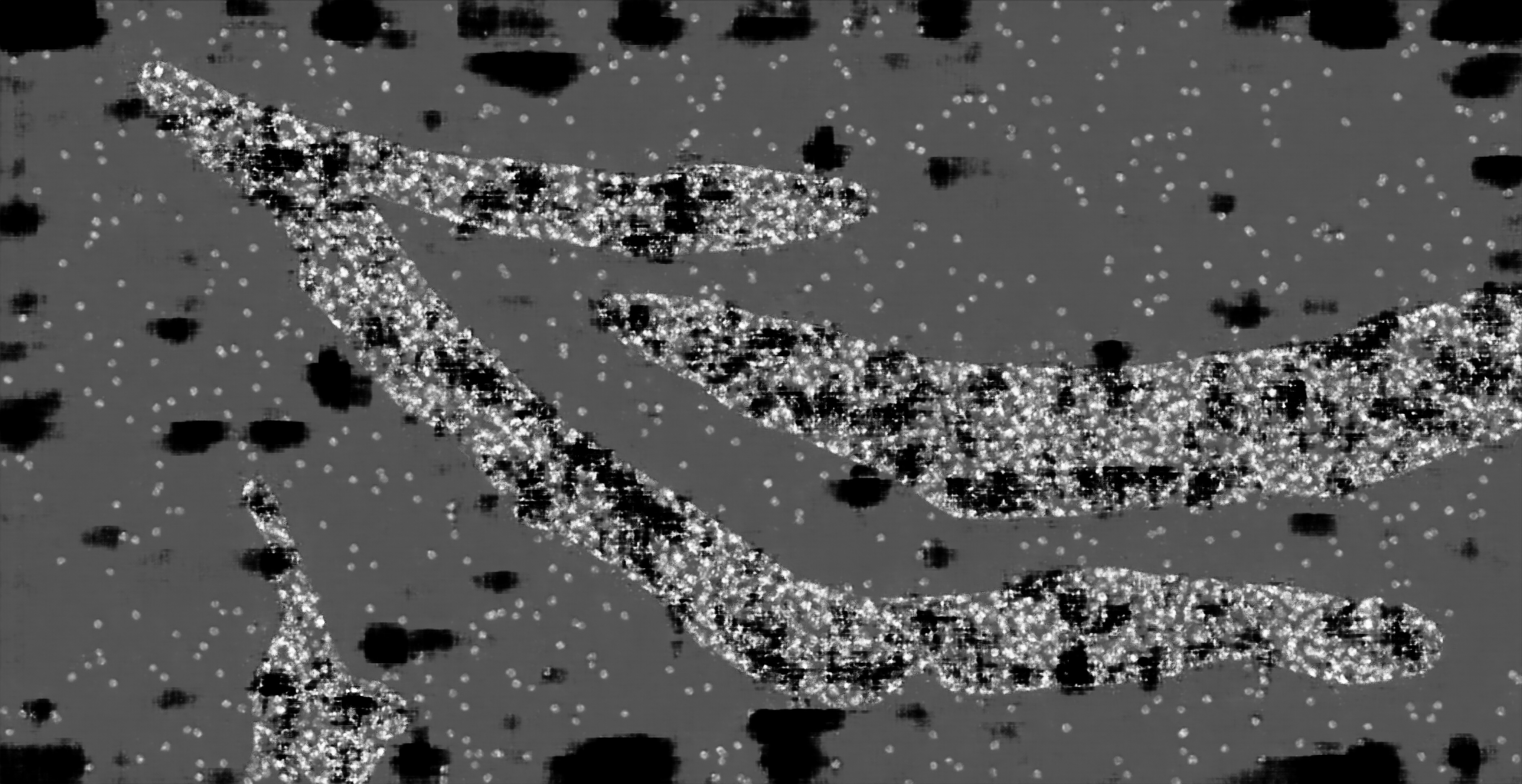

### hc_test_mask2dapi_002b_px_FS_16_KS_5_ML_True_4.jpg

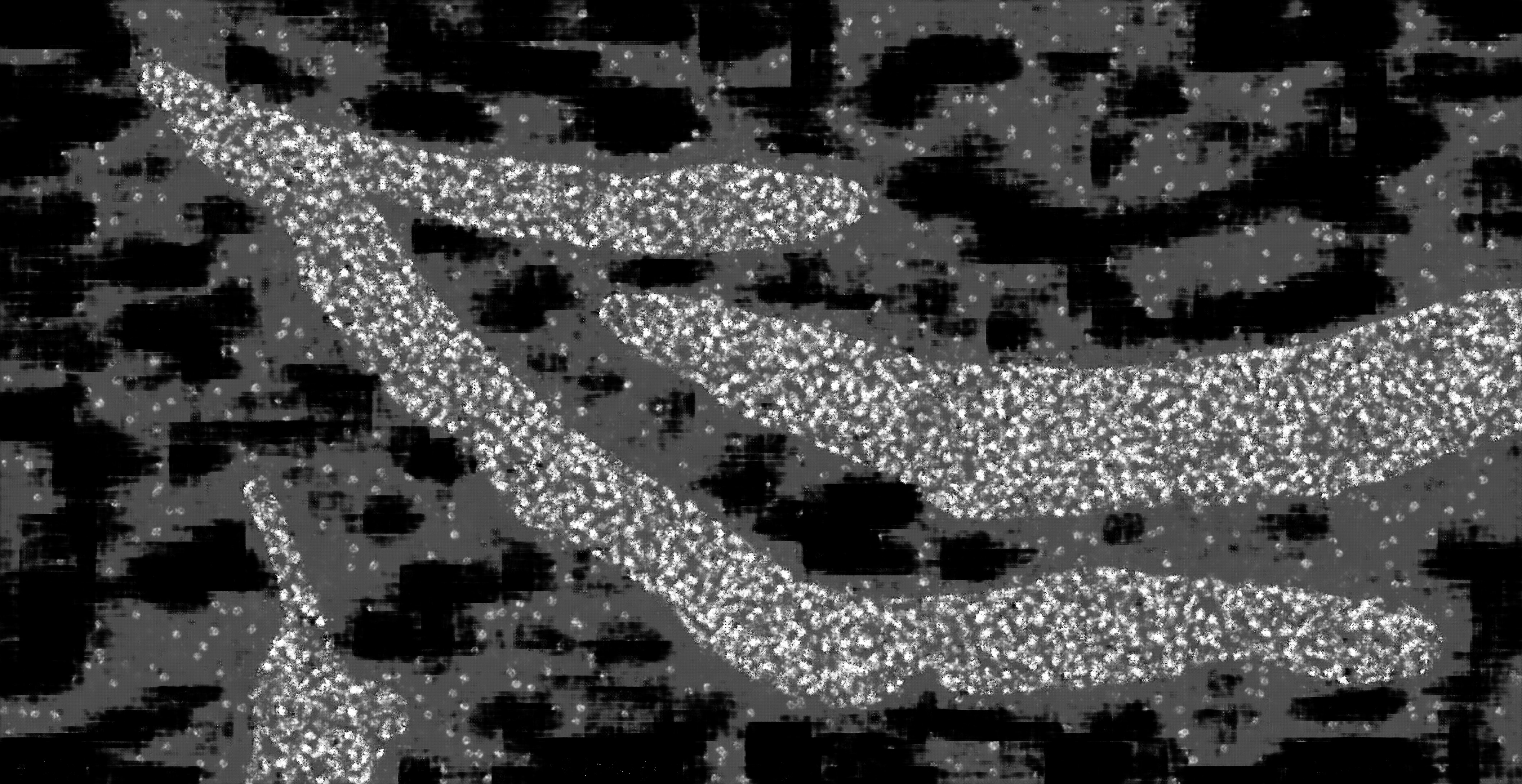

### hc_test_mask2dapi_002b_px_FS_16_KS_7_ML_False_4.jpg

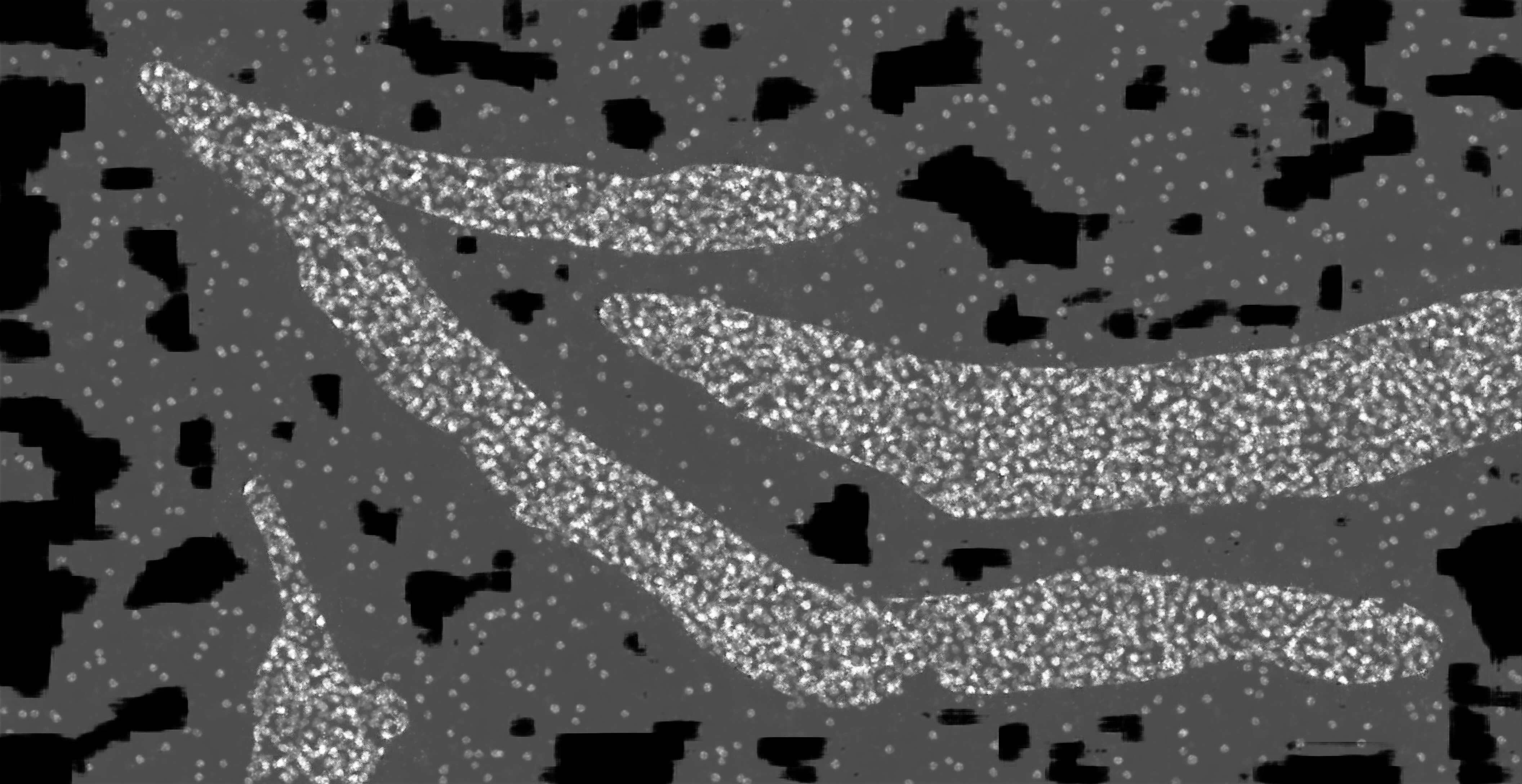

### hc_test_mask2dapi_002b_px_FS_16_KS_7_ML_True_4.jpg

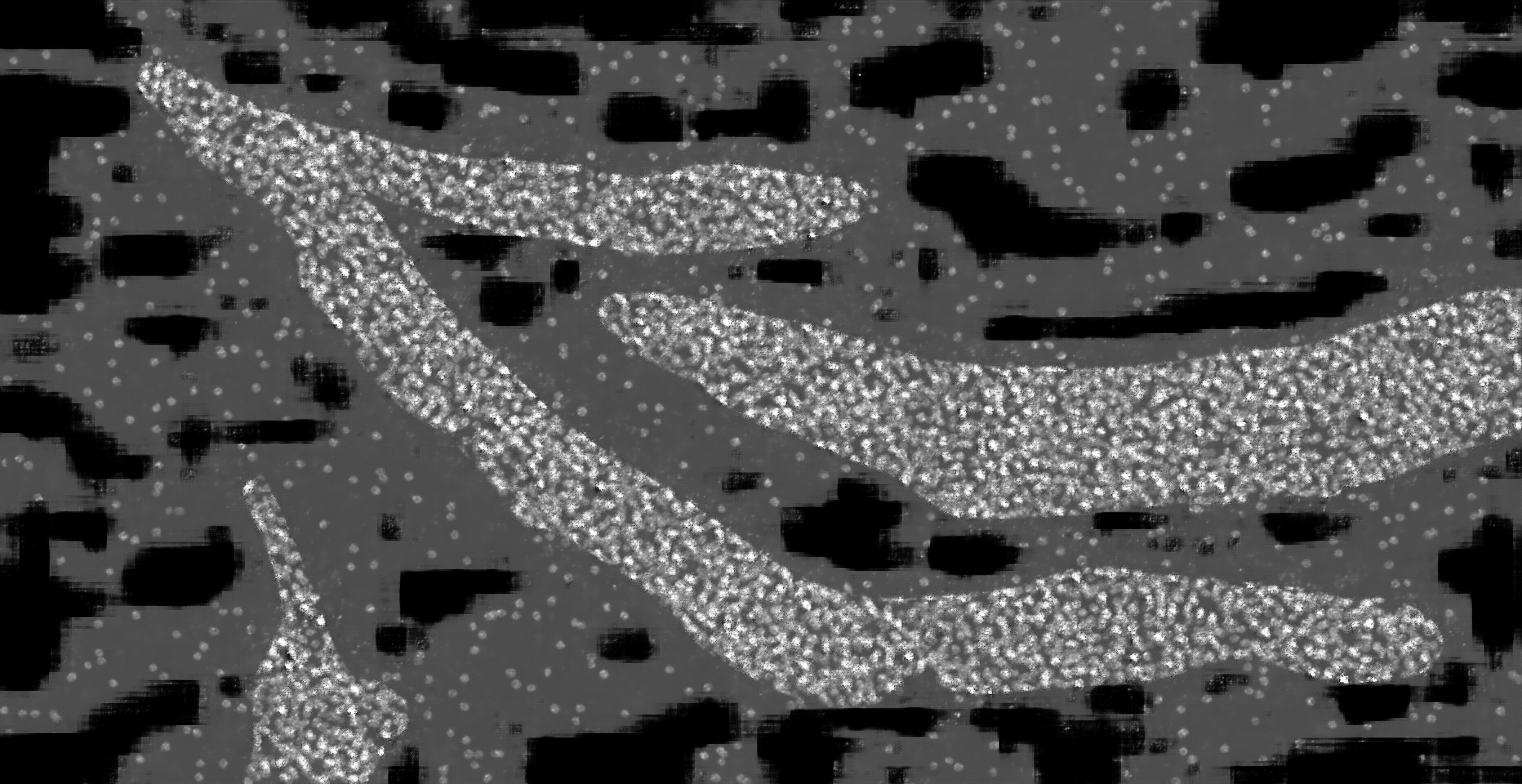

### hc_test_mask2dapi_003a_px_FS_16_KS_3_ML_False_4.jpg

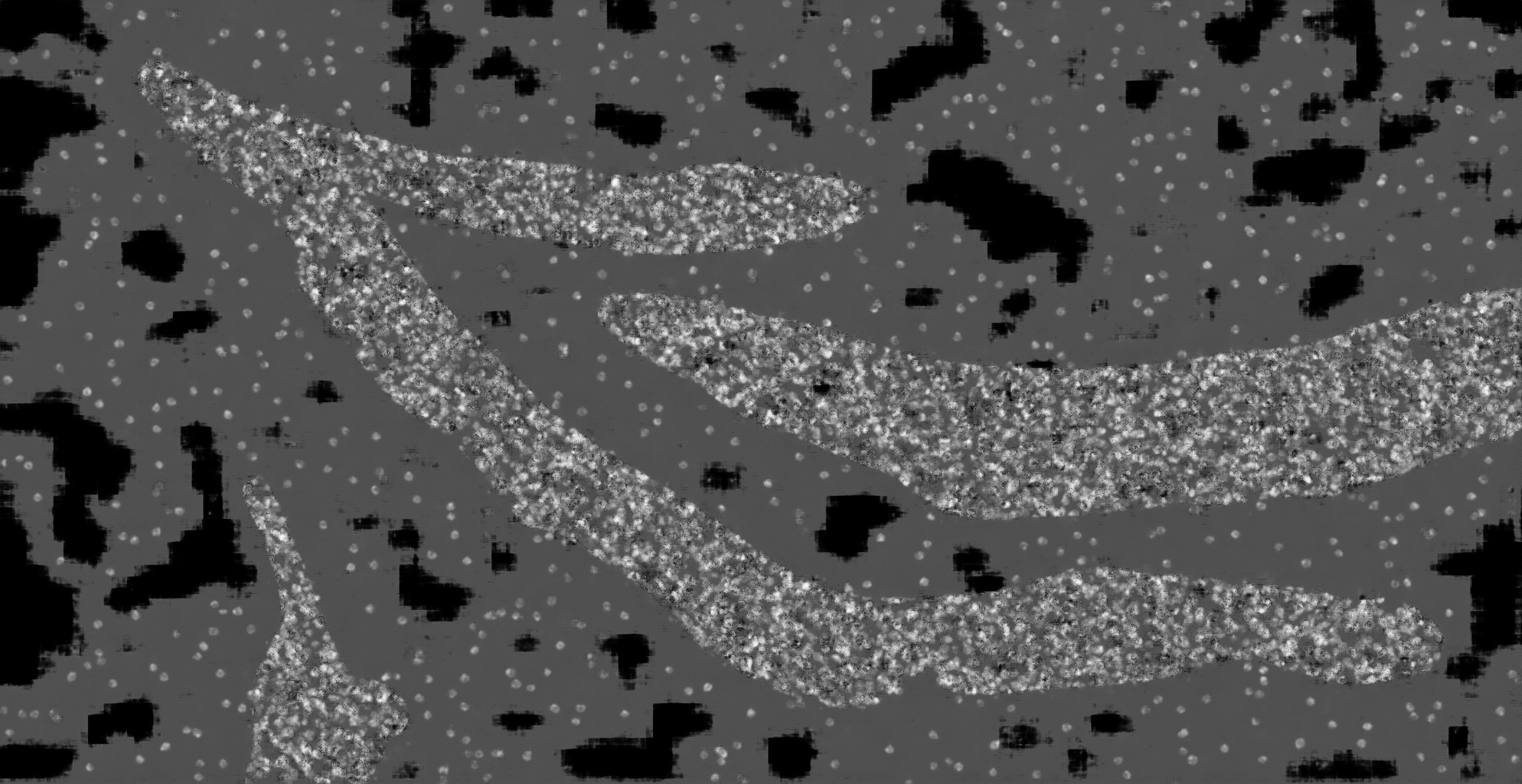

### hc_test_mask2dapi_003a_px_FS_16_KS_3_ML_True_4.jpg

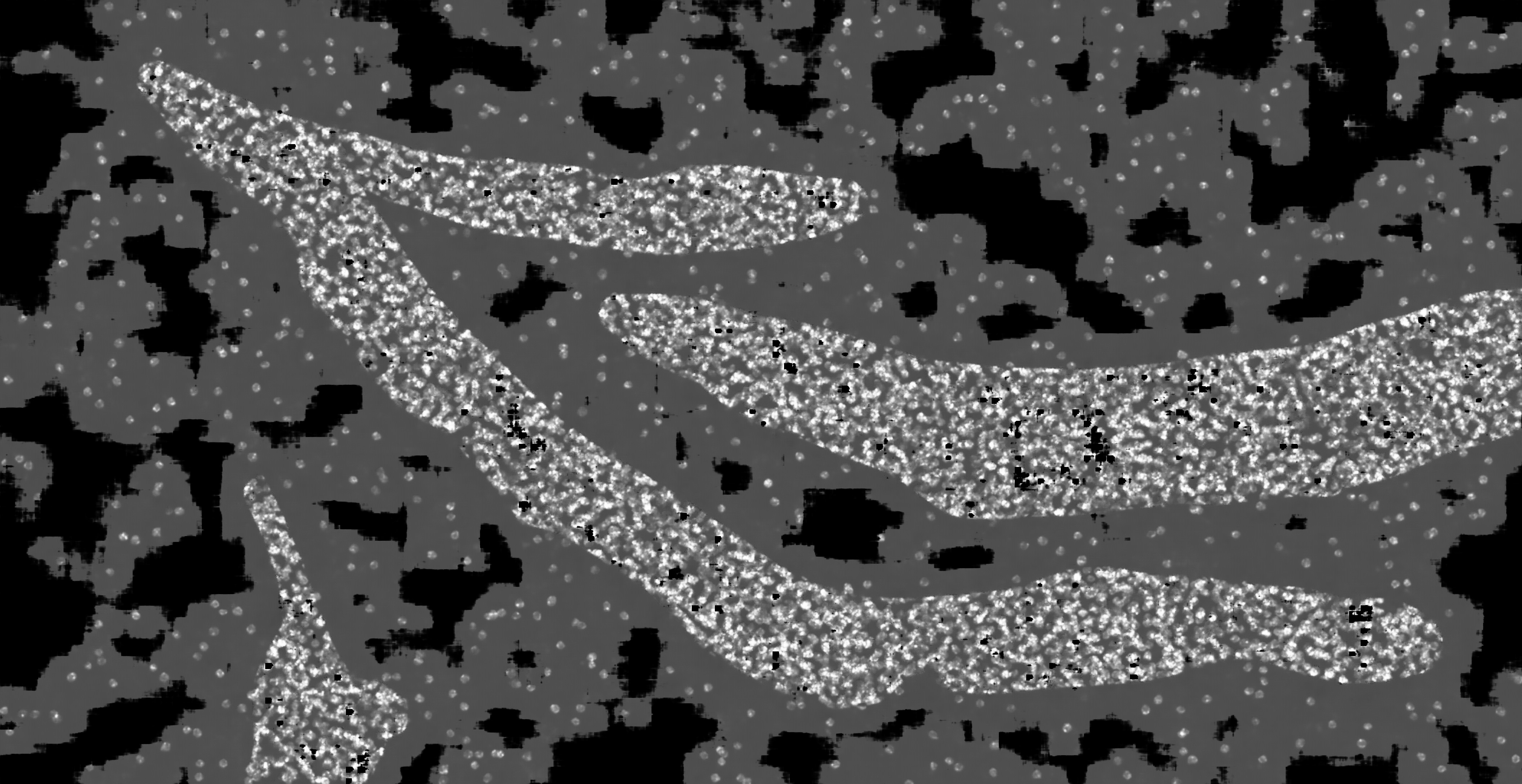

### hc_test_mask2dapi_003a_px_FS_16_KS_5_ML_False_4.jpg

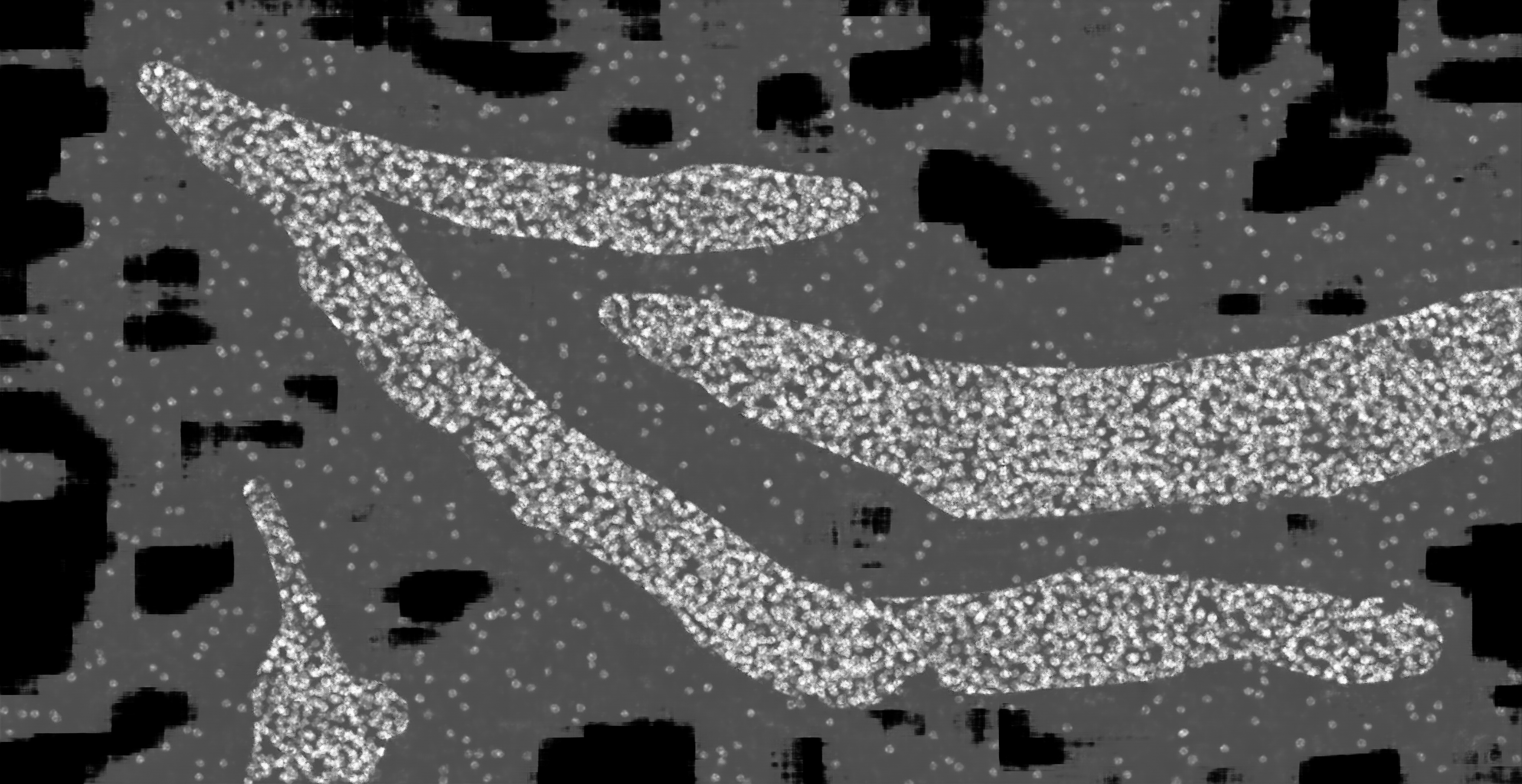

### hc_test_mask2dapi_003a_px_FS_16_KS_5_ML_True_4.jpg

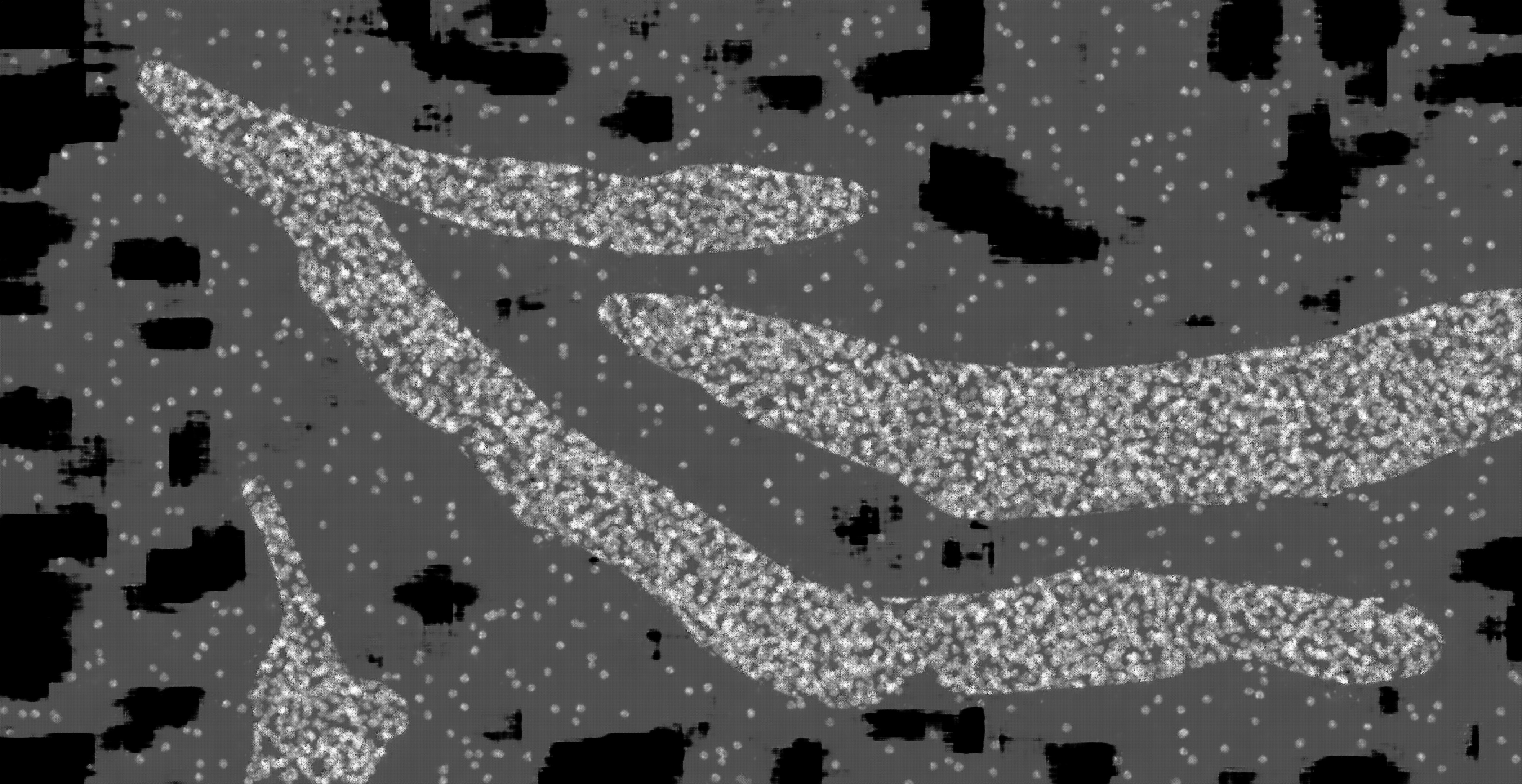

### hc_test_mask2dapi_003a_px_FS_16_KS_7_ML_False_4.jpg

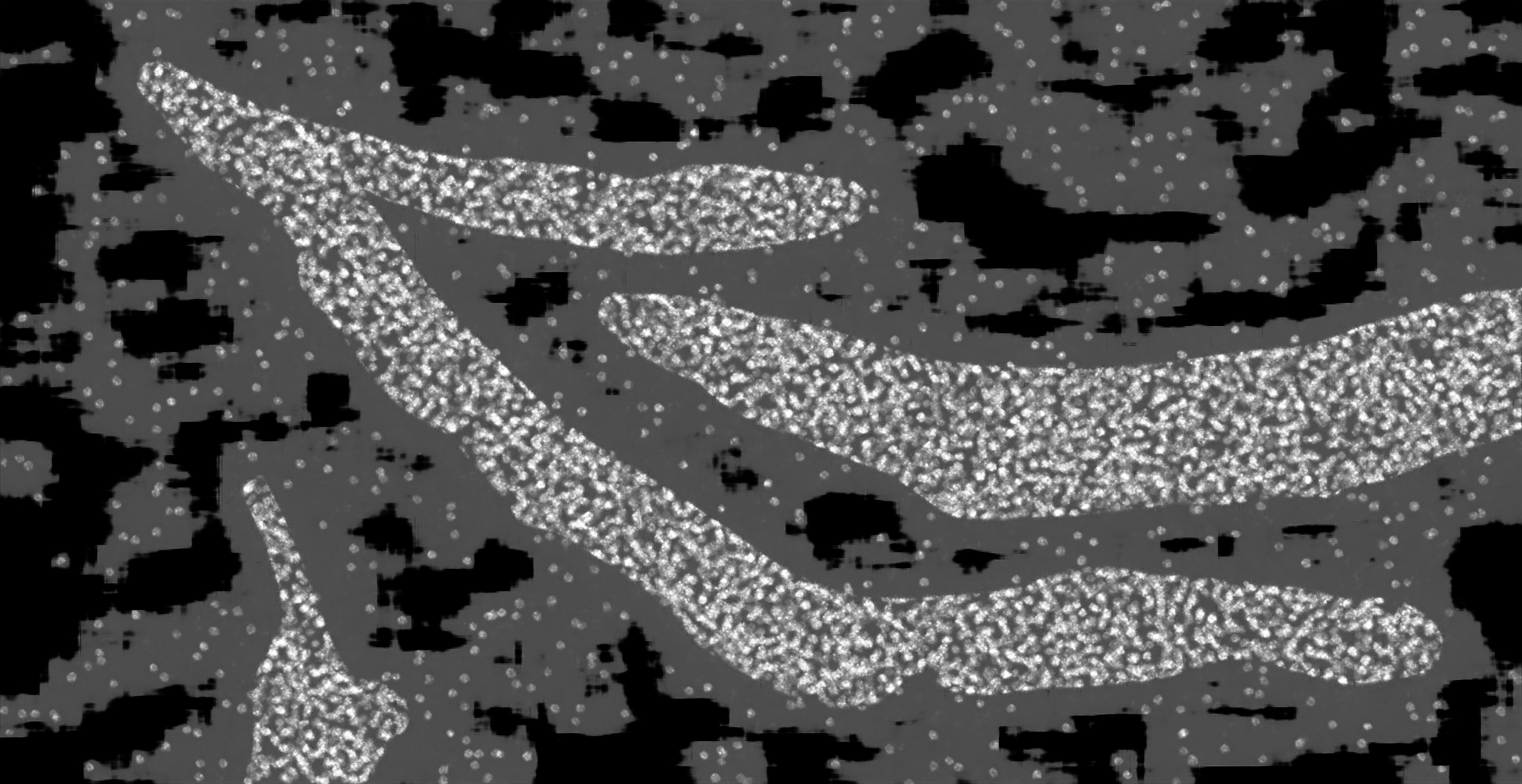

### hc_test_mask2dapi_003a_px_FS_16_KS_7_ML_True_4.jpg

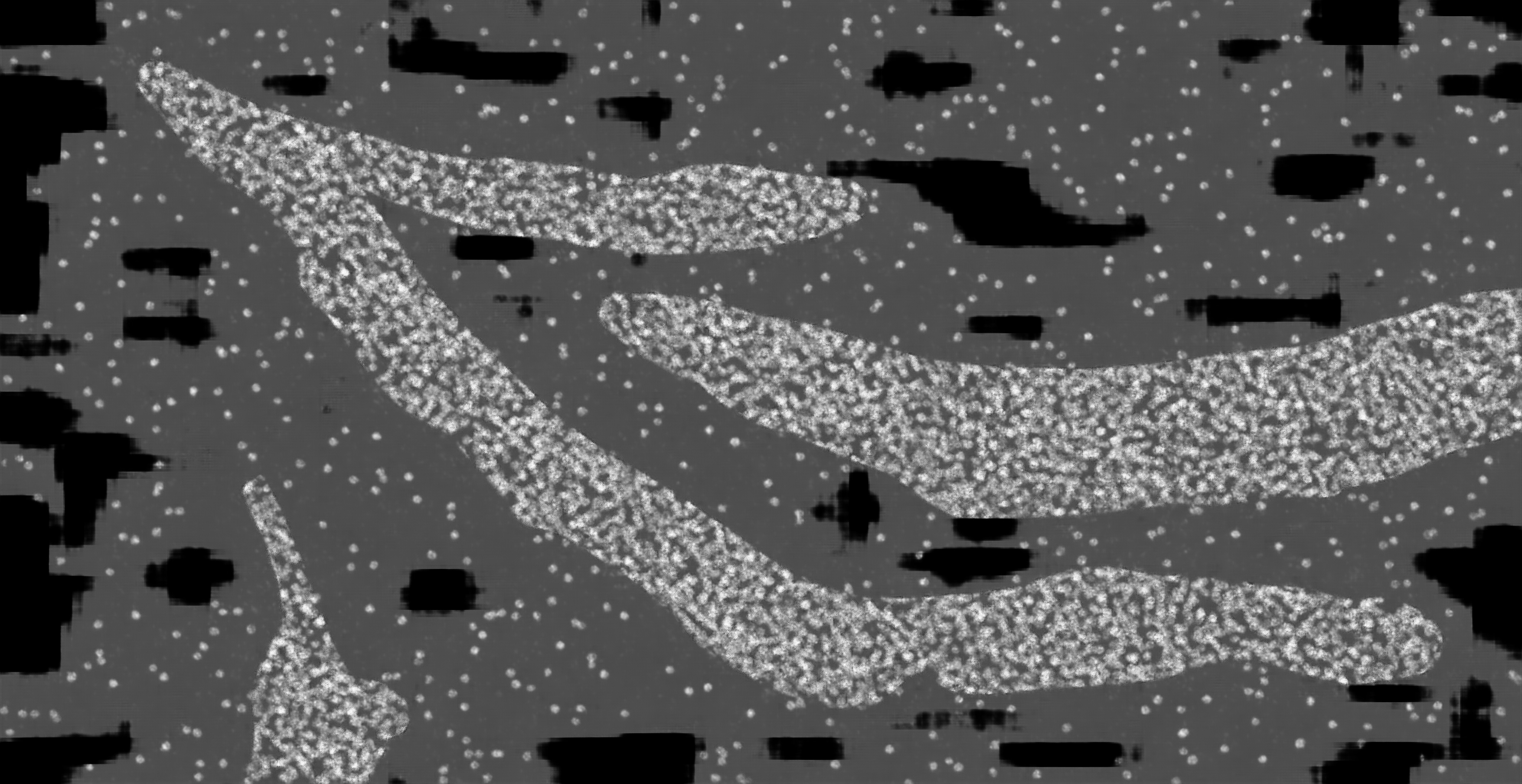

### hc_test_mask2dapi_003b_px_FS_16_KS_3_ML_False_4.jpg

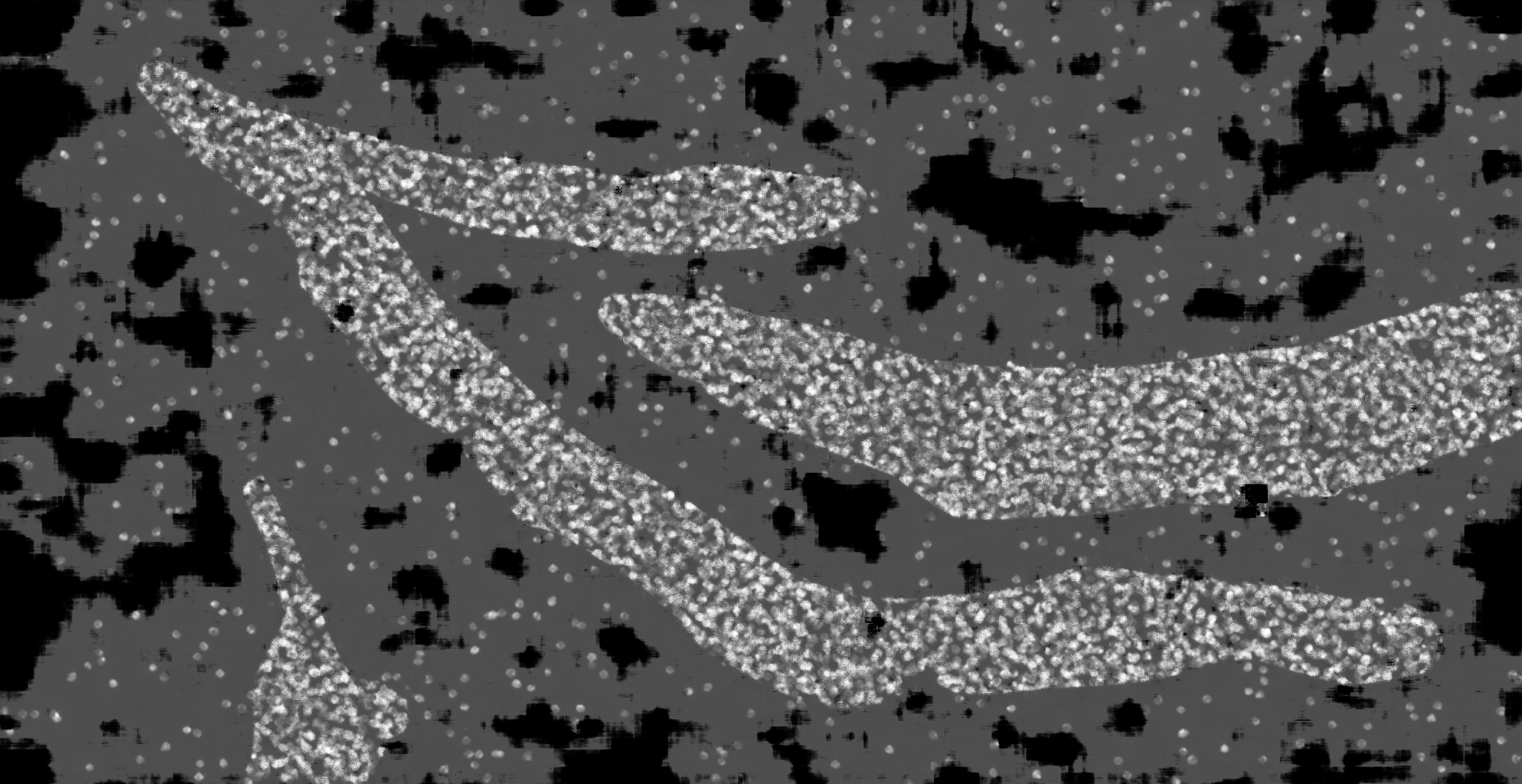

### hc_test_mask2dapi_003b_px_FS_16_KS_3_ML_True_4.jpg

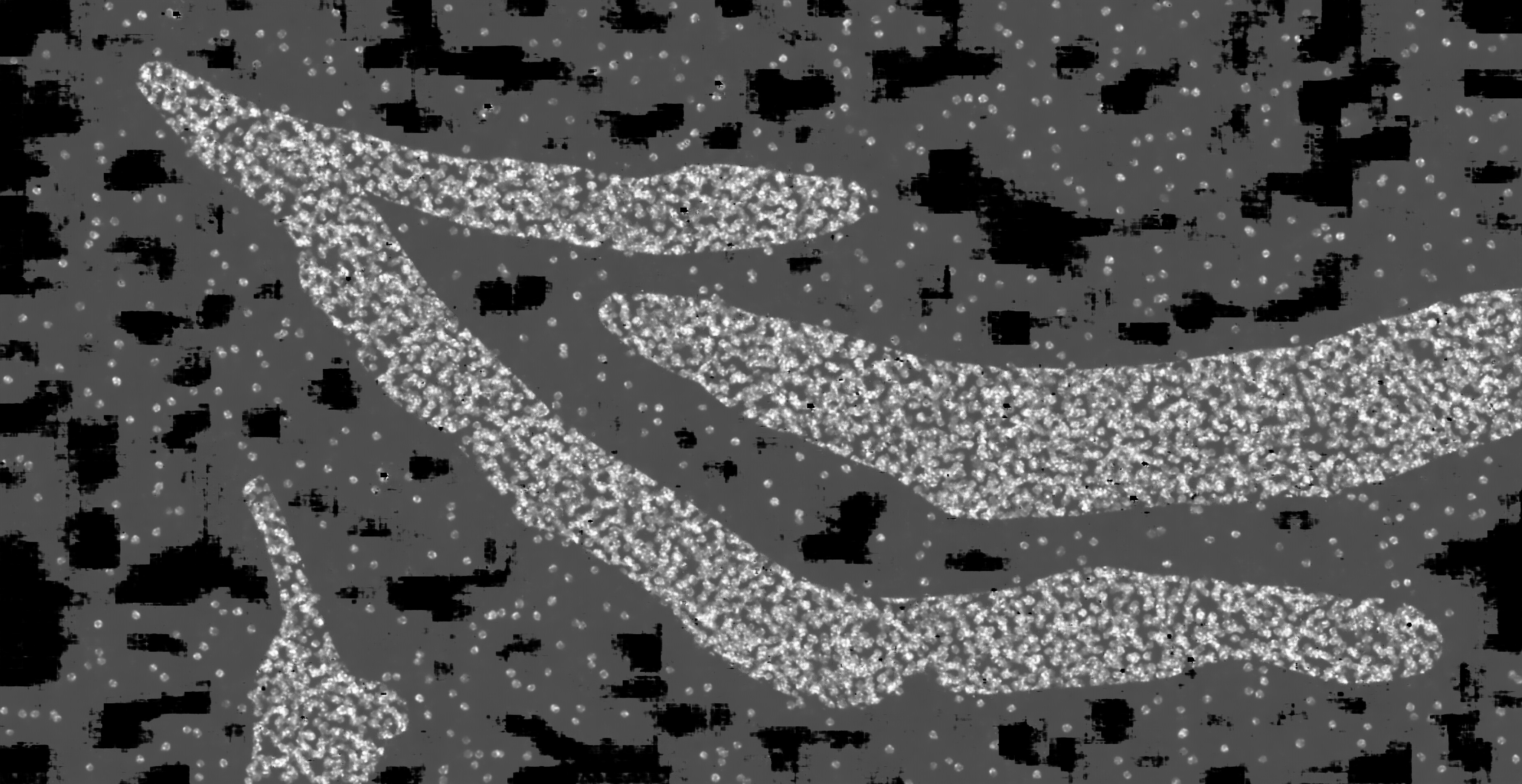

### hc_test_mask2dapi_003b_px_FS_16_KS_5_ML_False_4.jpg

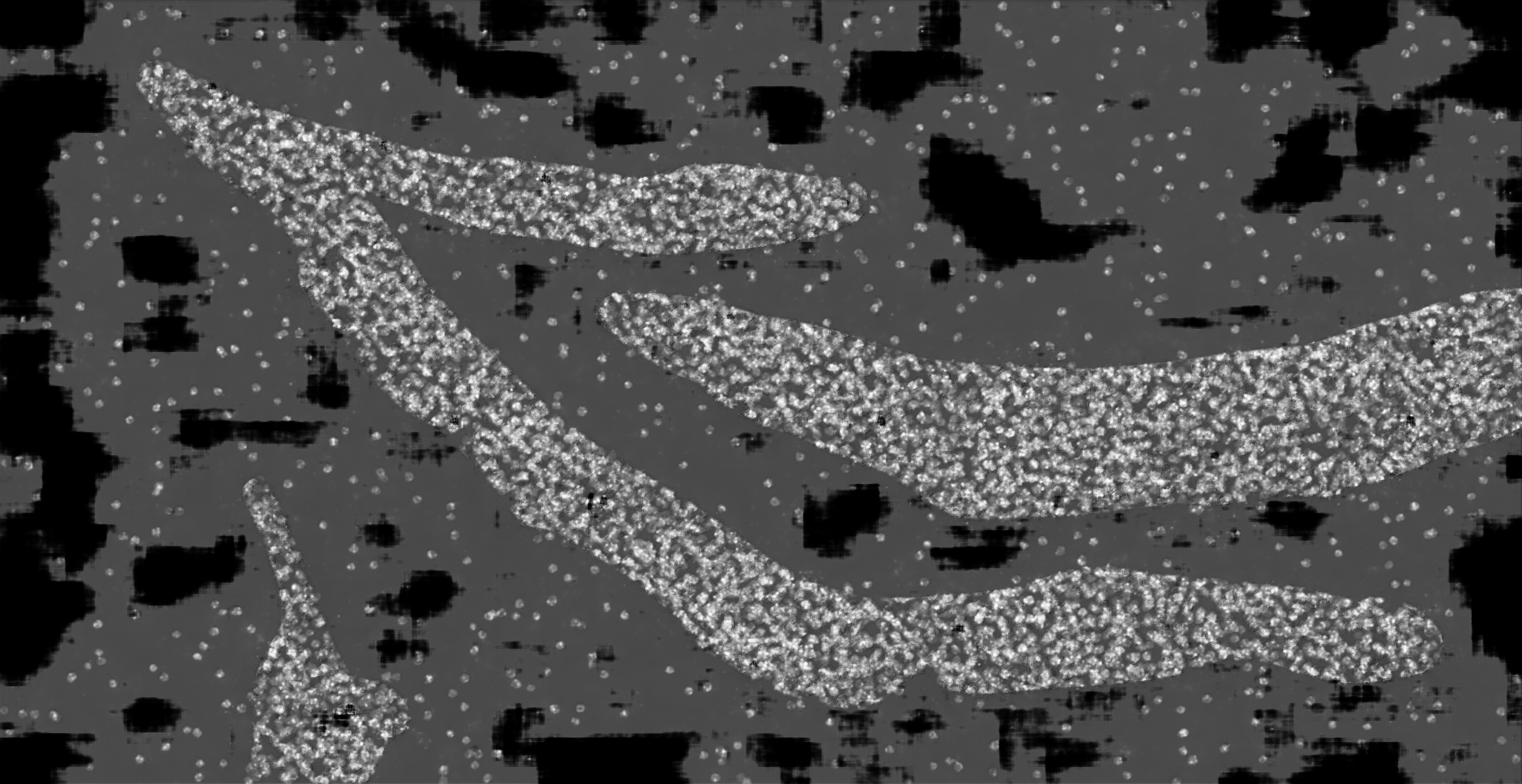

### hc_test_mask2dapi_003b_px_FS_16_KS_5_ML_True_4.jpg

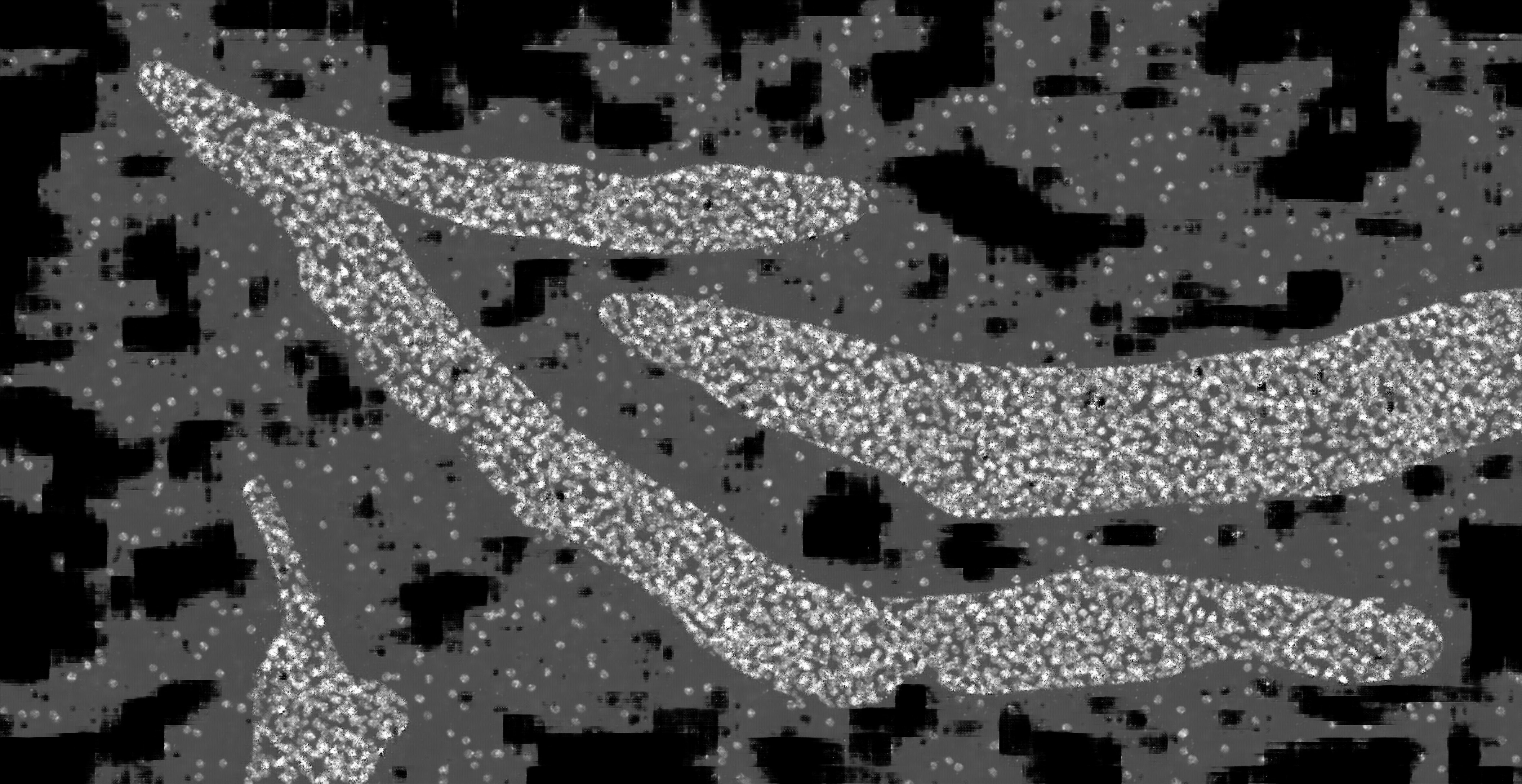

### hc_test_mask2dapi_003b_px_FS_16_KS_7_ML_False_4.jpg

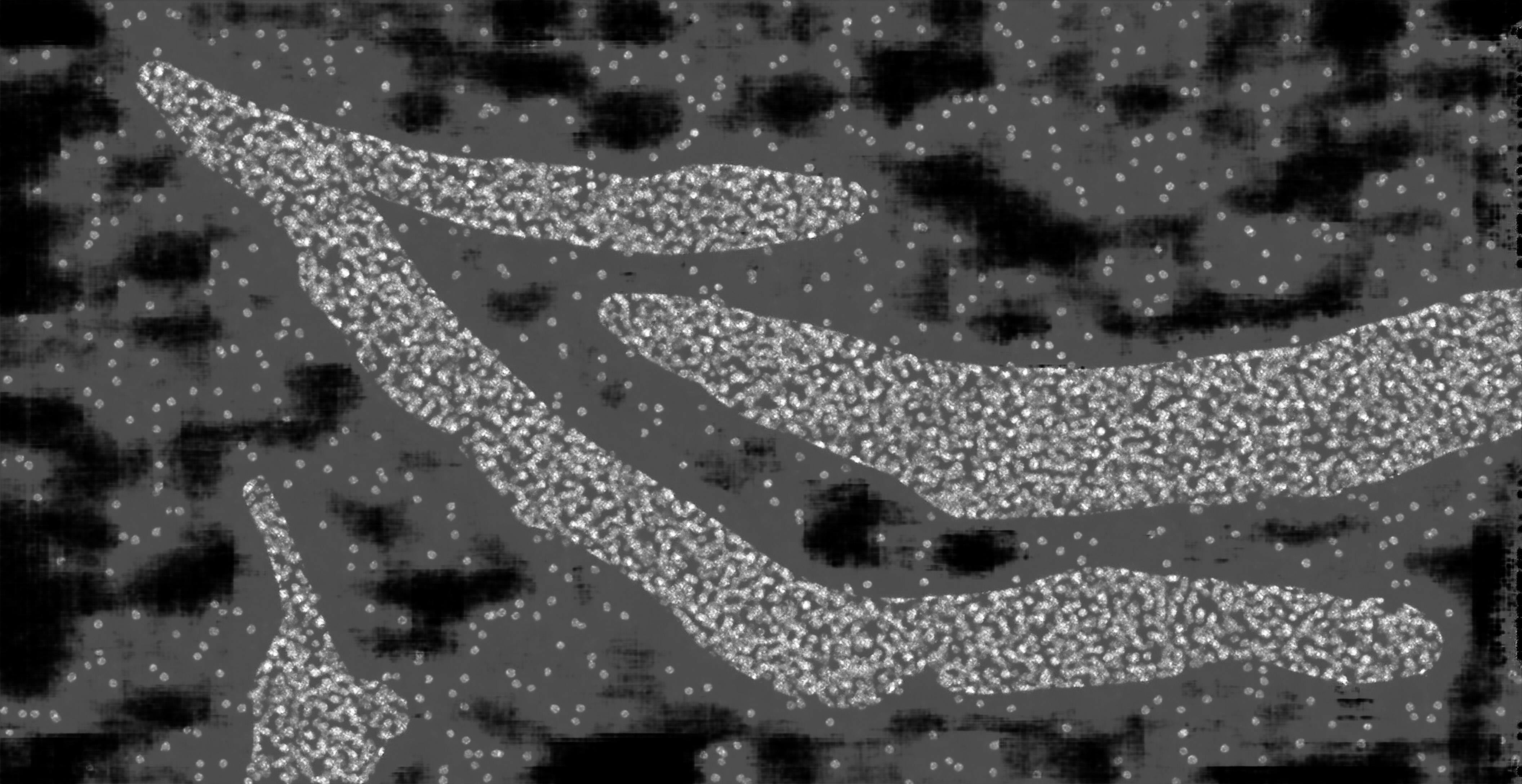

### hc_test_mask2dapi_003b_px_FS_16_KS_7_ML_True_4.jpg

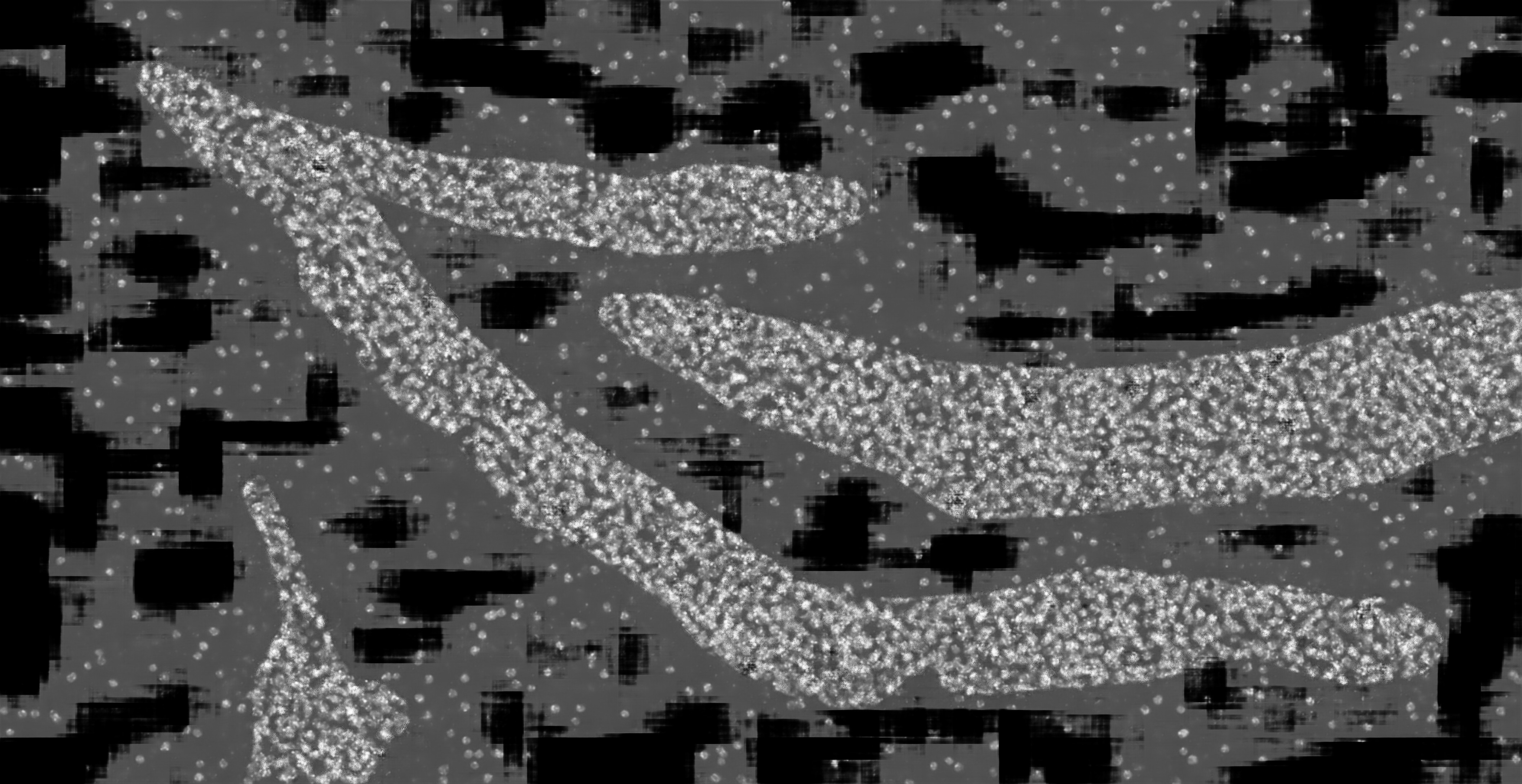

### Supp_Fig_1_zebrafish.jpg

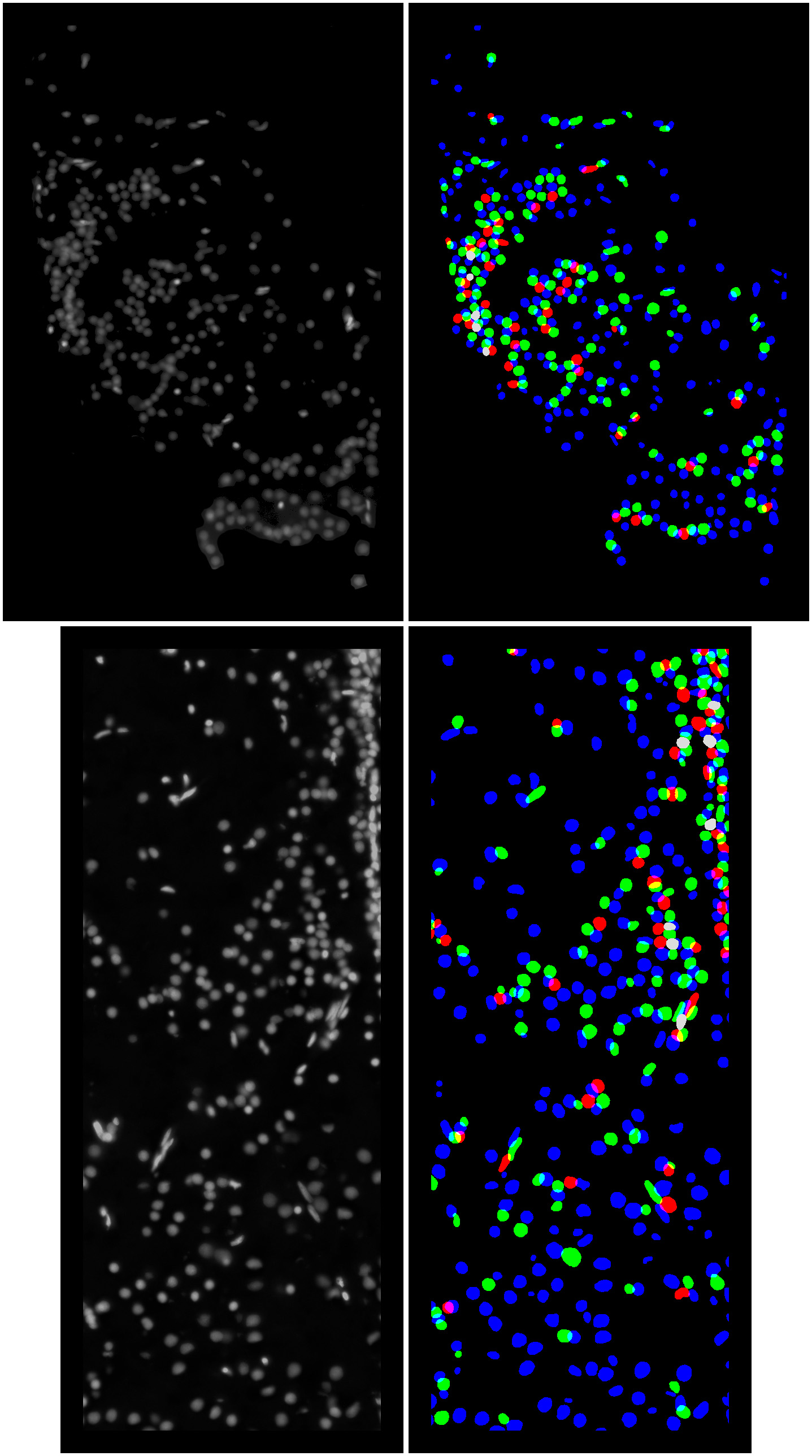

### Supp_Fig_2a_mouse.jpg

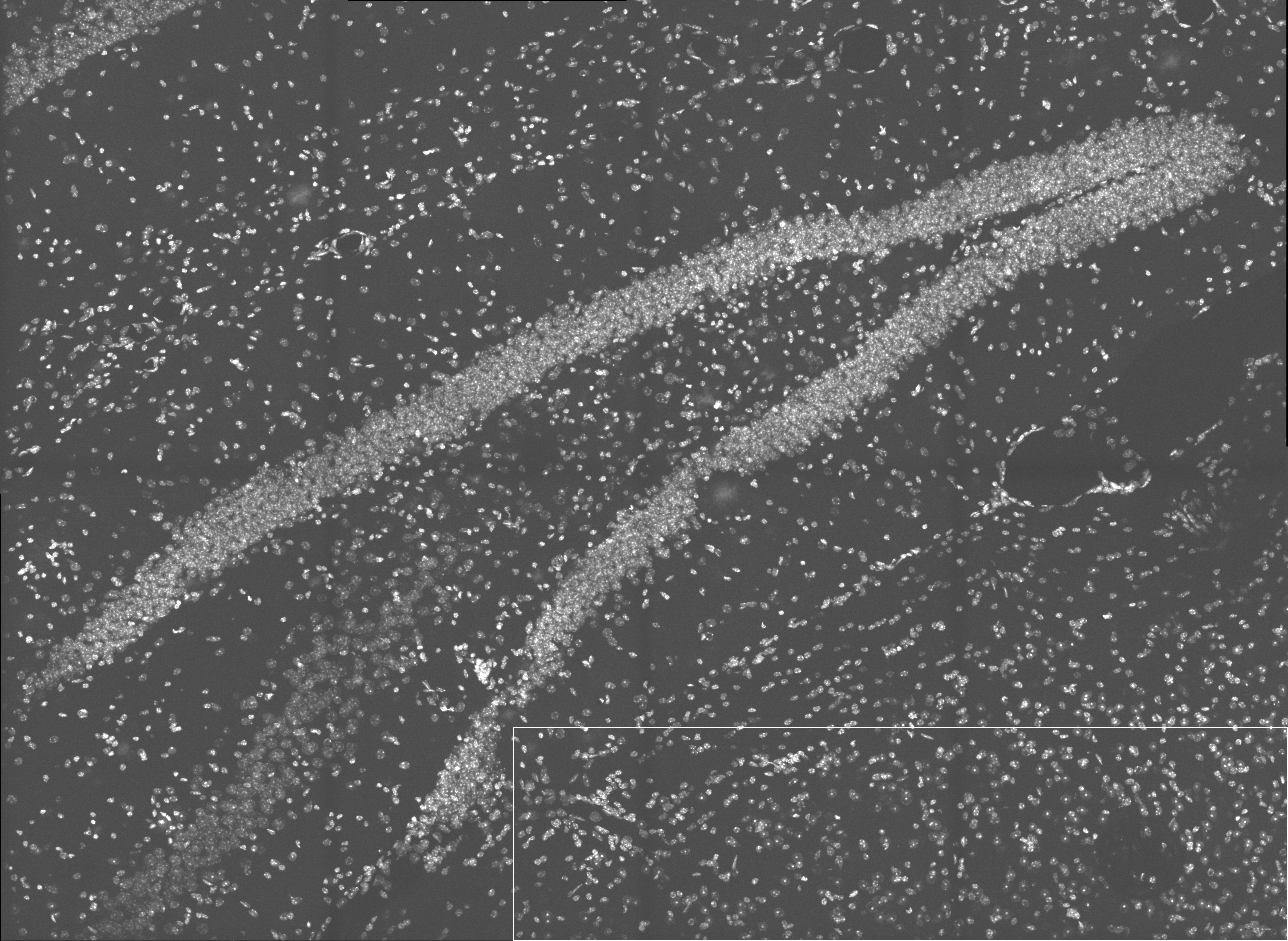

### Supp_Fig_2b_mouse_0_dapi.jpg

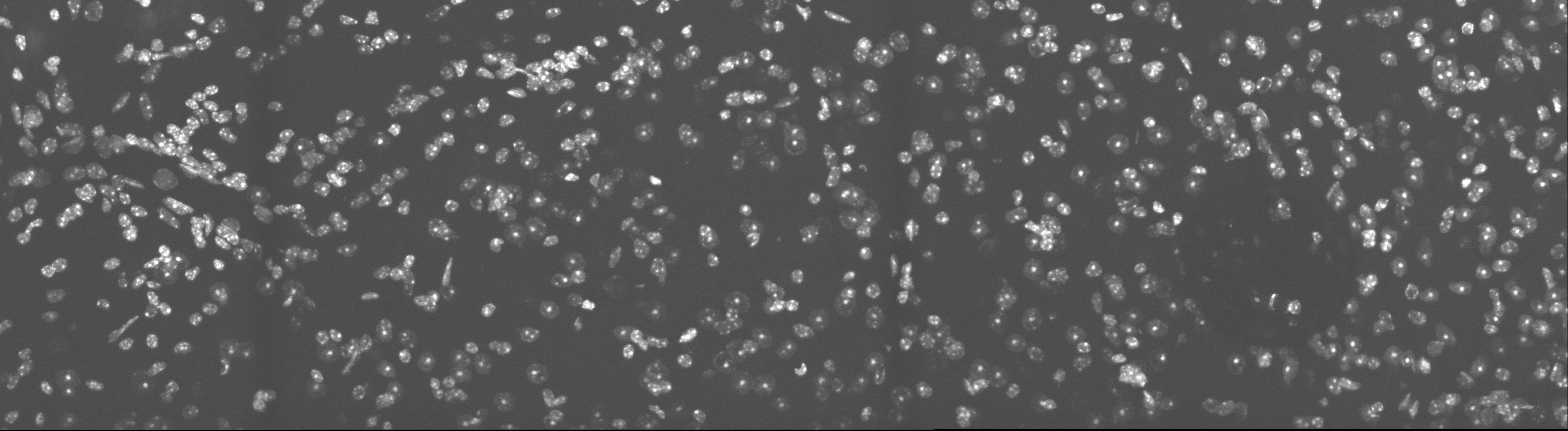

### Supp_Fig_2b_mouse_1_msk0.jpg

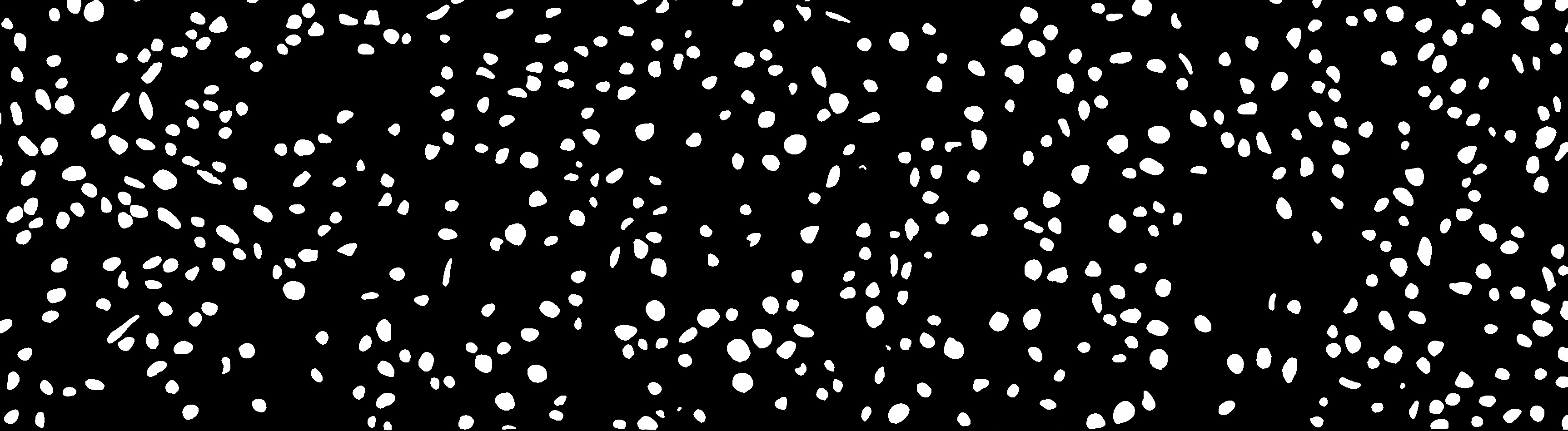

### Supp_Fig_2b_mouse_1_msk1.jpg

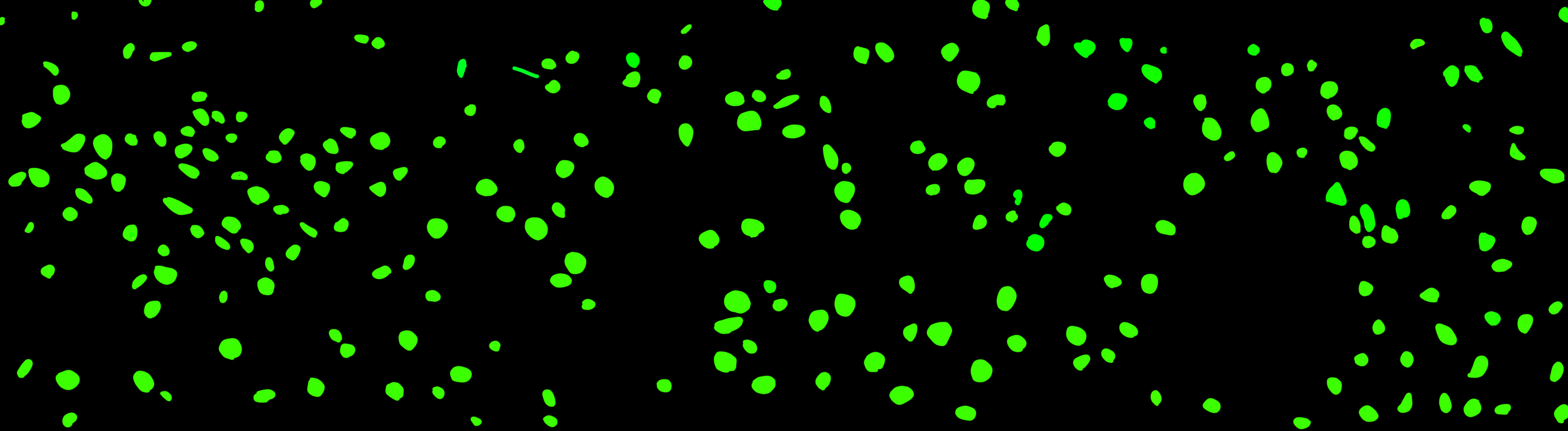

### Supp_Fig_2b_mouse_1_msk2.jpg

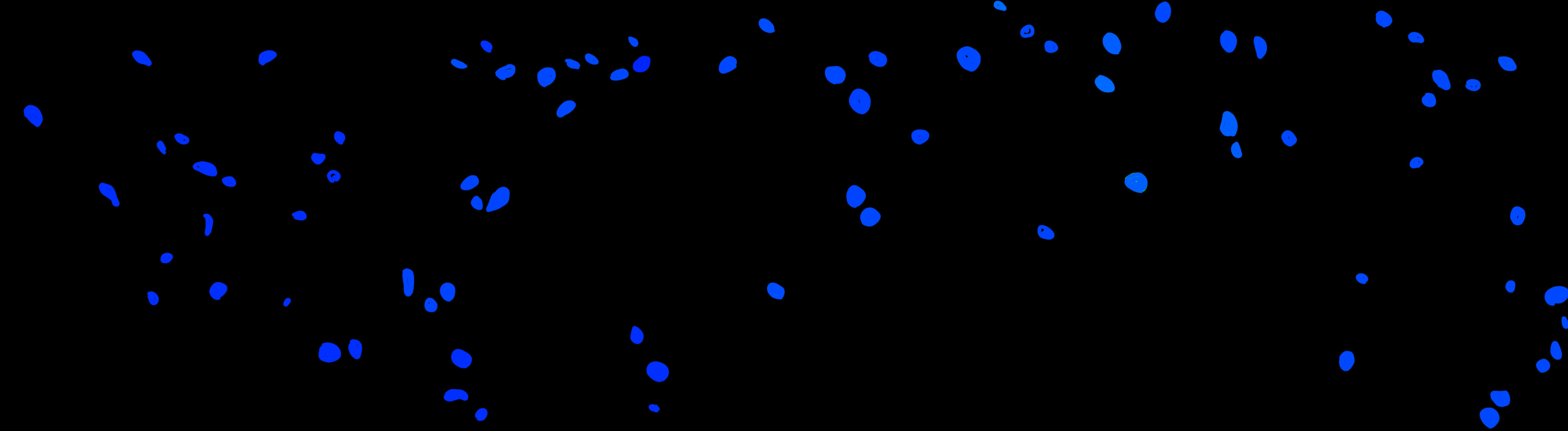

### Supp_Fig_2b_mouse_1_msk3.jpg

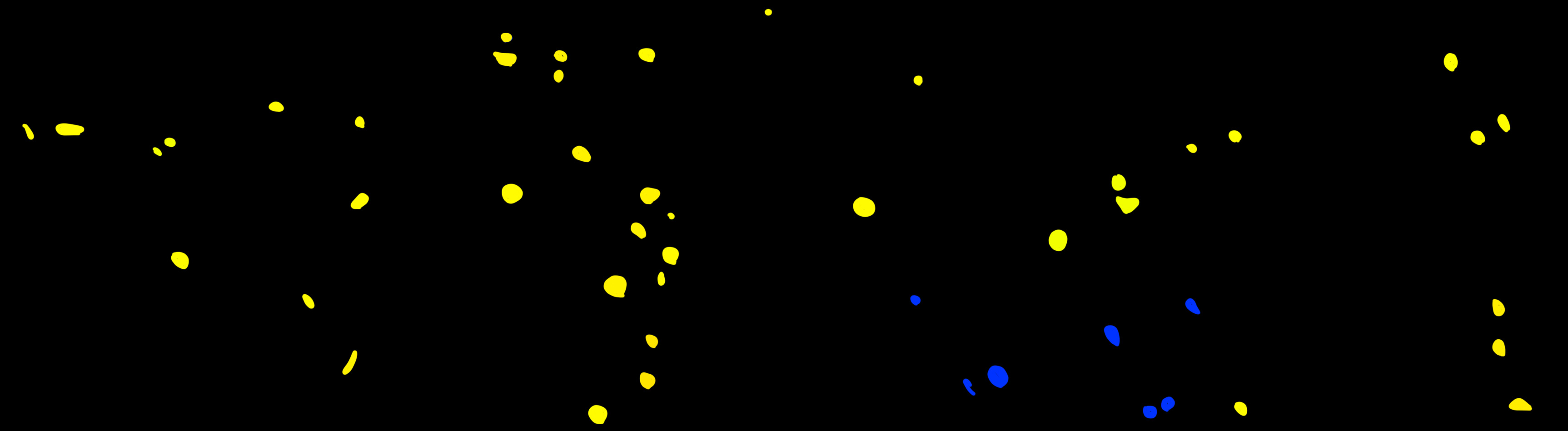

### Supp_Fig_3a_mouse.jpg

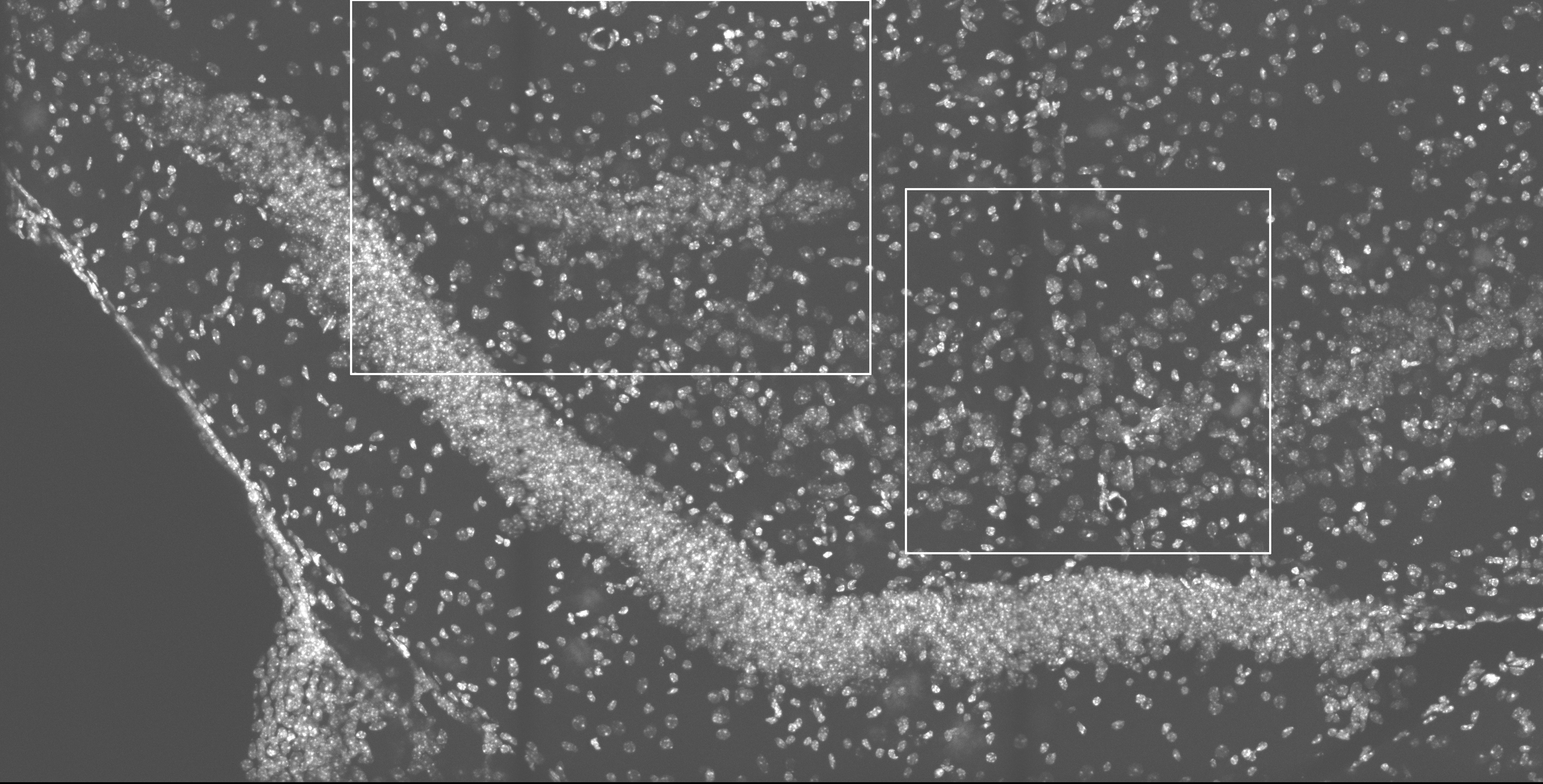

### Supp_Fig_3b_mouse_0_dapi.jpg

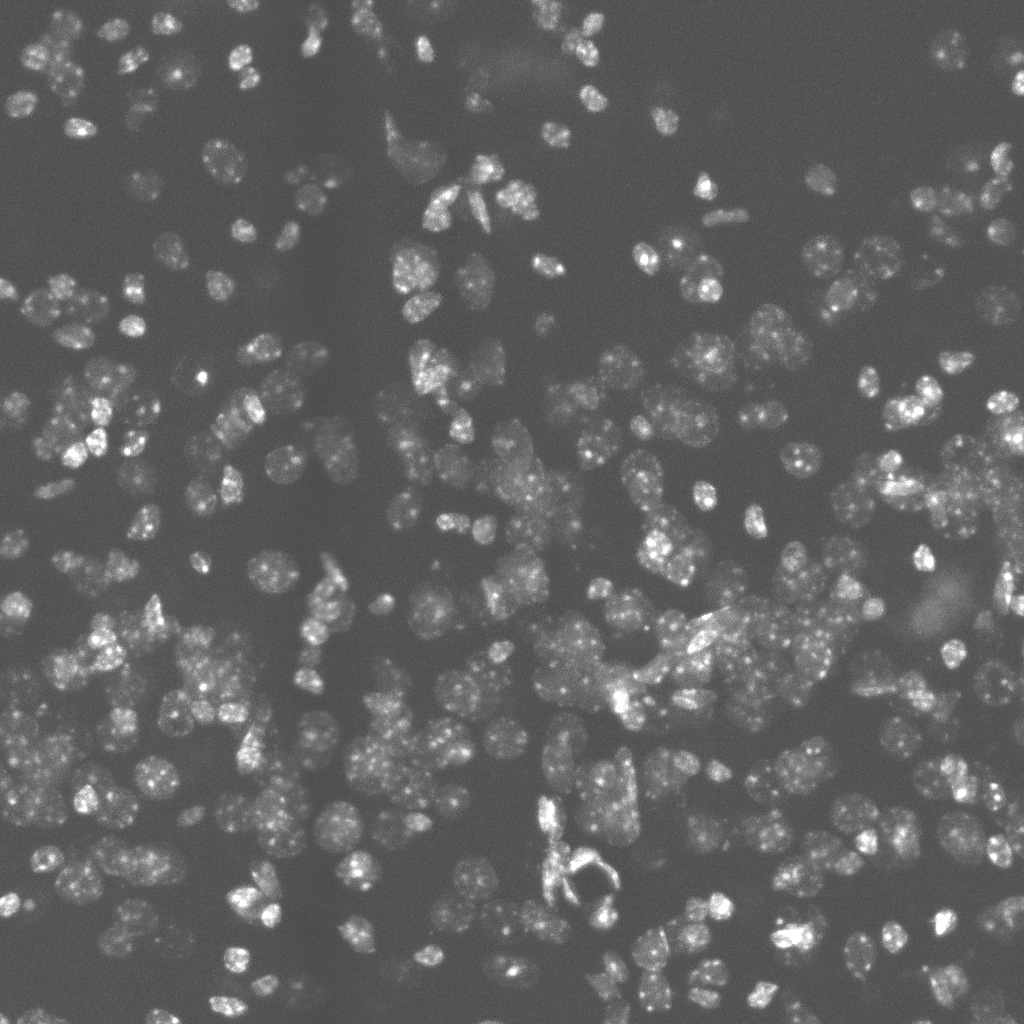

### Supp_Fig_3b_mouse_0_dapi.tif

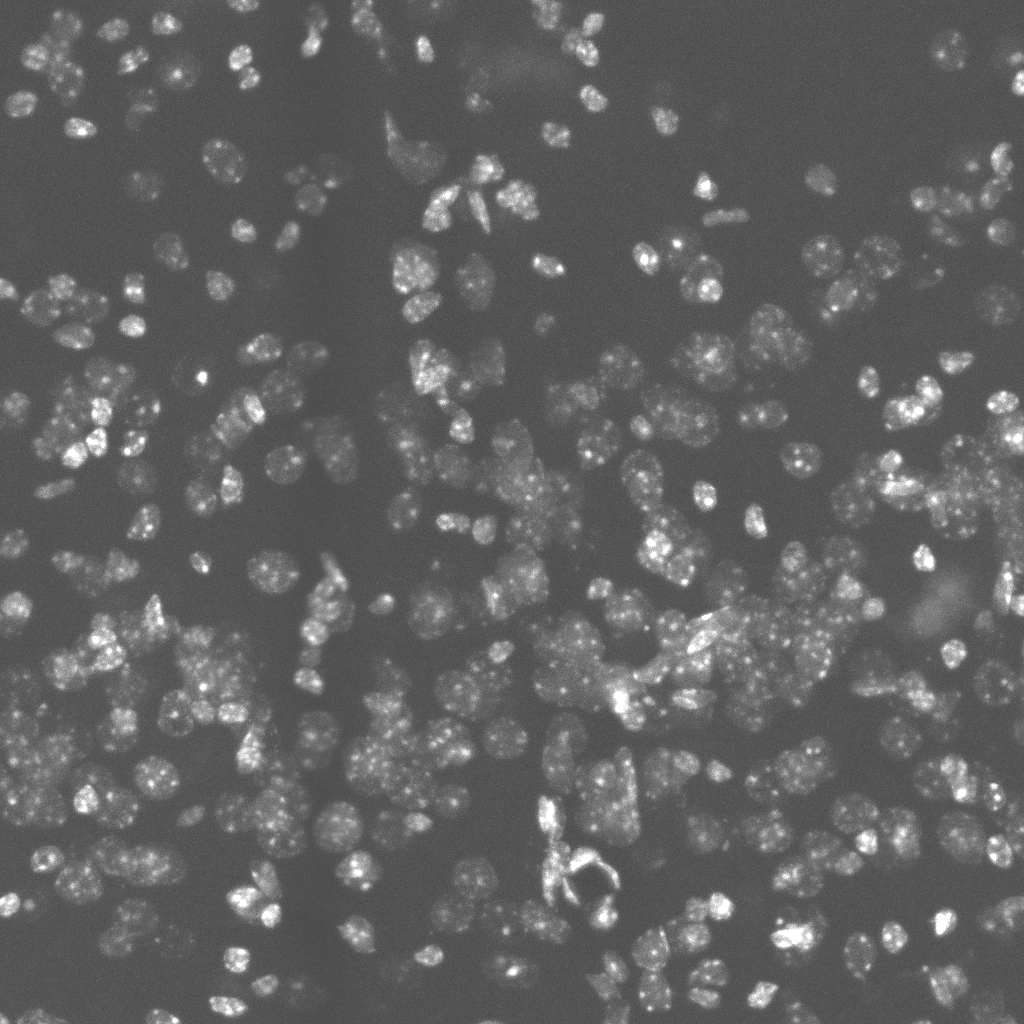

### Supp_Fig_3b_mouse_1_msk0.jpg

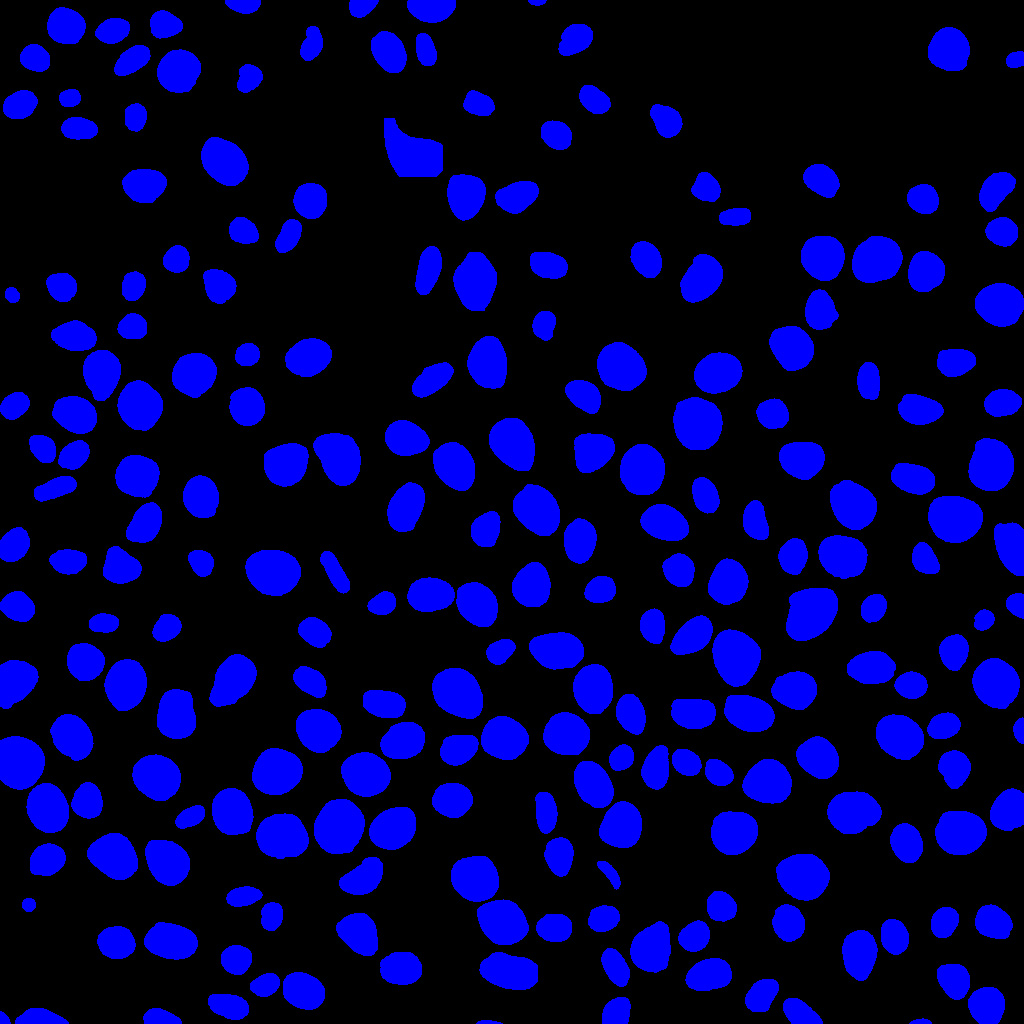

### Supp_Fig_3b_mouse_1_msk1.jpg

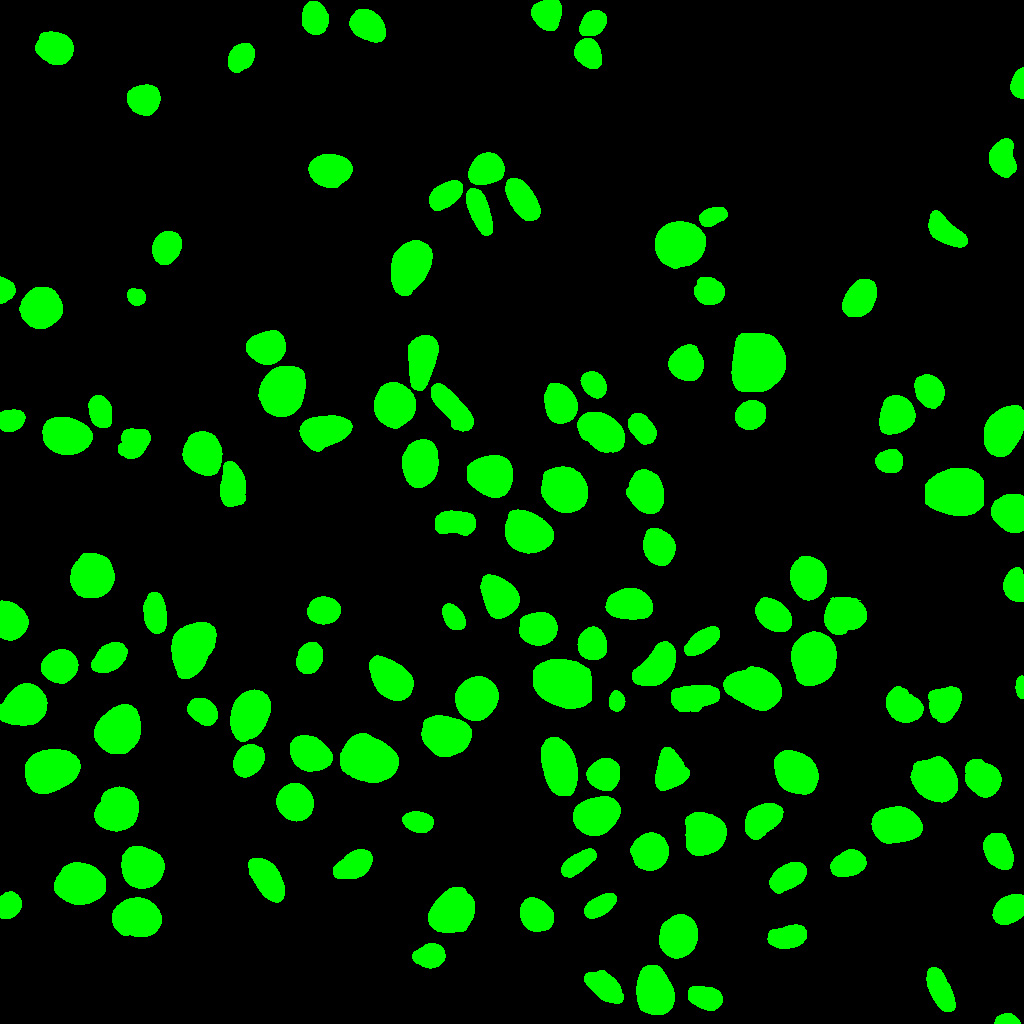

### Supp_Fig_3b_mouse_1_msk2.jpg

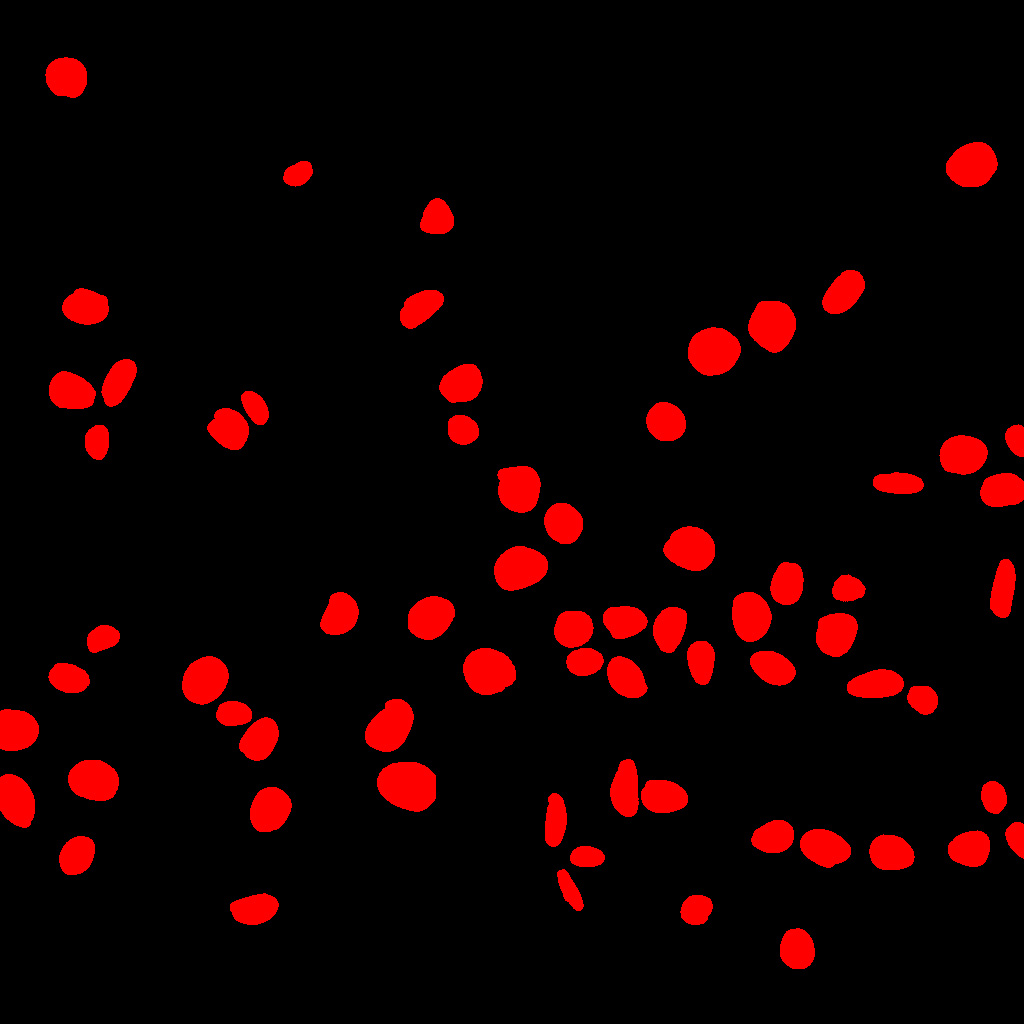

### Supp_Fig_3b_mouse_1_msk3.jpg

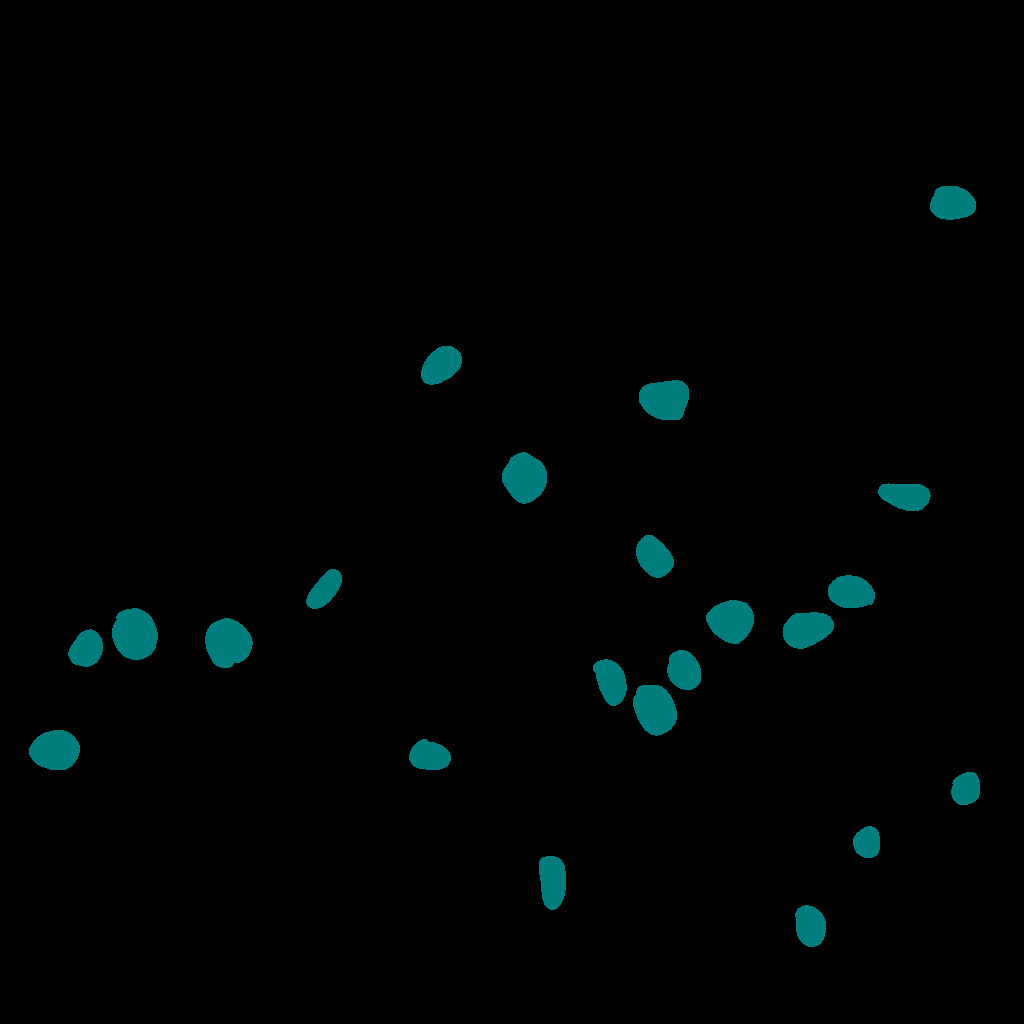

### Supp_Fig_3c_mouse_0_dapi.jpg

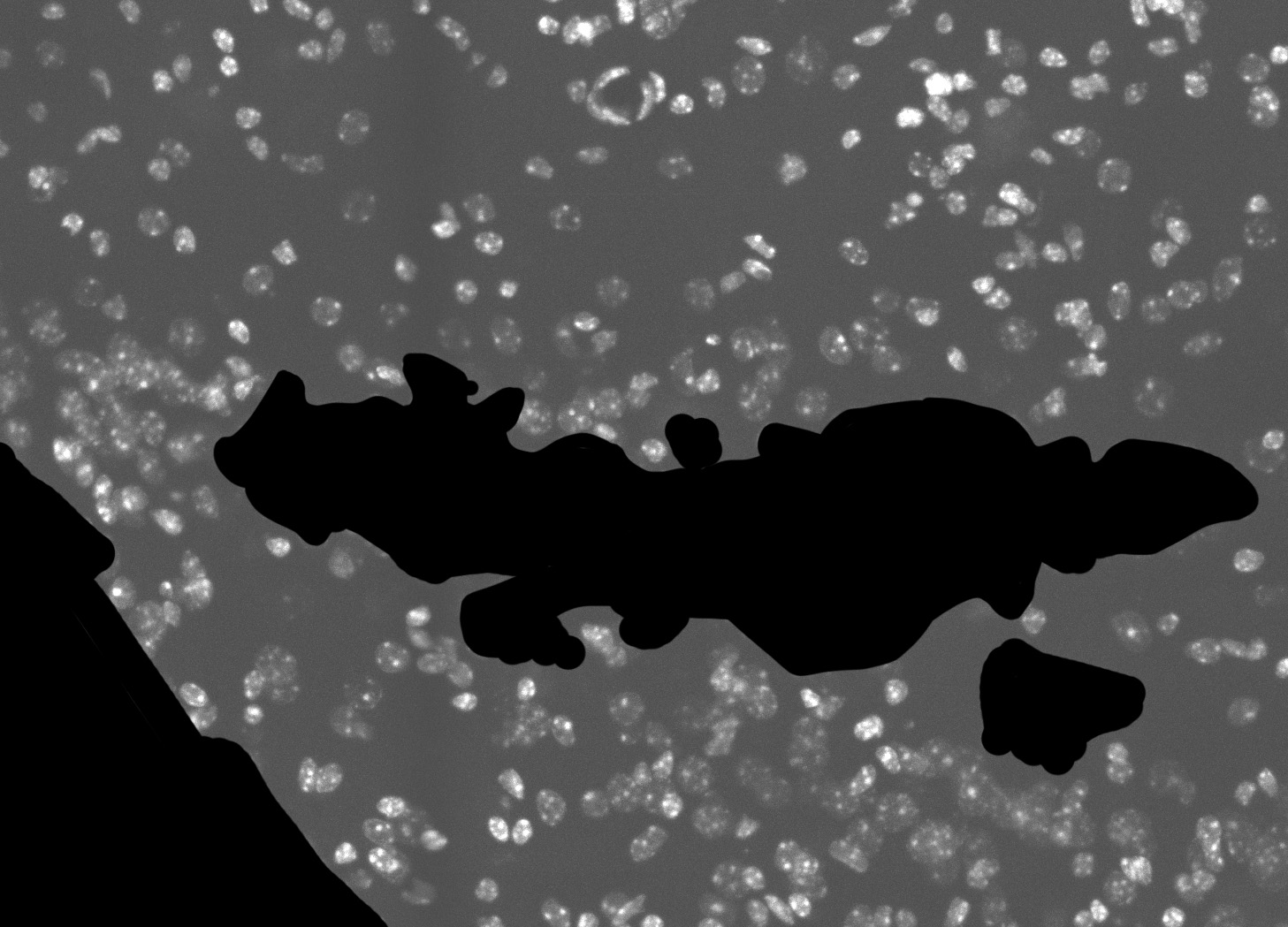

### Supp_Fig_3c_mouse_1_msk0.jpg

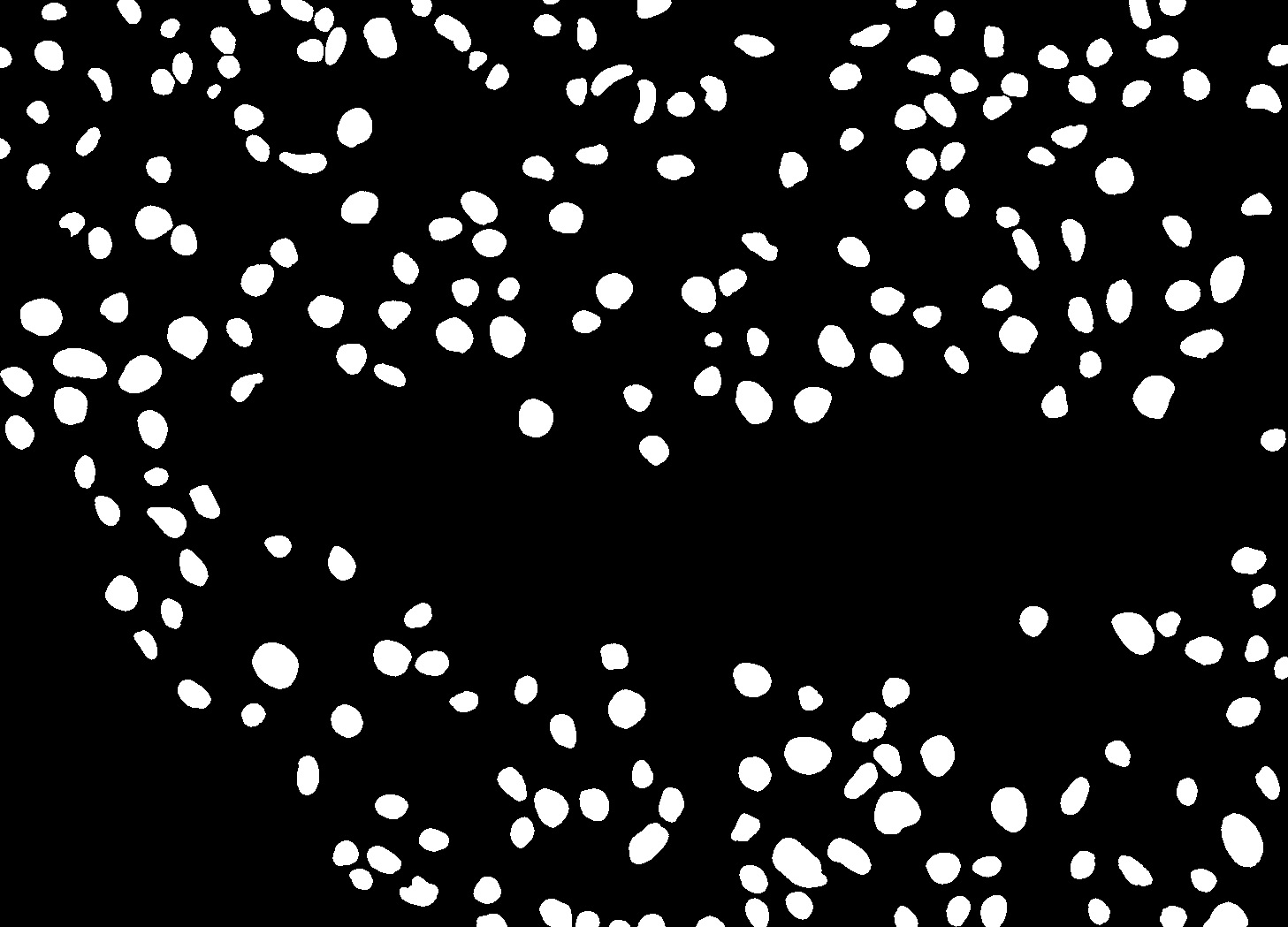

### Supp_Fig_3c_mouse_1_msk1.jpg

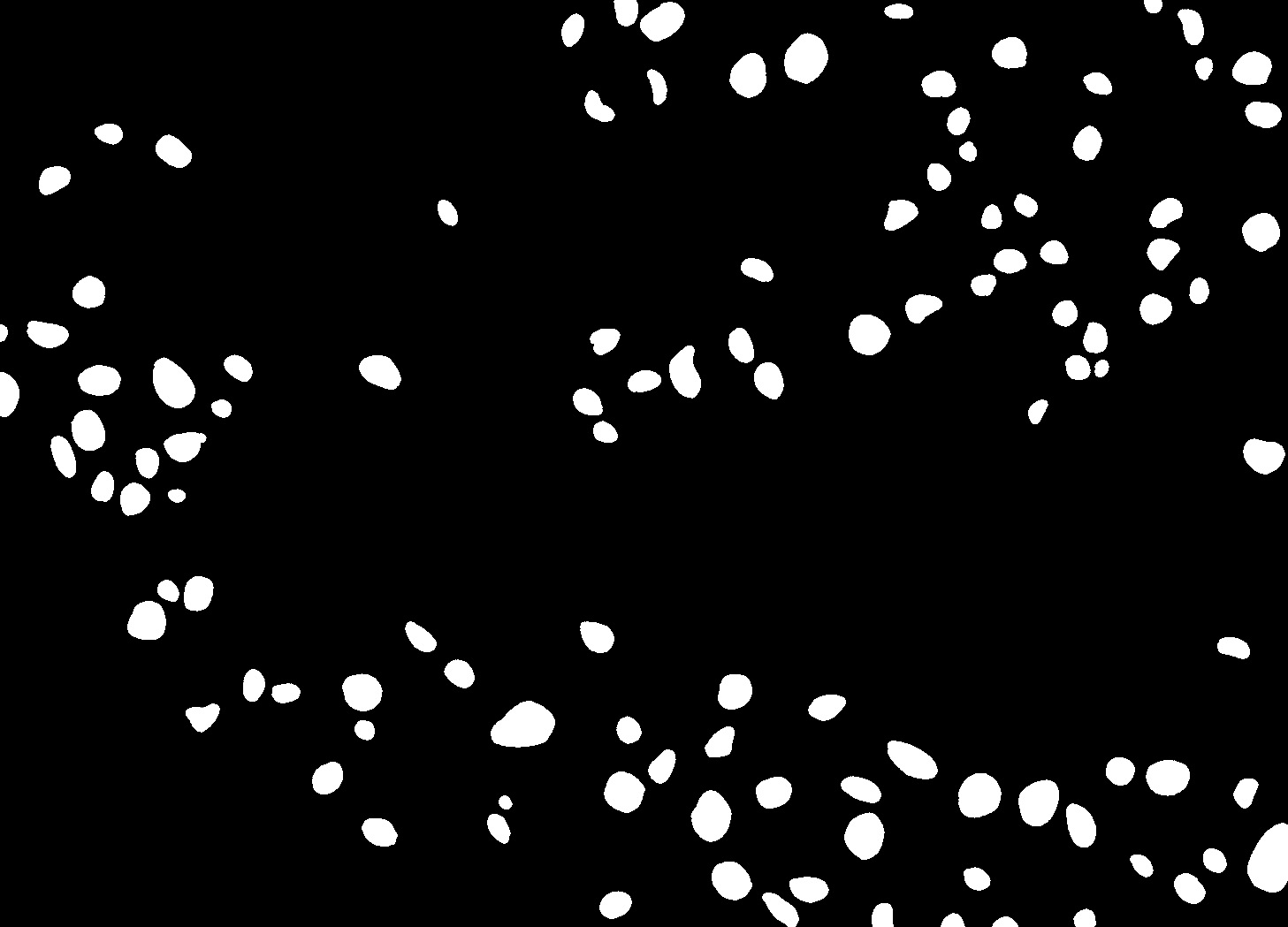

### Supp_Fig_3c_mouse_1_msk2.jpg

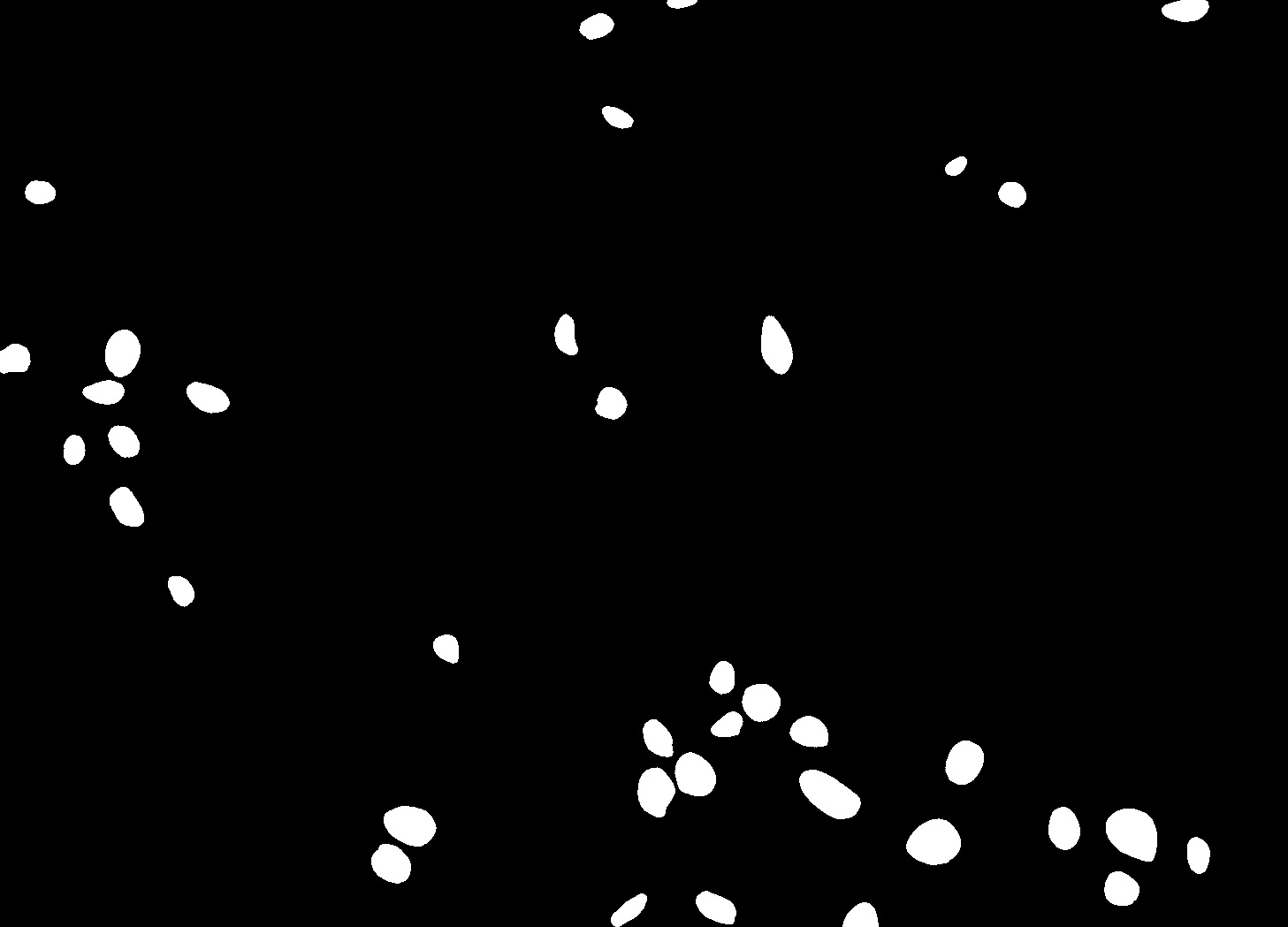

### Supp_Fig_3c_mouse_1_msk3.jpg

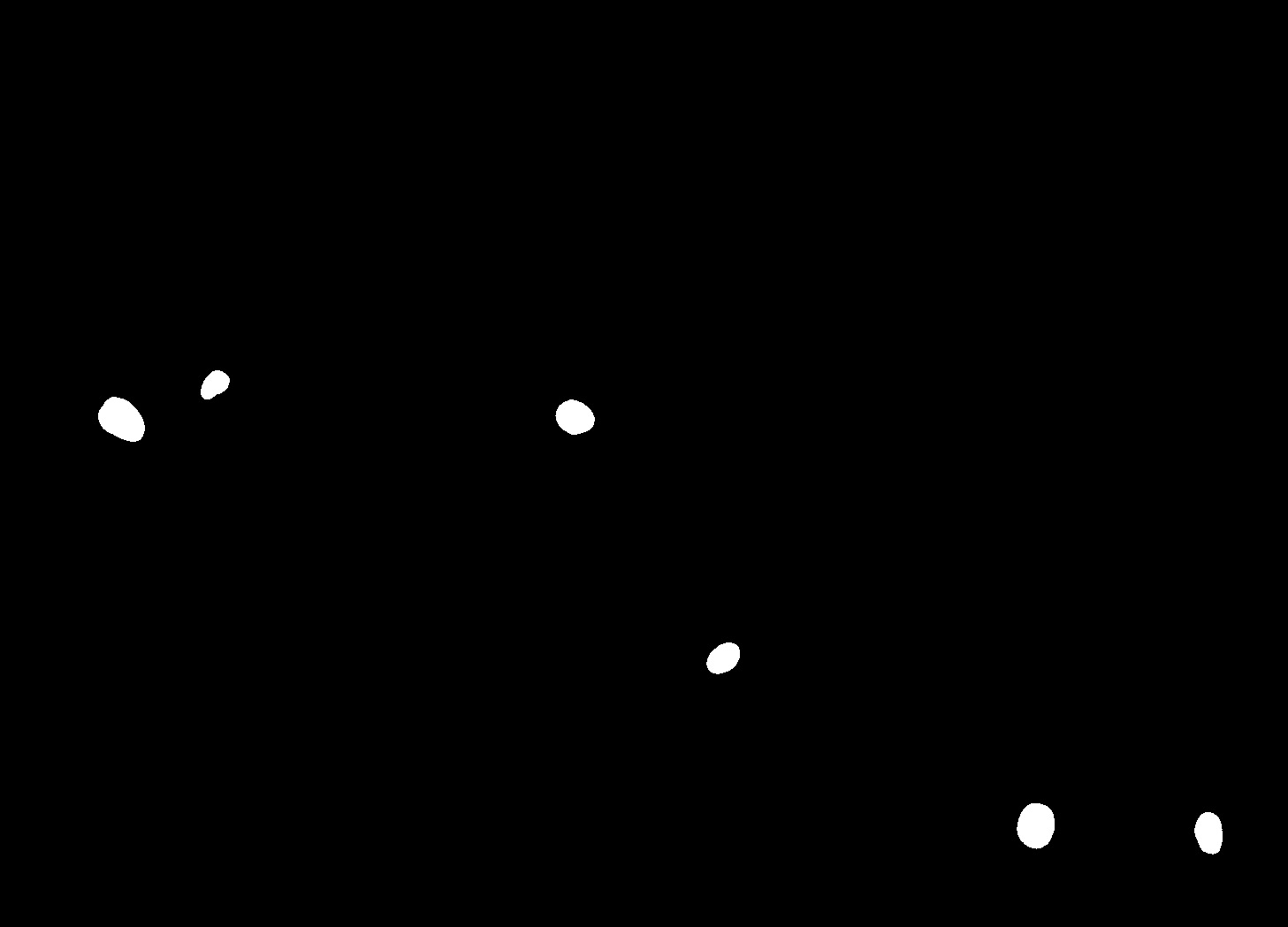

### Supp_Fig_4a_4ch.jpg

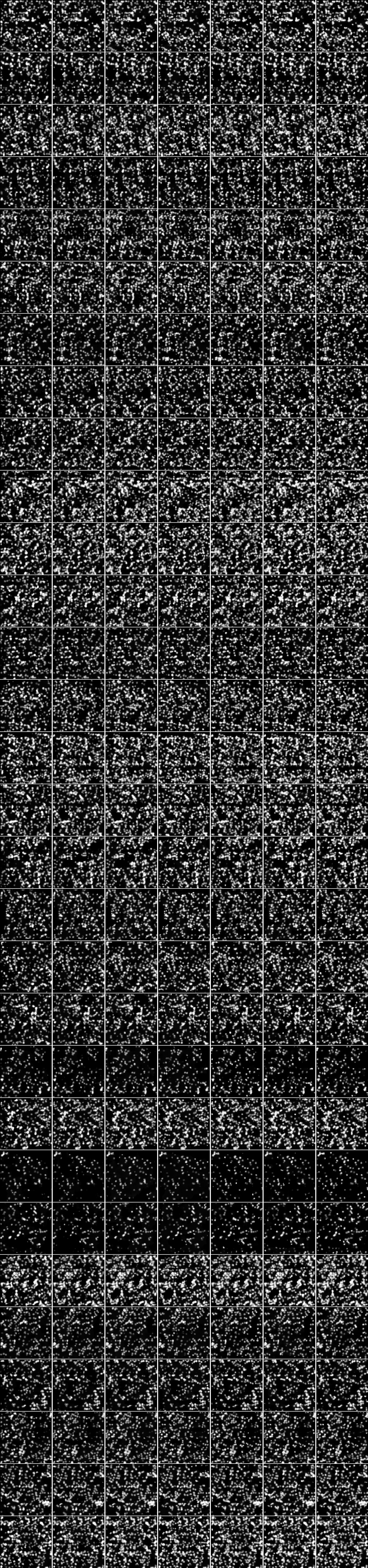

### Supp_Fig_4a_5ch.jpg

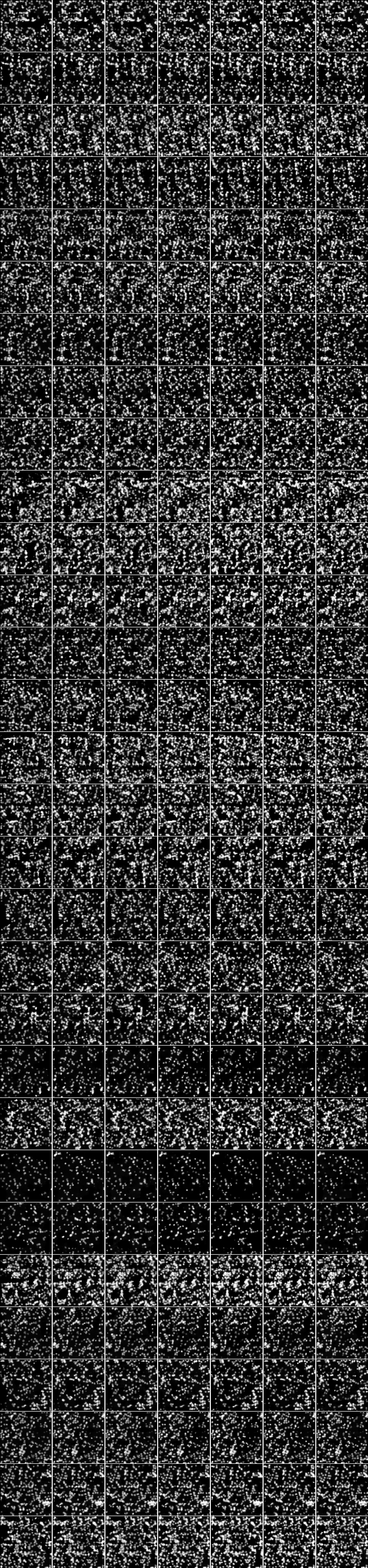

### Supp_Fig_4b_4ch.jpg

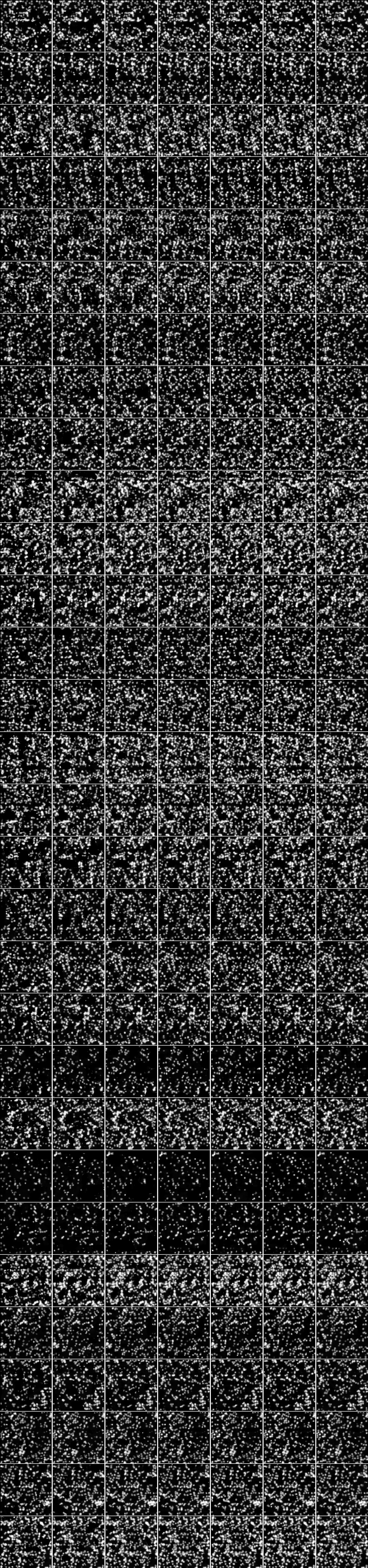

### Supp_Fig_4b_5ch.jpg

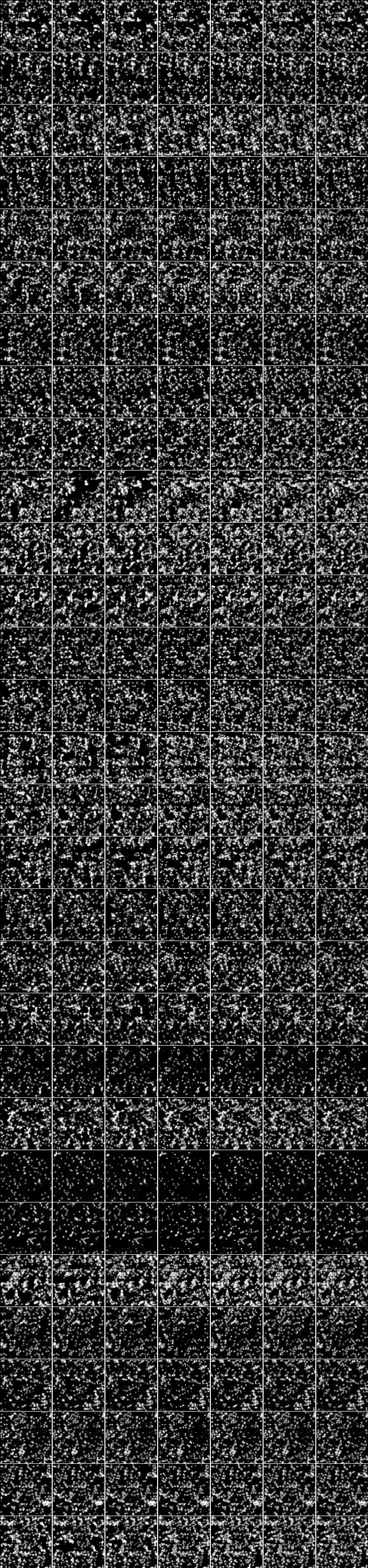

### Supp_Fig_4c_4ch.jpg

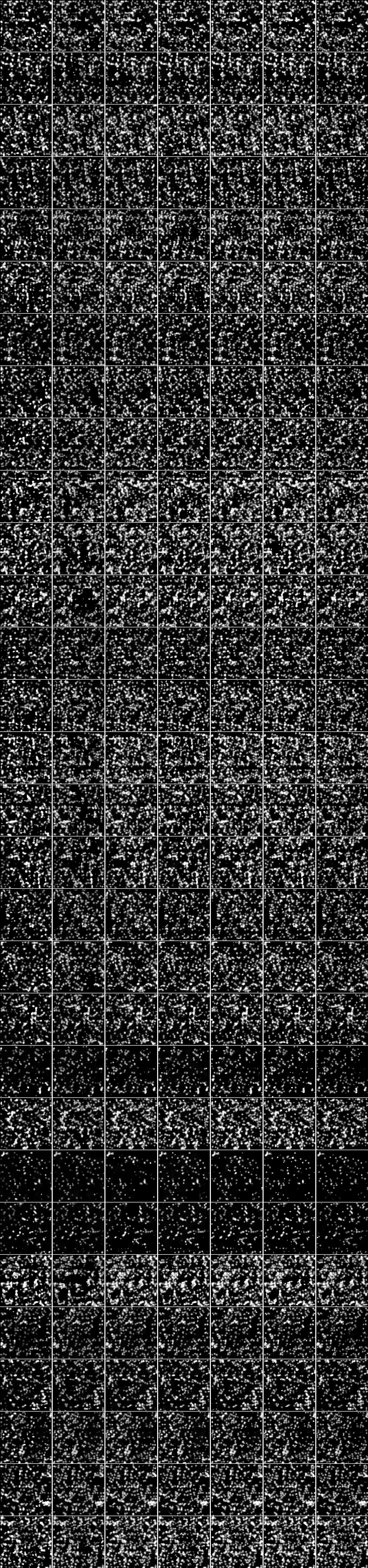

### Supp_Fig_4c_5ch.jpg

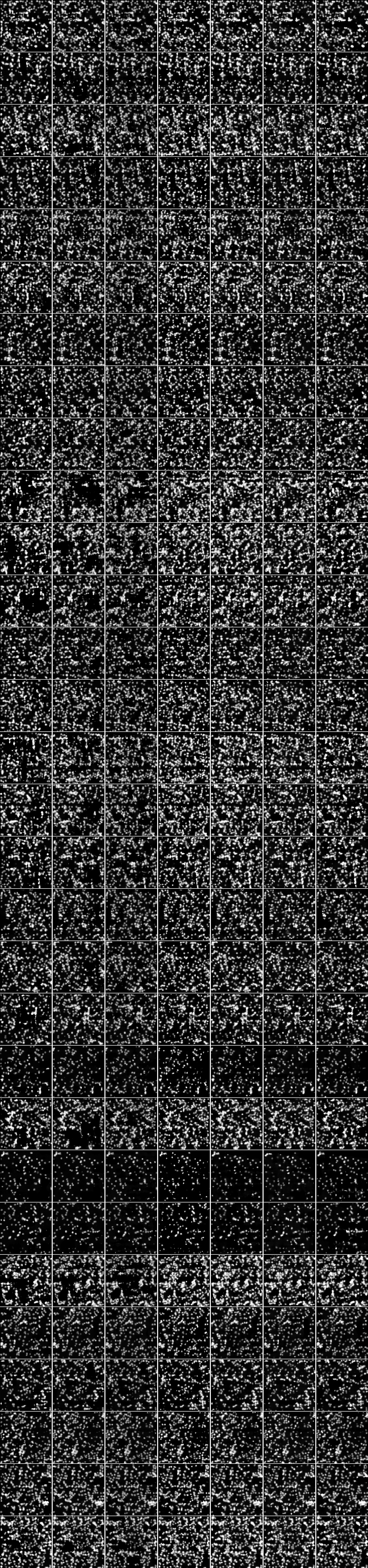

### Supp_Fig_4d_4ch.jpg

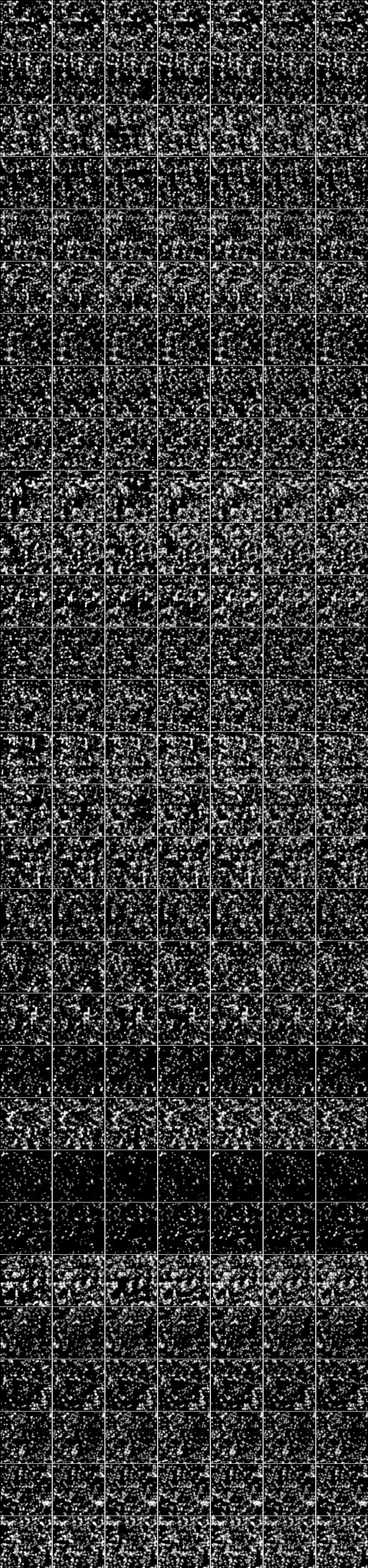

### Supp_Fig_4d_5ch.jpg

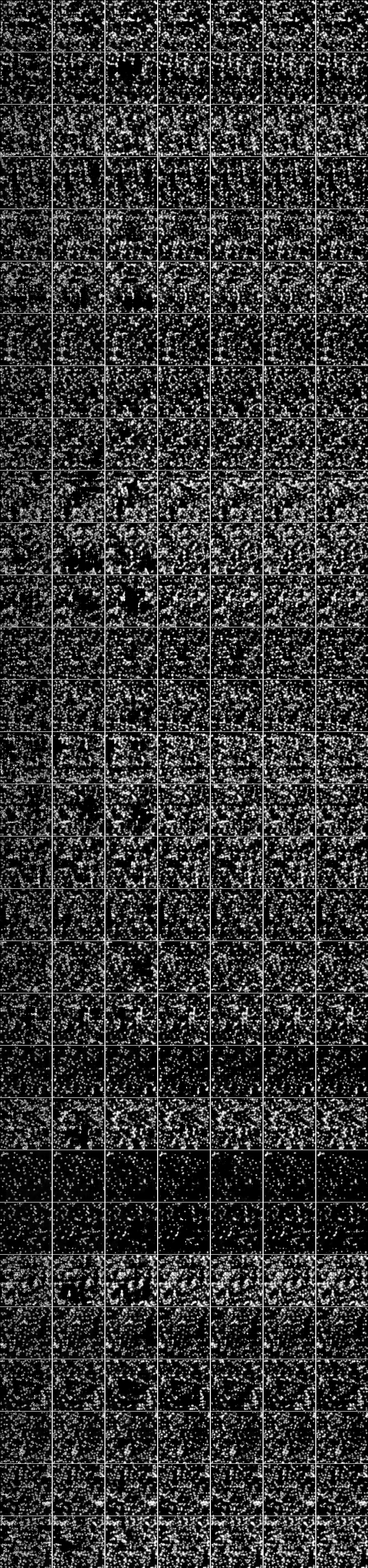

### Supp_Fig_5_unet4.jpg

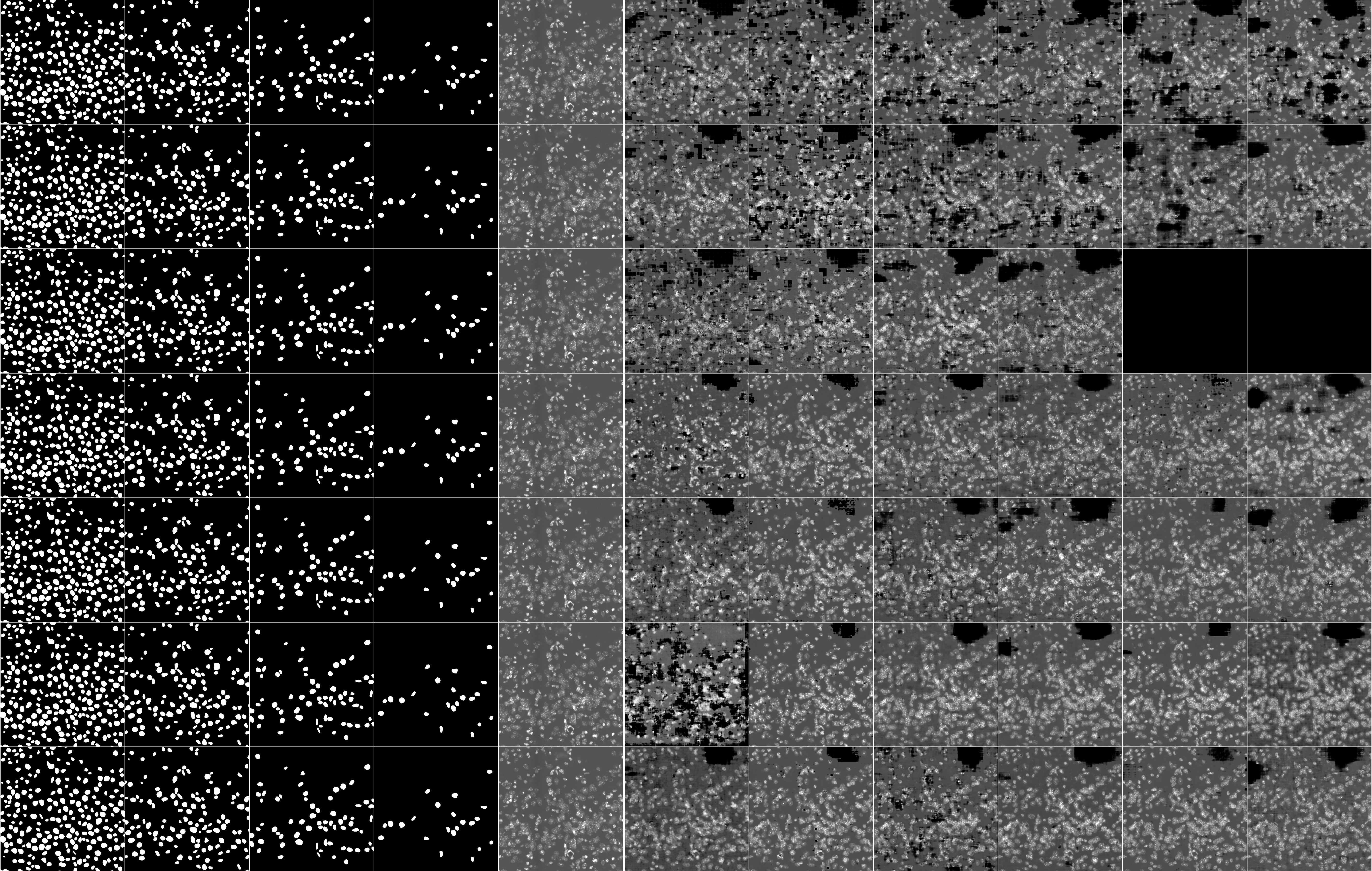

### Supp_Fig_5_unet6.jpg

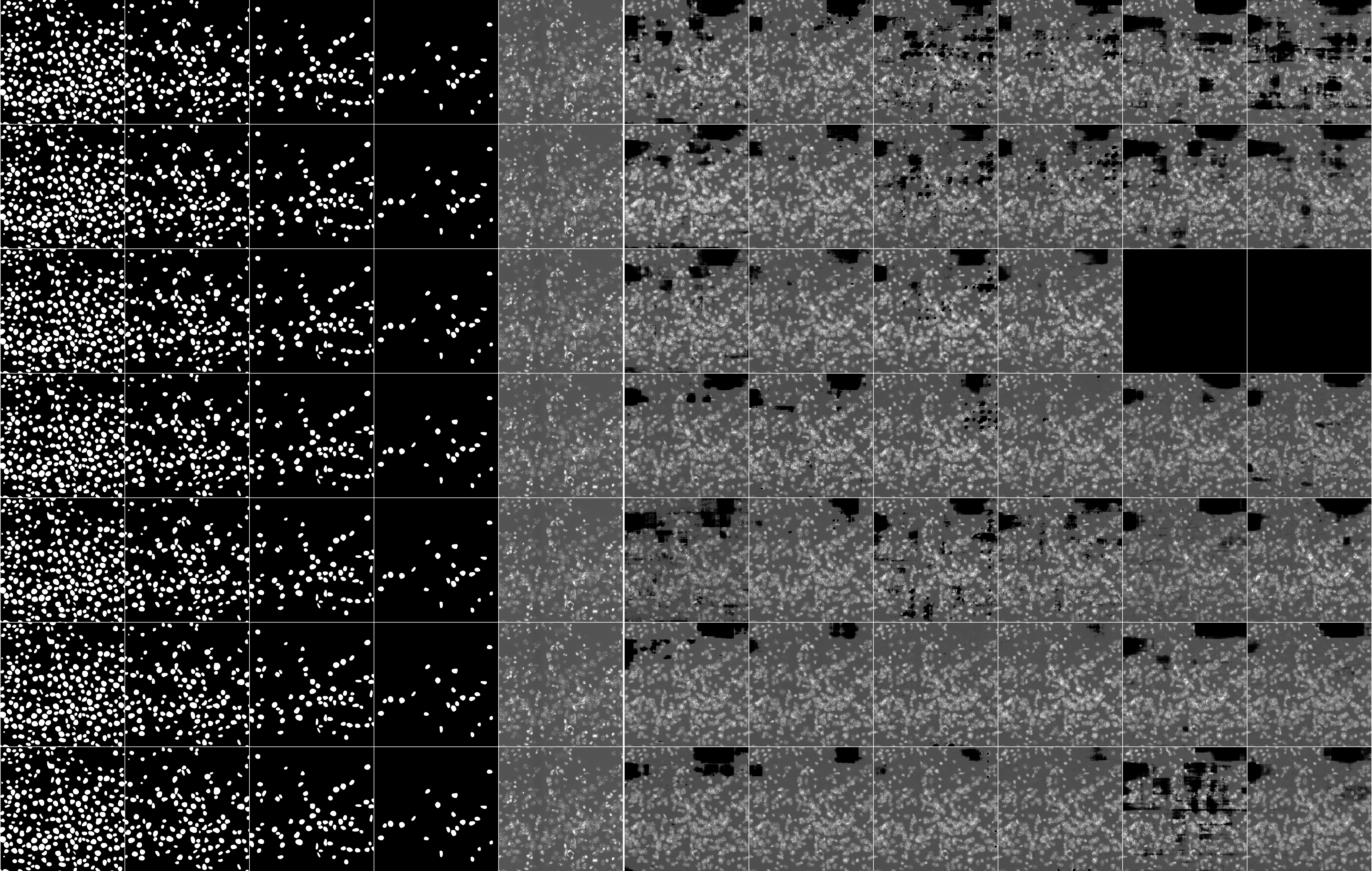

### Supp_Fig_6a_unet4_1024x1024customcells.jpg

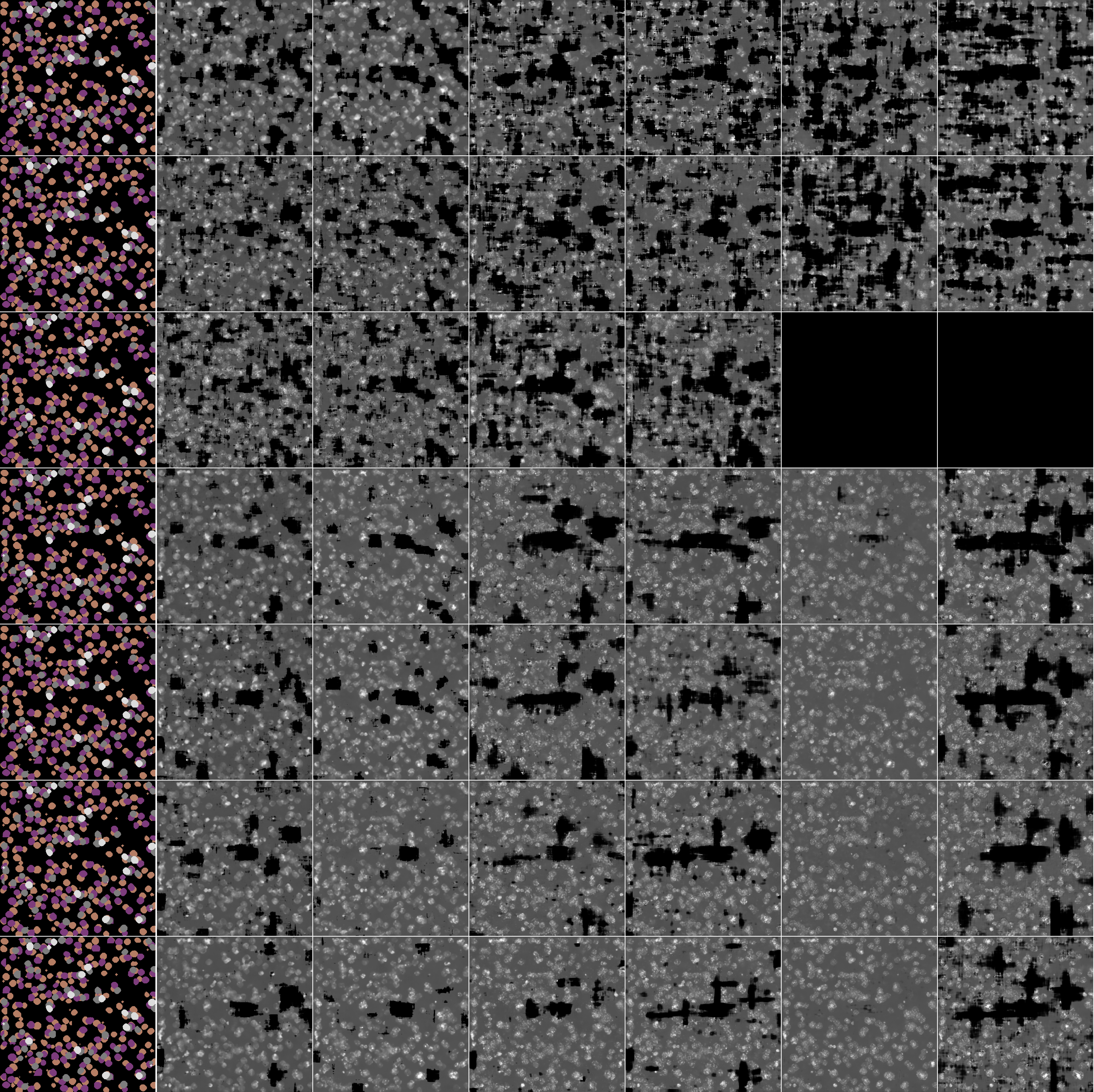

### Supp_Fig_6b_unet6_1024x1024customcells.jpg

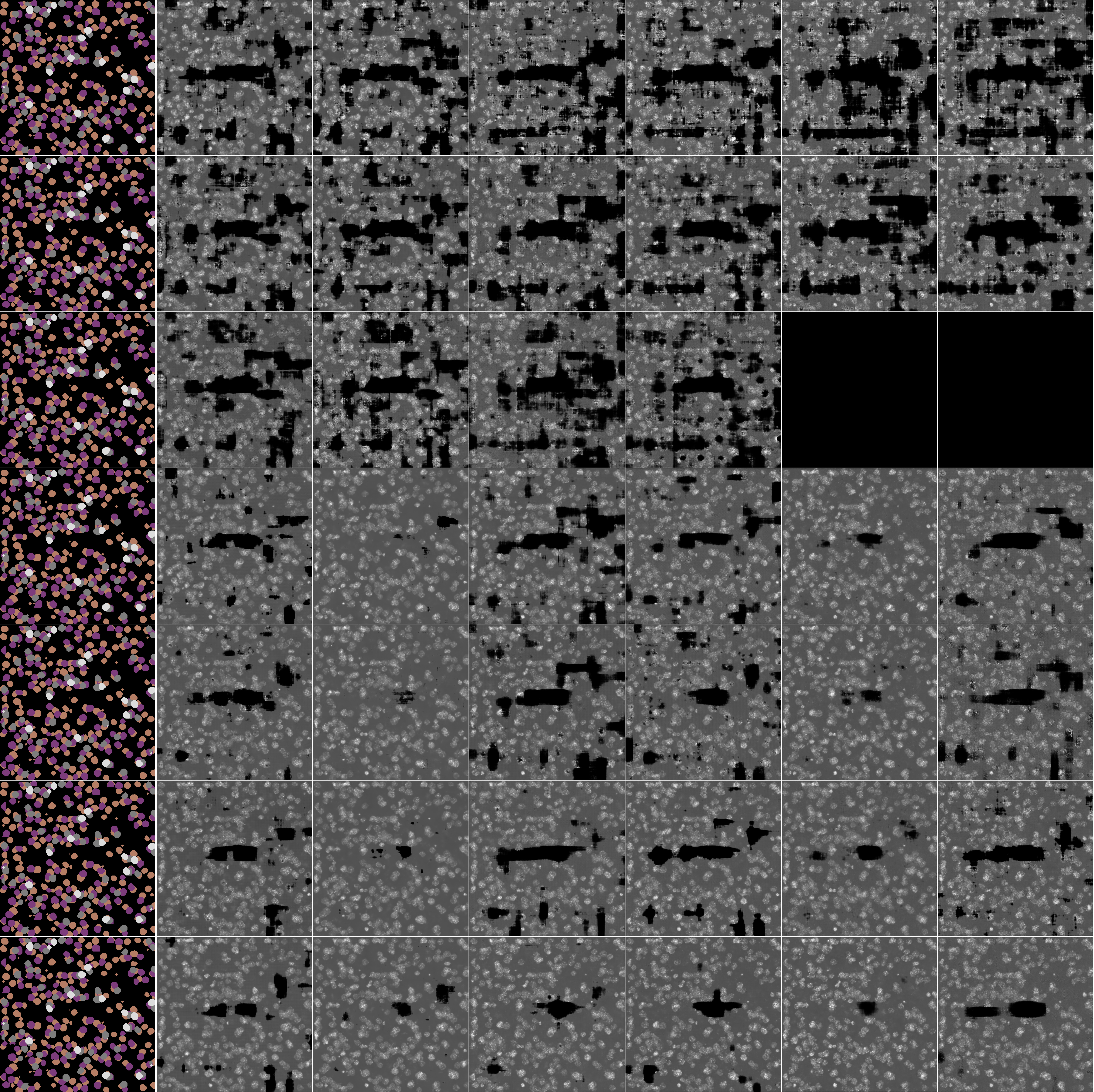

### Supp_Fig_7a_unet4_1024x1024customcells.jpg

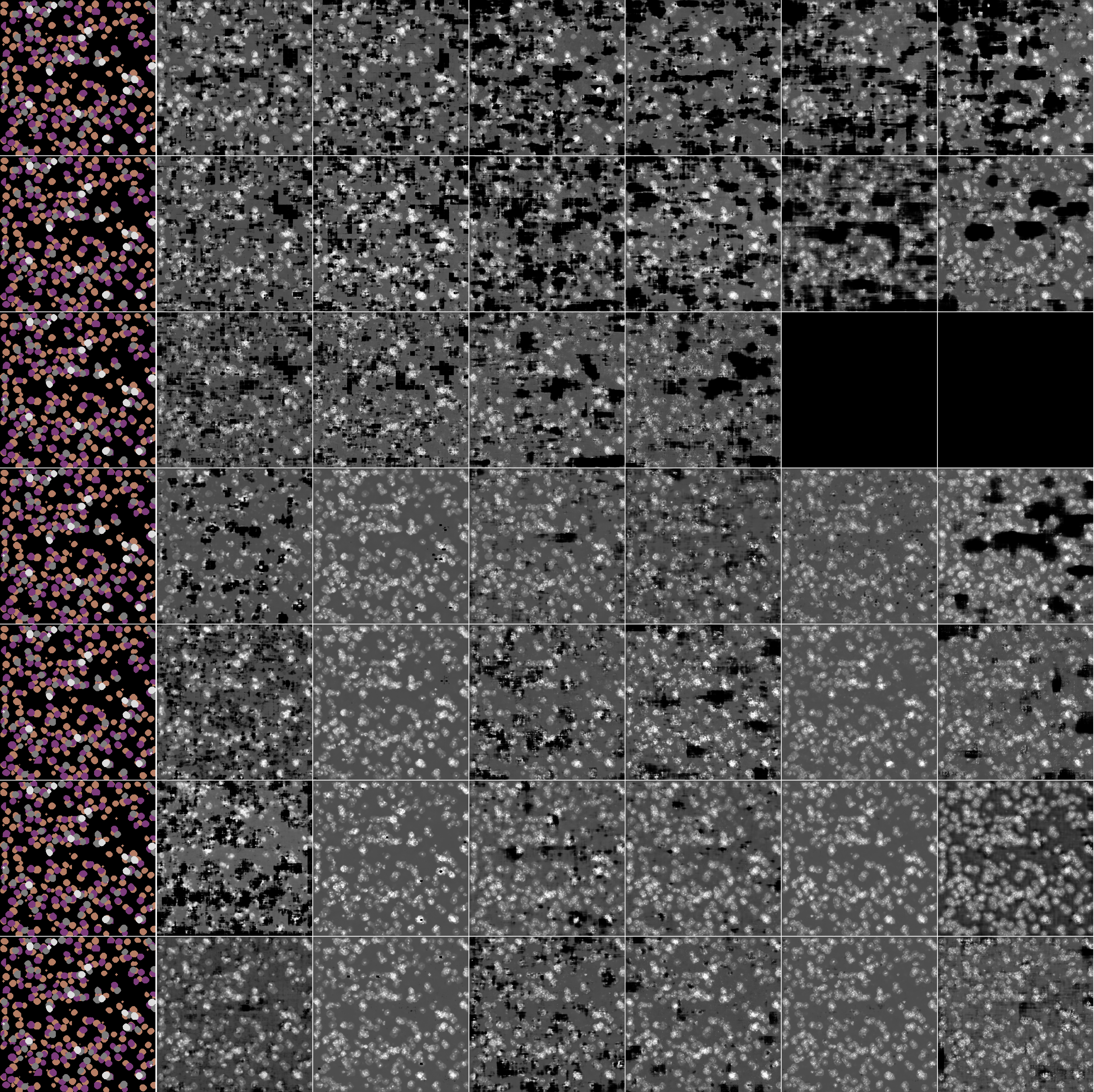

### Supp_Fig_7b_unet6_1024x1024customcells.jpg

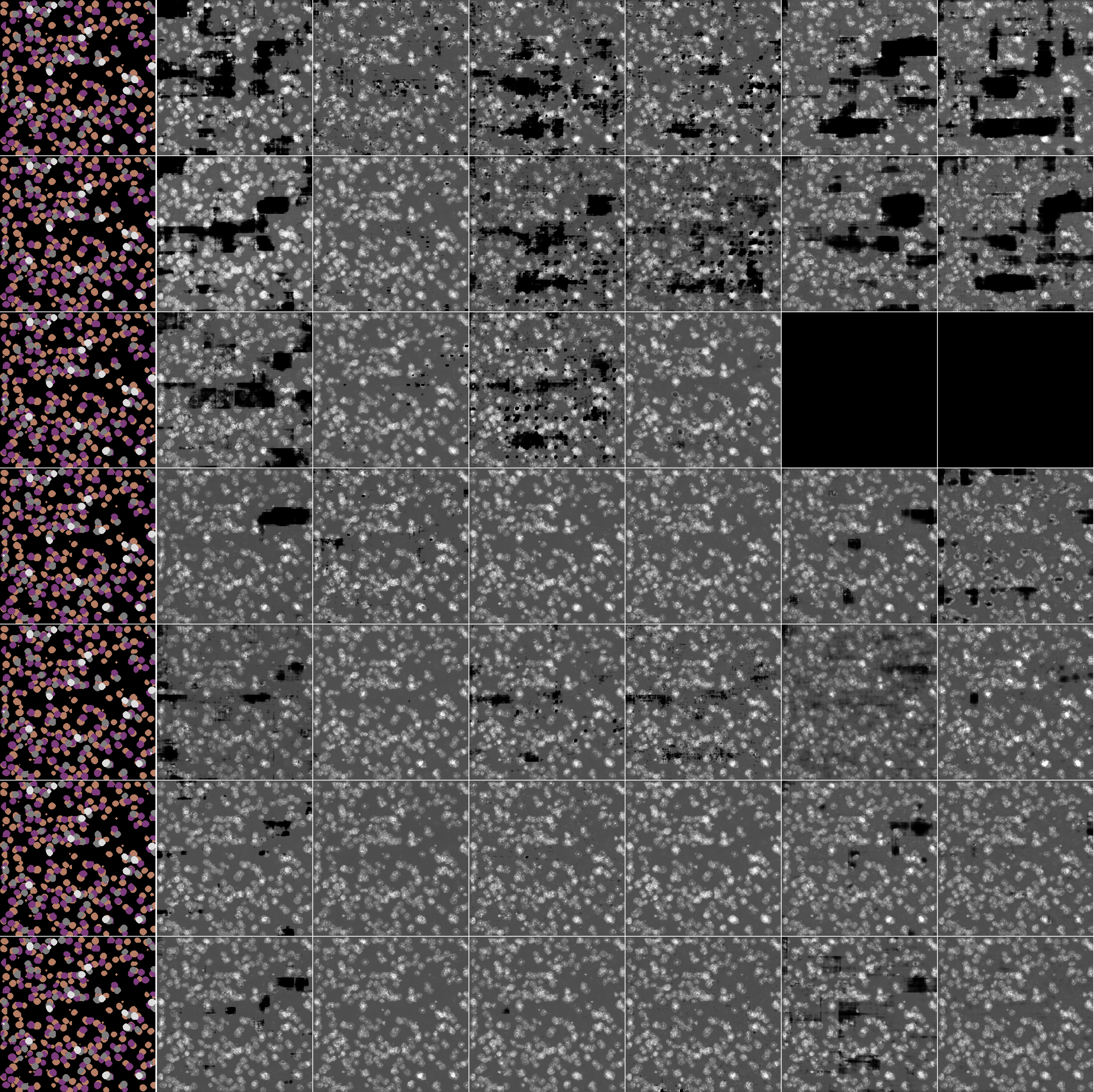

### Supp_Fig_8a_unet4_1024x1024.jpg

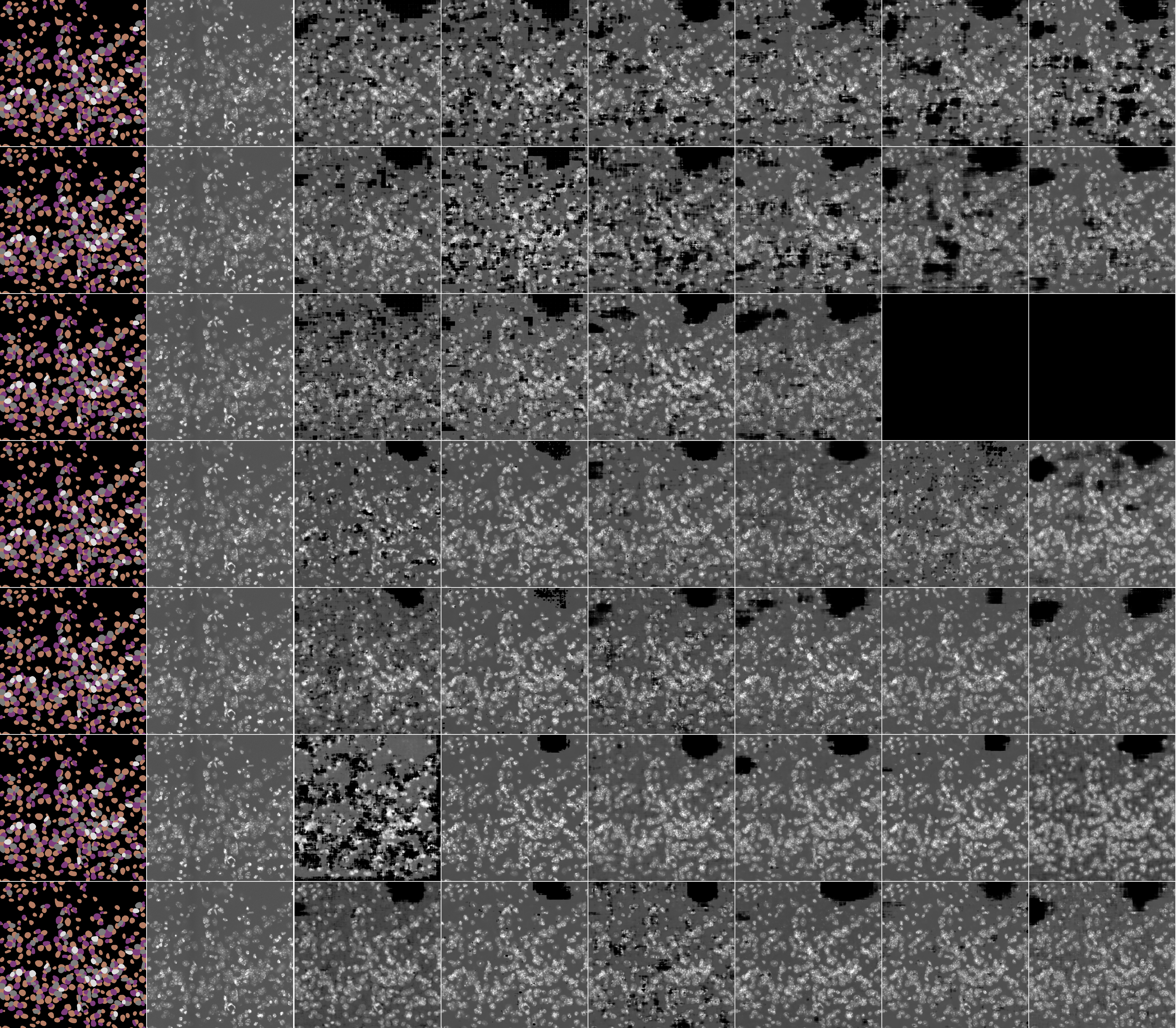

### Supp_Fig_8b_unet6_1024x1024.jpg

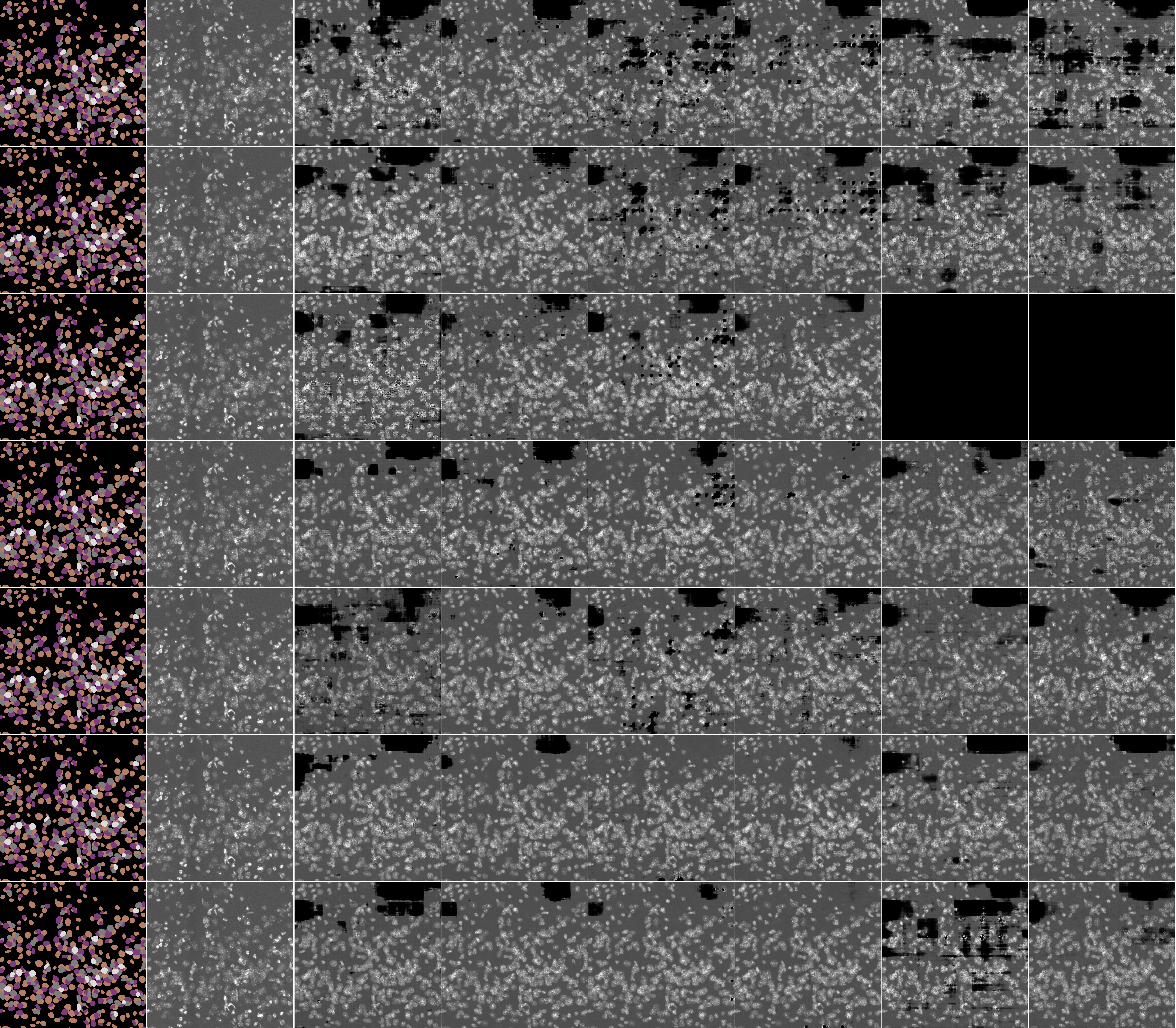

### Supp_Fig_9a_unet4.jpg

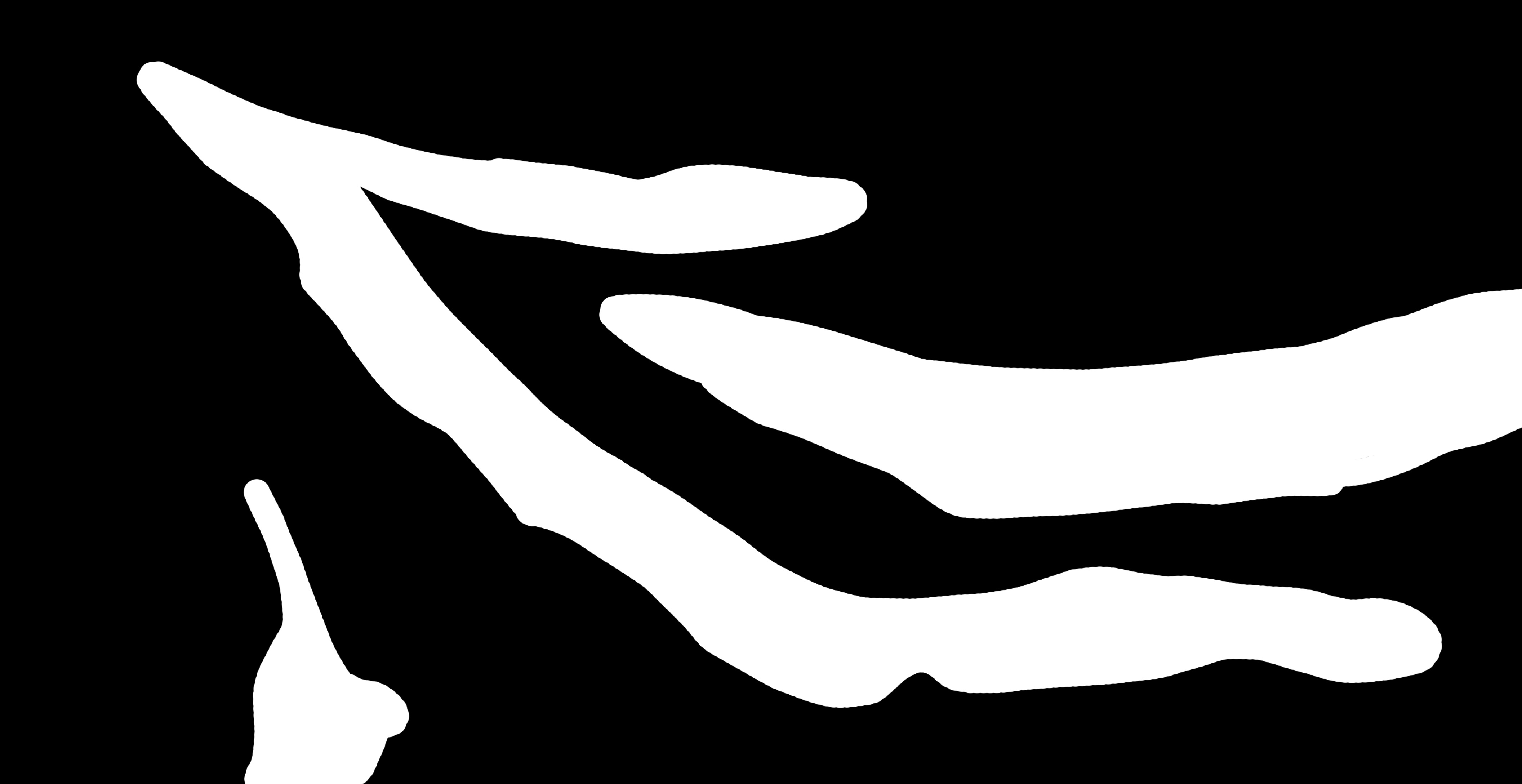

### Supp_Fig_9b_unet4.jpg

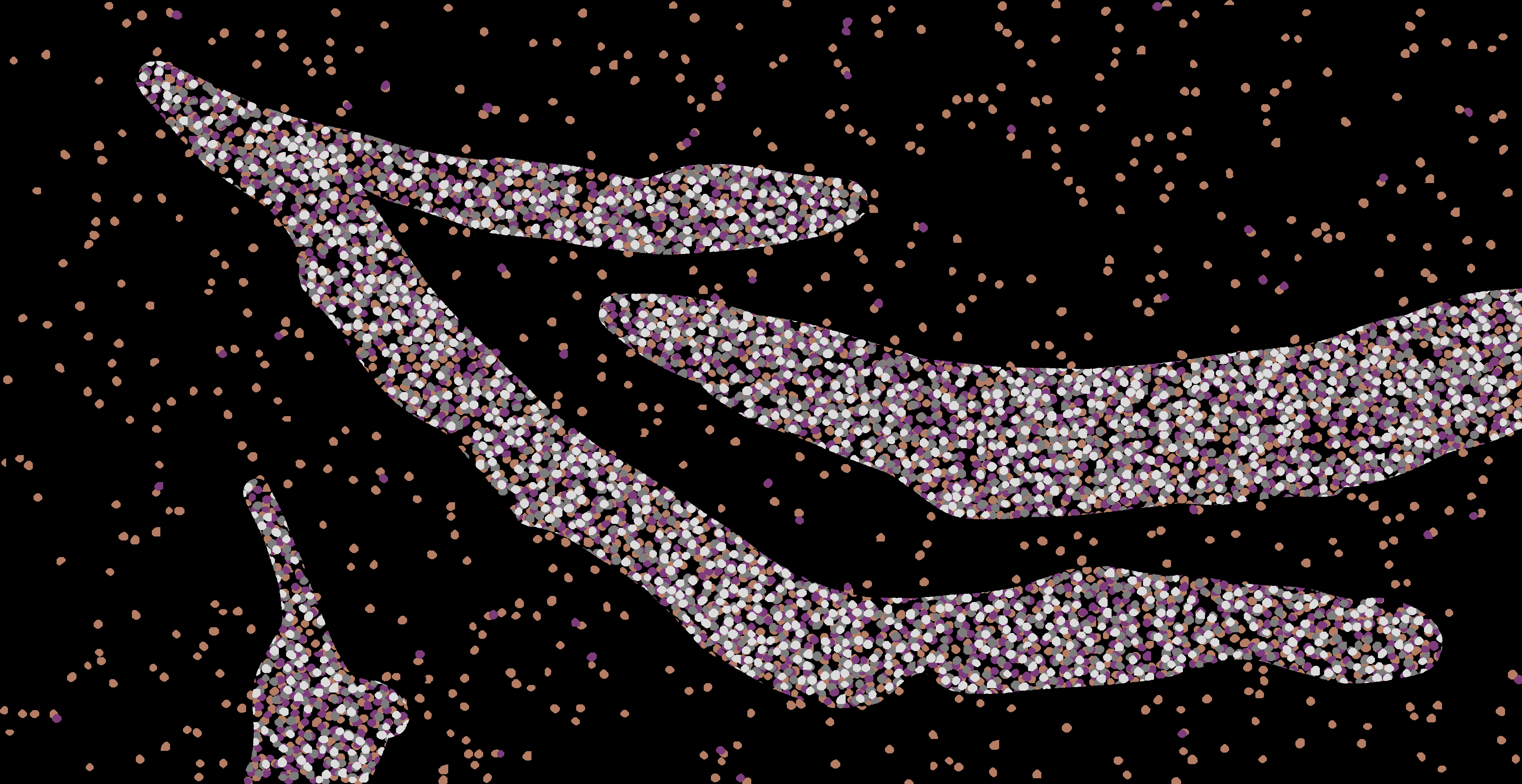

### Supp_Fig_9c_unet4.jpg

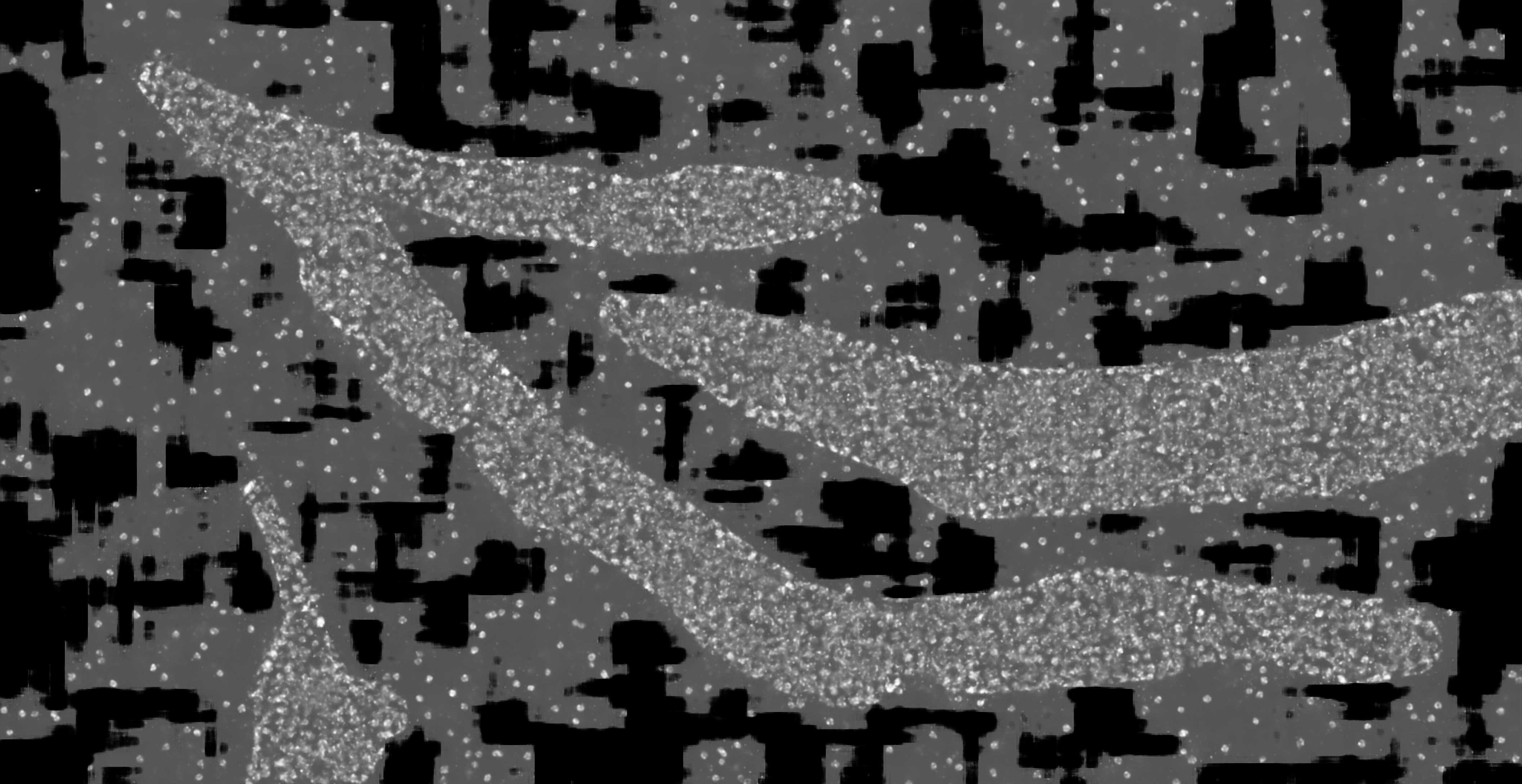

### Supp_Fig_9d_unet4.jpg

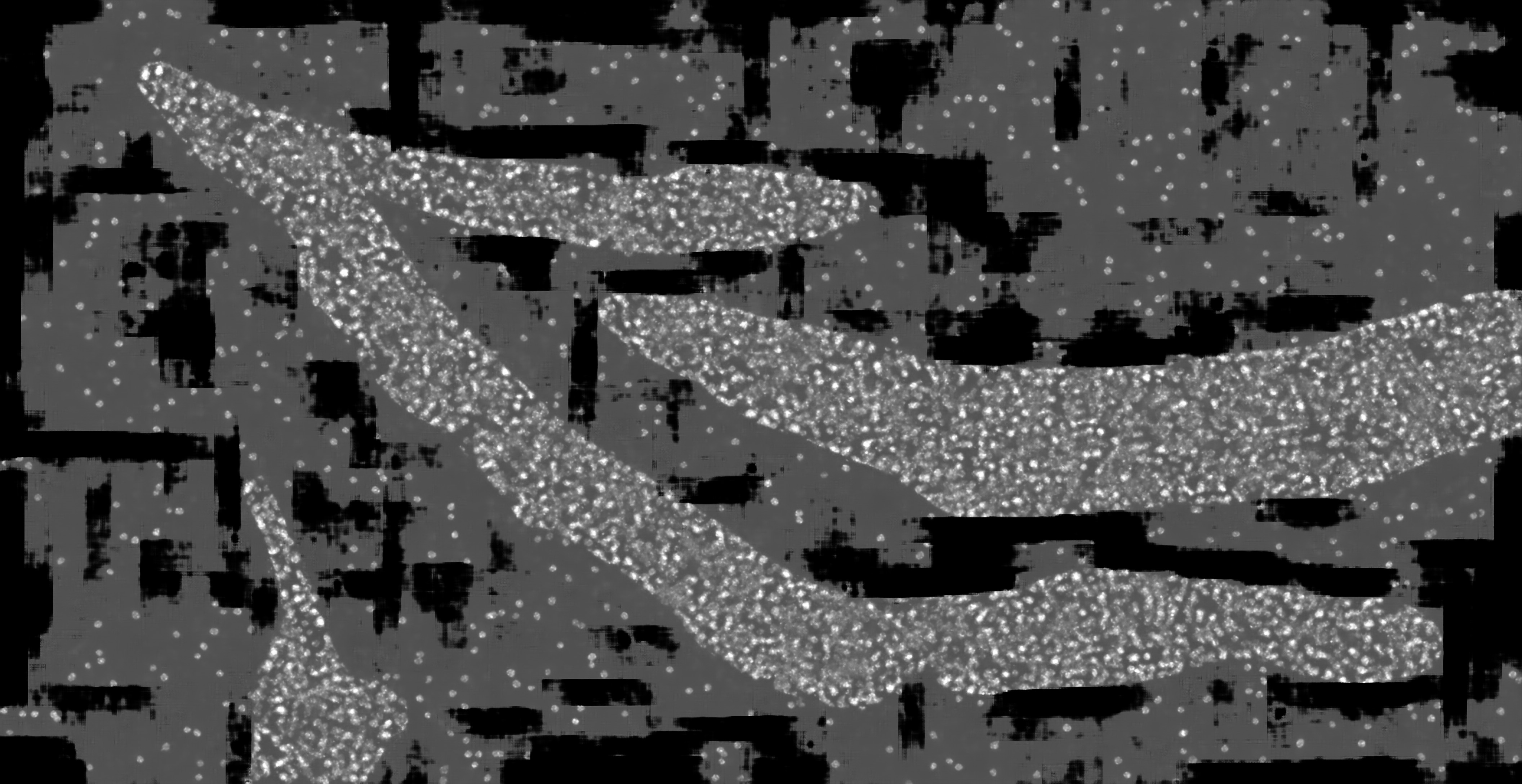
