## Supplementary figures and images for "Using conditional Generative Adversarial Networks (GAN) to generate *de novo* synthetic cell nuclei for training machine learning-based image segmentation"

### hc_more_test_mask2dapi_001_px_FS_16_KS_3_ML_False_4.jpg

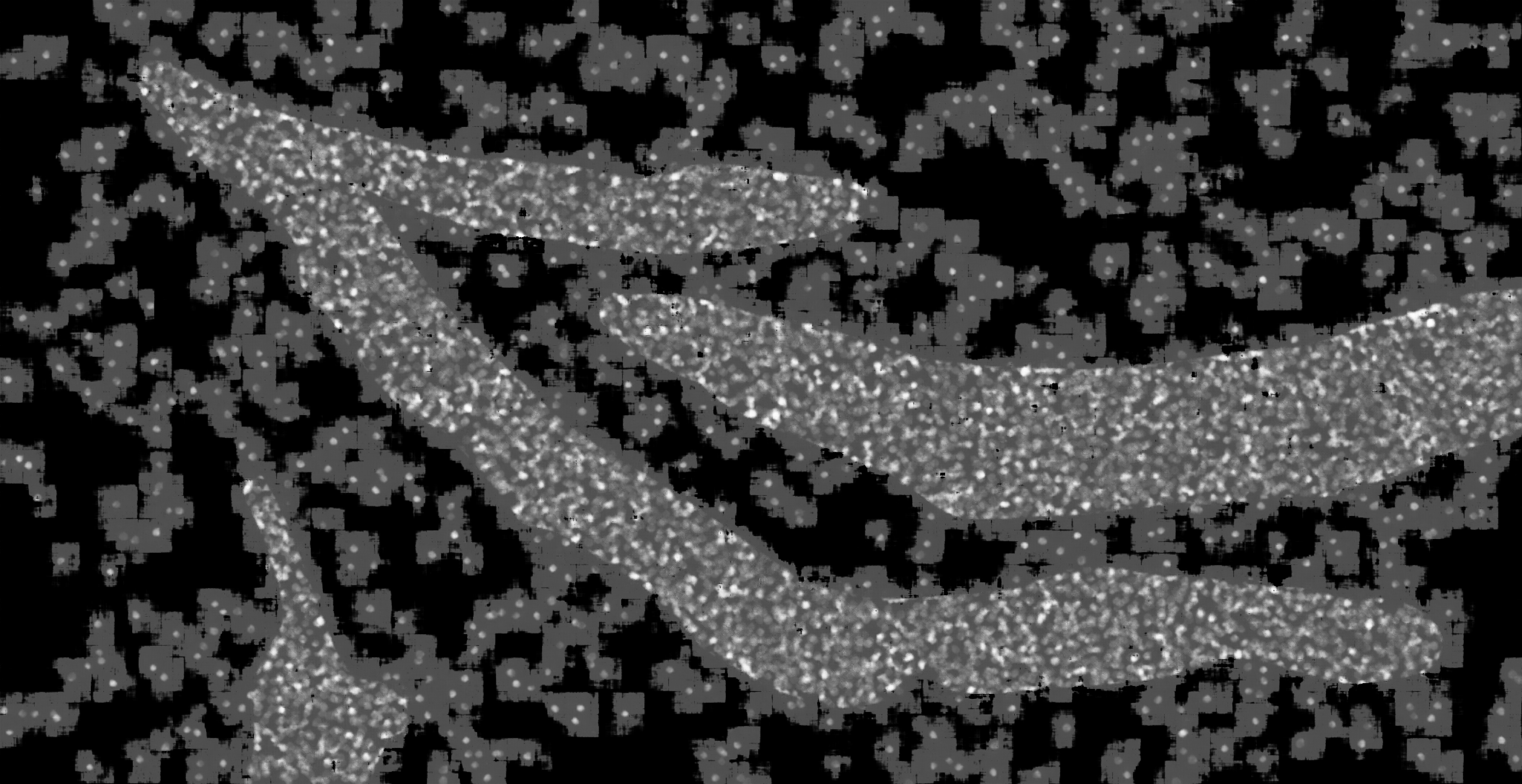

### hc_more_test_mask2dapi_001_px_FS_16_KS_5_ML_True_4.jpg

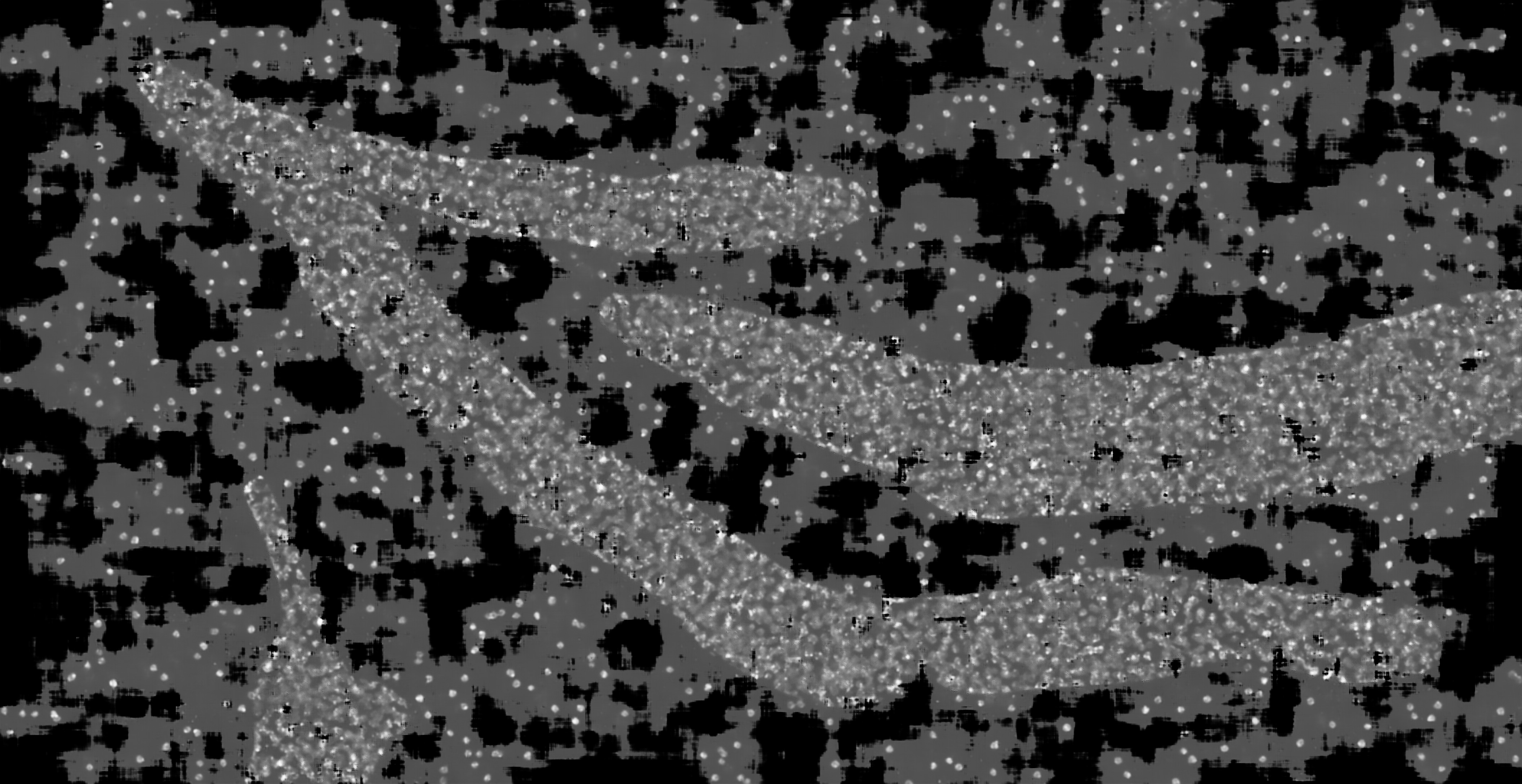

### hc_more_test_mask2dapi_001_px_FS_16_KS_7_ML_False_4.jpg

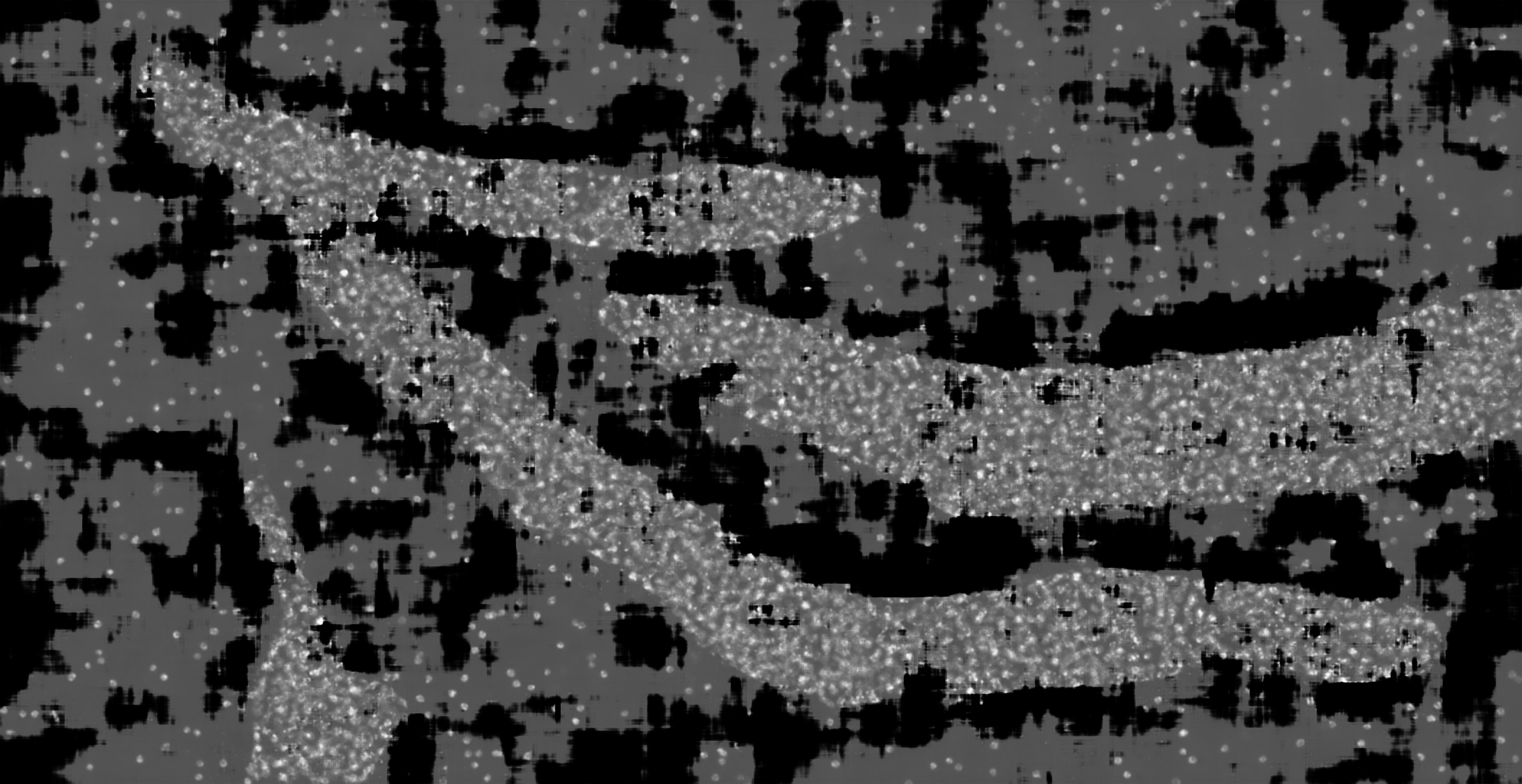

### hc_more_test_mask2dapi_001_px_FS_32_KS_3_ML_True_4.jpg

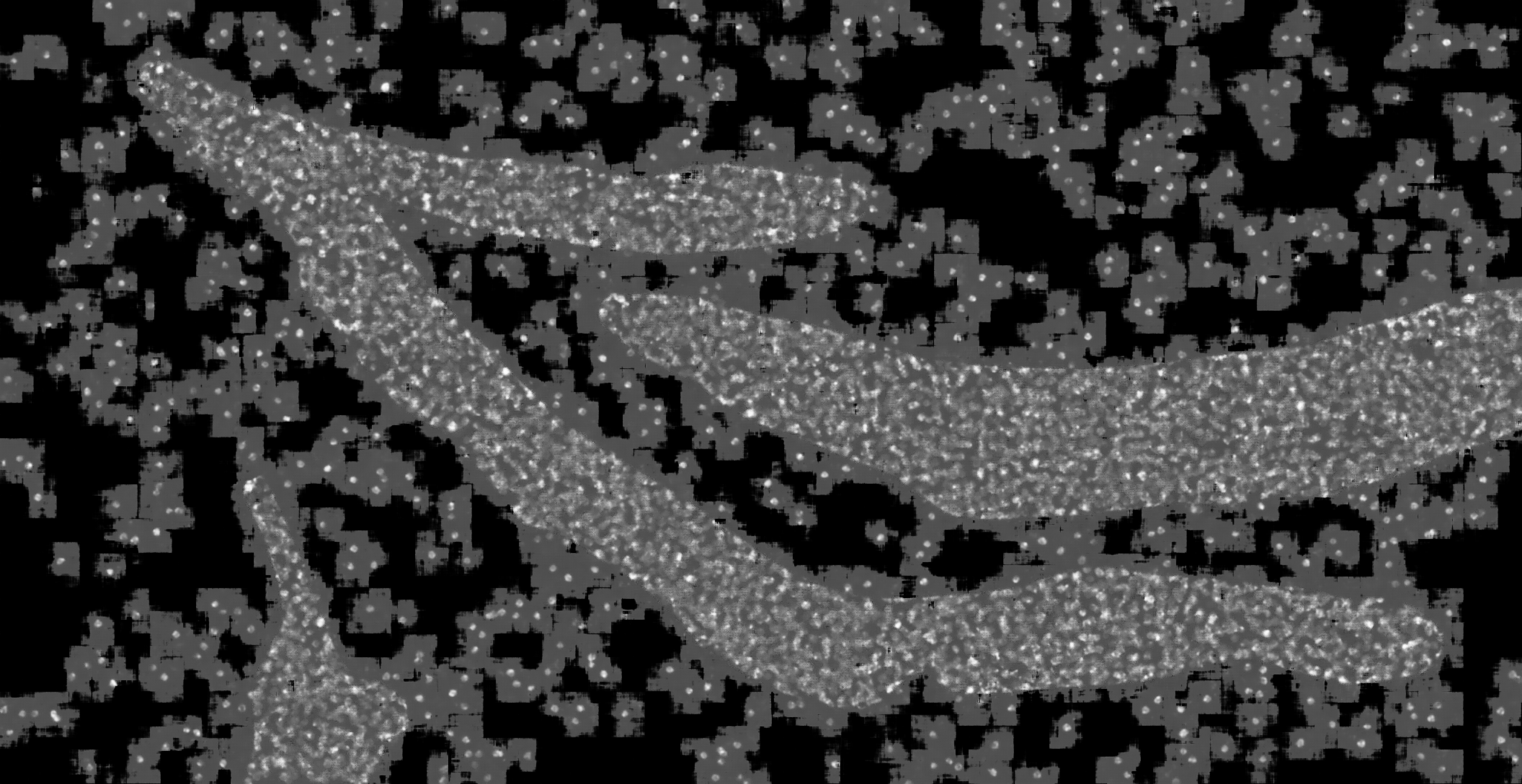

### hc_more_test_mask2dapi_001_px_FS_32_KS_7_ML_True_4.jpg

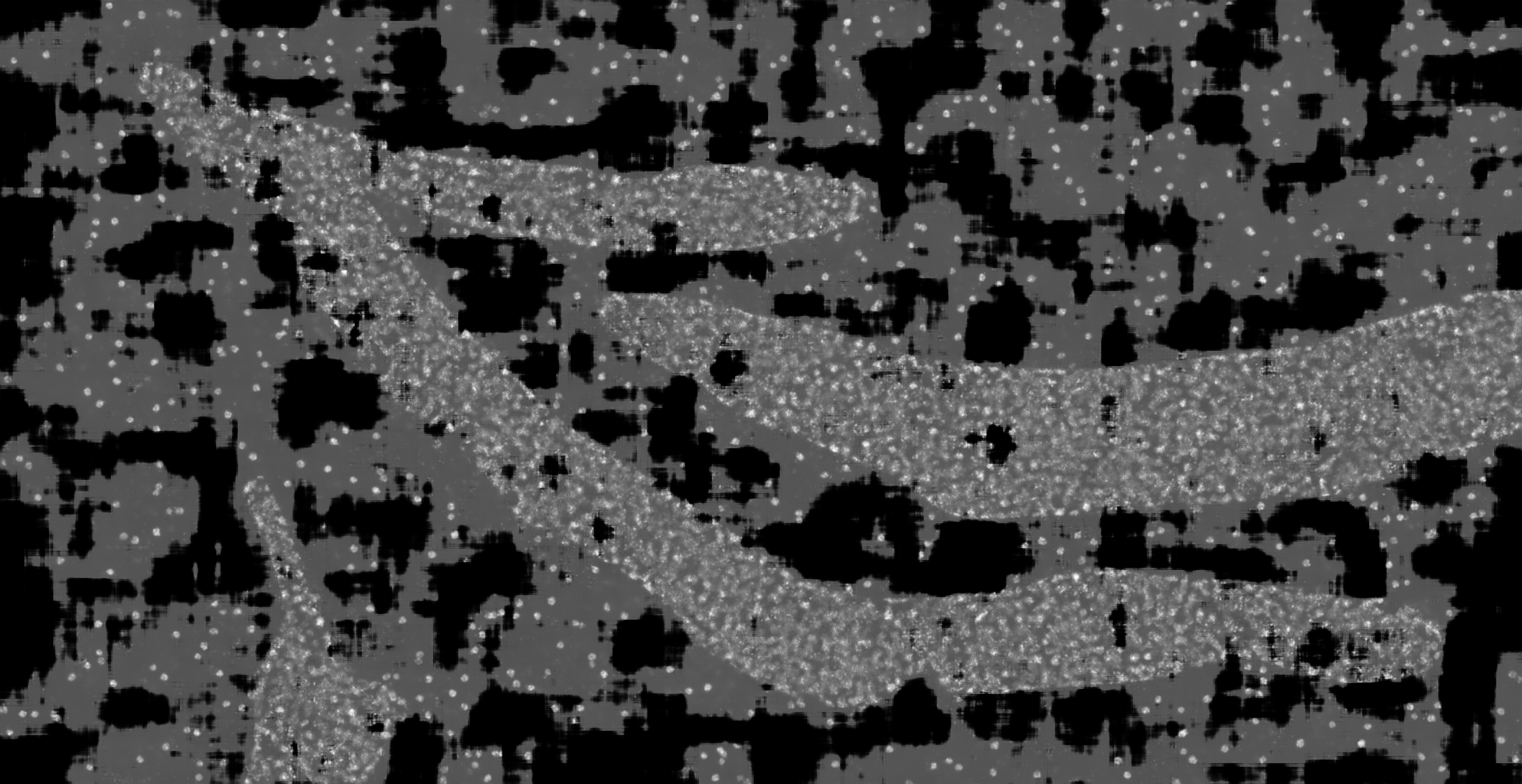

### hc_more_test_mask2dapi_002a_px_FS_16_KS_3_ML_True_4.jpg

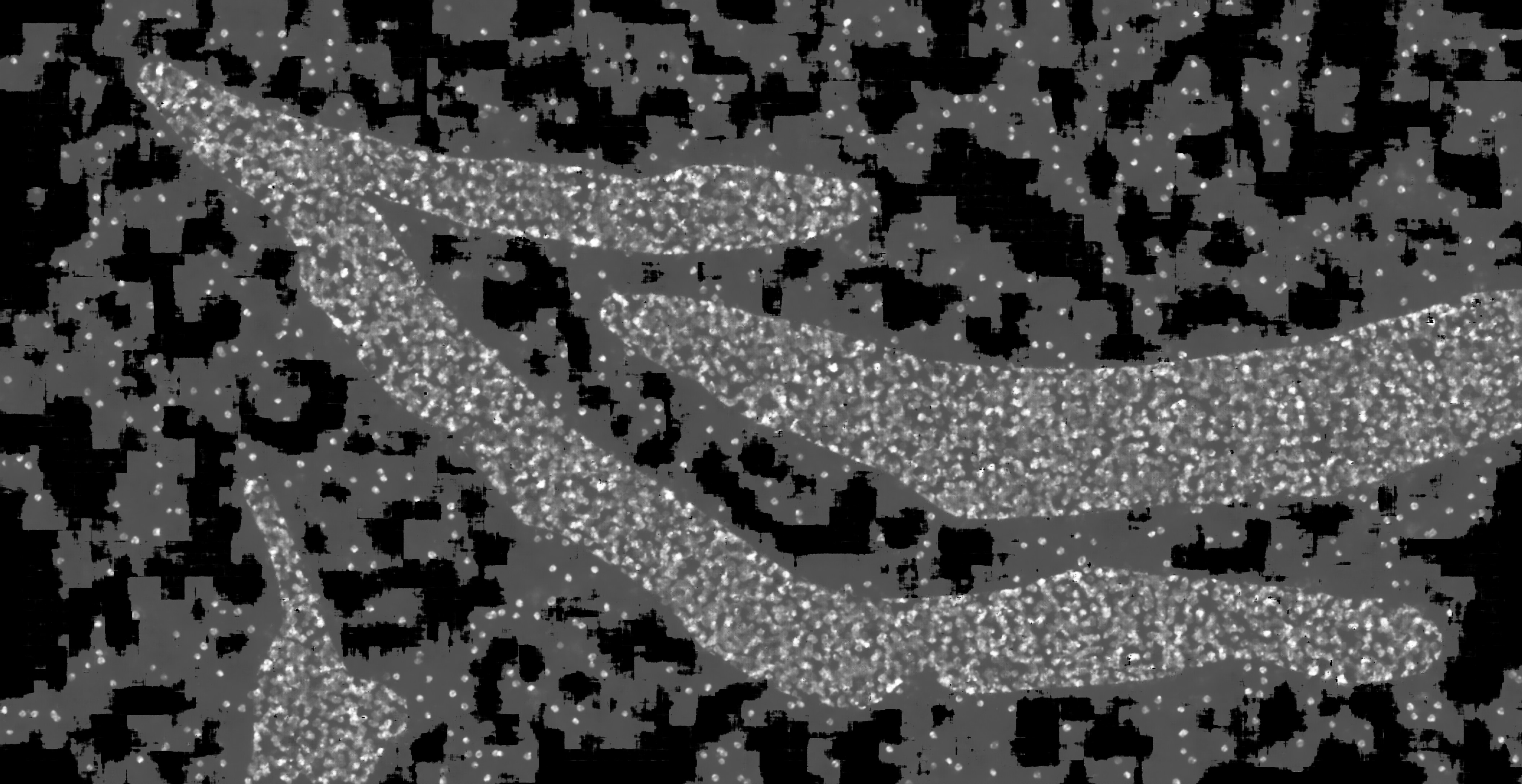

### hc_more_test_mask2dapi_002a_px_FS_16_KS_5_ML_False_4.jpg

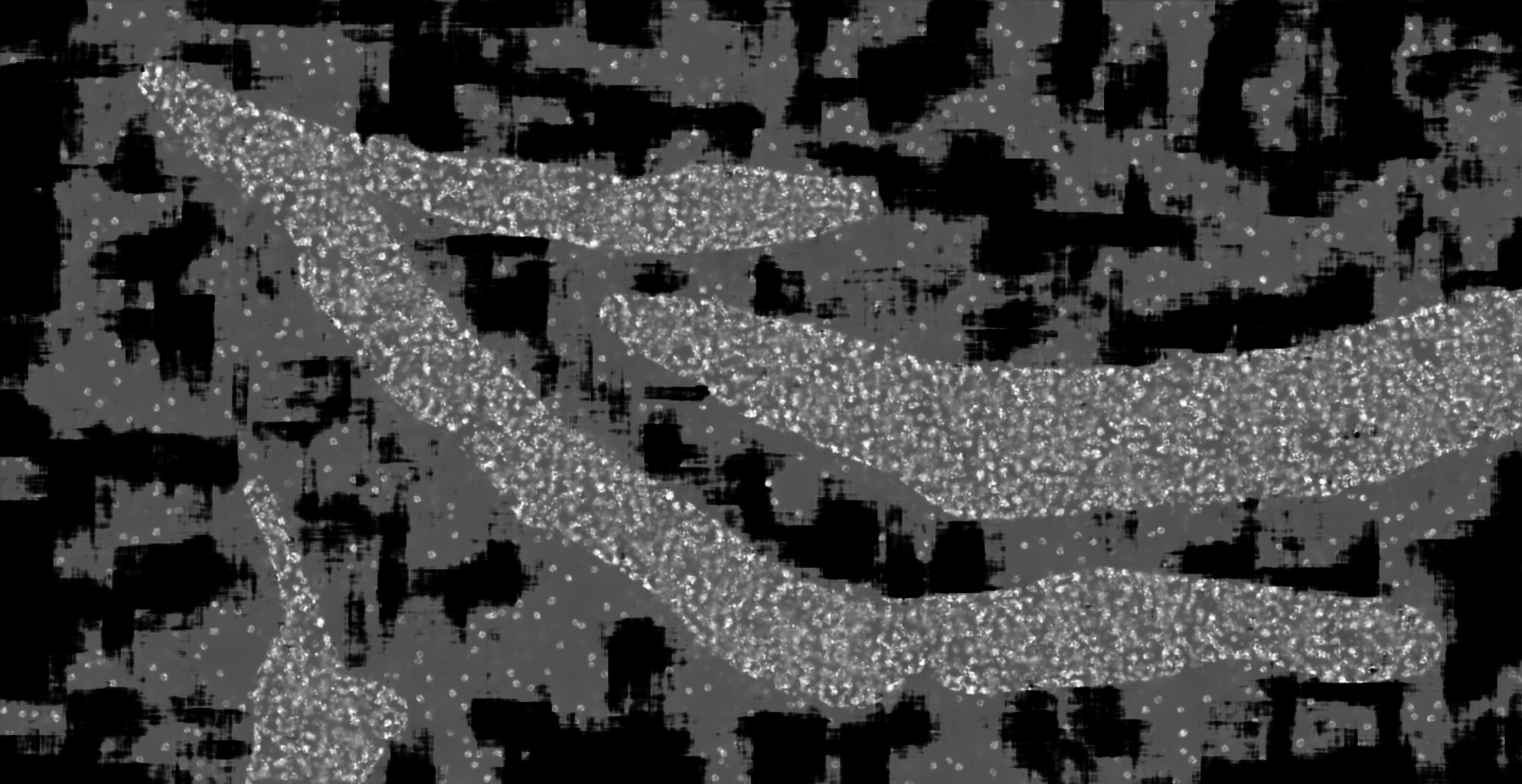

### hc_more_test_mask2dapi_002a_px_FS_16_KS_7_ML_False_4.jpg

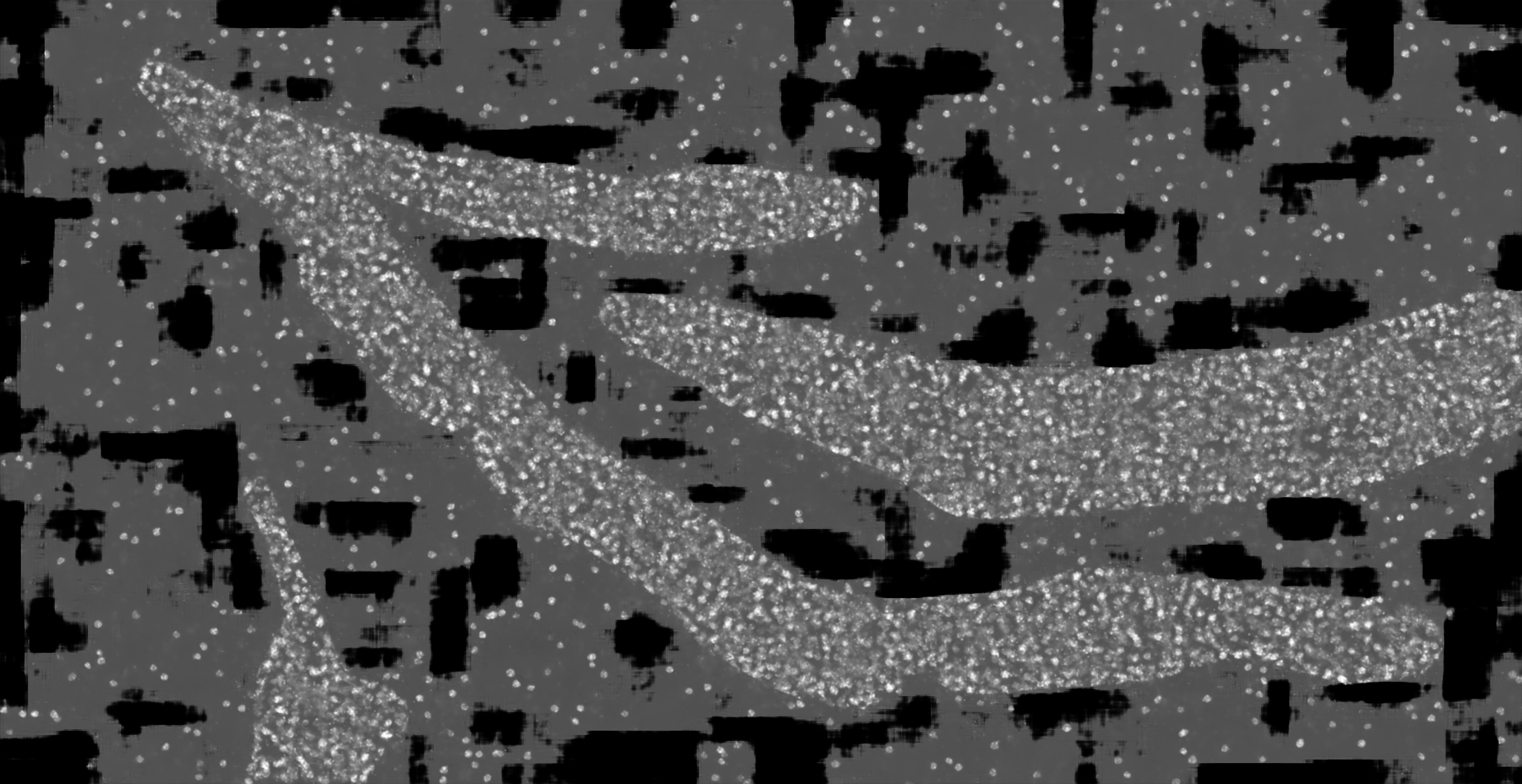

### hc_more_test_mask2dapi_002a_px_FS_16_KS_7_ML_True_4.jpg

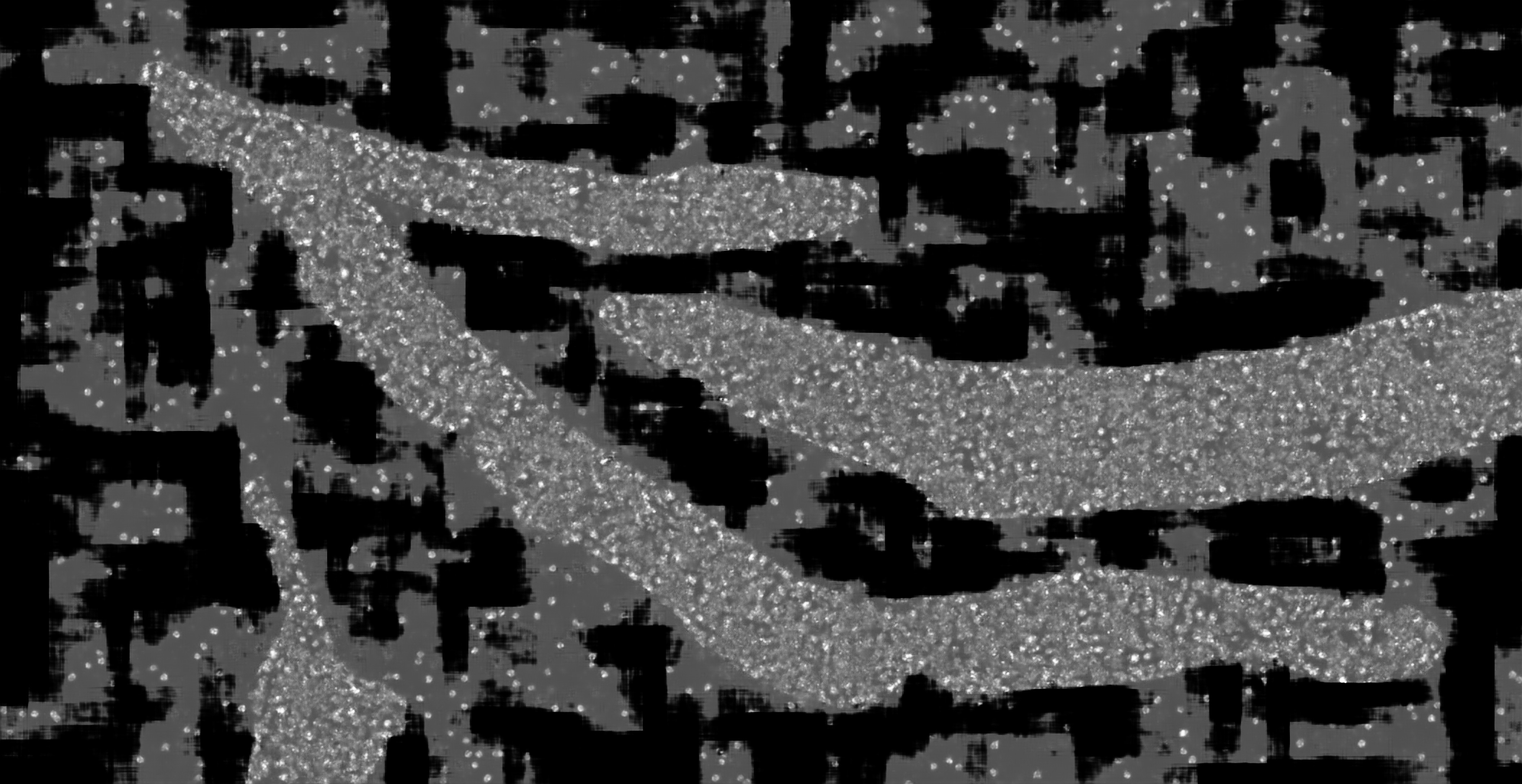

### hc_more_test_mask2dapi_002b_px_FS_16_KS_3_ML_False_4.jpg

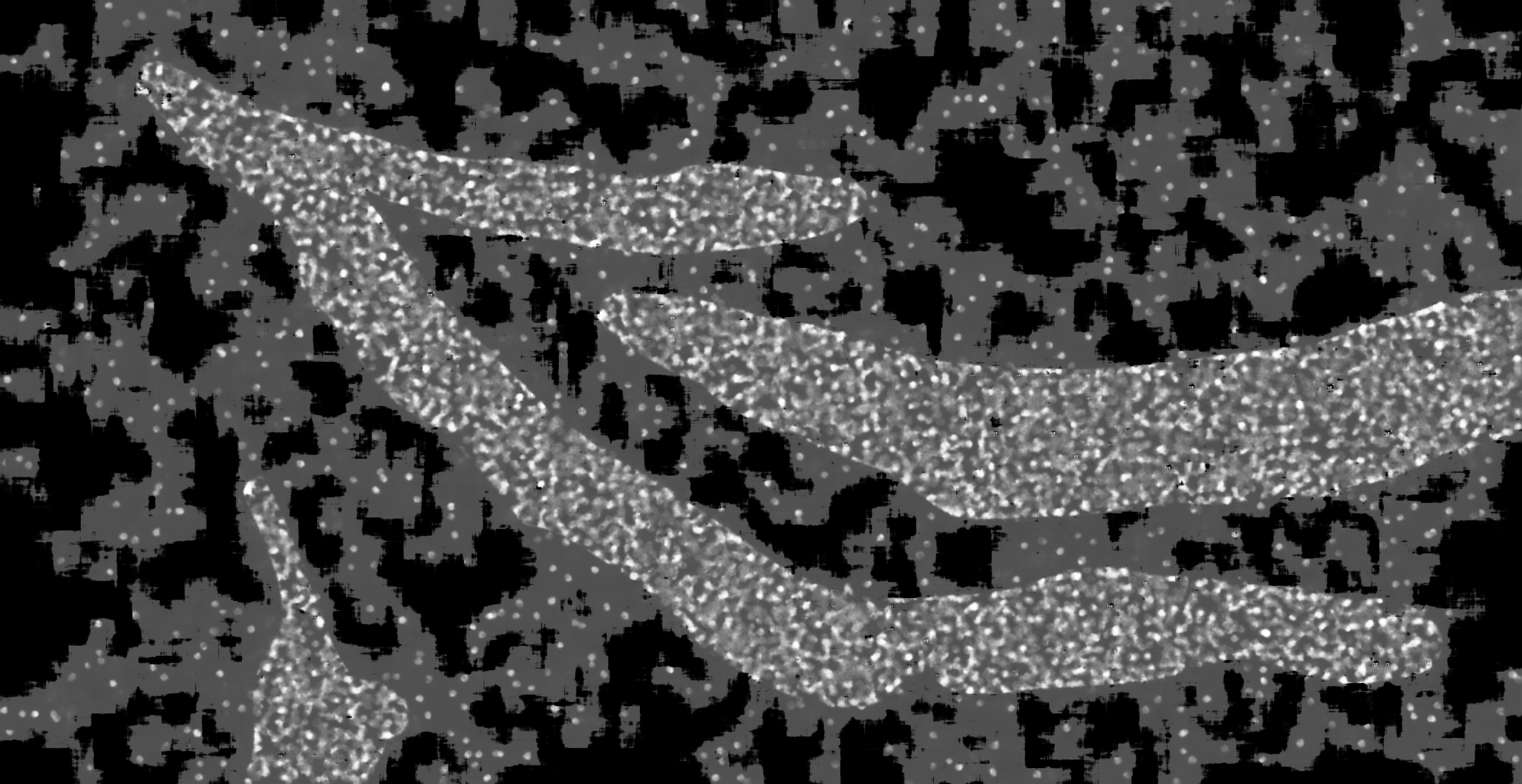

### hc_more_test_mask2dapi_002b_px_FS_16_KS_5_ML_True_4.jpg

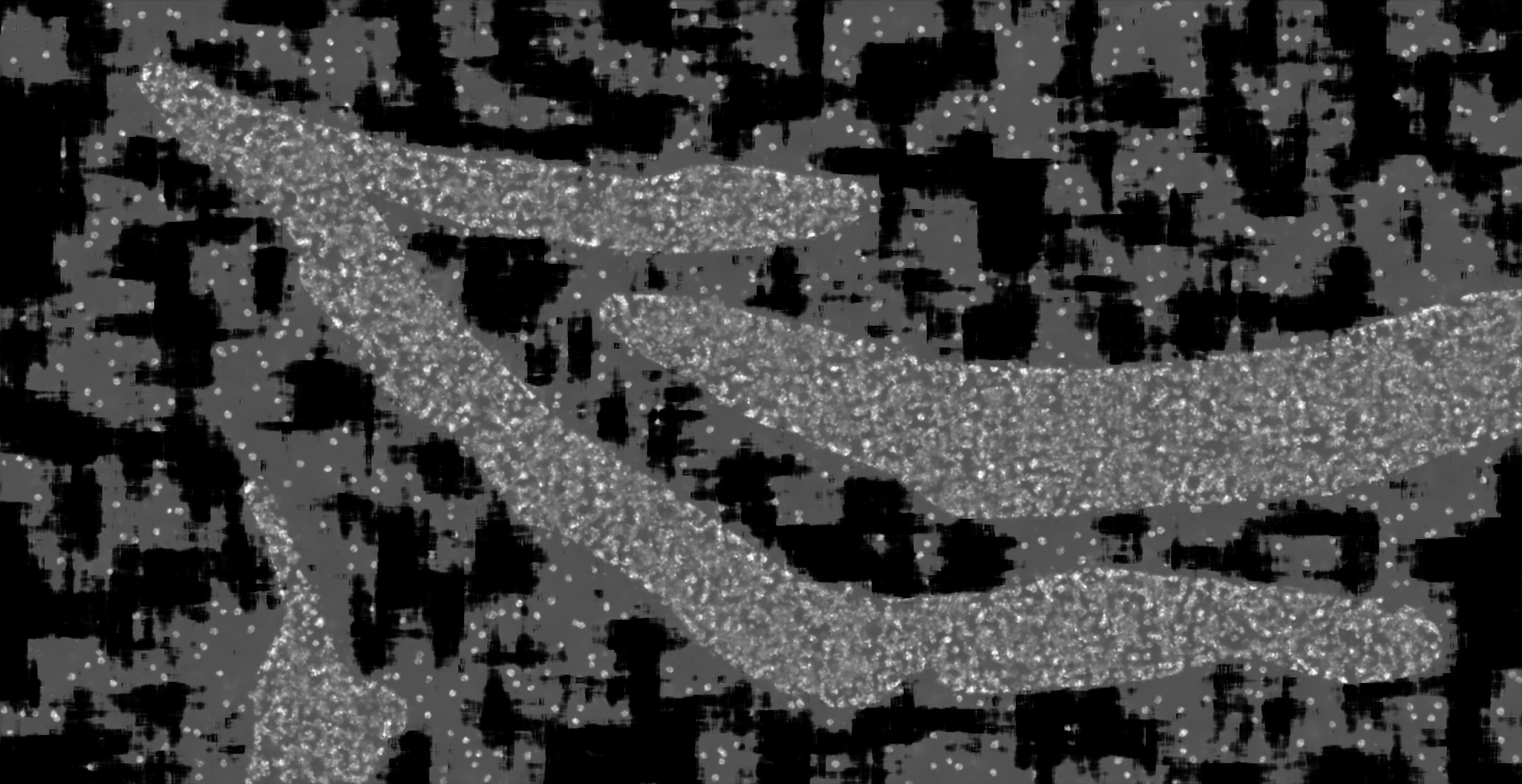

### hc_more_test_mask2dapi_002b_px_FS_16_KS_7_ML_False_4.jpg

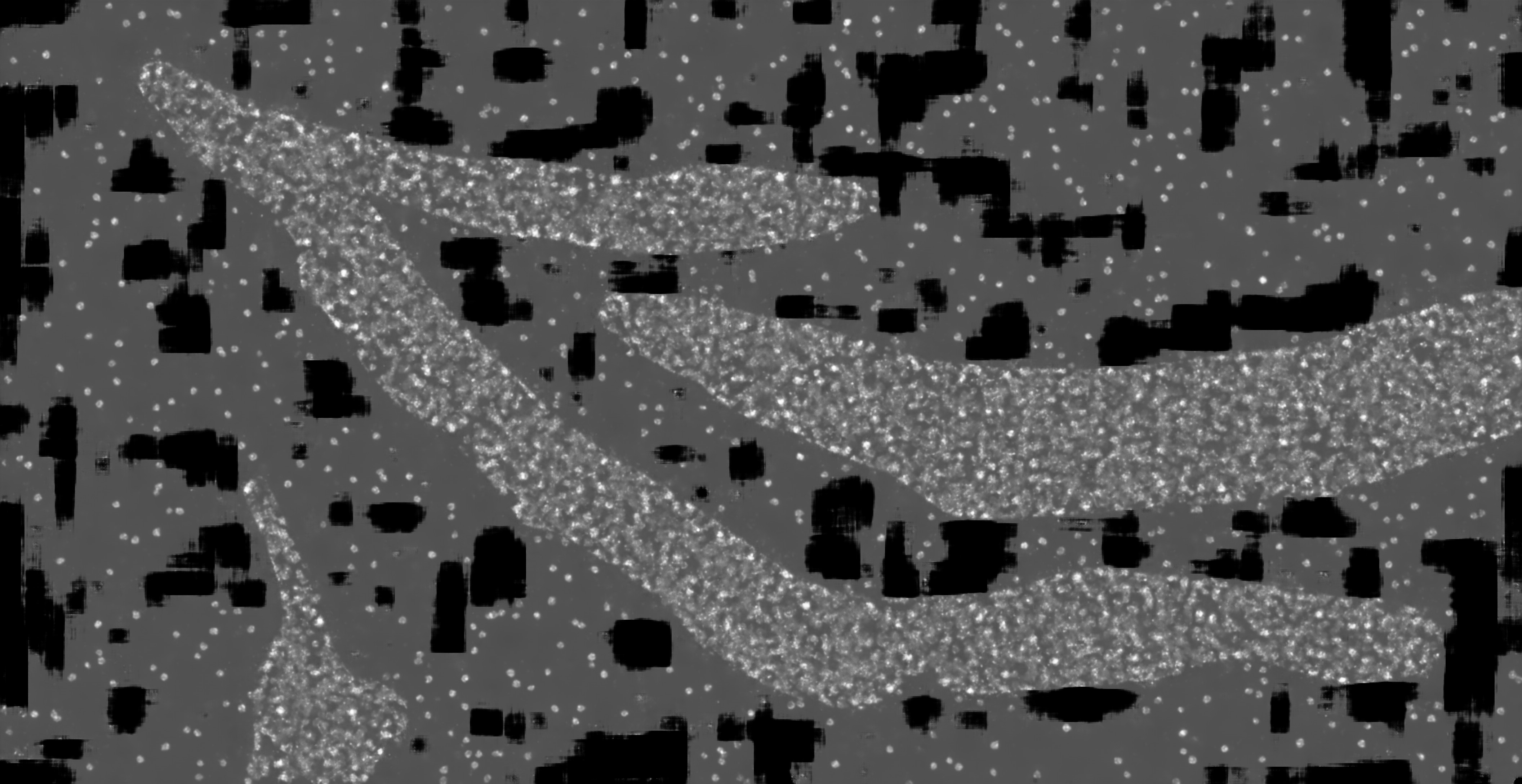

### hc_more_test_mask2dapi_003a_px_FS_16_KS_3_ML_False_4.jpg

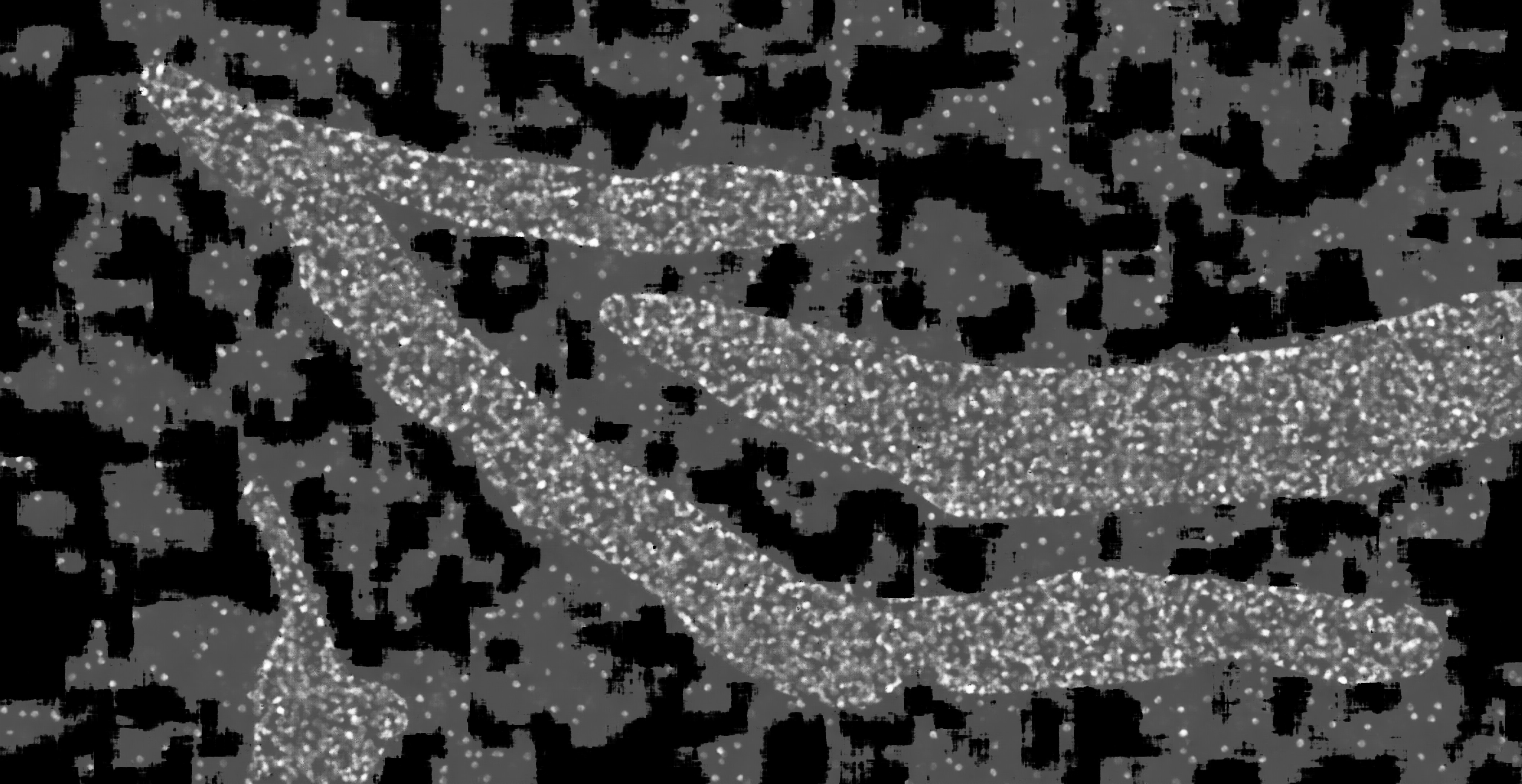

### hc_more_test_mask2dapi_003a_px_FS_16_KS_5_ML_False_4.jpg

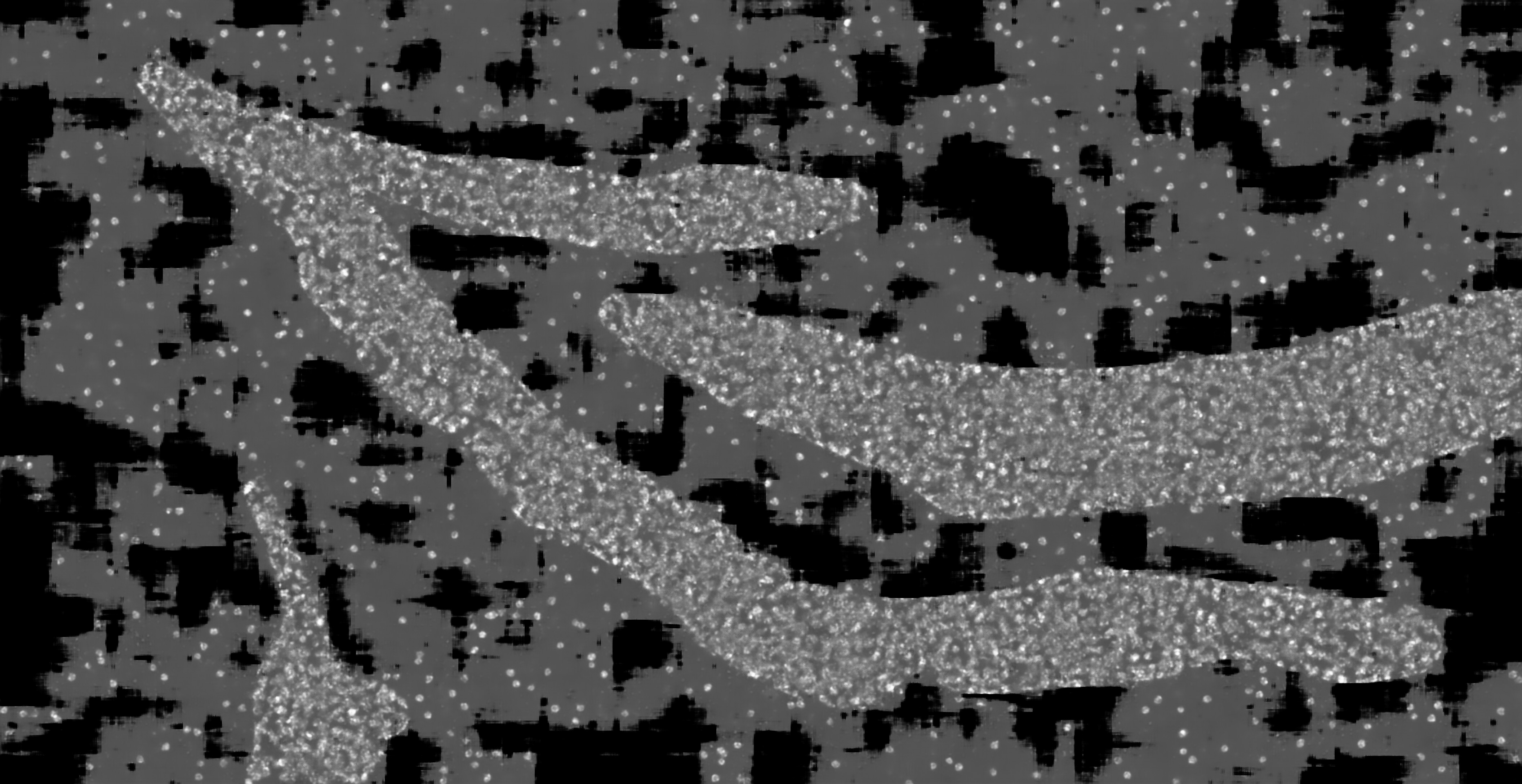

### hc_more_test_mask2dapi_003b_px_FS_16_KS_3_ML_False_4.jpg

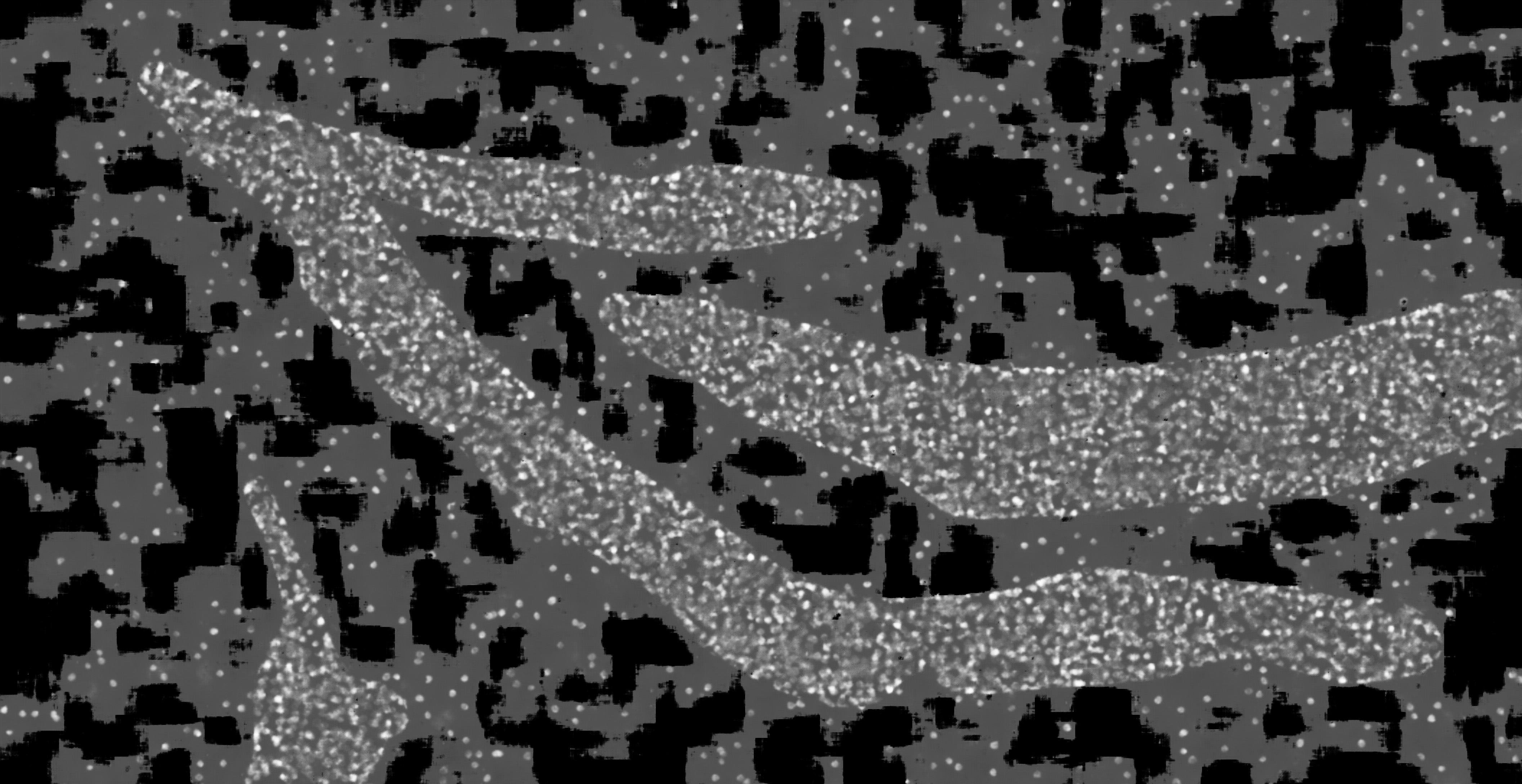

### hc_more_test_mask2dapi_003b_px_FS_16_KS_3_ML_True_4.jpg

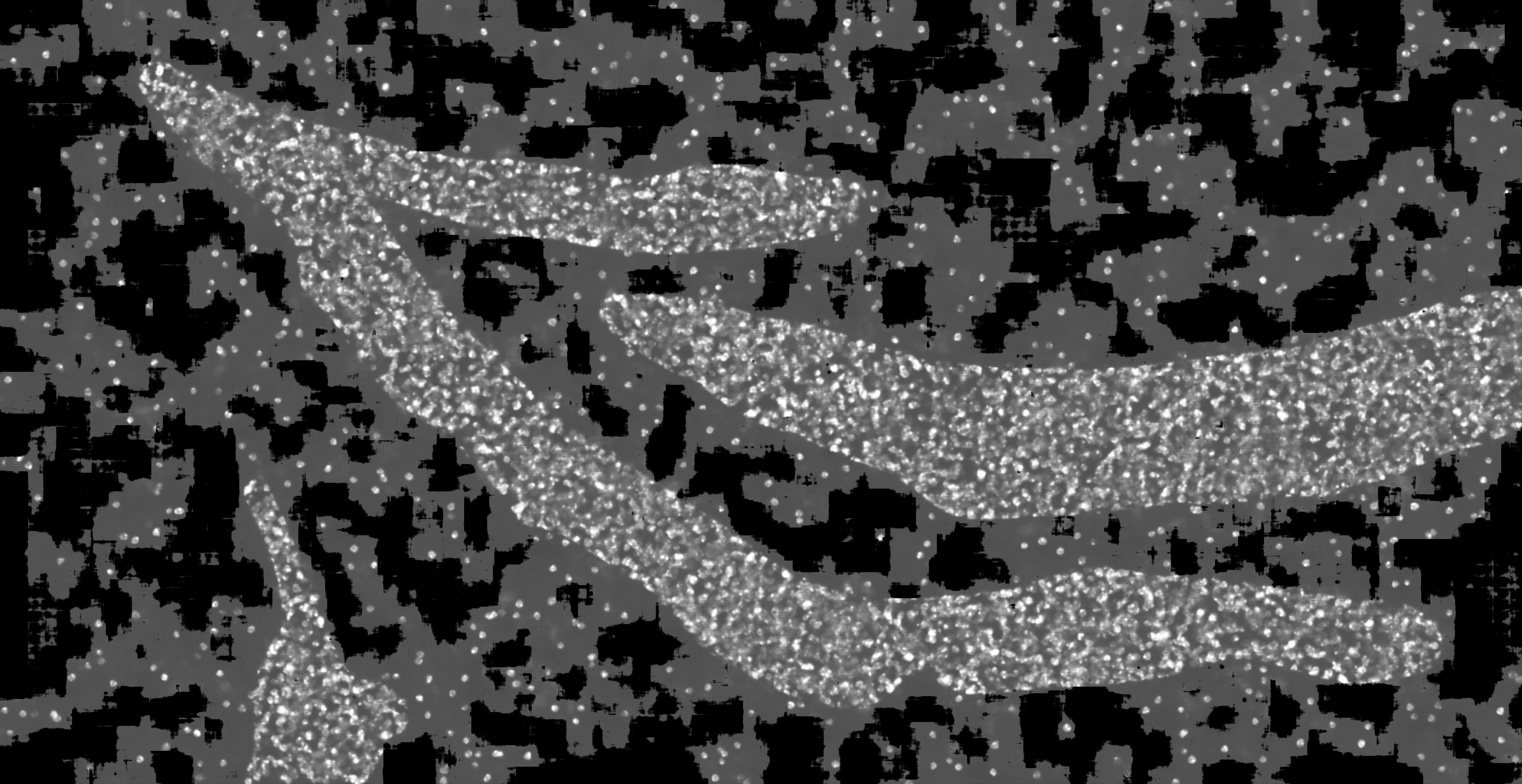

### hc_more_test_mask2dapi_003b_px_FS_16_KS_5_ML_False_4.jpg

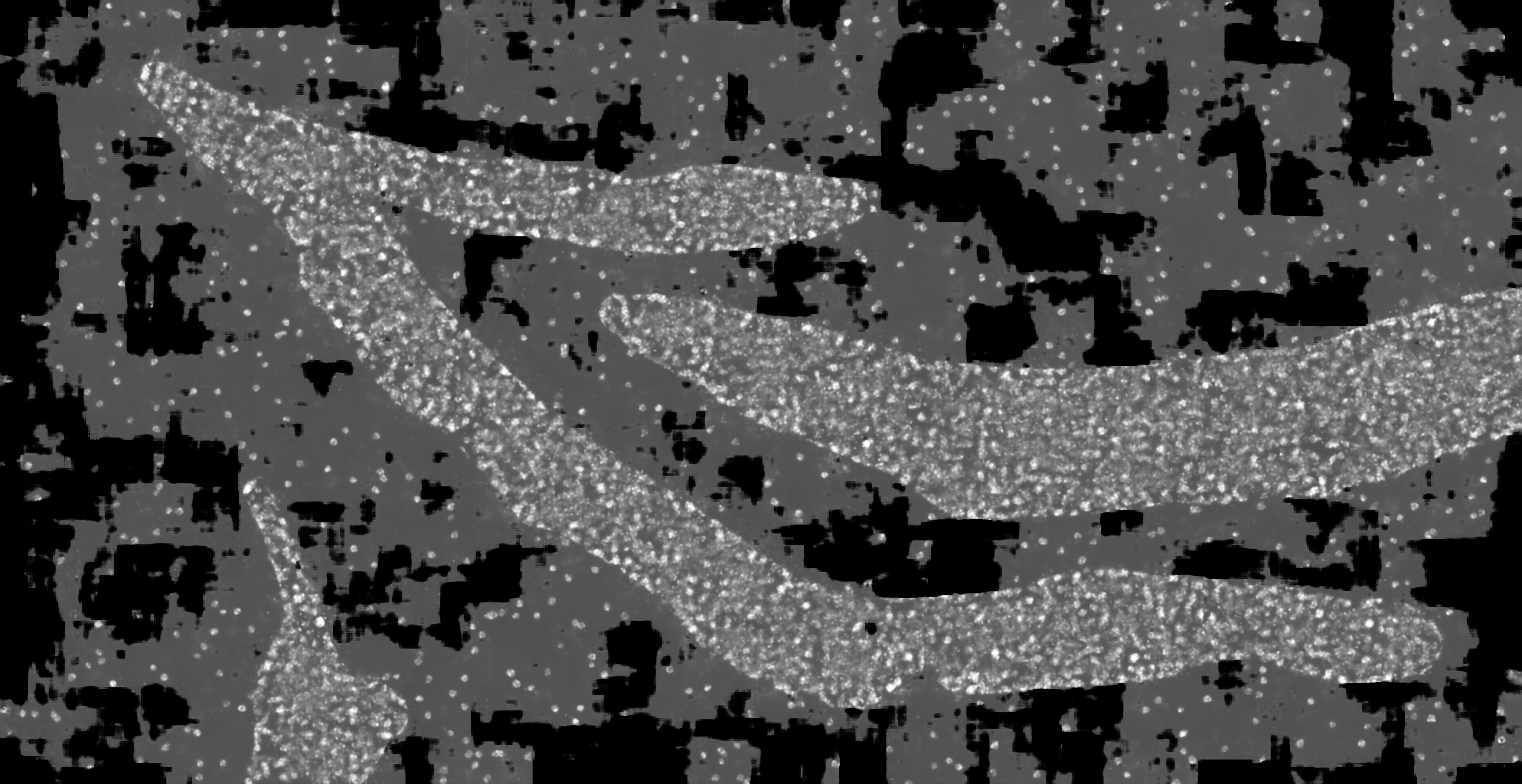

### hc_more_test_mask2dapi_003b_px_FS_16_KS_7_ML_False_4.jpg

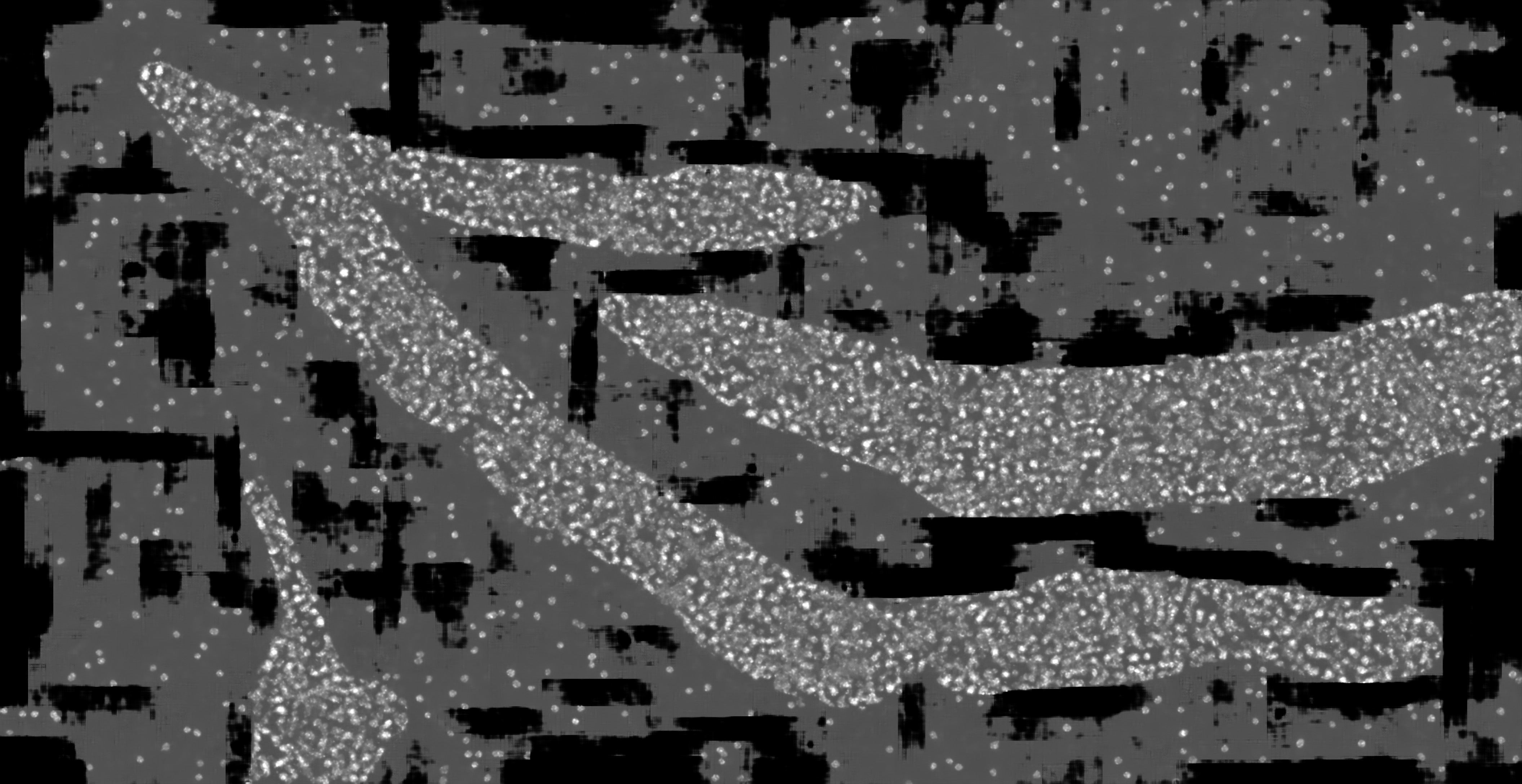

### hc_more_test_mask2dapi_003b_px_FS_16_KS_7_ML_True_4.jpg

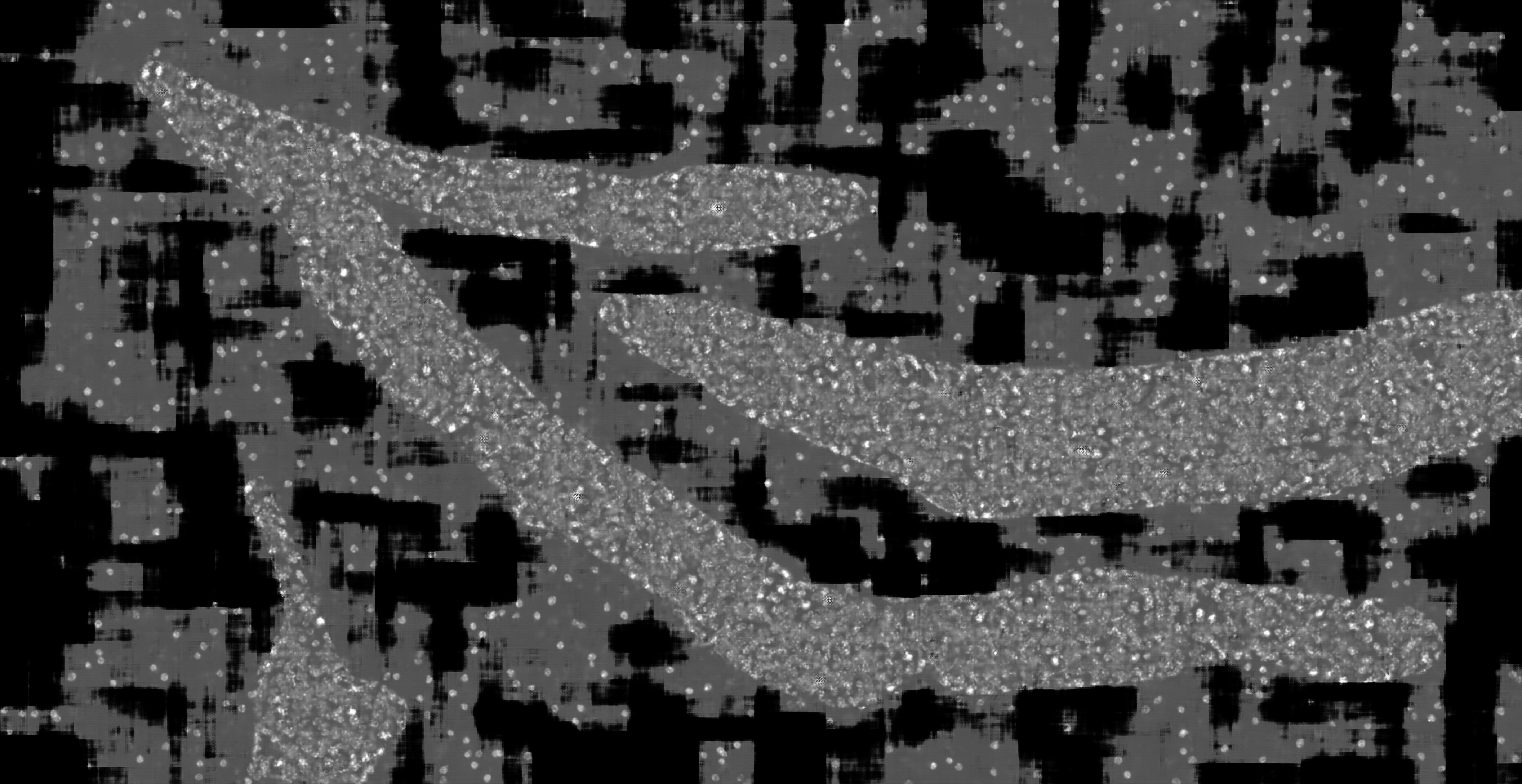
